# Discovery of novel 1,4-dicarbonylthiosemicarbazides as DNA gyrase inhibitors for the treatment of MRSA infection

**DOI:** 10.1101/2024.09.05.611349

**Authors:** Gao Zhang, Jiaxin Liang, Gang Wen, Mingli Yao, Yuqing Jia, Bo Feng, Jishun Li, Zunsheng Han, Qingxin Liu, Tianlei Li, Wenxuan Zhang, Hongwei Jin, Jie Xia, Liang Peng, Song Wu

## Abstract

Antibiotic resistance has become a serious threat to public health, thus novel antibiotics are urgently needed to combat drug-resistant bacteria including MRSA (methicillin-resistant *S. aureus*). The 1,4-dicarbonylthiosemicarbazide is an interesting chemotype that could exhibit antibacterial activity. However, the currently available compounds are not as potent as clinical antibiotics. Herein, we adopted the computer-aided drug design strategy, substructure search, to retrieve antibacterial 1,4-dicarbonylthiosemicarbazide derivatives, and identified compound **B5** (Specs ID: AG-690/15432331) from the Specs chemical library that exhibited moderate activity (minimum inhibitory concentration (MIC): 6.25 µg/mL) against *S. aureus* (ATCC 29213). Based on that compound, we further designed and synthesized 45 derivatives, and evaluated their antibacterial activity. Eight derivatives were more potent than or equivalent to vancomycin (MIC: 1.56 µg/mL). We compared the three most potent ones for their cytotoxicity to HepG2 and HUVEC cells and selected compound **1b** as our lead compound for comprehensive biological evaluation. As a result, compound **1b** exhibited a bacteriostatic mode, and was active against a panel of gram-positive bacteria strains, metabolically stable, and effective to protect the mice from MRSA infection. More importantly, we applied 2D similarity calculation and reverse docking to predict potential targets of compound **1b**. Through experimental validation and molecular dynamics simulation, we were able to confirm that compound **1b** inhibited *S. aureus* DNA gyrase (IC_50_: 1.81 µM) and DNA supercoiling, potentially by binding to the ATPase domain, where ASP81, GLU58 and GLN91 formed key hydrogen bonds. Taken together, we have discovered a new class of DNA gyrase inhibitors represented by compound **1b** for the treatment of MRSA infection, through the design, synthesis, and biological evaluation of novel 1,4-dicarbonylthiosemicarbazides.

**Graphic Abstract:** 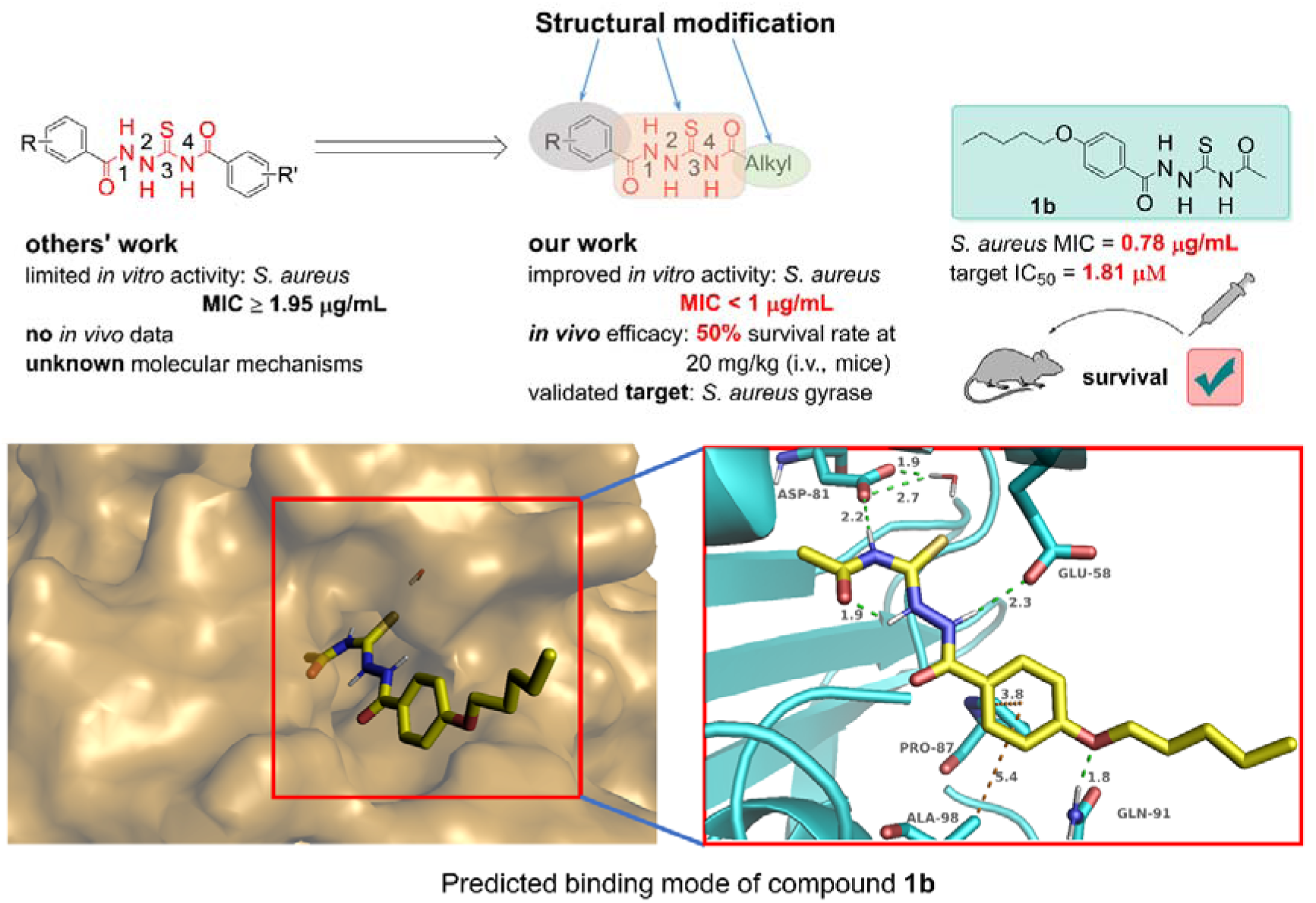

**Highlights:**

- An interesting 1,4-dicarbonylthiosemicarbazide derivative **B5** with moderate antibacterial activity for *S. aureus* was discovered by substructure search.
- Based on **B5**, 45 new 1,4-dicarbonylthiosemicarbazides were designed, synthesized and biologically evaluated, which led to the discovery of a promising lead compound **1b**.
- Compound **1b** exhibited good antibacterial activity for a panel of gram-positive bacteria strains, and was effective to protect mice from MRSA infection.
- The 1,4-dicarbonylthiosemicarbazides represented by compound **1b** were discovered as a new class of DNA gyrase inhibitors, by target prediction and experimental validation.

## 1. Introduction

Methicillin-resistant *S. aureus* (MRSA) is the leading bacterium responsible for nosocomial infections, and may cause bacteraemia, endocarditis, skin and soft tissue infections, bone and joint infections, and hospital-acquired infections [1,2]. In many places of the world, community-acquired MRSA infections tend to emerge rapidly [3]. Vancomycin is last-line treatment for most MRSA infections seen in hospitalized patients [4]. However, risk of treatment failure has already been evidenced by the presence of bacterial strains that develop intermediate or full resistance to vancomycin [5]. To address this dilemma, new antibiotics were approved by FDA against MRSA in the past 20 years, e.g., linozolid [6]. Unfortunately, linezolid resistance has also appeared, which may cause treatment failure for linezolid derivatives in nosocomial infection due to cross-resistance [7]. To prepare for the emerging of MRSA strains that are likely to be resistant to those recently approved chemotypes, discovery of new classes of anti-MRSA agents is necessary.

Semicarbazide is a biologically active moiety due to the presence of ureido unit (-NH-CO-NH-NH-), which acts as a pseudodipeptide mimic. Its thione form, thiosemicarbazide, is considered biologically important, as compounds with this scaffold may show antibacterial [8], anti-tubercular [9,10], anticancer [11], or antiviral [12] activities. As a subtype of thiosemicarbazides, 1,4-dicarbonylthiosemicarbazides were previously used as antibacterial agents for *S. aureus* (cf. **Fig. 1**). For instance, Ragavan *et al.* (2010) reported that compound **A1** showed moderate activity against *S. aureus* with minimum inhibitory concentration (MIC) value of 12.5 μg/mL [13]. Dunman *et al.* (2012) disclosed a patent, in which compound **A2** showed MIC of 16 μg/mL against MRSA. Although compound **A2** was proposed as an inhibitor of *S. aureus* ribonuclease-P-protein subunit mediated RNA degradation process, its activity in terms of half-maximal inhibitory concentration (IC_50_) was greater than 50 μM, implying there may be other molecular mechanisms related to its antibacterial activity [14]. Chudzik-Rząd and co-workers (2020) discovered compound **A3** that exhibited activity against *S. aureus* with MIC of 1.95 μg/mL, and MRSA with MIC of > 62.5 μg/mL. For molecular-level antibacterial mechanism, they tested activities of compound **A3** against *S. aureus* bacterial gyrase and topoisomerase IV with the supercoiling and decatenation assays, but compound **A3** showed no inhibition for these two enzymes (IC_50_ > 100 μM) [15]. Ameryckx and co-workers (2020) reported that compound **A4** with 2-hydroxybenzoyl group at position 1 and 3,4-dichlorobenzoyl group at position 4 is moderately active against *S. aureus*, with MIC of 9.6 μg/mL. They tested a series of derivatives containing this scaffold for their inhibition of D-alanyl-D-alanine ligase (Ddl), however, the Ddl inhibition of this type (i.e., the 1,4-dicarbonylthiosemicarbazides) was less than 50% at 50 μM [16]. Kowalczyk *et al.* reported that compound **A5** with 1*H*-indole-2-carbonyl group at position 1 and benzoyl group at position 4 exhibited moderate activity against *S. aureus*, with MIC values ranging from 8 to 32 μg/mL for a panel of strains. They determined the IC_50_ values of its derivatives for *S. aureus* gyrase, and DNA topoisomerase IV as > 50 μM, and > 10 μM, respectively [17]. To the best of our knowledge, (1) the currently available 1,4-dicarbonylthiosemicarbazides merely showed moderate antibacterial activity (MIC: > 1 μg/mL). (2) there is no systematic structure-activity relationship (SAR) exploration to uncover the effect of substitutions at two sides of the 1,4-dicarbonylthiosemicarbazide scaffold. (3) no *in vivo* data for anti-MRSA efficacy was reported. (4) more importantly, the protein target of this chemical series has not been truly uncovered.

**Fig. 1.**
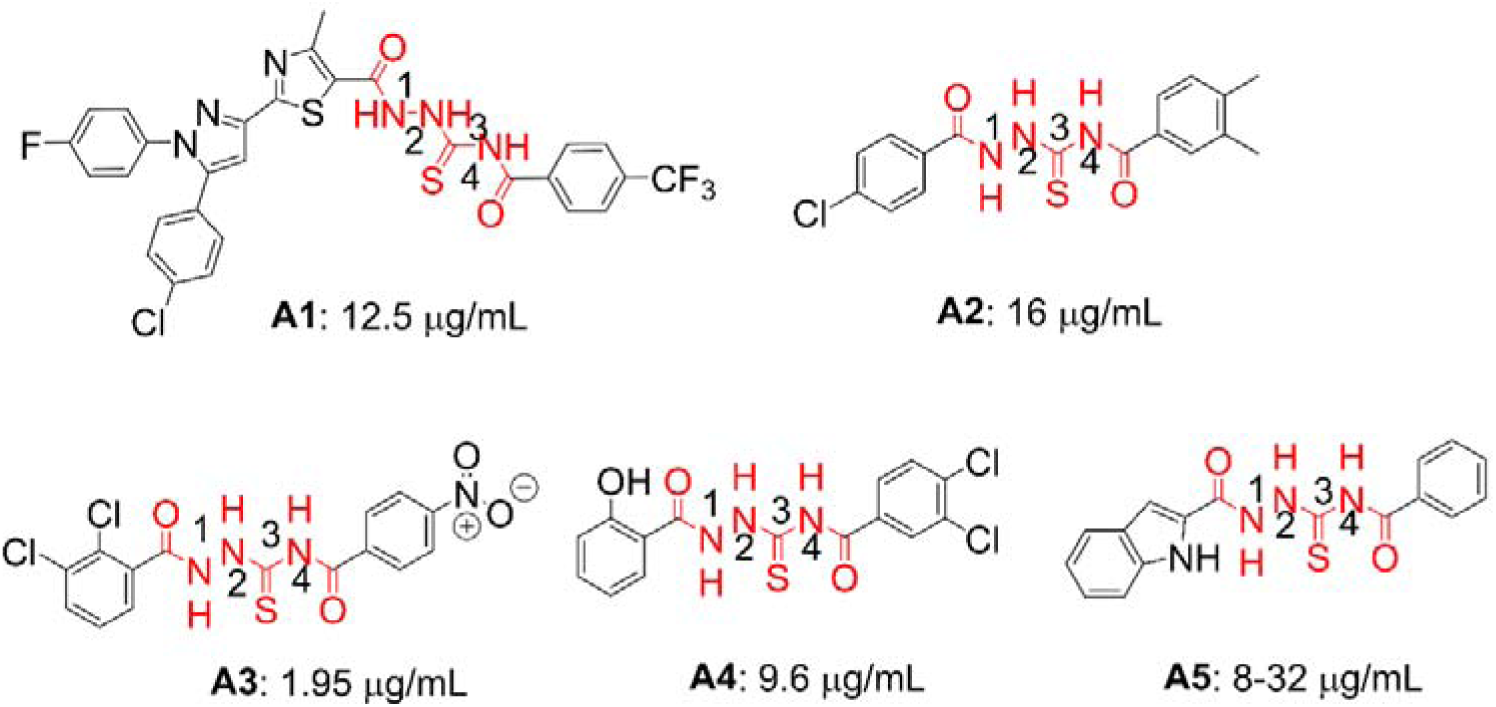
Representative small-molecule antibacterial agents containing the 1,4-dicarbonylthiosemicarbazide moiety.

In this study, we aim to address the issues listed above, by SAR exploration and biological evaluation of 1,4-dicarbonylthiosemicarbazides. One unique substructure (i.e., with 1-arylcarbonyl substituent and 4-alkylcarbonyl substituent at two different sides), which was different from the published structures (i.e., arylcarbonyl substituents at both sides), caught our attention. To be specific, we used 1,4-dicarbonylthiosemicarbazide core moiety for substructure search to identify diverse compounds. One derivative, **B5** (Specs ID: AG-690/15432331), was found to be a moderately active compound. It was then selected as our hit compound for further structural optimization. We synthesized a total of 45 new 1,4-dicarbonylthiosemicarbazide derivatives and evaluated their activities against *S. aureus* (ATCC 29213). The most promising compound, **1b**, was selected for further biological evaluation, including *in vivo* efficacy studies. We then performed target prediction by the strategy of pairwise similarity calculation and reverse docking, and validated our prediction by enzymatic experiments. We identified DNA gyrase as the target for this chemical series. The binding mode of the compound to DNA gyrase was revealed by molecular dynamics simulation.

## 2. Results and discussion

### 2.1. Molecular Design

We carried out substructure search, a commonly used computer-aided drug design strategy, to identify diverse 1,4-dicarbonylthiosemicarbazide derivatives (cf. **Fig. 2A**). We found that 1,4-dicarbonylthiosemicarbazide derivatives with aryl substituents at both sides had been identified as antibacterial agents [18,19], however, the derivatives with an alkyl group on one side and an aryl group on the other side had not been explored in any of the previous studies. Herein, we focused on this type of derivative by selecting and purchasing eight representative compounds from the Specs chemical library (**B1–B8**, cf. **Fig. 2B**) that were commercially available at that time. By conducting a literature search using SciFinder (https://scifinder.cas.org), we confirmed that their antibacterial activity had never been reported before.

**Fig. 2.**
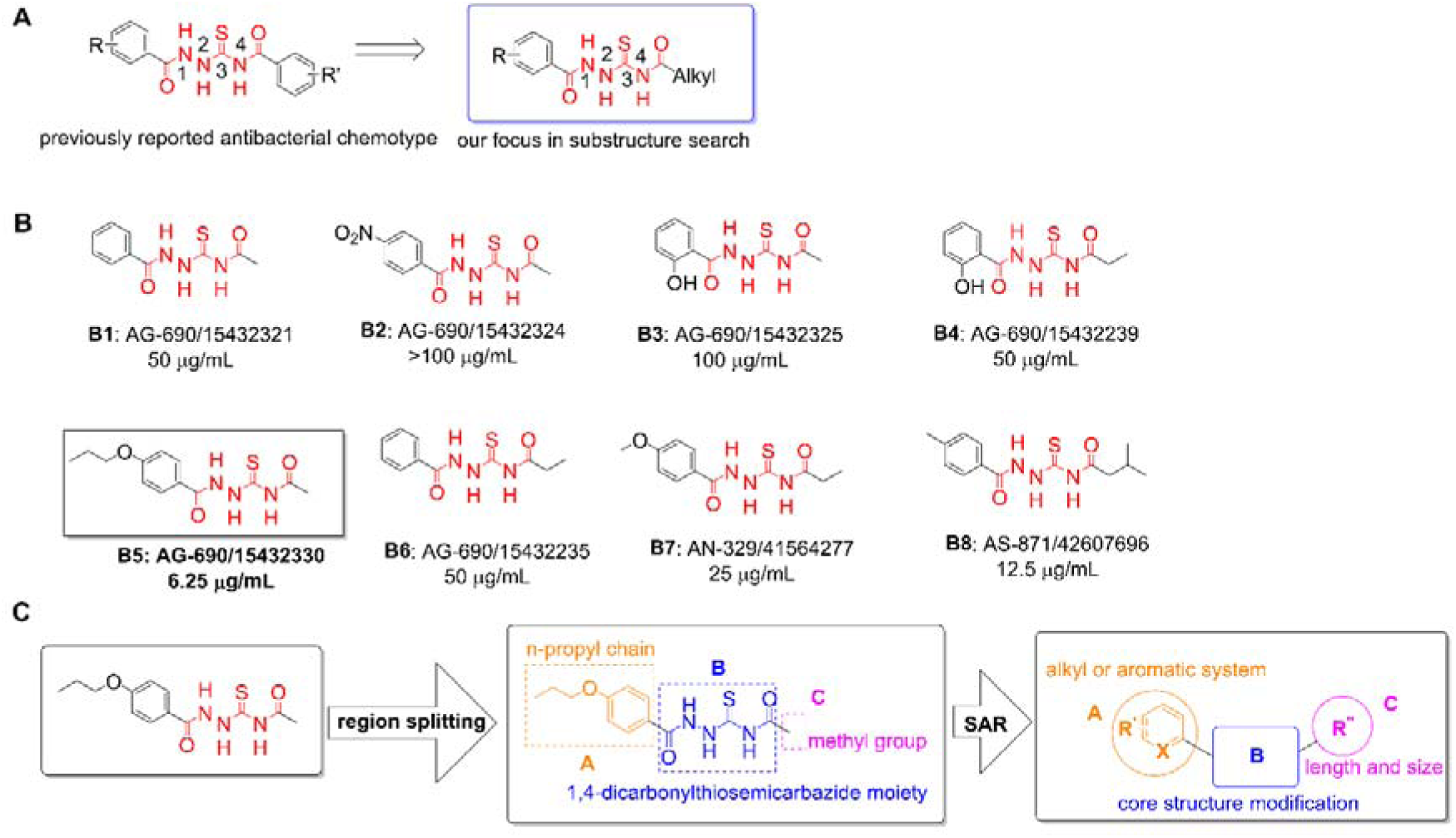
Substructure search-based virtual screening from the Specs chemical library for diverse 1,4-dicarbonylthiosemicarbazide derivatives and their antibacterial activities in terms of MIC. (A) The substructure search strategy. (B) Chemical structures and antibacterial activities (MICs) against *S. aureus* (ATCC 29213). The MIC of vancomycin (positive drug) was 1.56 μg/mL. (C) Three regions defined in molecular design and the outline of structure-antibacterial activity relationship study.

Then, their antibacterial activities in terms of MIC for *S. aureus* (ATCC 29213) were tested by the broth microdilution method [20] (cf. **Fig. 2B**). As the molecular mechanism/target was unknown for this type of antibacterial agent, the commonly used marketed drug, vancomycin, for gram-positive bacteria was selected as the positive drug. According to **Fig. 2B**, five compounds lacked antibacterial activity (MIC: ≥ 50 μg/mL), while three compounds showed moderate activity (6.25–25 μg/mL). Based on such data, a preliminary structure-activity relationship was concluded: (1) Electron-donating groups such as alkyl and alkoxy at *para-* position of the benzene ring were favorable for antibacterial activity (**B5**: 6.25 μg/mL, **B7**: 25 μg/mL, and **B8**: 12.5 μg/mL vs. **B2**: > 100 μg/mL); (2) The alkylcarbonyl group attached to position 4 had no significant effect on antibacterial activity as observed from **B1** (methylcarbonyl, 50 μg/mL) vs. **B6** (ethylcarbonyl, 50 μg/mL), as well as **B3** (methylcarbonyl, 50 μg/mL) vs. **B4** (ethylcarbonyl, 50 μg/mL). Among these derivatives, compound **B5** (Specs ID: AG-690/15432330) was the most potent (MIC: 6.25 μg/mL), whose activity value was close to vancomycin (MIC: 1.56 μg/mL).

As mentioned above, 1,4-dicarbonylthiosemicarbazides with alkylcarbonyl substituents at position 4 and arylcarbonyl groups at position 1 were not studied, in particular, for structure-antibacterial activity relationship. Thus, it was interesting to perform further modification on this class of antibacterial agents. According to the chemical features of compound **B5**, we divided it into three regions for structural modification (cf. **Fig. 2C**). As region A contained a *para*-substituted alkoxy group attached to the benzene ring, we firstly designed molecules with a variety of alkoxy groups and other substituents (e.g., phenyl, cyclohexyl). After the optimal group was determined, we designed molecules with substituents at region C such as alkyl groups, cycloalkyl groups, aryl groups, and polar carboxyl groups. To explore whether region B (the core scaffold, 1,4-dicarbonylthiosemicarbazide) was essential for antibacterial activity, we designed new molecules by replacing it with heterocycles such as 1,3,4-oxadiazole, 1,3,4-thiadiazole, or other thiosemicarbazide-containing groups.

### 2.2. Chemical synthesis

The 1,4-dicarbonylthiosemicarbazide derivatives were prepared according to previous publications [21–26]. The synthesis of key intermediates **b1–b13** is described in **Scheme 1**. The end products were synthesized according to the routes described in **Scheme 2–4**.

Generally, all the intermediates were prepared as follows (cf. **Scheme 1–2**). Intermediates **b1–b11** were obtained from the corresponding acyl chlorides **a1–a11**, with KSCN as the reactant. Since the acyl chlorides **a8** and **a11** were not commercially available, they were individually prepared from the corresponding acids. Acyl chloride **a12** as the starting material for the synthesis of **3d** was also obtained in this way. The intermediate **b12** was obtained from acetamide and oxalyl chloride. Intermediate **b13** was obtained from the following method: **a13** was prepared from methanesulfonamide in CS_2_, with KOH as the base, then, **a13** reacted with triphosgene undergoing elimination reaction to give **b13**. Phenols (**c1–c5, c8–c10**) were treated with iodoalkanes (or bromoalkanes), and K_2_CO_3_ in acetone under reflux, or carboxylic acid (**c6, c7, c11–c17**) were treated with concentrated sulfuric acid in methanol to yield ester intermediates (**d1–d24**). Hydrazine hydrate (80%) was then added to these esters in methanol under reflux to afford aryl hydrazines (**e1–e24**).

The majority of end products were prepared as follows (cf. **Scheme 2–3**). Aryl hydrazines (**e1–e24**) were added to the acetyl isothiocyanate **b1** in acetone under reflux for 6 h to give the products **1a–1x** (cf. **Scheme 2**). As shown in **Scheme 3**, aryl hydrazines **e2** were added to the corresponding acyl isothiocyanate **b2–b10** to give the products **2a–2i**, respectively. The derivatives with carboxyl groups (**2j**, **2k**) in region C were prepared from the corresponding ester derivatives (**2h**, **2i**) under basic condition, i.e., sodium hydroxide in methanol under reflux (cf. **Scheme 3**).

The derivatives containing variants of the 1,4-dicarbonylthiosemicarbazide moiety were readily obtained under different conditions (cf. **Scheme 4**). The 1,3,4-oxadiazole derivative (**3a**) was obtained from method A, in which product **1c** was treated with KIO_3_ and water under reflux. The 1,3,4-thiadiazole derivative (**3b)** was synthesized under microwave irradiation, i.e., **1c** was dissolved in acetic acid at 125 Cand under 300 W irradiation for 5 min. The methylation of nitrogen atoms at different positions (**3c** and **3d**) was achieved under different reaction conditions. The reduction of carbonyl groups (**3e** and **3f**) was achieved with lithium aluminum hydride as a reductant in the yields of 29% and 14%, respectively. Compounds **3g** or **3h** were prepared from acetyl isocyanate or methanesulfonyl isothiocyanate through nucleophilic attack by the corresponding hydrazine derivatives **e3** and **e2**, respectively. Compound **3i** was synthesized with benzoyl isothiocyanate **b11** and acetylhydrazine as reactants through the same nucleophilic addition reaction. Compound **3j** was synthesized from intermediate **e3** and *S*-methylisothiourea.

The three N-H protons of the 1,4-dicarbonylthiosemicarbazide moiety revealed peaks in ^1^H NMR spectra at δ 12.11, 11.57, and 10.89 ppm, respectively. Its two carbonyl signals and one thiocarbonyl signal were observed at δ 180.92, 172.52, and 164.67 ppm in ^13^C NMR spectra, respectively. The corresponding aromatic C–H protons of these compounds displayed in ^1^H NMR spectra as the signals with chemical shifts from δ 6.5 to 8.5 ppm. The aromatic carbon signals with chemical shifts ranged from δ 110 to 165 ppm. Based on this information, the structures of all the compounds were easily validated.

**Scheme 1.**
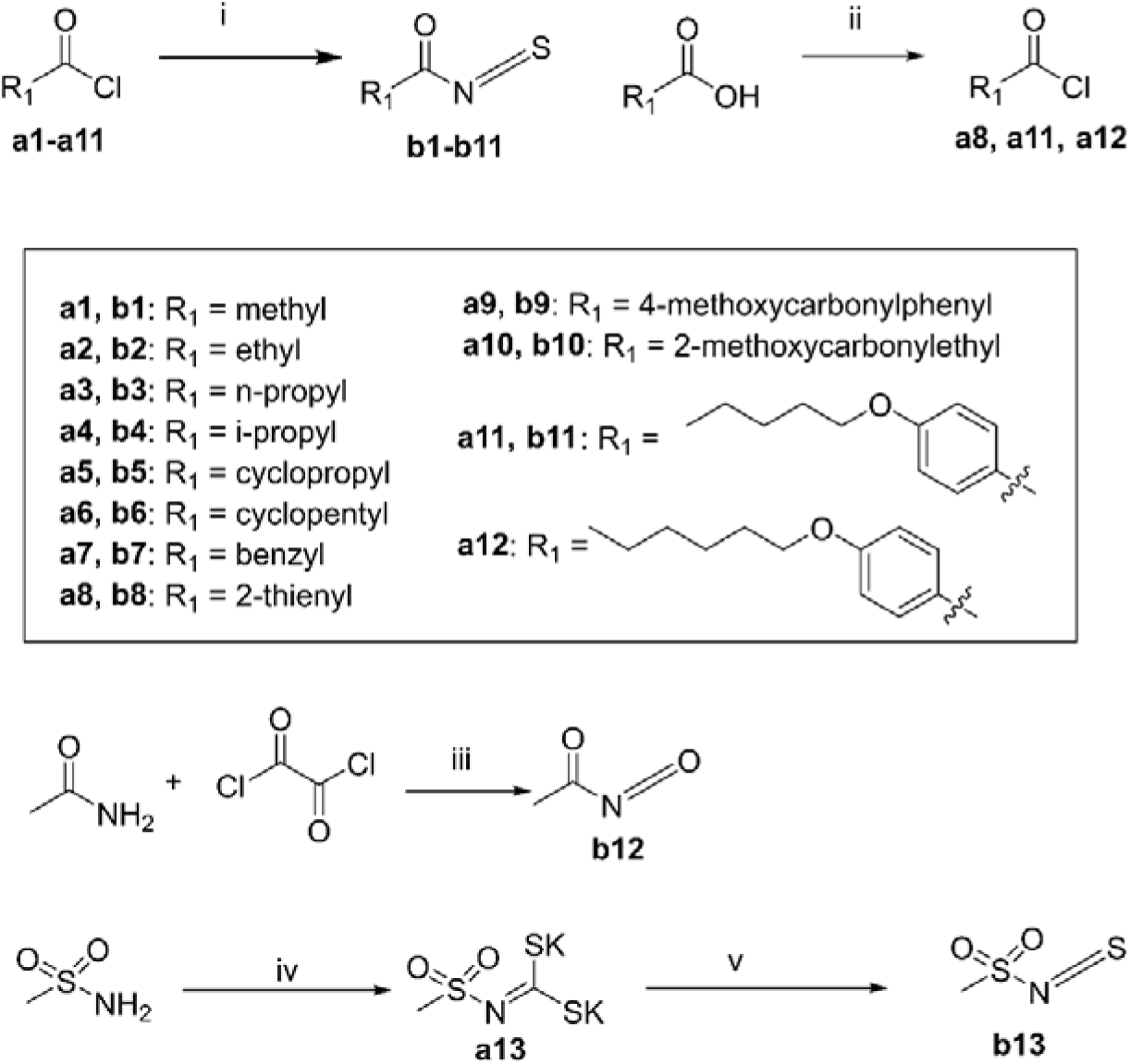
Reagents and conditions: (i) KSCN, acetone, r.t., 0.5 h; (ii) excess SOCl_2_, reflux, 3 h; (iii) 1,2-dichloroethane, reflux, 3 h; (iv) CS_2_, KOH, DMF, r.t., 2 h; (v) triphosgene, benzene, r.t., 6 h.

**Scheme 2.**
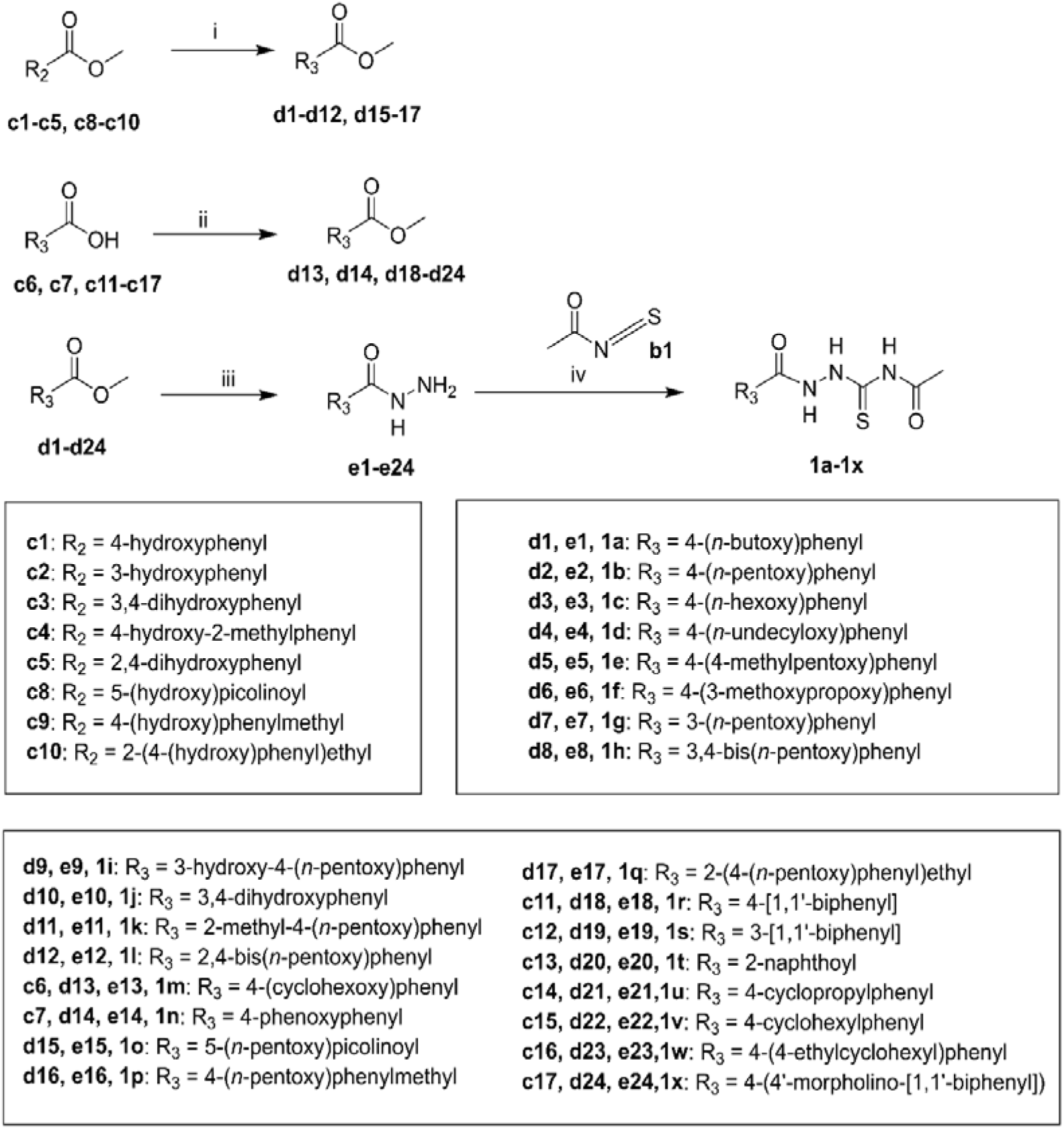
Reagents and conditions: (i) X-Alkyl (X = Br or I), K_2_CO_3_, acetone, reflux, 8 h; (ii) drops of concentrated H_2_SO_4_, MeOH, 80 °C, 10 h; (iii) hydrazine hydrate, MeOH, reflux, 6 h (iv) acetone, reflux, 6 h.

**Scheme 3.**
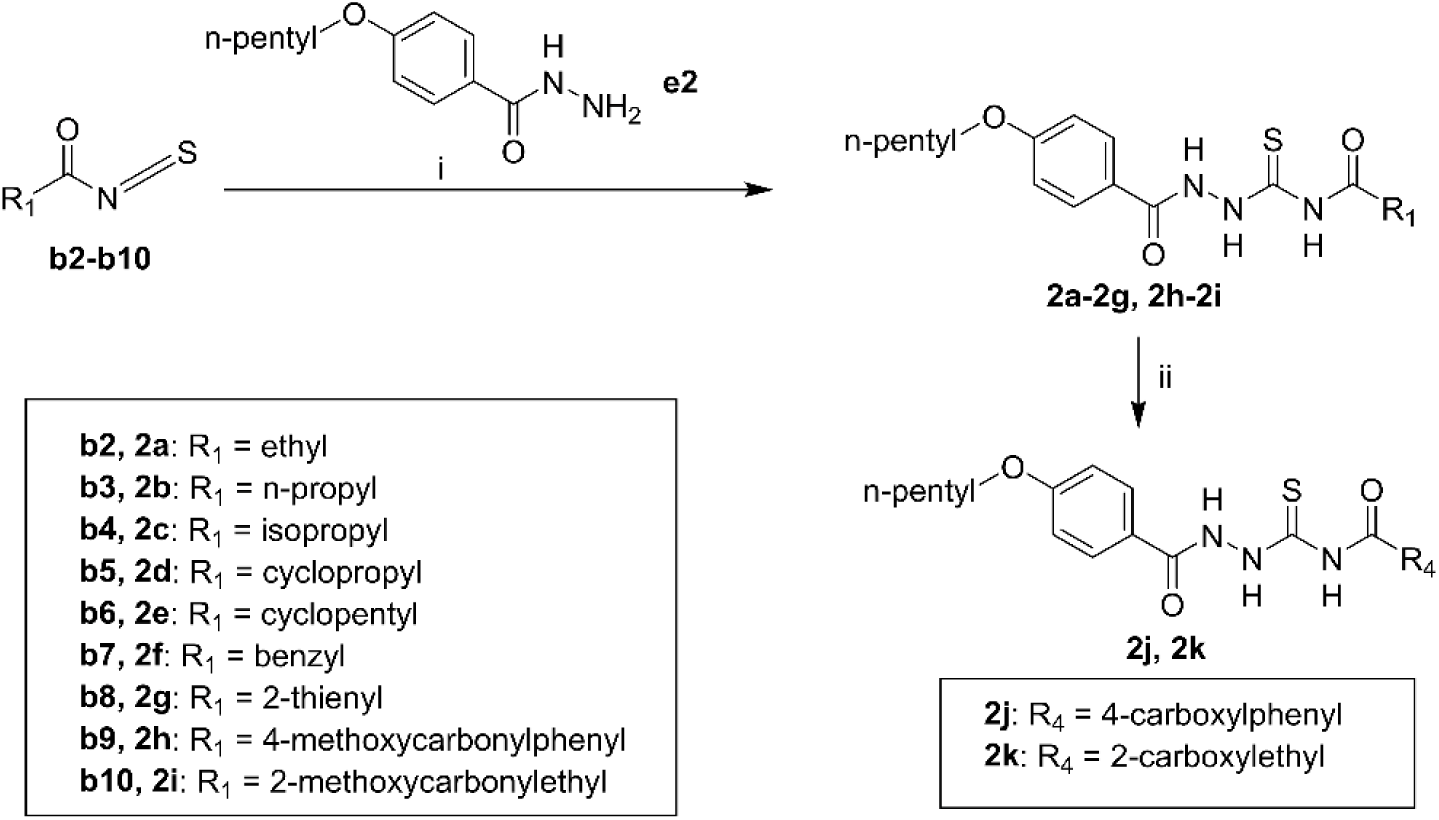
Reagents and conditions: (i) acetone, reflux, 6 h; (ii) 2.0 M of NaOH in MeOH, reflux, 2 h.

**Scheme 4.**
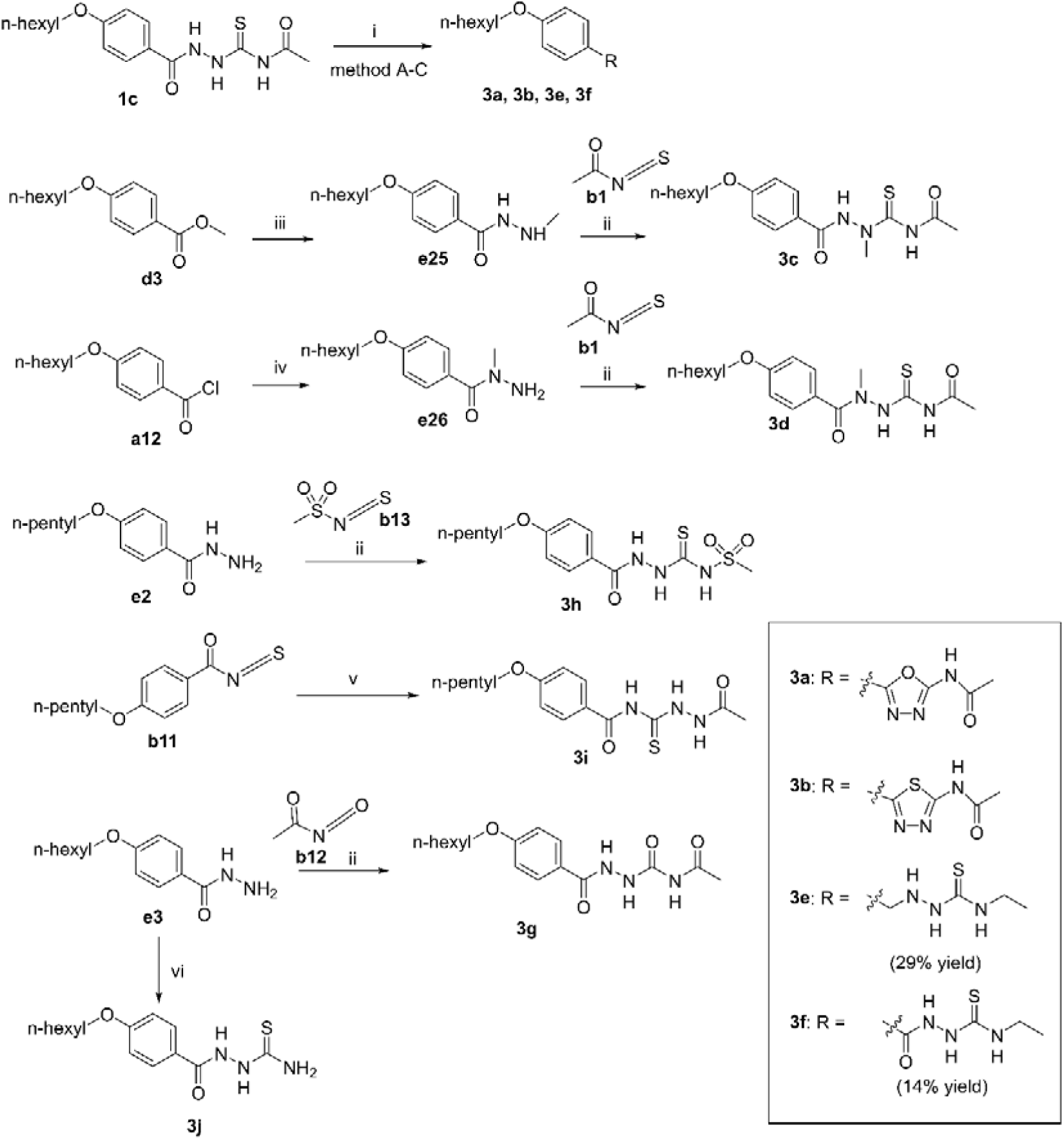
Reagents and conditions: (i) method A (**3a**): KIO_3_, water, reflux; method B (**3b**): acetic acid, microwave, T = 125 C, 5 min, 300 W; method C (**3e**, **3f**): LiAlH_4_, THF, reflux, 6 h; (ii) acetone, reflux, 6 h; (iii) methylhydrazine, MeOH, reflux, 6 h; (iv) methylhydrazine, DCM, r.t., 3 h; (v) acetyl hydrazide, acetone, reflux, 6 h; (vi) *S*-methylisothiourea, H_2_O, reflux, 6 h.

### 2.3 In vitro antibacterial activity and SAR

We determined the MICs of these synthesized derivatives containing the 1,4-dicarbonylthiosemicarbazide scaffold against *S. aureus* (ATCC 29213) by the broth microdilution method [20]. Vancomycin was used as the positive drug, since the mechanism was unknown when we performed the SAR study. The chemical structures and their MIC values of the compounds from three rounds of structural modification are listed in **Table 1**, **Table 3** and **Table 4**, respectively. As shown in the tables, three derivatives (**1b, 1v, 1w**) showed good *in vitro* antibacterial activity, with MIC values less than 1 μg/mL, and more potent than vancomycin (1.56 μg/mL).

**Table 1.**
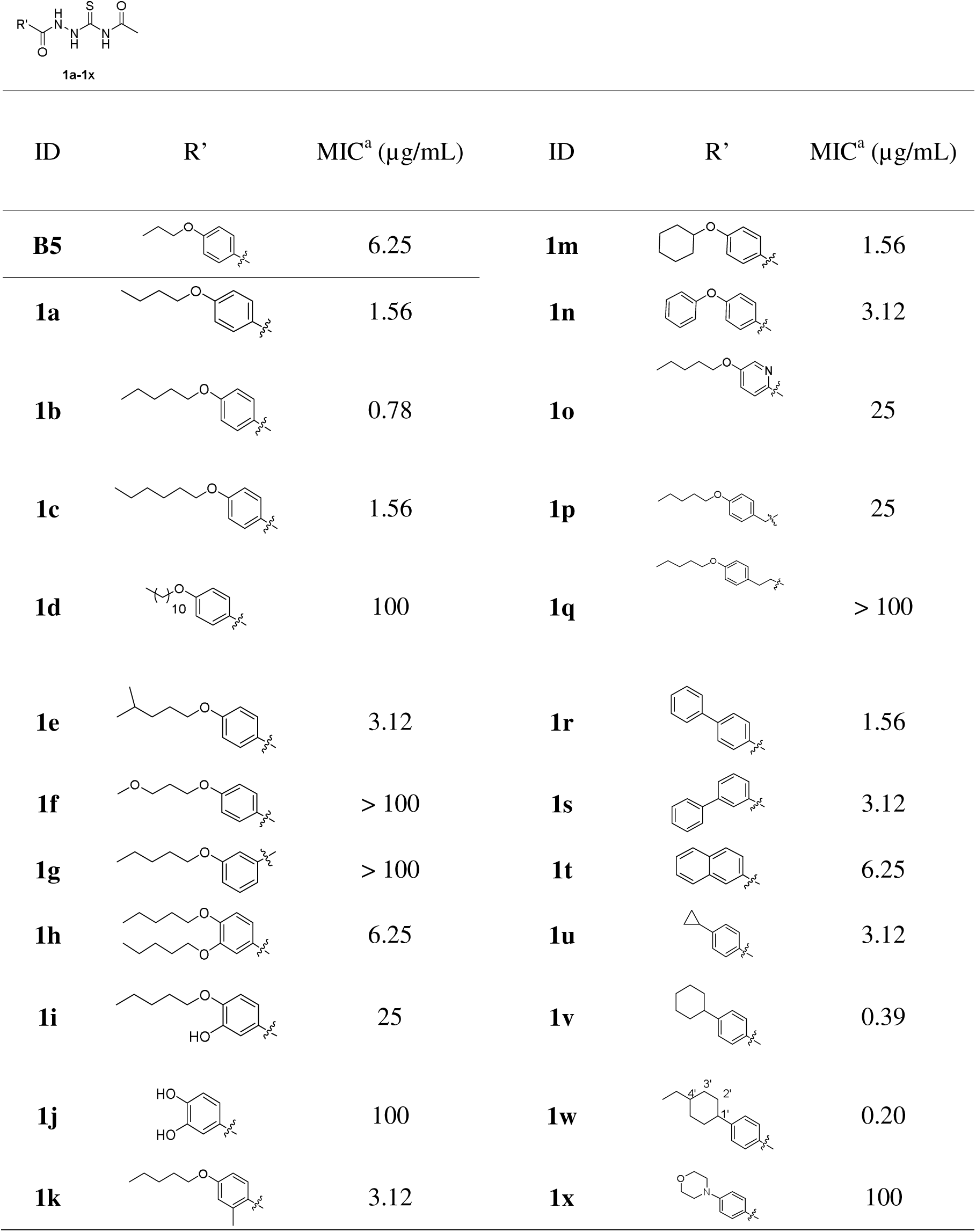
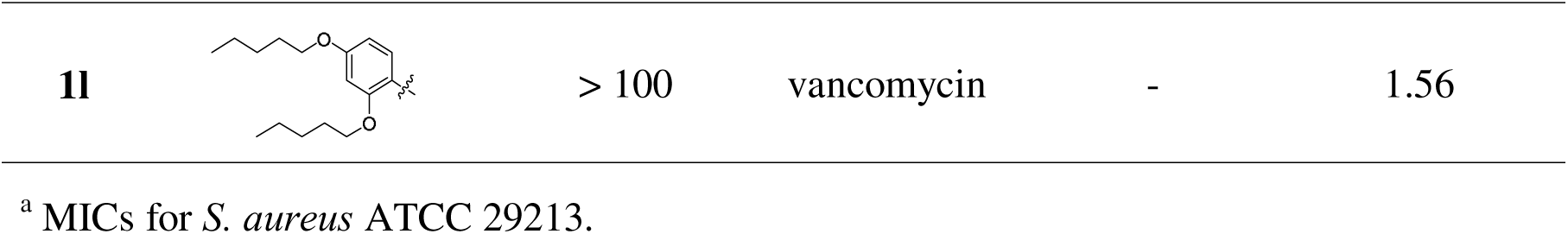
SAR of region A. Chemical structures of the synthesized compounds, and their antibacterial activities in terms of MIC (μg/mL) for *S. aureus* ATCC 29213.

**Table 2.** Comparison of three potent antibacterial compounds, **1b**, **1v**, and **1w** in terms of selectivity index, SI (CC_50_ for HepG-2 and HUVEC, and MIC for *S. aureus* ATCC 29213).

| compounds | CC<sub>50</sub> (μM, mean ± SD)<sup>a</sup> |  | MIC<br>(μM) | SI<sup>b</sup> |  |
| --- | --- | --- | --- | --- | --- |
|  | HepG-2 | HUVEC |  | HepG-2 | HUVEC |
| **1b** | 2.370 ± 0.130 | 6.460 ± 0.980 | 2.41 | 0.98 | 2.68 |
| **1v** | 0.120 ± 0.020 | 0.790 ± 0.080 | 1.22 | 0.10 | 0.65 |
| **1w** | 0.026 ± 0.013 | 0.067 ± 0.041 | 0.58 | 0.05 | 0.12 |
| Paclitaxel | 0.010 ± 0.003 | 0.016 ± 0.012 | n.d.<sup>c</sup> | n.d. | n.d. |

<sup>a</sup> values indicate cytotoxicity after 48-h treatment with the compounds. mean, the average of duplicates; SD, standard deviation.
<sup>b</sup> Selectivity index, SI = CC<sub>50</sub>/MIC<sub>29213</sub>.
<sup>c</sup> n.d., not determined.

**Table 3.** SAR of region C. Chemical structures of the synthesized compounds, and their antibacterial activities in terms of MIC (μg/mL) for *S. aureus* ATCC 29213.

| ID | R'' | MIC<sup>a</sup> (µg/mL) | ID | R'' | MIC<sup>a</sup> (µg/mL) |
| --- | --- | --- | --- | --- | --- |
| **1b** | -CH<sub>3</sub> | 0.78 | **2g** |  | 3.12 |
| **2a** |  | 1.56 | **2h** |  | > 100 |
| **2b** |  | 12.5 | **2i** |  | 12.5 |
| **2c** |  | 12.5 | **2j** |  | > 100 |
| **2d** |  | 6.25 | **2k** |  | 50 |
| **2e** |  | 50 | vancomycin | - | 1.56 |
| **2f** |  | > 100 |  |  |  |

<sup>a</sup> MICs for *S. aureus* ATCC 29213.

**Table 4.**
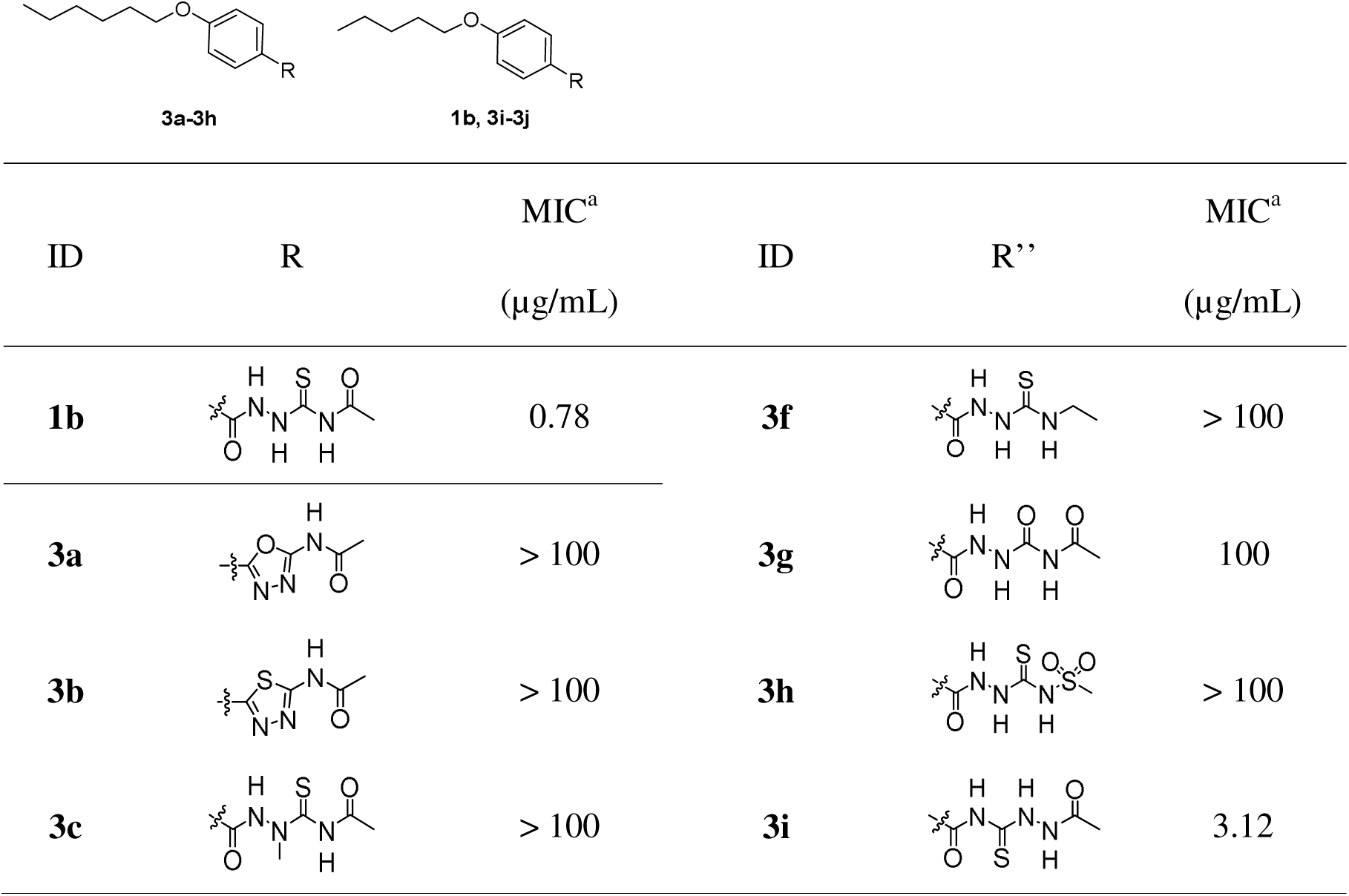
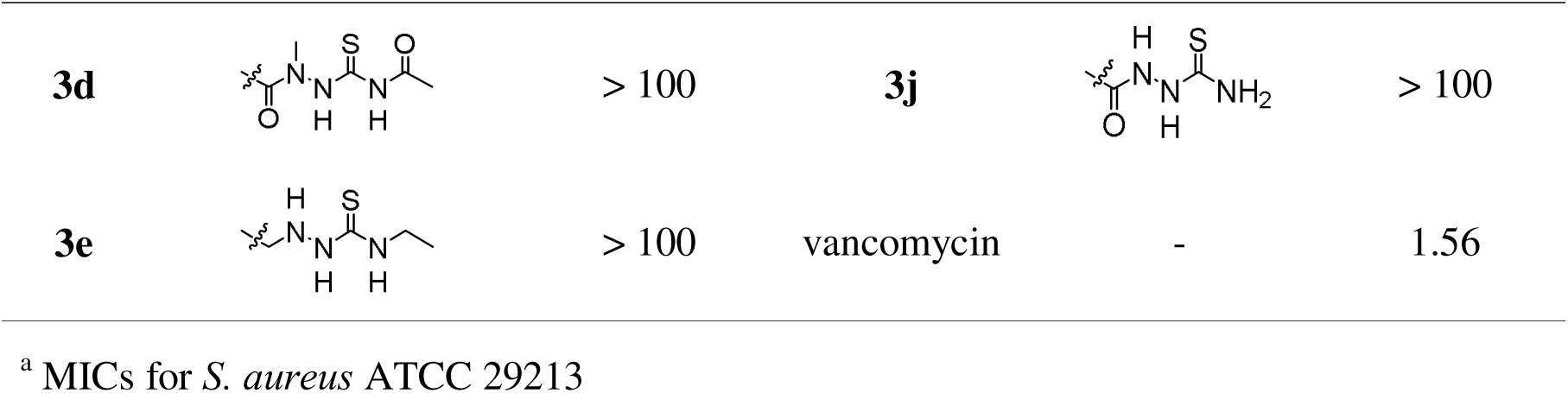
SAR in region B. Chemical structures of the synthesized compounds, and their antibacterial activities in terms of MIC (μg/mL) for *S. aureus* ATCC 29213.

|  |  |  |  |  |  |
| --- | --- | --- | --- | --- | --- |
| **3a-3h** |  | **1b, 3i-3j** |  |  |  |
| ID | R | MIC<sup>a</sup><br>(μg/mL) | ID | R'' | MIC<sup>a</sup><br>(μg/mL) |
| **1b** |  | 0.78 | **3f** |  | > 100 |
| **3a** |  | > 100 | **3g** |  | 100 |
| **3b** |  | > 100 | **3h** |  | > 100 |
| **3c** |  | > 100 | **3i** |  | 3.12 |
| **3d** |  | > 100 | **3j** |  | > 100 |
| **3e** |  | > 100 | vancomycin | - | 1.56 |

<sup>a</sup> MICs for *S. aureus* ATCC 29213

#### 2.3.1 Substitutions at region A and SAR

As mentioned in the “molecular design” section, we firstly synthesized four derivatives with alkoxy chains in different lengths at the *para-* position of the middle benzene ring. Apparently, compound **1b** with the *n*-pentoxy group was the most potent (MIC: 0.78 μg/mL, cf. **1a–1d** in **Table 1**), showing a nearly 10-fold potency increase compared with **B5** (MIC: 6.25 μg/mL), the hit compound from substructure search–based virtual screening. Accordingly, *n*-pentoxy group was the most suitable substituent. Shorter or longer carbon chains were detrimental for the activity.

By the incorporation of branches or oxygen atom to the carbon chain, antibacterial activity for *S. aureus* decreased. To be specific, the MIC values of **1e** and **1f** were 3.12 μg/mL and > 100 μg/mL, respectively.

As **1g** with *meta*-substitution lost its antibacterial activity (MIC: > 100 μg/mL), *para*-substitution seemed better for antibacterial activity.

Furthermore, we studied the effect of di-substitution at the benzene ring. The introduction of two *n*-pentoxy groups at 3,4-positions significantly decreased the antibacterial activity (MIC: 6.25 μg/mL for **1h** vs. 0.78 μg/mL for **1b**). The replacement of the *n*-pentoxy group at 3-position of **1h** by the hydroxy group, **1i** (MIC: 25 μg/mL), or the replacement of both *n*-pentoxy groups at the 3-and 4-positions, **1j** (MIC: 100 μg/mL), further impaired antibacterial activity. With di-substituents at 2,4-positions, the antibacterial activity also decreased, with **1b** as the reference (0.78 μg/mL). To be specific, **1k** with the small methyl group at 2-position still showed some antibacterial activity (MIC: 3.12 μg/mL), whereas **1l** with the large *n*-pentoxy group at 2-position completely lost activity (MIC: > 100 μg/mL). It should be noted that the six-membered ring substituents did not lead to obvious loss of activity, as shown by **1m** (cyclohexyl, MIC: 1.56 μg/mL) and **1n** (phenyl, MIC: 3.12 μg/mL). Additionally, when the middle benzene ring was replaced by the pyridine ring, the resulting derivative **1o** showed lower activity (MIC: 25 μg/mL) than **1b**. When one or two carbon atoms were inserted between the *n*-pentoxyphenyl group and the 1,4-dicarbonylthiosemicarbazide moiety, there was a decrease in antibacterial activity, as shown by **1p** (MIC: 25 μg/mL) or **1q** (MIC: > 100 μg/mL). Derivative **1r**, with the *n*-pentoxy group replaced by the phenyl group, retained antibacterial activity (MIC: 1.56 μg/mL). The derivative with 3-phenyl substituent at the benzene ring, **1s** (MIC: 3.12 μg/mL), and the derivative with naphthalene ring, **1t** (MIC: 6.25 μg/mL), showed decreased antibacterial activity. When the *n*-pentoxy group was replaced with the cyclopropyl group, as shown by **1u** (MIC: 3.12 μg/mL), the antibacterial activity was inferior to that of **1b** (MIC: 0.78 μg/mL). Interestingly, **1v** (MIC: 0.39 μg/mL), with the *n*-pentoxy group replaced by the cyclohexyl group, showed an improved potency, compared with **1b** (MIC: 0.78 μg/mL). Further modification with an ethyl group at the cyclohexyl ring led to further increase in potency (**1w**: 0.2 μg/mL), while 4-morpholino group as a substituent at the *para*-position of the middle benzene ring rendered the compound less active (**1x**: 100 μg/mL).

Mammalian cell toxicity is a property that should be avoided for antibiotics. To determine the best substituent for the follow-up SAR study, we compared three highly potent compounds **1b, 1v,** and **1w** (MIC for *S. aureus* ATCC 29213: < 1 μg/mL) by testing their toxicity to two mammalian cell lines, including human hepatocellular liver carcinoma cells (HepG-2) and human umbilical vein endothelial cells (HUVEC). With the CCK-8 assay, cell viability was measured after the compound treatment for 48 h. According to the concentration of cytotoxicity 50% (CC_50_) and MIC values, we were able to use selectivity index (SI, CC_50_/MIC) to choose the most promising compound. Compound **1b** exhibited the highest SI value (cf. **Table 2**), 0.98 for HepG-2, and 2.68 for HUVEC, respectively, and promising antibacterial activity (MIC: 2.41 μM or 0.78 µg/mL for *S. aureus* ATCC 29213). Although compounds **1v** and **1w** showed slightly better *in vitro* antibacterial activity (1.22 μM/0.39 µg/mL and 0.58 μM/0.2 µg/mL, respectively), they were too toxic to these two mammalian cell lines with low SI values (0.098 and 0.045 for HepG-2, 0.65 and 0.12 for HUVEC, respectively). It indicated that these two compounds were inferior to **1b** in terms of safety. Therefore, the *n*-pentoxybenzoyl group of **1b** was kept for the following SAR exploration.

#### 2.3.2 Substitutions at region C and SAR

Next, we studied whether the methyl group at region C could be extended or modified (cf. **Table 3**). Unfortunately, extending the alkyl chains or introducing cycloalkyl groups led to a decline of potency against *S. aureus*. To be specific, **2a** with the ethyl group (MIC: 1.56 μg/mL), **2b** with the *n*-propyl group (MIC: 12.5 μg/mL), **2c** with the isopropyl group (MIC: 12.5 μg/mL), **2d** with the cyclopropyl group (MIC: 6.25 μg/mL), and **2e** with the cyclopentyl group (MIC: 50 μg/mL), were all inferior to **1b** (MIC: 0.78 μg/mL). Aromatic substituents also resulted in decreased activity, as the MIC values of compounds **2f**, **2g** were 3.12 μg/mL and > 100 μg/mL, respectively. Hydrophilic carboxyl groups were not allowed in this region, as both **2j** (MIC: > 100 μg/mL) and **2k** (MIC: 50 μg/mL) were even less active. Their methyl esters, **2h**: (MIC: > 100 μg/mL) and **2i** (MIC: 12.5 μg/mL) were also less potent than **1b** (MIC: 0.78 μg/mL). Accordingly, the methyl group was considered as the best substituent and should be preserved.

#### 2.3.3 Substitutions at region B and SAR

At last, we studied the SAR of region B to assess the importance of the 1,4-dicarbonylthiosemicarbazide moiety. As a result, this 1,4-dicarbonylthiosemicarbazide moiety was found to be highly conservative and only the switch of thiocarbonyl group from 3-position to 2-position in this moiety was tolerated, as evidenced by compound **3i** that showed the MIC value of 3.12 μg/mL (cf. **Table 4**). Any ring closure into 1,3,4-oxadiazole (**3a**) or 1,3,4-thiadiazole (**3b**), substitution of -NH-with a methyl group (**3c, 3d**), or reduction of carbonyl groups (**3e, 3f**), rendered the compounds completely inactive, with MICs greater than 100 μg/mL.

#### 2.3.4 Overall SAR

According to the **Table 1**, **3–4**, the corresponding structure-activity relationship of this chemical series in different regions is summarized in **Fig. 3**. In region A: (1) *para*-position substituents at the middle benzene ring were favorable for antibacterial activity, and *n*-pentoxy group was the best alkoxy substituent for antibacterial activity; (2) phenyl and cyclohexyl groups were tolerated in this region, and ethyl substitution at 4’-position of the cyclohexane ring could further enhance antibacterial activity while increasing the cytotoxicity at the same time. In region C: (1) small alkyl groups were favored in this region, such as methyl and ethyl; (2) as this region might be hydrophobic, polar carboxylic groups were detrimental for the potency. In region B: (1) the 1,4-dicarbonylthiosemicarbazide moiety was essential for antibacterial activity, and this moiety should not be replaced by heterocycles; (2) the position of thiocarbonyl group could be switched from 3-position to 2-position.

**Fig. 3.**
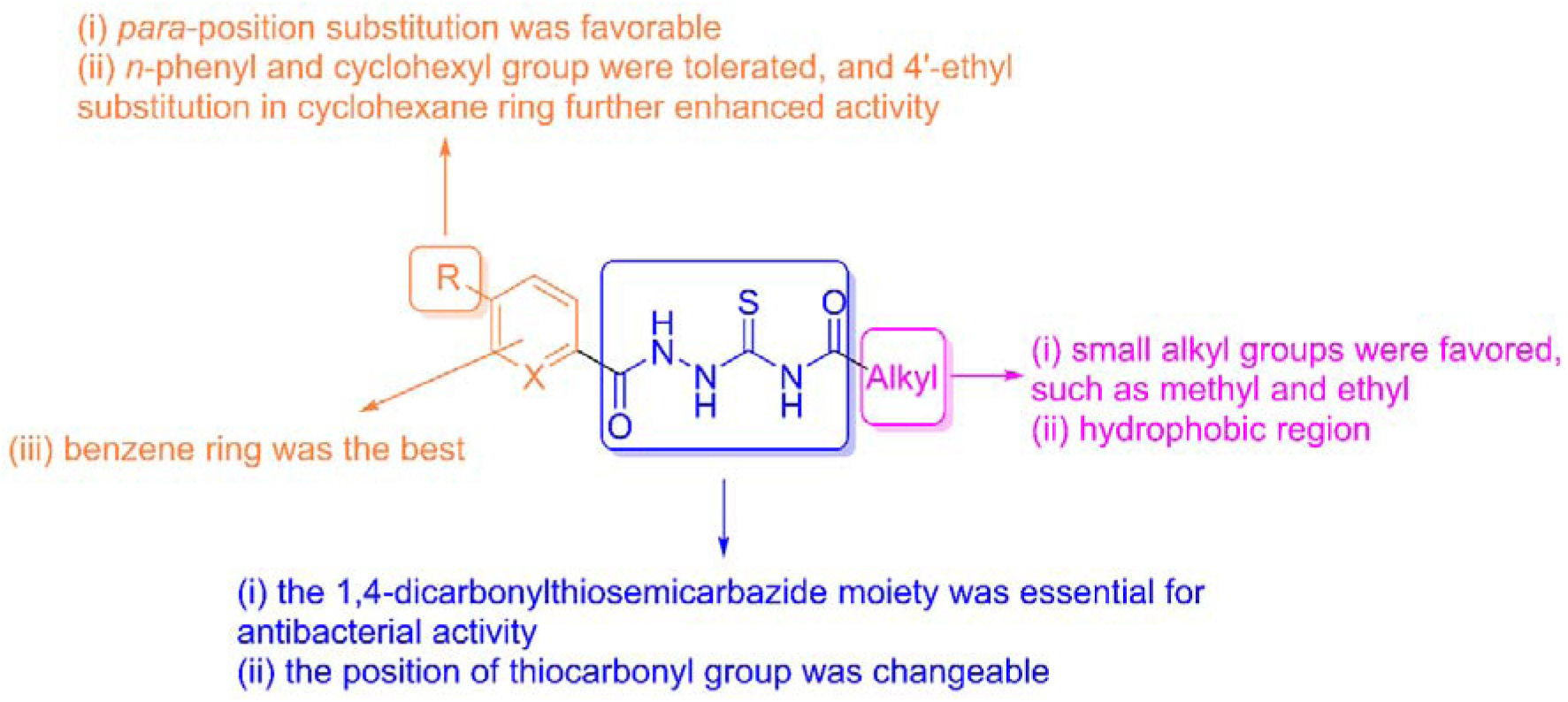
Overall SAR of the 1,4-dicarbonylthiosemicarbazide derivatives.

### 2.4 The biological profile of compound **1b**

#### 2.4.1 General

Among these derivatives, compound **1b** displayed good activity against *S. aureus* and the best safety profile. Therefore, it was selected for follow-up research to explore the antibacterial mechanism of the 1,4-dicarbonylthiosemicarbazides. We performed several biological assays to assess its potential as a small-molecule antibiotic, including: (1) time-kill analysis of compound **1b** against *S. aureus*; (2) antibacterial activity against a panel of gram-positive bacterial strains; (3) *in vitro* metabolic stability; and (4) *in vivo* efficacy in MRSA-infected mouse model.

#### 2.4.2 Time-kill analysis of compound **1b** against S. aureus (ATCC 29213)

To understand the mode of action (bactericidal or bacteriostatic) by which compound **1b** killed *S. aureus* (ATCC 29213), a time-kill kinetics assay was performed, covering three concentrations (1 × MIC, 4 × MIC, and 16 × MIC) and four time points (0, 4, 8, and 24 h). As shown in **Fig. 4A**, compound **1b** exhibited a bacteriostatic mode at the 1 × MIC, 4 × MIC concentrations and bactericidal mode at the 16 × MIC concentration. At the 1 × MIC concentration, **1b** inhibited bacterial growth for 24 h. At the 4 × MIC and 16 × MIC concentrations, **1b** reduced the bacteria colony counts by 2.51 and 6.41 log units after 24 h, respectively. As a positive control, the time-kill kinetics study for vancomycin was also performed. The results showed that vancomycin was bactericidal as the starting bacteria concentration (6.41 log_10_CFU/mL) decreased by more than 3 log units after treatment for 24 h at the concentration of 1 × MIC, 4 × MIC, and 16 × MIC. Therefore, **1b** showed a bacteriostatic effect at the low concentrations (1 × MIC, 4 × MIC), while exhibiting a bactericidal effect at the high concentration (16 × MIC).

**Fig. 4.**
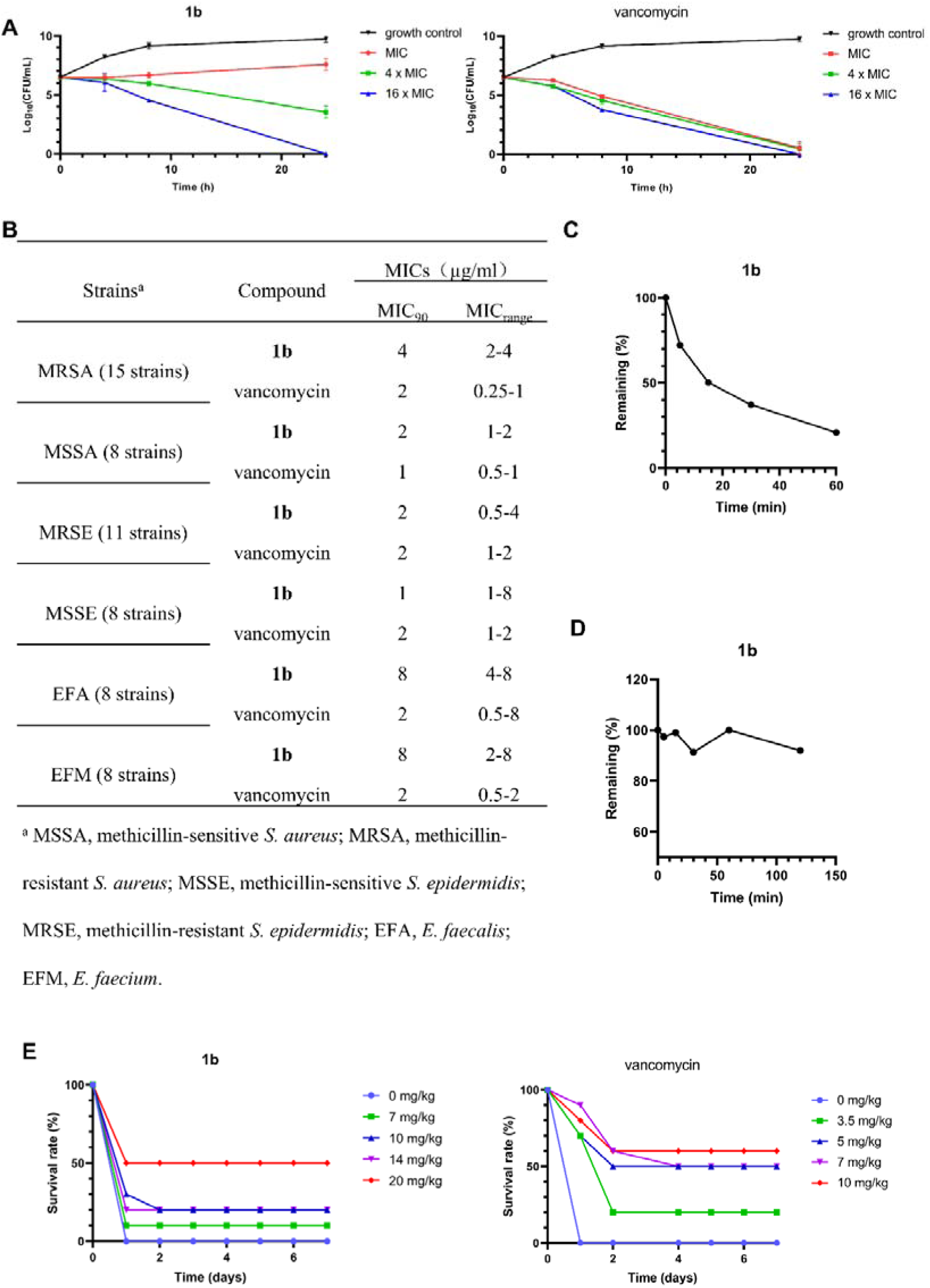
The biological evaluation of compound **1b**. (A) Time-kill analysis of compound **1b** against *S. aureus* ATCC 29213, with vancomycin as the positive control. (B) Antibacterial activity of compound **1b** for a panel of gram-positive bacteria strains. (C) Metabolic stability of **1b** in mouse liver microsomes. (D) Metabolic stability of **1b** in mouse plasma. (E) Survival rate curves of the MRSA 16-1 (clinical isolate)–infected mice treated by compound **1b** or vancomycin.

#### 2.4.3 Antibacterial activity against gram-positive bacteria

We further tested **1b** for its activity against 58 gram-positive bacterial strains, including 15 MRSA, 8 methicillin-sensitive *S. aureus* (MSSA), 11 methicillin-resistant *S. epidermidis* (MRSE), 8 methicillin-sensitive *S. epidermidis* (MSSE), 8 *E. faecalis* (EFA), and 8 *E. faecium* (EFM) strains, with vancomycin as the positive control. Compared with its activity against ATCC 29213 (MIC: 0.78 μg/mL) in the primary SAR study, **1b** showed similar antibacterial potency for the tested strains, with MICs ranging from 0.5 μg/mL to 8 μg/mL (cf. **Fig. 4B** and **Table S1, S2**). Its MIC values were close to those of vancomycin (MICs: 0.5–8 μg/mL).

#### 2.4.4 In vitro metabolic stability

Metabolic stability of small molecules is an important property for drug design, as it affects *in vivo* pharmacokinetic profile and thus *in vivo* efficacy. Before *in vivo* efficacy evaluation, we studied metabolic stability properties of **1b** by incubating the compound with mouse liver microsomes or mouse plasma and analyzed it with LC-MS/MS method (cf. **Table S3**). As shown in **Fig. 4C** and **Table S4**, the value of T_1/2_ (i.e., half-life) and intrinsic clearance (CL_int_) for **1b** in mouse liver microsomes was estimated as 28.3 min and 49.0 mL/min/mg protein, respectively. It indicated that **1b** was a compound of slow clearance [27]. Besides, the estimated T_1/2_ value in mouse plasma exceeded 2 h, indicating **1b** was stable in mouse plasma (cf. **Fig. 4D** and **Table S4**). These data demonstrated that **1b** was stable in terms of *in vitro* metabolism.

#### 2.4.5 In vivo efficacy in MRSA-infected mouse model

Due to good *in vitro* antibacterial activity and metabolic stability, compound **1b** was further assessed for its *in vivo* efficacy by treating the infected mice (clinical isolate MRSA 16-1) and measuring the survival rate. Prior to the efficacy study, we determined the minimum lethal dose (MLD) as 1.30 × 10^6^ CFU/mL, by the intraperitoneal injection of the MRSA 16-1 inoculum (0.5 mL). After being infected by MRSA 16-1, we administered the compound **1b** at different doses by intravenous injection, with vancomycin as the positive control. The mice treated with only vehicle control (saline, 0 mg/kg drug) showed a survival rate of 0%. A dose-response relationship of **1b** or vancomycin was observed from **Fig. 4E**. Both compounds displayed protective effects and led to survival of the mice. As shown in **Fig. 4E**, the survival rates were 50%, 20%, 20% and 10%, when the infected mice were treated with the compound **1b** at the doses of 20 mg/kg, 14 mg/kg, 10 mg/kg and 7 mg/kg, respectively. The survival rates of the infected mice treated with vancomycin were 60%, 50%, 50%, 20% at the doses of 10 mg/kg, 7 mg/kg, 5 mg/kg and 3.5 mg/kg, respectively. These results demonstrated that compound **1b** was effective to protect the mice from MRSA infection, which might be further optimized into a new class of antibiotic.

### 2.5 Molecular mechanism: target prediction and experimental validation

#### 2.5.1 General workflow

To the best of our knowledge, the protein target for antibacterial activity of 1,4-dicarbonylthiosemicarbazides had not been uncovered, though a few attempts were made to address it [13–17]. Herein, we applied 2D similarity calculation and reverse docking to identify the most plausible protein target for our chemical series (cf. **Fig. 5A** and **Fig. S1**), and then we performed *in vitro* assays to validate it.

**Fig. 5.**
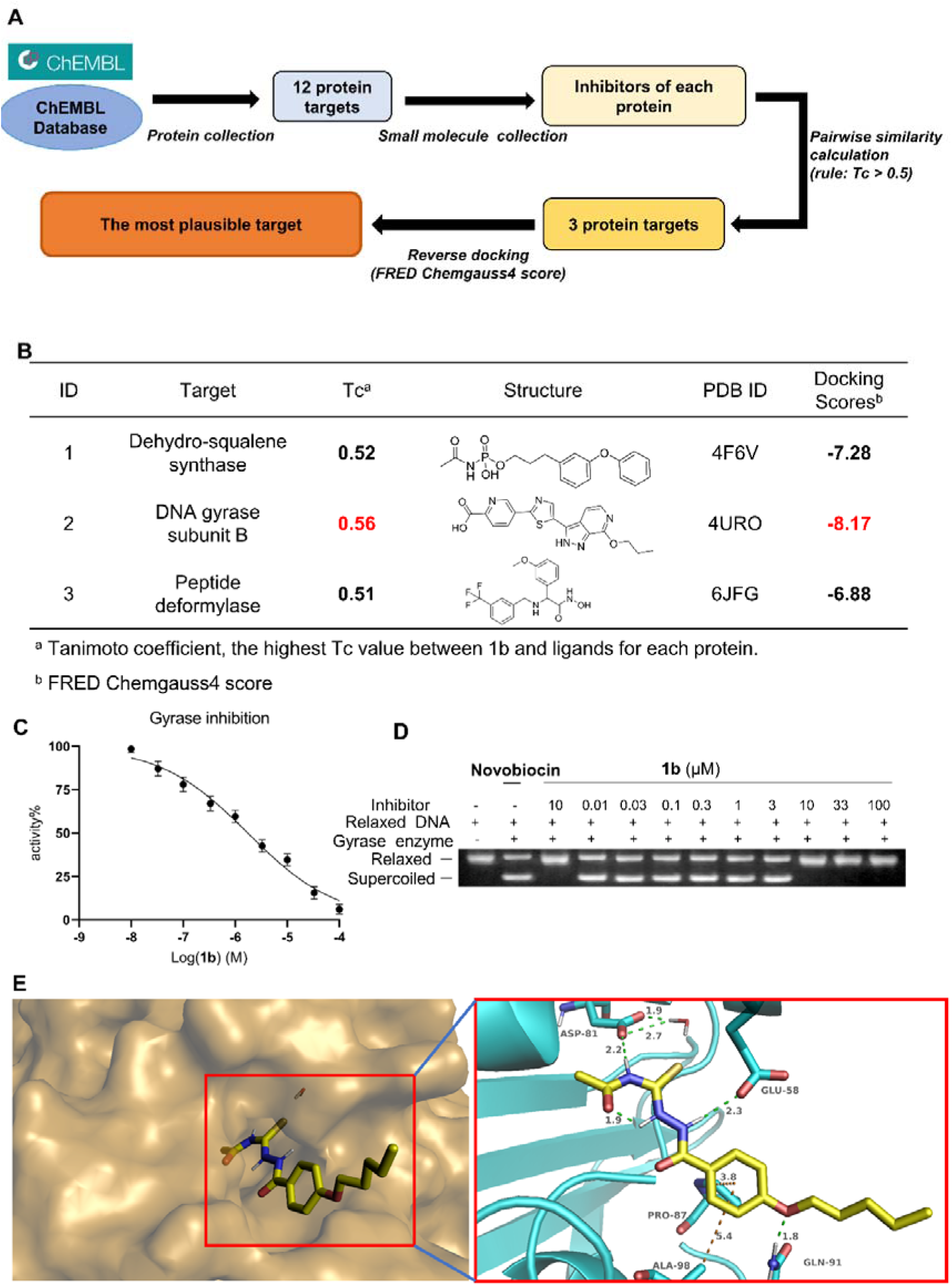
Computer-aided target identification. (A) Our work-flow of target prediction. (B) Three potential target proteins for **1b**, with similarity values and docking scores listed. (C) The concentration-dependent ATPase inhibition of *S. aureus* gyrase. The IC_50_ value of **1b** was 1.81 μM. (D) Agarose gel electrophoresis assay. **1b** inhibited the formation of supercoiled DNA. (E) The binding pocket of gyrase ATPase domain (left, shown as a surface representation) and the protein-ligand interaction mode between gyrase ATPase domain and **1b (**right, the binding mode was optimized by molecular dynamics simulation).

To begin with, we searched the ChEMBL database (ChEMBL34, accessed in March 2024) for all protein targets related to *S. aureus*. For “single protein” targets, we chose 12 proteins that had at least one co-crystal structure with a bound ligand. They were: cell division protein FtsZ, dehydrosqualene synthase, dihydrofolate reductase, dihydroneopterin aldolase, DNA gyrase subunit A, DNA gyrase subunit B, DNA ligase, enoyl-[acyl-carrier-protein] reductase (FABI), methionine aminopeptidase, peptide deformylase, pyruvate kinase, and UDP-*N*-acetylmuramate dehydrogenase (cf. **Table S5**). Next, we curated all the inhibitors of each protein (cf. **Table S5** for details) and performed pairwise similarity calculation between **1b** and inhibitors of each protein using “Find Similar Molecules by Fingerprints (MDLPublicKeys)” module of Discovery Studio (version 19.1.0). Three proteins had inhibitors that were highly similar to **1b** (Tanimoto coefficient [28] > 0.5). To be specific, the protein of dehydrosqualene synthase had inhibitors similar to **1b** with Tc value of 0.52. The DNA gyrase subunit B had the most similar inhibitor to **1b** with Tc value of 0.56. The inhibitors of peptide deformylase also exhibited high similarity to **1b** with Tc value of 0.51 (cf. **Fig. 5B**). Compound **1b** was further docked with OEDocking (version 3.3.1.2, OpenEye Scientific Software, Santa Fe, NM, USA) against the pocket of each of the three protein structures. To be specific, 4F6V, 4URO, and 6JFG (PDB ID) were used as the representative crystal structures to represent each of the three proteins, and the binding sites were defined by their cognate ligands, respectively. The pose similar to that of the cognate ligand was regarded as the binding pose and its Chemgauss4 score was used to measure binding affinity. The predicted binding modes and scores by molecular docking are shown in **Fig. S2**.

As shown in **Fig. 5B**, the predicted binding affinity of **1b** to the ATP binding site of DNA gyrase was the highest (Chemgauss4 score: -8.17). Therefore, it was very likely that **1b** could inhibit gyrase ATPase activity by binding to the ATP binding site.

#### 2.5.2 **1b** inhibited S. aureus gyrase ATPase and DNA supercoiling

To verify our prediction, we performed experimental validations, with ATP hydrolysis inhibition assay and DNA supercoiling inhibition assay. As shown in **Fig. 5C–D**, (1) compound **1b** inhibited the gyrase ATP hydrolysis activity, with IC_50_ value of 1.81 μM (cf. **Fig. 5C**). (2) Since gyrase ATPase inhibition eventually impair DNA supercoiling [29], we tested that effect of **1b** by agarose gel electrophoresis. As a result, **1b** inhibited the production of supercoiled DNA in a dose-response manner (cf. **Fig. 5D**). We further tested other potent antibacterial derivatives for their inhibitory activity for *S. aureus* gyrase. The outcome was expected, **1c, 1v**, and **1w** all showed potent gyrase inhibition (IC_50_ values: **1c**: 1.01 μM; **1v**: 1.89 μM; and **1w**: 0.23 μM). These data further confirmed DNA gyrase as the target of this chemical series.

#### 2.5.3 Binding mode

By a 200-ns molecular dynamics simulation with GROMACS (version 2019.4) (cf. **Fig. S4**), we were able to propose the binding mode of compound **1b** to gyrase ATPase domain. **Fig. 5E** (left) showed the binding mode of **1b** to gyrase ATP binding site. To be specific, compound **1b** well occupied the gyrase ATP binding pocket (the conserved water molecule was preserved) and mainly formed two types of interactions with gyrase ATPase domain (cf. **Fig. 5E**, right): (1) Hydrogen bonding: Two -NH-groups of the 1,4-dicarbonylthiosemicarbazide moiety formed hydrogen bonds with ASP81 and GLU58, respectively; another hydrogen bond was also observed between oxygen atom of alkoxy group and GLN91. (2) π-alkyl interactions: the benzene ring of **1b** could form π-alkyl interactions with ALA98 and PRO87. The aforementioned interactions were consistent with those between novobiocin and gyrase ATPase domain (cf. **Fig. S5**, PDB code: 4URO [30]).

## 3. Conclusion

Due to the development of antibiotic resistance in *S. aureus*, *i.e.* the emerging of MRSA or even vancomycin-resistant *S. aureus*, new classes of antibiotics are urgently needed. The 1,4-dicarbonylthiosemicarbazides represent a potential chemotype for novel antibacterial drug discovery. However, the previously reported compounds are not as potent as clinical drugs, e.g. vancomycin, and their true antibacterial mechanism remains to be elucidated.

In this study, we firstly used 1,4-dicarbonylthiosemicarbazide moiety for substructure search to identify diverse antibacterial agents. Compound **B5** (Specs ID: AG-690/15432331), the 1,4-dicarbonylthiosemicarbazide as our hit compound bearing the *n*-propoxybenzoyl group, showed moderate antibacterial activity for *S. aureus* (MIC: 6.25 μg/mL). We then synthesized 45 new 1,4-dicarbonylthiosemicarbazides with alkylcarbonyl substituents at position 4 and arylcarbonyl groups at position 1 by using various routes and evaluated their activities against *S. aureus* (ATCC 29213). Among these derivatives, compound **1b** exhibited better antibacterial activity than vancomycin (0.78 μg/mL vs. 1.56 μg/mL for vancomycin), the highest SI value (selective index, 0.98 for HepG-2, and 2.68 for HUVEC, respectively), and also good *in vivo* activity (50% survival rate at the dose of 20 mg/kg). To address the issue of molecular mechanism, we performed target prediction by similarity calculation and reverse docking, and proposed DNA gyrase as its target protein. Through *S. aureus* gyrase ATPase inhibition assay, the IC_50_ values of four potent derivatives were determined to be 0.23–1.89 μM. The following agarose gel electrophoresis assay further supported our hypothesis that **1b** inhibited DNA supercoiling via affecting ATPase activity of *S. aureus* gyrase. We also proposed a possible interaction mode between **1b** and gyrase ATP binding site via molecular docking and a 200-ns molecular dynamics simulation, where strong hydrogen bonds and π-alkyl interactions were observed, which were consistent with the binding mode of the classical gyrase inhibitor, novobiocin. Nevertheless, it should be noted that the SI of **1b** was not high enough, and thus optimization to improve selectivity will be our future work.

In summary, we discovered a novel type of DNA gyrase inhibitors as promising antibacterial agents. The mechanism study facilitated by computational modeling provides new insight into the 1,4-dicarbonylthiosemicarbazides and may facilitate further lead optimization.

## 4. Experimental

### 4.1. Computation

#### 4.1.1 Substructure search

The “substructure search” protocol implemented in Pipeline Pilot (version 19.1.0; Dassault Systèmes Biovia Corp.) was used for our purpose. In this protocol, 1,4-dicarbonylthiosemicarbazide was set as the substructure, and the Specs chemical library (http://www.specs.net, > 210,000 compounds) was set as the screening database. The 1,4-dicarbonylthiosemicarbazide derivatives obtained were further selected mainly based on structure diversity as well as commercial availability.

#### 4.1.2. Target prediction

##### Data collection and curation

In the “Targets” section of the ChEMBL database (ChEMBL34, accessed in March 2024), data with the organism taxonomy of “Staphylococcus” were retrieved. On the target list, only when a target belonged to the type of “SINGLE PROTEIN” and the organism of “*Staphylococcus aureus”* (including different strains), it was collected. Then, the names of these targets were searched from RCSB PDB (https://www.rcsb.org/). Among them, the proteins that had been resolved in complex with at least one cognate ligand were included for the following analysis. For each of the above mentioned proteins, small molecules and their activity data (IC_50_, K_d_, K_i_) were obtained from the ChEMBL database. The molecules that had the activity value less than 10 μM were considered as inhibitors of that protein, which comprised the corresponding inhibitor dataset.

##### 2D Similarity calculation

The “Find Similar Molecules by Fingerprints” module in Discovery Studio (v19.1.0, Dassault Systèmes Biovia Corp) was used for 2D similarity calculation. In this module, the structure of compound **1b** was set as the reference ligand. MDLPublicKeys was the molecular fingerprinting algorithm, while Tanimoto coefficient was the metric to measure 2D similarity. The similarity of every compound in the inhibitor dataset to **1b** was calculated. The most similar one was recorded and its similarity was used to evaluate the likeliness of **1b** being an inhibitor of that protein.

##### Reverse docking

Prior to molecular docking, the identical protein chain and nonessential cofactors were removed from the protein crystal structure. All the co-crystallized water molecules, except for the conserved one potentially involved in ligand binding, were removed. Then, the “Clean Protein” module of Discovery Studio was used to correct all potential problems with amino acids such as alternative conformations, nonstandard names, incomplete residues and terminus, incorrect bonds and bond orders as well as atom orders, and add hydrogen atoms as well as generate a protonation state at pH 7.0. The “Prepare Ligands” module of Discovery Studio (v19.1.0, Dassault Systèmes Biovia Corp) was used to ionize the structure of compound **1b** at pH 7.4. The compound was docked against the binding site defined by the cognate ligand from the co-crystal structure of each of the proteins, by sequentially using OMEGA (version 3.1.1.2, OpenEye Scientific Software, Inc., Santa Fe, NM, USA) [32] and OEDocking (version 3.3.1.2, OpenEye Scientific Software, Inc., Santa Fe, NM, USA) [40]. The binding affinity was scored by the Chemgauss4 scoring function.

#### 4.1.3 Molecular dynamics simulation

Molecular dynamics (MD) simulations were performed using GROMACS (version 2019.4) [33]. The GROMOS96 43A1 force field [34] was used to generate the topology file for the protein (herein, DNA gyrase subunit B, PDB ID: 4URO), while the topology file of the compound was generated by using the LigParGen server (https://zarbi.chem.yale.edu/ligpargen/) [35]. The geometry of the compound was optimized with the HF/6-31G* level of theory and basis set by Gaussian 09W, and atomic charges were assigned based on the molecular electrostatic potentials, by the RESP module of AMBER21. The whole system was put in a cubic box, with the edge 10 Å distant from the periphery of the system in each dimension and then solvated with 12,887 simple point charge (SPC) water [46]. Then, 10 Na^+^ ions were added to neutralize the whole system. The simulation began with 5000-step energy minimization based on the steepest descent algorithm, followed by the equilibration phase composed of 500-ps simulation under NVT condition and 500-ps simulation under NPT condition. The Parrinello-Rahman was applied to maintain the system at the pressure of 1 atm. The V-rescale was applied to maintain the system at the temperature of 300 K. Lastly, a 200-ns MD simulation was performed under NPT, during which the coordinates of the complex were saved every 100 ps.

### 4.2. Chemistry

#### 4.2.1. General methods

All of the reagents were obtained from commercial sources and used without further purification. Thin-layer chromatography (TLC) on the silica gel plates GF_254_ (200–300 mm; Qingdao Haiyang Chemical Co., Ltd., Qingdao, China) with UV light illumination was used to monitor chemical reactions. ^1^H NMR (500 MHz) and ^13^C NMR (125 MHz) spectra were measured by Avance spectrometer (Bruker, Varian Mercury, USA). Chemical shifts were reported in δ values (ppm) with tetramethylsilane as the internal standard. High-resolution mass spectrometry (HRMS) was performed using the Thermo Scientific™ Exactive™ Plus mass spectrometer (Thermo, USA). The purity was determined by high-performance liquid chromatography (HPLC) on a Waters Acquity machine with a BEH C_18_ column (1.7 µm, 50 × 2.1 mm); mobile phase A = water (containing 0.1% formic acid) and mobile phase B = acetonitrile; the flow rate was 0.25 mL/min.

#### 4.2.2 General procedures for the preparation of intermediates **a8, a11, a12, b1–b13**

##### Preparation of **a8**, **a11** and **a12**

They were not commercially available and thus prepared according to the following method. 2-thienyl chloride **a8,** was prepared from 2-thienylcarboxylic acid (1.0 equiv) and excessive SOCl_2_ (as solvent) under reflux to give the product. **a11** and **a12** were prepared via the same method as that for **a8**, with 4-(*n*-pentoxy)benzoic acid and 4-(*n*-hexoxy)benzoic acid as the starting material, respectively.

##### Preparation of **b1–b11**

Potassium thiocyanate (2.8 mmol, 1.0 equiv) was dissolved in 10 mL of acetone in a 50-mL flask, then acetyl chloride **a1** (2.8 mmol, 1.0 equiv) was then added dropwise. The reaction was carried out for 0.5 h. Then solvent was kept and the formed solution of intermediate **b1** was directly used for the next chemical reaction without further purification. Intermediates **b2–b11** were obtained using the same method as that for **b1** with acyl chlorides **a2–a11**.

##### Preparation of **b12**

To a solution of 0.15 g (2.5 mmol, 1.0 equiv) of acetamide in 15 mL of 1,2-dichloroethane in a 50 mL flask, 0.37 mL (4.3 mmol, 1.7 equiv) of oxalyl chloride was added dropwise and stirred under reflux for 3 h. After the reaction, the solvent was removed to give the intermediate **b12**, which was used for the next chemical reaction without further purification.

##### Preparation of **b13**

To a mixture of methylsulfonamide (1.0 g, 10.5 mmol, 1.0 equiv) and carbon disulfide (0.63 mL, 10.5 mmol, 1.0 equiv) in DMF (7 mL) in a 25 mL flask, KOH (0.59 g, 10.5 mmol, 1.0 equiv) was added. The reaction was carried out for 2 h at room temperature. After that, the other portion of KOH (0.59 g, 10.5 mmol, 1.0 equiv) was added, and the solid was filtered to afford yellow solid **a13**, 2.5 g, yield 96%. The mixture of the yellow solid **a13** (1.0 g, 4.0 mmol, 1.0 equiv) and triphosgen (0.44 g, 1.4 mmol, 0.35 equiv) in 25 mL of benzene was stirred at room temperature. When completed, the solid was filtered out, and the solvent was concentrated to give brown oil intermediate **b13**.

#### 4.2.3. General procedures for the preparation of intermediates **d1–d24**

##### 4.2.3.1. General procedures for the preparation of intermediates **d1–d12, d15–d17**

###### Preparation of **d1–d12**, **d15–d17**

To a 100-mL flask, methyl 4-hydroxybenzoate (**c1**, 5.0 mmol, 1.0 equiv), and potassium carbonate (15 mmol, 3.0 equiv) were added along with 50 mL acetone. The mixture was heated to reflux, then butyl iodide (7.5 mmol, 1.5 equiv) was added. TLC was used to monitor the reaction until the reaction was completed. Upon completion, the solvent was removed under reduced pressure. Then 100 mL of water was added and the aqueous solution was extracted with ethyl acetate (30 mL × 3). The organic layers were combined and dried over anhydrous sodium sulfate. The solvent was removed to give the intermediate **d1**, which was purified by silica column chromatography. Intermediates **d2–d12**, **d15–d17** were obtained via the same method as that for **d1** with different alkyl iodides or bromides.

##### 4.2.3.2. General procedures for the preparation of intermediates **d13–d14, d18–d24**

###### Preparation of **d13–d14**, **d18–d24**

To a solution of 4-phenoxybenzoic acid (**c6**, 1.97 mmol, 1.0 equiv) in 20 mL of methanol, several drops of concentrated sulfuric acid (98%) were added, and the resulting solution was refluxed for 8 h until the methylation was completed. The solvent was then removed to afford crude intermediate **d13**, which was used for the next step without further purification. Intermediates **d14, d18–d24** were obtained via the same method as that for **d13**.

#### 4.2.4. General procedures for the preparation of intermediates **e1–e26**

##### Preparation of **e1–e25**

To a 100-mL round bottom flask, intermediate **d1** (3 mmol, 1.0 equiv), and hydrazine hydrate (80%) aqueous solution (30 mmol, 10.0 equiv) were added to 30 mL of methanol. The solution was heated to reflux for 8 h. TLC was used to monitor the reaction until the reaction was completed. The solvent was then removed under reduced pressure to give solid product. The solid was washed with water for 3 times, recrystallized with methanol, and dried to give intermediate **e1**. Intermediates **e2–e24** were obtained via the same method as that for **e1**.

**e25** was synthesized via the same method as that for **e1**, except that methylhydrazine (180 mmol, 60.0 equiv) was used instead.

##### Preparation of **e26**

To a 50-mL flask, 4-(*n*-hexoxy)benzoic acid (4.5 mmol, 1.0 equiv), and 10 mL of SOCl_2_ were mixed and refluxed at 80 C for 2 h to give intermediate **a12**. Upon completion, thionyl chloride was evaporated. Next, 4-(*n*-hexoxy)benzoyl chloride **a12** (4.5 mmol, 1.0 equiv) and methylhydrazine (45 mmol, 10.0 equiv) were added to 20 mL of tetrahydrofuran (THF), stirred at room temperature. Upon completion, the solvent was removed and the mixture was further purified by silica column chromatography to give intermediate **e26**.

#### 4.2.5 General procedures for the preparation of products **1a–1x, 2a–2k, 3a–3j**

##### Preparation of **1a–1x** (except for **1i**), **2a–2i** and **3c–3d**

Intermediate **e1** (2.3 mmol, 0.8 equiv) and another 10 mL of acetone were added to the solution of intermediate **b1** *in situ*, heated to reflux. TLC was used to monitor the reaction until the reaction was completed. Upon completion, the solvent was removed under reduced pressure. The crude solid product was washed with water and then recrystallized with methanol to give pure product **1a**. The products **1b–1h, 1j–1x, 2a–2i, 3c–3d** were prepared via the same method as that for **1a**. The products, which could not be recrystallized, were purified by silica column chromatography.

##### Preparation of **2j–2k**

To a solution of **2h** (1.25g, 2.82 mmol) in 15 mL of methanol, 2.0 M sodium hydroxide aqueous solution (2.8 mL, 5.6 mmol) was added. Then the solution was refluxed for 4 h. The reaction was monitored by TLC. Upon completion, the reaction mixture was acidified and precipitated with dilute hydrochloride solution. The crude solid product was recrystallized from methanol/H_2_O and dried to give **2j**. The preparation method of **2k** was the same as that of **2j**, except for **2i** as the precursor.

##### Preparation of other products

The detailed preparation methods for **1i, 3a–3b, 3e–3j** were described individually, along with their structural characterization data.

##### 4.2.5.1 N-(2-(4-butoxybenzoyl)hydrazine-1-carbonothioyl)acetamide (**1a**)

White solid, 398 mg (yield 46%). ^1^H NMR (500 MHz, DMSO*-d_6_*) δ 12.11 (s, 1H), 11.57 (s, 1H), 10.89 (s, 1H), 7.85 (d, *J* = 8.8 Hz, 2H), 7.03 (d, *J* = 8.9 Hz, 2H), 4.04 (t, *J* = 6.5 Hz, 2H), 2.14 (s, 3H), 1.77–1.64 (m, 2H), 1.44 (h, *J* = 7.4 Hz, 2H), 0.94 (t, *J* = 7.4 Hz, 3H). ^13^C NMR (125 MHz, DMSO*-d_6_*) δ 180.68, 172.50, 164.42, 162.15, 130.06, 124.33, 114.62, 67.91, 31.07, 24.00, 19.17, 14.17. HRMS Calcd for C_14_H_19_N_3_O_3_S [M+Na^+^]: 332.1039; Found:332.1028.

##### 4.2.5.2 N-(2-(4-(n-pentoxy)benzoyl)hydrazine-1-carbonothioyl)acetamide (**1b**)

White solid, 360 mg (yield 40%). ^1^H NMR (500 MHz, DMSO*-d_6_*) δ 12.10 (s, 1H), 11.54 (s, 1H), 10.85 (s, 1H), 7.84 (d, *J* = 8.3 Hz, 2H), 7.02 (d, *J* = 8.3 Hz, 2H), 4.03 (t, *J* = 6.5 Hz, 2H), 2.13 (s, 3H), 1.72 (t, *J* = 7.1 Hz, 2H), 1.36 (m, 4H), 0.89 (t, *J* = 6.6 Hz, 3H). ^13^C NMR (125 MHz, DMSO*-d_6_*) δ 180.32, 172.18, 164.11, 161.85, 129.75, 124.05, 114.33, 67.91, 28.42, 27.83, 23.71, 22.05, 14.10. HRMS Calcd for C_15_H_21_N_3_O_3_S [M+H^+^]: 324.1377; Found: 324.1367. HPLC purity: > 98% (cf. **Fig. S7**).

##### 4.2.5.3 N-(2-(4-(n-hexoxy)benzoyl)hydrazine-1-carbonothioyl)acetamide (**1c**)

White solid, 300 mg (yield 32%). ^1^H NMR (500 MHz, DMSO*-d_6_*) δ 12.10 (d, *J* = 2.8 Hz, 1H), 11.54 (s, 1H), 10.85 (d, *J* = 3.0 Hz, 1H), 7.84 (d, *J* = 8.4 Hz, 2H), 7.02 (d, *J* = 8.4 Hz, 2H), 4.03 (t, *J* = 6.5 Hz, 2H), 2.13 (s, 3H), 1.71 (p, *J* = 6.9 Hz, 2H), 1.41 (t, *J* = 7.5 Hz, 2H), 1.30 (p, *J* = 3.8 Hz, 4H), 0.87 (t, *J* = 6.75 Hz, 3H). ^13^C NMR (125 MHz, DMSO*-d_6_*) δ 180.33, 172.20, 164.12, 161.86, 129.76, 124.05, 114.34, 67.93, 31.15, 28.69, 25.31, 23.70, 22.24, 14.09. HRMS Calcd for C_16_H_23_N_3_O_3_S [M+H^+^]: 338.1533; Found: 338.1522.

##### 4.2.5.4 N-(2-(4-(n-undecoxy)benzoyl)hydrazine-1-carbonothioyl)acetamide (**1d**)

White solid, 364 mg (yield 32%). ^1^H NMR (500 MHz, DMSO*-d_6_*) δ 12.10 (s, 1H), 11.54 (s, 1H), 10.85 (s, 1H), 7.84 (d, *J* = 8.4 Hz, 2H), 7.02 (d, *J* = 8.5 Hz, 2H), 4.02 (t, *J* = 6.5 Hz, 2H), 2.13 (s, 3H), 1.71 (p, *J* = 6.9 Hz, 2H), 1.39 (q, *J* = 7.4 Hz, 2H), 1.30–1.19 (m, 14H), 0.84 (t, *J* = 6.7 Hz, 3H). ^13^C NMR (125 MHz, DMSO*-d_6_*) δ 180.33, 172.20, 164.12, 161.87, 129.76, 124.04, 114.33, 67.92, 32.12, 29.18, 28.92, 28.89, 28.72, 25.62, 23.70, 22.29, 14.15. HRMS Calcd for C_21_H_33_N_3_O_3_S [M+H^+^]: 408.2315; Found: 408.2306.

##### 4.2.5.5 N-(2-(4-((4-methylpentyl)oxy)benzoyl)hydrazine-1-carbonothioyl)acetamide (**1e**)

White solid, 112 mg (yield 32%). ^1^H NMR (500 MHz, DMSO*-d_6_*) δ 12.11 (d, *J* = 2.6 Hz, 1H), 11.55 (s, 1H), 10.87 (d, *J* = 2.7 Hz, 1H), 7.85 (d, *J* = 8.4 Hz, 2H), 7.03 (d, *J* = 8.4 Hz, 2H), 4.03 (t, *J* = 6.5 Hz, 2H), 2.14 (s, 3H), 1.73 (m, 2H), 1.59 (m, 1H), 1.31 (m, 2H), 0.89 (d, 6H). ^13^C NMR (125 MHz, DMSO*-d_6_*) δ 180.55, 172.53, 164.44, 162.18, 130.05, 124.35, 114.61, 68.50, 35.11, 27.73, 26.95, 23.99, 22.91. HRMS calcd for C_16_H_23_N_3_O_3_S [M+H^+^]: 338.1538, found: 338.1519.

##### 4.2.5.6 N-(2-(4-(3-methoxypropoxy)benzoyl)hydrazine-1-carbonothioyl)acetamide (**1f**)

White solid, 250 mg (yield 48%). ^1^H NMR (400 MHz, DMSO*-d_6_*) δ 12.11 (s, 1H), 11.56 (s, 1H), 10.89 (s, 1H), 7.86 (d, *J* = 8.5 Hz, 2H), 7.03 (d, *J* = 7.9 Hz, 2H), 4.09 (t, *J* = 6.0 Hz, 2H), 3.48 (d, *J* = 6.6 Hz, 2H), 3.24 (s, 3H), 2.14 (s, 3H), 1.96 (t, *J* = 8.2 Hz, 2H). ^13^C NMR (100 MHz, DMSO*-d_6_*) δ 180.64, 172.51, 164.43, 162.05, 130.08, 124.47, 114.63, 68.84, 65.38, 58.43, 29.29, 23.99. HRMS calcd for C_14_H_19_N_3_O_4_S [M+H^+^]: 326.1175, found: 326.1154.

##### 4.2.5.7 N-(2-(3-(n-pentoxy)benzoyl)hydrazine-1-carbonothioyl)acetamide (**1g**)

White solid, 282 mg (yield 36%). ^1^H NMR (400 MHz, DMSO*-d_6_*) δ 12.11 (s, 1H), 11.61 (s, 2H), 11.02 (s, 1H), 7.45 (m, 3H), 7.16 (d, *J* = 8.0 Hz, 1H), 4.03 (t, *J* = 6.5 Hz, 2H), 2.17 (s, 3H), 1.75 (p, *J* = 6.4 Hz, 2H), 1.40 (m, 4H), 0.92 (t, *J* = 7.0 Hz, 3H). ^13^C NMR (100 MHz, DMSO*-d_6_*) δ 181.08, 172.49, 164.75, 159.06, 133.85, 130.14, 120.24, 118.74, 113.78, 68.12, 28.78, 28.17, 23.99, 22.36, 14.38. HRMS calcd for C_15_N_21_N_3_O_3_S [M+H^+^]: 324.1382, found: 324.1364.

##### 4.2.5.8 N-(2-(3,4-bis(n-pentoxy)benzoyl)hydrazine-1-carbonothioyl)acetamide (**1h**)

White solid, 400 mg (yield 43%). ^1^H NMR (500 MHz, DMSO*-d_6_*) δ 12.08 (s, 1H), 11.54 (s, 1H), 10.84 (s, 1H), 7.49 (d, *J* = 8.3 Hz, 1H), 7.46 (s, 1H), 7.05 (d, *J* = 8.4 Hz, 1H), 4.00 (dt, *J* = 12.8, 6.3 Hz, 4H), 2.13 (s, 3H), 1.72 (p, *J* = 6.8 Hz, 4H), 1.41 (m, 4H), 1.35 (m, 4H), 0.89 (t, *J* = 7.2 Hz, 6H). ^13^C NMR (125 MHz, DMSO*-d_6_*) δ 180.48, 172.16, 164.13, 152.05, 148.08, 124.17, 121.49, 112.86, 112.65, 68.68, 68.45, 28.56, 28.46, 27.93, 27.90, 23.69, 22.05, 22.03, 14.10. HRMS Calcd for C_20_H_31_N_3_O_4_S [M+H^+^]: 410.2108; Found: 410.2096.

##### 4.2.5.9 N-(2-(3-hydroxy-4-(n-pentoxy)benzoyl)hydrazine-1-carbonothioyl)acetamide (**1i**)

To a mixture of methyl 3,4-dihydroxybenzoate (1.0 g, 5.9 mmol) and potassium carbonate (2.7 g, 19.7 mmol) in acetone (60 mL), 1-bromo-*n*-pentane (1.8 mL, 14.5 mmol) was added. The reaction mixture was heated to 60 C, carefully monitored via TLC. When the partial alkylation finished, the solvent was removed. Water was added and ethyl acetate (30 mL × 3) was used to extract the aqueous phase. The combined organic layer was dried over anhydrous sodium sulfate. The solvent was removed and recrystallization from methanol was performed to afford the product.

White solid, 700 mg (yield 49%). ^1^H NMR (500 MHz, DMSO*-d_6_*) δ 12.11 (s, 1H), 11.53 (s, 1H), 10.74 (s, 1H), 9.19 (s, 1H), 7.43–7.24 (m, 2H), 6.99 (d, *J* = 8.4 Hz, 1H), 4.01 (t, *J* = 6.6 Hz, 2H), 2.13 (s, 3H), 1.73 (t, *J* = 7.2 Hz, 2H), 1.37 (m, 4H), 0.89 (t, *J* = 7.1 Hz, 3H). ^13^C NMR (125 MHz, DMSO*-d_6_*) δ 180.06, 172.22, 164.30, 150.51, 146.50, 124.30, 119.63, 115.07, 112.49, 68.40, 28.52, 27.81, 23.71, 22.11, 14.13. HRMS Calcd for C_15_H_21_N_3_O_4_S [M+H^+^]: 340.1326; Found: 340.1316.

##### 4.2.5.10 N-(2-(3,4-dihydroxybenzoyl)hydrazine-1-carbonothioyl)acetamide (**1j**)

Pale yellow solid, 111 mg (yield 30%). ^1^H NMR (400 MHz, DMSO*-d_6_*) δ 12.14 (s, 1H), 11.55 (s, 1H), 10.70 (s, 1H), 9.72 (s, 1H), 9.32 (s, 1H), 7.33 (s, 1H), 7.28 (d, *J* = 8.2 Hz, 1H), 6.84 (d, *J* = 8.2 Hz, 1H), 2.16 (s, 3H). ^13^C NMR (100 MHz, DMSO*-d_6_*) δ 179.62, 172.09, 164.25, 149.37, 145.04, 122.82, 119.73, 115.33, 115.15, 23.53. HRMS calcd for C_10_H_11_N_3_O_4_S [M+H^+^]: 270.0549, found: 270.0531.

##### 4.2.5.11 N-(2-(2-methyl-4-(n-pentoxy)benzoyl)hydrazine-1-carbonothioyl)acetamide (**1k**)

White solid, 200 mg (yield 43%). ^1^H NMR (500 MHz, DMSO*-d_6_*) δ 12.05 (s, 1H), 11.55 (s, 1H), 10.64 (s, 1H), 7.51 (d, *J* = 8.2 Hz, 1H), 6.82 (d, *J* = 9.0 Hz, 2H), 3.99 (t, *J* = 6.9 Hz, 2H), 2.39 (s, 3H), 2.14 (s, 3H), 1.71 (m, 2H), 1.36 (m, 4H), 0.90 (t, *J* = 7.0 Hz, 3H). ^13^C NMR (125 MHz, DMSO*-d_6_*) δ 180.80, 172.49, 166.70, 160.57, 139.53, 130.32, 126.06, 117.13, 111.57, 67.97, 28.76, 28.14, 24.00, 22.34, 20.37, 14.13. HRMS calcd for C_16_H_23_N_3_O_3_S [M+Na^+^]: 360.1358, found: 360.1340.

##### 4.2.5.12 N-(2-(2,4-bis(n-pentoxy)benzoyl)hydrazine-1-carbonothioyl)acetamide (**1l**)

White solid, 400 mg (yield 43%). ^1^H NMR (400 MHz, DMSO*-d_6_*) δ 13.53 (d, *J* = 8.5 Hz, 1H), 11.67 (s, 1H), 11.26 (d, *J* = 8.6 Hz, 1H), 7.97 (d, *J* = 8.6 Hz, 1H), 6.69 (d, *J* = 8.5 Hz, 2H), 4.21 (t, *J* = 6.5 Hz, 2H), 4.04 (t, *J* = 6.6 Hz, 2H), 2.15 (s, 3H), 1.96 (p, *J* = 6.8 Hz, 2H), 1.80–1.62 (m, 2H), 1.56–1.26 (m, 8H), 0.90 (t, *J* = 5.8 Hz, 6H). ^13^C NMR (100 MHz, DMSO*-d_6_*) δ 172.95, 170.05, 164.07, 159.26, 159.00, 133.32, 110.45, 107.42, 100.01, 70.11, 68.46, 28.86, 28.72, 28.48, 28.10, 25.58, 23.83, 22.38, 22.35, 14.32. HRMS Calcd for C_20_H_31_N_3_O_4_S [M+H^+^]: 410.2108; Found: 410.2095.

##### 4.2.5.13 N-(2-(4-(cyclohexoxy)benzoyl)hydrazine-1-carbonothioyl)acetamide (**1m**)

White solid, 114 mg (yield 22%). ^1^H NMR (400 MHz, DMSO*-d_6_*) δ 12.13 (s, 1H), 11.58 (s, 1H), 10.88 (s, 1H), 7.86 (d, *J* = 8.4 Hz, 2H), 7.05 (d, *J* = 8.3 Hz, 2H), 4.48 (s, 1H), 2.16 (s, 3H), 1.95 (s, 2H), 1.74 (s, 2H), 1.65–1.13 (m, 6H). ^13^C NMR (100 MHz, DMSO*-d_6_*) δ 180.63, 172.50, 164.42, 160.90, 130.09, 124.14, 115.60, 74.91, 31.62, 25.51, 23.99, 23.57. HRMS calcd for C_16_H_21_N_3_O_3_S [M+Na^+^]: 358.1201, found: 358.1186.

##### 4.2.5.14 N-(2-(4-phenoxybenzoyl)hydrazine-1-carbonothioyl)acetamide (**1n**)

White solid, 321 mg (yield 35%). ^1^H NMR (500 MHz, DMSO*-d_6_*) δ 12.13 (s, 1H), 11.60 (s, 1H), 11.01 (s, 1H), 7.95 (d, *J* = 8.4 Hz, 2H), 7.48 (t, *J* = 7.7 Hz, 2H), 7.26 (s, 1H), 7.13 (dd, *J* = 19.4, 8.2 Hz, 4H), 2.18 (s, 3H).^13^C NMR (125 MHz, DMSO*-d_6_*) δ 180.43, 172.03, 163.78, 160.20, 155.40, 130.32, 129.96, 126.55, 124.50, 119.69, 117.45, 23.54. HRMS calcd for C_16_H_15_N_3_O_3_S [M+Na^+^]: 352.0726, found: 352.0711.

##### 4.2.5.15 N-(2-(5-(n-pentoxy)picolinoyl)hydrazine-1-carbonothioyl)acetamide (**1o**)

White solid, 107 mg (yield 15%). ^1^H NMR (400 MHz, DMSO*-d_6_*) δ 12.91 (d, *J* = 6.2 Hz, 1H), 11.73 (s, 1H), 11.00 (d, *J* = 6.0 Hz, 1H), 8.39 (s, 1H), 8.03 (d, *J* = 8.7 Hz, 1H), 7.59 (dd, *J* = 8.6, 2.8 Hz, 1H), 4.14 (t, *J* = 6.6 Hz, 2H), 2.17 (s, 3H), 1.77 (p, *J* = 6.7 Hz, 2H), 1.39 (m, 4H), 0.91 (t, *J* = 7.0 Hz, 3H). ^13^C NMR (125 MHz, DMSO*-d_6_*) δ 175.24, 173.02, 160.03, 158.20, 140.25, 138.04, 124.33, 121.89, 68.95, 28.56, 27.99, 23.94, 22.30, 14.36. HRMS calcd for C_14_H_20_N_4_O_3_S [M+H^+^]: 325.1329, found: 325.1334.

##### 4.2.5.16 N-(2-(2-(4-(n-pentoxy)phenyl)acetyl)hydrazine-1-carbonothioyl)acetamide (**1p**)

White solid, 300 mg (yield 42%). ^1^H NMR (500 MHz, DMSO*-d_6_*) δ 12.35 (s, 1H), 7.21 (d, *J* = 8.0 Hz, 2H), 6.87 (d, *J* = 8.1 Hz, 2H), 4.23 (s, 2H), 3.92 (t, *J* = 6.6 Hz, 2H), 2.12 (s, 3H), 1.68 (p, *J* = 6.7 Hz, 2H), 1.34 (tt, *J* = 14.6, 7.6 Hz, 4H), 0.88 (t, *J* = 7.0 Hz, 3H). ^13^C NMR (125 MHz, DMSO*-d_6_*) δ 168.63, 164.30, 158.77, 157.87, 130.07, 129.73, 114.82, 114.82, 67.56, 2856, 27.90, 22.55, 22.08, 14.12. HRMS Calcd for C_16_H_23_N_3_O_3_S [M+H^+^]: 338.1533; Found: 338.1523.

##### 4.2.5.17 N-(2-(3-(4-(n-pentoxy)phenyl)propanoyl)hydrazine-1-carbonothioyl)acetamide (**1q**)

White solid, 300 mg (yield 43%). ^1^H NMR (500 MHz, DMSO*-d_6_*) δ 12.35 (d, *J* = 4.6 Hz, 1H), 11.46 (s, 1H), 10.76 (d, *J* = 4.6 Hz, 1H), 7.11 (d, *J* = 8.0 Hz, 2H), 6.81 (d, *J* = 8.0 Hz, 2H), 3.90 (t, *J* = 6.7 Hz, 2H), 2.78 (d, *J* = 7.8 Hz, 2H), 2.09 (s, 3H), 1.67 (t, *J* = 7.1 Hz, 2H), 1.34 (m, 4H), 0.88 (t, *J* = 7.1 Hz, 3H). ^13^C NMR (125 MHz, DMSO*-d_6_*) δ 176.58, 172.30, 168.85, 157.18, 132.72, 129.59, 129.36, 114.45, 67.49, 29.96, 28.62, 27.93, 23.64, 22.09, 14.13. HRMS Calcd for C_17_H_25_N_3_O_3_S [M+H^+^]: 352.1689; Found: 352.1681.

##### 4.2.5.18 N-(2-([1,1’-biphenyl]-4-carbonyl)hydrazine-1-carbonothioyl)acetamide (**1r**)

White solid, 220 mg (yield 47%). ^1^H NMR (500 MHz, DMSO*-d_6_*) δ 12.13 (s, 1H), 11.59 (s, 1H), 11.09 (s, 1H), 7.99 (d, *J* = 8.1 Hz, 2H), 7.83 (d, *J* = 8.0 Hz, 2H), 7.75 (d, *J* = 7.7 Hz, 2H), 7.51 (t, *J* = 7.6 Hz, 2H), 7.43 (d, *J* = 7.4 Hz, 1H), 2.16 (s, 3H). ^13^C NMR (125 MHz, DMSO*-d_6_*) δ 180.92, 172.52, 164.67, 144.07, 139.51, 131.30, 129.55, 128.84, 128.70, 127.42, 127.18, 24.03. HRMS calcd for C_16_H_15_N_3_O_2_S [M+H^+^]: 314.0963, found: 314.0947.

##### 4.2.5.19 N-(2-([1,1’-biphenyl]-3-carbonyl)hydrazine-1-carbonothioyl)acetamide (**1s**)

White solid, 196 mg (yield 42%). ^1^H NMR (500 MHz, DMSO*-d_6_*) δ 12.12 (s, 1H), 11.60 (s, 1H), 11.17 (s, 1H), 8.19 (s, 1H), 7.89 (dd, J = 14.9, 7.7 Hz, 2H), 7.75 (d, J = 7.4 Hz, 2H), 7.62 (t, *J* = 7.7 Hz, 1H), 7.52 (t, *J* = 7.5 Hz, 2H), 7.43 (d, *J* = 7.3 Hz, 1H), 2.16 (s, 3H). ^13^C NMR (125 MHz, DMSO*-d_6_*) δ 180.65, 172.03, 164.46, 140.32, 139.34, 132.76, 130.23, 129.29, 129.07, 127.94, 126.87, 126.83, 125.84, 23.56. HRMS calcd for C_16_H_15_N_3_O_2_S [M+Na^+^]: 336.0777, found: 336.0775.

##### 4.2.5.20 N-(2-(2-naphthoyl)hydrazine-1-carbonothioyl)acetamide (**1t**)

White solid, 222 mg (yield 46%). ^1^H NMR (500 MHz, DMSO*-d_6_*) δ 12.18 (s, 1H), 11.60 (s, 1H), 11.19 (s, 1H), 8.53 (s, 1H), 8.04 (dt, *J* = 17.6, 8.6 Hz, 4H), 7.95 (d, *J* = 8.6 Hz, 1H), 7.64 (m, 2H), 2.16 (s, 3H). ^13^C NMR (100 MHz, DMSO) δ 180.90, 172.59, 165.09, 135.01, 132.53, 129.88, 129.50, 128.86, 128.67, 128.57, 128.24, 127.49, 124.63, 24.07. HRMS calcd for C_14_H_13_N_3_O_2_S [M+Na^+^]: 392.0621, found: 392.0620.

##### 4.2.5.21 N-(2-(4-cyclopropylbenzoyl)hydrazine-1-carbonothioyl)acetamide (**1u**)

White solid, 254 mg (yield 40%). ^1^H NMR (500 MHz, DMSO*-d_6_*) δ 12.11 (s, 1H), 11.56 (s, 1H), 10.92 (s, 1H), 7.78 (d, *J* = 7.8 Hz, 2H), 7.20 (d, *J* = 7.8 Hz, 2H), 2.14 (s, 3H), 2.04–1.93 (m, 1H), 1.02 (d, *J* = 7.9 Hz, 2H), 0.75 (d, *J* = 5.2 Hz, 2H). ^13^C NMR (125 MHz, DMSO*-d_6_*) δ 180.98, 172.69, 164.94, 149.34, 129.49, 128.34, 125.83, 24.19, 15.88, 10.98. HRMS calcd for C_13_H_15_N_3_O_2_S [M+H^+^]: 278.0958, found: 278.0951.

##### 4.2.5.22 N-(2-(4-cyclohexylbenzoyl)hydrazine-1-carbonothioyl)acetamide (**1v**)

White solid, 250 mg (yield 45%). ^1^H NMR (500 MHz, DMSO*-d_6_*) δ 12.10 (s, 1H), 11.56 (s, 1H), 10.92 (s, 1H), 7.81 (d, *J* = 8.0 Hz, 2H), 7.36 (d, *J* = 8.0 Hz, 2H), 2.62–2.53 (m, 1H), 2.14 (s, 3H), 1.80 (dd, *J* = 11.9, 4.7 Hz, 4H), 1.78 (d, *J* = 13.0 Hz, 1H), 1.48–1.34 (m, 4H), 1.37 (m, 1H). ^13^C NMR (100 MHz, DMSO) δ 180.83, 172.56, 164.91, 152.45, 130.15, 128.29, 127.30, 44.24, 34.17, 26.75, 26.03, 24.04. HRMS calcd for C_16_H_21_N_3_O_2_S [M+Na^+^] 342.1247, found: 342.1242.

##### 4.2.5.23 N-(2-(4-(4-ethylcyclohexyl)benzoyl)hydrazine-1-carbonothioyl)acetamide (**1w**)

White solid, 128 mg (yield 39%). ^1^H NMR (500 MHz, DMSO*-d_6_*) δ 12.14 (s, 1H), 11.60 (s, 1H), 10.96 (s, 1H), 7.85 (d, *J* = 7.9 Hz, 2H), 7.39 (d, *J* = 7.9 Hz, 2H), 2.58 (d, *J* = 12.2 Hz, 1H), 2.18 (s, 3H), 1.86 (t, *J* = 12.6 Hz, 4H), 1.49 (qd, *J* = 12.9, 12.3, 3.5 Hz, 2H), 1.26 (dt, *J* = 20.2, 10.0 Hz, 3H), 1.07 (q, *J* = 11.0, 10.3 Hz, 2H), 0.92 (t, *J* = 7.2 Hz, 3H). ^13^C NMR (125 MHz, DMSO*-d_6_*) δ 180.30, 172.03, 164.37, 151.76, 129.63, 127.75, 126.80, 43.80, 36.28, 33.54, 32.92, 23.53, 19.49, 14.28. HRMS calcd for C_18_H_25_N_3_O_2_S [M+H^+^]: 348.1740, found: 348.1733.

##### 4.2.5.24 N-(2-(4-morpholinobenzoyl)hydrazine-1-carbonothioyl)acetamide (**1x**)

White solid, 298 mg (yield 39%). ^1^H NMR (400 MHz, DMSO*-d_6_*) δ 12.10 (s, 1H), 11.57 (s, 1H), 10.77 (s, 1H), 7.82 (d, *J* = 8.4 Hz, 2H), 7.03 (d, *J* = 8.5 Hz, 2H), 3.76 (s, 4H), 3.27 (s, 4H), 2.17 (s, 3H). ^13^C NMR (100 MHz, DMSO) δ 180.17, 172.52, 164.50, 153.95, 129.51, 121.50, 113.80, 66.36, 47.60, 23.99. HRMS calcd for C_14_H_18_N_4_O_3_S [M+H^+^]: 323.1173, found: 323.1164.

##### 4.2.5.25 N-(2-(4-(n-pentoxy)benzoyl)hydrazine-1-carbonothioyl)propionamide (**2a**)

White solid, 198 mg (yield 30%). ^1^H NMR (500 MHz, DMSO*-d_6_*), δ 12.15 (s, 1H), 11.55 (s, 1H), 10.89 (s, 1H), 7.89 (d, *J*=8.0Hz, 2H), 7.06 (d, *J*=8.0Hz, 2H), 4.07 (t, *J* = 6.0 Hz, 2H), 2.48 (t, *J*=7.5 Hz, 2H), 1.76 (m, 2H), 1.40 (m, 4H), 1.08 (t, *J*=7.5 Hz, 3H), 0.93 (t, *J*=7.5 Hz, 3H). ^13^C NMR (125 MHz, DMSO*-d_6_*) δ 180.72, 175.91, 164.44, 162.16, 130.06, 124.36, 114.63, 68.21, 29.38, 28.71, 28.12, 22.34, 14.39. HRMS calcd for C_16_H_23_N_3_O_3_S [M+Na^+^]: 360.1352, found: 360.1333.

##### 4.2.5.26 N-(2-(4-(n-pentoxy)benzoyl)hydrazine-1-carbonothioyl)butyramide (**2b**)

White solid, 86 mg (yield 13%). ^1^H NMR (500 MHz, DMSO*-d_6_)*, δ 12.18 (s, 1H), 11.55 (s, 1H), 10.89 (s, 1H), 7.89 (d, *J*=8.0Hz, 2H), 7.06 (d, *J*=8.0Hz, 2H), 4.07 (t, *J* = 6.0 Hz, 2H), 2.45 (t, *J*=7.5Hz, 2H), 1.76 (m, 2H), 1.62 (m, 2H), 1.41 (m, 3H), 0.93 (m, 6H). ^13^C NMR (125 MHz, DMSO*-d_6_*) δ 180.44, 174.00, 164.42, 162.16, 130.06, 124.34, 114.63, 68.21, 34.52, 34.47, 28.71, 28.12, 27.72, 22.34, 14.39. HRMS calcd for C_16_H_23_N_3_O_3_S [M+H^+^]: 352.1690, found: 352.1694.

##### 4.2.5.27 N-(2-(4-(n-pentoxy)benzoyl)hydrazine-1-carbonothioyl)isobutyramide (**2c**)

White solid, 150 mg (yield 18%). ^1^H NMR (500 MHz, DMSO*-d_6_*), δ 12.18 (s,1H), 11.55 (s, 1H), 10.89 (s, 1H), 7.89 (d, *J* = 8.0 Hz, 2H), 7.06 (d, *J* = 8.0 Hz, 2H), 4.08 (t, *J* = 6.0 Hz, 2H), 2.83 (m, 1H), 1.76 (m, 2H), 1.41 (m, 4H), 1.12 (m, 6H), 0.93 (t, *J* = 7.5 Hz, 3H). ^13^C NMR (125 MHz, DMSO*-d_6_*) δ 180.86, 179.05, 164.43, 162.17, 130.06, 124.34, 114.63, 68.21, 34.63, 28.71, 28.12, 22.34, 19.39, 14.39. HRMS calcd for C_17_H_25_N_3_O_3_S [M+H^+^]: 352.1690, found: 352.1693.

##### 4.2.5.28 N-(2-(4-(n-pentoxy)benzoyl)hydrazine-1-carbonothioyl)cyclopropanecarboxamide (**2d**)

White solid, 100 mg (yield 13%). ^1^H NMR (500 MHz, DMSO*-d_6_*), δ 12.15 (s, 1H), 11.90 (s, 1H), 10.89 (s, 1H), 7.89 (d, *J*=8.0Hz, 2H), 7.06 (d, *J*=8.0Hz, 2H), 4.07 (t, *J* = 6.0 Hz, 2H), 2.11 (s, 1H), 1.76 (m, 2H), 1.42 (m, 4H), 1.01–0.93 (m, 7H). ^13^C NMR (125 MHz, DMSO*-d_6_*) δ 180.37, 175.63, 164.43, 162.16, 130.04, 124.33, 114.62, 68.21, 28.71, 28.12, 22.34, 14.38, 9.84. HRMS calcd for C_17_H_25_N_3_O_3_S [M+H^+^]: 350.1533, found: 350.1537.

##### 4.2.5.29 N-(2-(4-(n-pentoxy)benzoyl)hydrazine-1-carbonothioyl)cyclopentanecarboxamide (**2e**)

White solid, 270 mg (yield 32%). ^1^H NMR (500 MHz, DMSO*-d_6_*), δ 12.23 (s, 1H), 11.59 (s, 1H), 10.91 (s, 1H), 7.91 (d, *J* = 8.0 Hz, 2H), 7.09 (d, *J* = 8.0 Hz, 2H), 4.09 (t, *J* = 6.5 Hz, 2H), 3.03 (m, 1H), 1.92 (m, 2H), 1.60–1.79 (m, 8H), 1.41 (m, 4H), 0.96 (t, *J* = 7.0 Hz, 3H). ^13^C NMR (125 MHz, DMSO*-d_6_*) δ 180.73, 178.31, 164.42, 162.16, 130.05, 124.35, 114.63, 68.21, 44.73, 30.40, 28.71, 28.12, 26.23, 22.34, 14.39. HRMS calcd for C_19_H_27_N_3_O_3_S [M+H^+^]: 378.1846, found: 378.1844.

##### 4.2.5.30 N-(2-(4-(n-pentoxy)benzoyl)hydrazine-1-carbonothioyl)-3-phenylpropanamide (**2f**)

White solid, 690 mg (yield 77%). ^1^H NMR (400 MHz, DMSO*-d_6_*) δ 12.06 (s, 1H), 11.79 (s, 1H), 10.89 (s, 1H), 7.86 (d, *J* = 7.9 Hz, 2H), 7.35 (d, *J* = 6.4 Hz, 5H), 7.04 (d, *J* = 8.2 Hz, 2H), 4.05 (d, *J* = 5.8 Hz, 2H), 3.82 (d, *J* = 4.9 Hz, 2H), 1.75 (d, *J* = 6.5 Hz, 2H), 1.39 (s, 4H), 0.92 (d, *J* = 5.2 Hz, 3H). ^13^C NMR (100 MHz, DMSO*-d_6_*) δ 180.66, 172.91, 164.43, 162.17, 134.82, 130.06, 129.86, 128.93, 127.48, 124.33, 114.63, 68.21, 42.59, 28.70, 28.11, 22.34, 14.38. HRMS calcd for C_21_H_25_N_3_O_3_S [M+H^+^]: 400.1690, found: 400.1691.

##### 4.2.5.31 N-(2-(4-(n-pentoxy)benzoyl)hydrazine-1-carbonothioyl)thiophene-2-carboxamide (**2g**)

Pale yellow solid, 400 mg (yield 46%). ^1^H NMR (500 MHz, DMSO*-d_6_*), δ 12.26 (s, 1H), 11.82 (s, 1H), 10.97 (s, 1H), 8.41 (d, *J* = 2.8 Hz, 1H), 8.09 (d, *J* = 2.8 Hz, 1H), 7.92 (d, *J* = 8.4 Hz, 2H), 7.28 (dd, *J* = 2.8 Hz, *J* = 4.1 Hz, 1H), 7.07 (d, *J* = 8.4 Hz, 2H), 4.07 (t, *J* = 6.3 Hz, 2H), 1.80–1.74 (m, 2H), 1.41 (m, 4H), 0.93 (t, *J* = 6.9 Hz, 3H). ^13^C NMR (100 MHz, DMSO*-d_6_*) δ 180.53, 164.44, 162.18, 162.11, 136.91, 135.92, 133.29, 130.08, 129.26, 124.37, 114.65, 68.22, 28.72, 28.12, 22.35, 14.39. HRMS calcd for C_18_H_21_N_3_O_3_S_2_ [M+H^+^]: 392.1097, found: 392.1110.

##### 4.2.5.32 methyl 4-((2-(4-(n-pentoxy)benzoyl)hydrazine-1-carbonothioyl)carbamoyl)benzoate (**2h**)

White solid, 1250 mg (yield 63%). ^1^H NMR (500 MHz, DMSO*-d_6_*), δ 12.35 (s, 1H), 11.99 (s, 1H), 11.01 (s, 1H), 8.12 (s, 4H), 7.92 (d, *J* = 8.0 Hz, 2H), 7.07 (d, *J* = 8.0 Hz, 2H), 4.08 (t, *J* = 6.5 Hz, 2H), 3.94 (s, 3H), 1.77 (m, 2H), 1.40 (m, 4H), 0.94 (t, *J* = 7.0Hz, 3H). ^13^C NMR (125 MHz, DMSO*-d_6_*) δ 180.37, 167.53, 165.97, 164.44, 162.20, 136.56, 133.65, 130.10, 129.62, 129.51, 124.35, 114.65, 68.22, 53.13, 28.70, 28.11, 22.33, 14.36. HRMS calcd for C_22_H_25_N_3_O_5_S [M+H^+^]: 444.1588, found: 444.1591.

##### 4.2.5.33 methyl 4-oxo-4-(2-(4-(n-pentoxy)benzoyl)hydrazine-1-carbothioamido)butanoate (**2i**)

White solid, 500 mg (yield 28%). ^1^H NMR (500 MHz, DMSO*-d_6_*), δ 12.03 (s, 1H), 11.66 (s, 1H), 10.88 (s, 1H), 7.89 (d, *J* = 8.0 Hz, 2H), 7.06 (d, *J* = 8.0 Hz, 2H), 4.07 (t, *J* = 6.5 Hz, 2H), 3.64 (s, 3H), 2.77 (t, *J* = 6.5 Hz, 2H), 2.64 (t, *J* = 6.5 Hz, 2H), 1.77 (m, 2H), 1.41 (m, 4H), 0.94 (t, *J* = 7.0 Hz, 3H). ^13^C NMR (125 MHz, DMSO) δ 180.60, 174.01, 173.24, 164.66, 162.34, 130.21, 124.50, 114.81, 68.38, 52.20, 31.09, 28.88, 28.30, 22.54, 14.59. HRMS calcd for C_18_H_25_N_3_O_5_S [M+H^+^]: 396.1588, found: 396.1589.

##### 4.2.5.34 4-((2-(4-(n-pentoxy)benzoyl)hydrazine-1-carbonothioyl)carbamoyl)benzoic acid (**2j**)

White solid, 610 mg (yield 51%). ^1^H NMR (500 MHz, DMSO*-d_6_*), δ 13.35 (s, 1H), 12.36 (s, 1H), 11.95 (s, 1H), 11.01 (s, 1H), 8.09 (m, 4H), 7.92 (d, *J* = 8.0 Hz, 2H), 7.08 (d, *J* = 8.0 Hz, 2H), 4.08 (t, *J* = 6.5 Hz, 2H), 1.77 (m, 2H), 1.41 (m, 4H), 0.94 (t, *J* = 7.0 Hz, 3H). ^13^C NMR (125 MHz, DMSO*-d_6_*) δ 180.42, 167.68, 167.03, 164.45, 162.20, 136.18, 134.96, 130.11, 129.64, 129.49, 124.35, 114.65, 68.22, 28.72, 28.12, 22.35, 14.38. HRMS calcd for C_21_H_23_N_3_O_3_S [M+H^+^]: 430.1431, found: 430.1433.

##### 4.2.5.35 4-oxo-4-(2-(4-(n-pentoxy)benzoyl)hydrazine-1-carbothioamido)butanoic acid (**2k**)

White solid, 251 mg (yield 52%). ^1^H NMR (500 MHz, DMSO*-d_6_*), δ 12.18 (s, 1H), 11.55 (s, 1H), 10.89 (s, 1H), 7.89 (d, *J* = 8.0 Hz, 2H), 7.06 (d, *J* = 8.0 Hz, 2H), 4.07 (t, *J* = 6.5 Hz, 2H), 2.71 (t, *J* = 6.5 Hz, 2H), 1.78 (m, 2H), 1.42 (m, 4H), 0.93 (t, *J* = 7.0 Hz, 3H). ^13^C NMR (125 MHz, DMSO*-d_6_*) δ 180.47, 174.12, 173.93, 164.41, 162.15, 130.06, 124.37, 114.62, 68.21, 31.06, 28.71, 28.44, 28.12, 22.34, 14.39. HRMS calcd for C_21_H_23_N_3_O_3_S [M+H^+^]: 382.1431, found: 382.1436.

##### 4.2.5.36 N-(5-(4-(n-hexoxy)phenyl)-1,3,4-oxadiazol-2-yl)acetamide (**3a**)

0.3 g (0.8 mmol) of **1c**, and 0.29 g (1.3 mmol) of KIO_3_ were added to 30 mL of water in a 100-mL flask, refluxed at 100 C for h. TLC was used for monitoring. While completed, the mixture was cooled, Then, the precipitate was filtered and purified with silica column chromatography to give solid product, **3a**, 0.25g.

Yellow solid, 250 mg (yield 93%). ^1^H NMR (500 MHz, DMSO*-d_6_*) δ 11.62 (s, 1H), 7.82 (d, *J* = 8.5 Hz, 2H), 7.11 (d, *J* = 8.5 Hz, 2H), 4.04 (t, *J* = 6.5 Hz, 2H), 2.15 (s, 3H), 1.72 (m, 2H), 1.41 (m, 2H), 1.31 (m, 4H), 0.88 (d, *J* = 6.7 Hz, 3H). ^13^C NMR (125 MHz, DMSO*-d_6_*) δ 161.36, 160.72, 157.11, 127.94, 115.78, 115.47, 68.01, 31.15, 28.67, 25.30, 23.53, 22.25, 14.10. HRMS Calcd for C_16_H_21_N_3_O_3_ [M+H^+^]: 304.1656, Found: 304.1649.

##### 4.2.5.37 N-(5-(4-(n-hexoxy)phenyl)-1,3,4-thiadiazol-2-yl)acetamide (**3b**)

0.3 g (0.8 mmol) of **1c** was dissolved in 4 mL of acetic acid, and the reaction was carried out at microwave irradiation (T = 125 °C, t = 5 min, 300 W). While completed, the mixture was cooled. Then the precipitate was filtered and purified by silica column chromatography to give solid product, **3b**, 0.20 g.

White solid, 200 mg (yield 70%). ^1^H NMR (500 MHz, DMSO*-d_6_*) δ 12.53 (s, 1H), 7.84 (d, *J* = 8.3 Hz, 2H), 7.04 (d, *J* = 8.3 Hz, 2H), 4.02 (t, *J* = 6.4 Hz, 2H), 2.19 (s, 3H), 1.72 (m, 2H), 1.41 (t, *J* = 7.5 Hz, 2H), 1.30 (m, 4H), 0.87 (d, *J* = 6.6 Hz, 3H). ^13^C NMR (125 MHz, DMSO*-d_6_*) δ 168.71, 161.68, 160.64, 157.83, 128.60, 122.74, 115.33, 67.91, 28.71, 25.31, 22.60, 22.25, 14.10. HRMS Calcd for C_16_H_21_N_3_O_2_S [M+H^+^]: 320.1427, Found: 320.1419.

##### 4.2.5.38 N-(2-(4-(n-hexoxy)benzoyl)-1-methylhydrazine-1-carbonothioyl)acetamide (**3c**)

White solid, 400 mg (yield 57%). ^1^H NMR (500 MHz, DMSO*-d_6_*) δ 10.68 (s, 1H), 10.55 (s, 1H), 7.77 (d, *J* = 8.3 Hz, 2H), 7.02 (d, *J* = 8.3 Hz, 2H), 4.03 (t, *J* = 6.5 Hz, 2H), 3.51 (s, 3H), 1.96 (s, 3H), 1.71 (m, 2H), 1.39 (d, *J* = 8.5 Hz, 2H), 1.30 (m, 4H), 0.87 (t, *J* = 6.4 Hz, 3H). ^13^C NMR (125 MHz, DMSO*-d_6_*) δ 181.95, 171.37, 165.04, 161.63, 129.74, 123.71, 114.42, 67.95, 42.91, 31.14, 28.67, 25.30, 23.77, 22.24, 14.10. HRMS Calcd for C_17_H_25_N_3_O_3_S [M+H^+^]: 352.1689; Found: 352.1679.

##### 4.2.5.39 N-(2-(4-(n-hexoxy)benzoyl)-2-methylhydrazine-1-carbonothioyl)acetamide (**3d**)

White solid, 400 mg (yield 57%). ^1^H NMR (500 MHz, DMSO*-d_6_*) δ 12.12 (s, 1H), 11.29 (s, 1H), 7.49 (d, *J* = 8.3 Hz, 2H), 6.86 (d, *J* = 8.3 Hz, 2H), 3.96 (t, *J* = 6.5 Hz, 2H), 3.13 (s, 3H), 2.01 (s, 3H), 1.68 (m, 2H), 1.39 (m, 2H), 1.29 (m, 4H), 0.86 (t, *J* = 6.8 Hz, 3H). ^13^C NMR (125 MHz, DMSO*-d_6_*) δ 181.22, 171.40, 160.54, 130.02, 127.09, 113.74, 67.98, 31.45, 29.05, 25.63, 23.91, 22.53, 14.38. HRMS Calcd for C_17_H_25_N_3_O_3_S [M+H^+^]: 352.1689; Found: 352.1679.

##### 4.2.5.40 N-ethyl-2-(4-(n-hexoxy)benzyl)hydrazine-1-carbothioamide (**3e**)

0.3 g (0.8 mmol) of **1c**, 0.3 g (8.0 mmol) of LiAlH_4_, and 30 mL of THF were added to a 100-mL three-neck flask, stirred under reflux with nitrogen gas protection. TLC plate (DCM: MeOH = 15:1) was used for monitoring. When completed, iced water was added to quench the reaction and the aqueous phase was extracted with ethyl acetate for three times. The combined organic layer was dried over anhydrous sodium sulfate and purified by silica column chromatography to give white solid.

White solid, 80 mg (yield 29%). ^1^H NMR (500 MHz, DMSO*-d_6_*) δ 8.52 (s, 1H), 7.72 (s, 1H), 7.24 (d, *J* = 8.1 Hz, 2H), 6.83 (d, *J* = 8.1 Hz, 2H), 5.16 (s, 1H), 3.91 (t, *J* = 6.5 Hz, 2H), 3.72 (d, *J* = 5.0 Hz, 2H), 3.37 (m, 2H), 1.67 (m, 2H), 1.39 (m, 2H), 1.29 (m, 4H), 0.94 (t, *J* = 7.1 Hz, 3H), 0.87 (t, *J* = 6.6 Hz, 3H). ^13^C NMR (125 MHz, DMSO*-d_6_*) δ 180.34, 158.04, 130.18, 129.80, 114.18, 67.51, 53.96, 37.60, 31.18, 28.82, 25.37, 22.25, 14.83, 14.10. HRMS Calcd for C_16_H_27_N_3_OS [M+H^+^]: 310.1948; Found: 310.1939.

##### 4.2.5.41 N-ethyl-2-(4-(n-hexoxy)benzoyl)hydrazine-1-carbothioamide (**3f**)

The same preparation method as **3e** mentioned above was used to give white solid **3f**. White solid, 40 mg (yield 14%). ^1^H NMR (500 MHz, DMSO*-d_6_*) δ 10.12 (s, 1H), 9.15 (s, 1H), 8.03 (s, 1H), 7.86 (d, *J* = 8.4 Hz, 2H), 6.99 (d, *J* = 8.4 Hz, 2H), 4.02 (t, *J* = 6.5 Hz, 2H), 3.45 (m, 2H), 1.70 (m, 2H), 1.40 (t, *J* = 7.6 Hz, 2H), 1.30 (m, 4H), 1.04 (t, *J* = 7.1 Hz, 3H), 0.87 (t, *J* = 6.7 Hz, 3H). ^13^C NMR (125 MHz, DMSO*-d_6_*) δ 165.62, 161.64, 129.90, 124.67, 114.06, 67.89, 38.63, 31.16, 28.69, 25.30, 22.25, 14.70, 14.10. HRMS Calcd for C_16_H_25_N_3_O_2_S [M+H^+^]: 324.1740; Found: 324.1732.

##### 4.2.5.42 N-acetyl-2-(4-(n-hexoxy)benzoyl)hydrazine-1-carboxamide (**3g**)

The aforementioned **b12**, 0.5 g (2.1 mmol, 0.8 equiv) of **e3**, and 20 mL of acetone were mixed and stirred under reflux. Upon completion, the solvent was removed and the crude solid was further purified by silica column chromatography to give white solid **3g**.

White solid, 40 mg (yield 6%). ^1^H NMR (500 MHz, DMSO*-d_6_*) δ 10.63 (s, 1H), 10.35 (s, 1H), 9.73 (s, 1H), 7.82 (d, *J* = 8.2 Hz, 2H), 7.00 (d, *J* = 8.6 Hz, 2H), 4.02 (t, *J* = 6.6 Hz, 2H), 2.08 (s, 2H), 1.71 (t, *J* = 7.4 Hz, 2H), 1.40 (d, *J* = 10.2 Hz, 2H), 1.30 (m, 4H), 0.87 (t, *J* = 7.6 Hz, 3H). ^13^C NMR (125 MHz, DMSO*-d_6_*) δ 172.36, 165.19, 161.70, 153.71, 129.57, 124.29, 114.28, 67.89, 31.16, 28.70, 25.31, 23.64, 22.25, 14.09. HRMS Calcd for C_16_H_23_N_3_O_4_ [M+H^+^]: 322.1761; Found: 322.1753.

##### 4.2.5.43 N-(methylsulfonyl)-2-(4-(n-pentoxy)benzoyl)hydrazine-1-carbothioamide (**3h**)

To the aforementioned oil **b13**, the intermediate **e2** (0.8 g, 3.6 mmol, 0.9 equiv) and THF (15 mL) was added. The reaction was allowed to stir at room temperature until completed. The solvent was removed and the crude product was washed with dichloromethane for 3 times to afford pale yellow solid **3h**.

Pale yellow solid, 800 mg (yield 62%). ^1^H NMR (500 MHz, DMSO*-d_6_*) δ 10.61 (s, 1H), 10.33 (s, 1H), 7.85 (d, *J* = 8.4 Hz, 2H), 7.01 (d, *J* = 6.2 Hz, 2H), 4.03 (t, *J* = 6.6 Hz, 2H), 3.46 (d, *J* = 5.3 Hz, 3H), 1.71 (m, 2H), 1.36 (m, 4H), 0.89 (t, *J* = 7.0 Hz, 3H). ^13^C NMR (125 MHz, DMSO*-d_6_*) δ 181.07, 164.67, 161.82, 130.04, 129.69, 114.36, 114.07, 67.93, 41.41, 28.44, 27.85, 22.08, 14.13. HRMS Calcd for C_14_H_21_N_3_O_4_S_2_ [M+H^+^]: 360.1046; Found: 360.1059.

##### 4.2.5.44 N-(2-acetylhydrazine-1-carbonothioyl)-4-(n-pentoxy)benzamide (**3i**)

4-(*n*-pentoxy)benzoic acid (1.0 g, 4.8 mmol) was added to 10 mL of thionyl chloride under 85 C for 3 h and the solvent was removed to afford **a11**. The mixture of **a11** (0.6 g, 2.6 mmol), potassium thiocyanate (0.27 g, 2.7 mmol) in 10 mL of acetone was stirred at room temperature for 1 h to afford **b11**. Then acetyl hydrazide (0.24 g, 3.2 mmol) was added to the aforementioned solution. The reaction was carried out under reflux for several hours until completion. The mixture was concentrated in vacuum and the residue was recrystallized from methanol to give white solid.

White solid, 400 mg (yield 47%). ^1^H NMR (500 MHz, DMSO*-d_6_*) δ 12.69 (d, *J* = 4.6 Hz, 1H), 11.46 (s, 1H), 10.81 (d, *J* = 4.6 Hz, 1H), 7.96 (d, *J* = 8.5 Hz, 2H), 7.02 (d, *J* = 8.6 Hz, 2H), 4.05 (t, *J* = 6.5 Hz, 2H), 1.97 (s, 3H), 1.73 (m, 2H), 1.36 (m, 4H), 0.89 (t, *J* = 7.0 Hz, 3H). ^13^C NMR (100 MHz, DMSO) δ 177.24, 167.67, 166.89, 163.18, 131.49, 123.84, 114.65, 68.38, 28.68, 28.09, 22.33, 20.81, 14.37. HRMS Calcd for C_15_H_21_N_3_O_3_S [M+H^+^]: 324.1376. found: 324.1363.

##### 4.2.5.45 2-(4-(n-hexoxy)benzoyl)hydrazine-1-carbothioamide (**3j**)

0.3 g (1.2 mmol) of **e3**, 0.17 g (1.2 mmol) of *S*-methylisothiourea, and 20 mL of water were added to a 50-mL flask and stirred under 100 C. TLC plate (DCM: MeOH = 15:1) was used for monitoring. When completed, the mixture was filtered to give crude solid, which was further purified by silica column chromatography to give white solid.

White solid, 200 mg (yield 57%). ^1^H NMR (500 MHz, DMSO*-d_6_*) δ 7.86 (d, *J* = 8.2 Hz, 2H), 6.95 (d, *J* = 7.9 Hz, 2H), 4.00 (t, *J* = 6.5 Hz, 2H), 1.75 (m, 2H), 1.41 (t, *J* = 7.7 Hz, 2H), 1.30 (m, 4H), 0.86 (t, *J* = 6.6 Hz, 3H). ^13^C NMR (125 MHz, DMSO*-d_6_*) δ 165.68, 161.66, 129.90, 125.06, 114.06, 67.88, 31.09, 28.41, 27.82, 22.05, 14.09. HRMS Calcd for C_15_H_21_N_3_O_3_S [M+H^+^]: 296.1427; Found: 296.1418.

### 4.3. Biology

#### 4.3.1. Bacterial growth inhibition (MIC) assay

Broth microdilution assay recommended by the Clinical and Laboratory Standards Institute [20] was used to determine MIC, with vancomycin as the positive control. The compound was dissolved in DMSO to make a stock solution. The bacterial inoculum, *S. aureus* ATCC 29213, was incubated overnight at 37 C and transferred to new Cation-Adjusted Mueller-Hinton broth (CAMHB) until bacterial growth reached the logarithmic phase. The bacteria were then diluted to 10^5^–10^6^ colony forming units (CFU)/mL in CAMHB. Then, 200 μL of the bacteria suspension was added to the 1st well of the 96-well plate and 100 μL of suspension was added to the other 11 wells. The stock solution was added to the 1st well and 2-fold serial dilution was performed. Accordingly, compound solutions at the concentrations ranging from 100 to 0.05 μg/mL were made. The 96-well plate was incubated at 37 C for 18-24 h. Lastly, the concentration at which the bacterial growth was completely inhibited was determined as the MIC value of the compound. All the experiments were carried out for three times.

To determine the MICs of **1b** for a panel of bacterial strains, the same broth microdilution assay was performed, except that the maximum compound concentration was 128 μg/mL.

#### 4.3.2. Cytotoxicity assay

DMEM media with 10% fetal bovine serum was added to HepG-2 cells or HUVEC cells, and then the cells were cultured in the incubator (37 °C, 5% CO_2_), passaged for 3-5 generations before further experiments. The cells were cultured in a 96-well plate at the density of 40-50%. Each compound (Paclitaxel as a positive control) was added to the 96-well plate at the final concentrations of 100 nM, 333 nM, 1 μM, 3 μM, 10 μM, 33 μM, and 100 μM. After culture for 48 h, cell viability was measured by CCK-8 assay kits. Briefly, 10 μL of CCK-8 solution was added to the cells, and then incubated at 37 °C for 2 h. The optical density was measured at 450 nM by using a Microplate Reader (TECAN, Switzerland). Cell viability was expressed as percentage (%) of negative control (DMSO). With GraphPad Prism 8.4.0 (GraphPad Software Inc, La Jolla, CA), CC_50_ was calculated according to the dose-response curve from nonlinear regression. Data are presented as mean ± SEM. Each assay was repeated at least twice.

#### 4.3.3. Time-kill kinetics assay

By adding a certain amount of the compound to 2 mL of CAMHB in sterilized test tubes, the compound solutions at three concentrations (1 × MIC, 4 × MIC, 16 × MIC) were prepared, respectively. 2 mL of bacterial suspension (*S. aureus* ATCC 29213) was added to every tube so as to make bacterial concentration at 10^6^-10^7^ CFU/mL. As a growth control, a mixture of CAMH broth (2 mL) and bacterial suspension (2 mL) were also prepared in a sterilized test tube. All the tubes were placed in an incubator at 37 C. The assay was performed in triplicate. At each of the following 4 time points, i.e. 0 h, 4 h, 8 h and 24 h, 100 μL of bacterial suspension was taken and serially diluted by 10 folds with the sterilized physiological saline (0.9% NaCl). 100 μL of aliquots were taken from the dilutions and respectively placed on the Tryptose Soya Agar plates. The plates were incubated at 37 C for 24 h. The number of bacterial colonies on each plate was counted. According to the dilution rate, the surviving bacteria (log_10_CFU/mL) was counted. The bacterial counts (log_10_CFU/mL) from three measurements were averaged. For each concentration of compound solution, incubation time-bacterial count curve was plotted so as to determine mode of action (bactericidal or bacteriostatic).

#### 4.3.4. Mouse liver microsomes stability

The compound and multiple positive controls (phenacetin, diclofenac, testosterone, dextromethorphan, and omeprazole) at a concentration of 1 µM were respectively incubated with mice liver microsomes (0.5 mg protein/mL) in phosphate buffer (0.1 M, pH 7.4) at 37 C, supplemented with NADPH. At the time points of 0, 5, 15, 30, 60 min, the incubation mixtures (30 µL) were treated with 200 µL of a 4 C stop solution (acetonitrile containing 5 ng/mL terfenadine) to stop the reactions. The samples were vortexed for 1 min and then centrifuged at 13,000 rpm for 10 min at 4 C. For each sample, 100 μL of supernatant was pipetted and 100 μL of distilled water was added, for LC-MS/MS analysis. The LC-MS/MS method and data analysis method are listed in supporting information (cf. **Table S3**).

#### 4.3.5. Mouse plasma stability

The compound (1 µM) was incubated with mouse plasma at 37 C in Eppendorf tubes. Propantheline bromide (1 µM) in human plasma (2 mL) was used as a control. Samples were collected at 0, 5, 15, 30, 60, 120 min (50 µL at each time point). Reaction was terminated by adding ice-cold acetonitrile/methanol (v:v=50:50) containing the internal standard (50 ng/mL labetalol, 50 ng/mL tolbutamide). Each sample was vortexed for 1 min, then the supernatant was collected after centrifugation at 4000 rpm for 10 min. Then, 100 μL of the supernatant was pipetted and 100 μL of distilled water was added for LC-MS/MS analysis. The LC-MS/MS method and data analysis method are listed in supporting information (cf. **Table S3**)

#### 4.3.6. In vivo efficacy

KM mice (18-20 g) were randomly divided into 9 groups (10 mice per group, male/female = 1:1), i.e., a negative control group (mice administered with physiological saline), 4 compound treatment groups (mice administered with compound **1b** at 7 mg/kg, 10 mg/kg, 14 mg/kg and 20 mg/kg, i.v.), 4 vancomycin treatment groups (positive control, mice administered with vancomycin at 10 mg/kg, 7 mg/kg, 5 mg/kg and 3.5 mg/kg, i.v.). Prior to efficacy evaluation, all the mice were intraperitoneally injected with 0.5 mL (minimal lethal dose, MLD) of MRSA 16-1, a clinical isolate of MRSA. After 10 min, the compound (or vancomycin) at the aforementioned doses was administered via tail vein injection.

Following the compound treatment, the mice were observed for 7 consecutive days. The survival rate in each group was recorded.

#### 4.3.7 S. aureus gyrase inhibition assay

*S. aureus* gyrase was purchased from Inspiralis Ltd. (Norwich, United Kingdom). The gyrase inhibition assay required 10 mL of mixture that contained 5 nM enzyme (*S. aureus* gyrase), buffer (40 mM HEPES-KOH (pH 7.6), 10 mM magnesium acetate, 10 mM dithiothreitol, 50 g/L BSA, 500 mM potassium glutamate), the compound (0.001 μM-10 μM), 1% DMSO, 10 nM linear pBR322 DNA, and 100 mM ATP. Prior to the assay, the 3-fold serial dilutions of the compound at 10 times the specified concentrations were prepared, by being dissolved in DMSO (10 mM) and diluted with the assay buffer and DMSO. For the assay, the buffer (7 μL), the prepared compound dilution (1 μL), the linear pBR322 DNA (0.5 μL) and *S. aureus* gyrase (0.5 μL) were added to a PCR tube sequentially and mixed by vortex. To initiate the enzymatic reaction, ATP (1 μL) was added to the tube. The PCR tube was sealed and incubated at 37 C for 30 min. The generated ADP was quantified by using the ADP-Glo kits. 40 μL of ADP-Glo reagent was added to the reaction mixture and incubated at 37 C for 40 min, which was used to stop the ADP-generating reaction and use up the remaining ATP. 50 μL of the detection reagent was added and mixed for 5 min. 100 μL of the mixture was transferred to the 96-well plate and the luminescence (LU) was measured using a BioTek Synergy 2 microplate reader. The enzymatic activity was calculated based on LU. According to the activity (%) values of the *S. aureus* gyrase treated by the compound at nine concentrations, IC_50_ was determined using nonlinear regression with normalized dose-response fit implemented in GraphPad Prism 5 software (GraphPad Software Inc., La Jolla, CA). Herein, novobiocin (a classical gyrase inhibitor) was used as a positive control, and the assays were performed in duplicate.

#### 4.3.8 S. aureus DNA supercoiling assay

The *S. aureus* DNA supercoiling assay was mostly similar to the gyrase ATPase inhibition assay described above, except that agarose gel electrophoresis was used to quantify DNA. Using the same mixture (10 μL) as that for the gyrase ATPase inhibition assay in the PCR tube, the enzymatic reaction was carried out at 37 C for 30 min. Subsequently, the sample in the PCR tube was centrifuged at 1500 rpm for 1 min and then mixed with 2 μL of DNA loading buffer. The mixture was run through a 1% agarose gel in TAE buffer (40 mM Trisacetate, 2 mM EDTA) for 2 h at 80V. The gel was stained with ethidium bromide for 60 min, visualized, and photographed under UV light. Likewise, novobiocin (a classical gyrase inhibitor) was used as the positive control.

## AUTHOR INFORMATION

### CRediT authorship contribution statement

Gao Zhang: Data curation, Formal analysis, Investigation, Methodology, Software, Validation, Visualization, Writing–original draft. Jiaxin Liang: Investigation, Methodology, Resources, Validation, Writing–review & editing. Gang Wen: Formal analysis, Investigation, Methodology, Resources. Mingli, Yao: Data curation, Investigation, Software, Visualization. Yuqing Jia: Investigation, Resources. Bo Feng: Investigation, Resources. Jishun Li: Investigation, Resources. Zunsheng Han: Formal analysis, Investigation, Resources. Qingxin Liu: Data curation, Software. Tianlei Li: Project administration, Supervision. Wenxuan Zhang: Project administration, Supervision. Hongwei Jin: Software. Jie Xia: Conceptualization,

Funding acquisition, Project administration, Supervision, Writing–review & editing. Liang Peng: Funding acquisition, Project administration, Supervision. Song Wu: Conceptualization, Funding acquisition, Project administration, Supervision.

## Acknowledgements

This study was funded by the CAMS Innovation Fund for Medical Sciences (2021-I2M-1-069), Guangdong Basic and Applied Basic Research Foundation-Enterprise Joint Project (2023A1515220222), and the Program for Foreign Talent of Ministry of Science and Technology of the People’s Republic of China (G2021194015L).

## Appendix A. Supplementary data

Supplementary data related to this article can be found at xxx.

## Abbreviations

ATCC: American Type Culture Collection
CAMHB: Cation-Adjusted Mueller-Hinton broth
CC_50_: Concentration of cytotoxicity 50%
CFU: colony forming unit
CL_int_: intrinsic clearance
DCM: dichloromethane
DMF: *N,N*-dimethylmethanamide
DMSO: dimethyl sulfoxide
EFA: *E. faecalis*
EFM: *E. faecium*
HepG2: hepatocellular carcinoma
HUVEC: human umbilical vein endothelial cells
IC_50_: Half-maximal inhibitory concentration
LU: luminescence
MD: molecular dynamics
MIC: minimum inhibitory concentration
MLD: minimum lethal dose
MRSA: methicillin-resistant *S. aureus*
MRSE: methicillin-resistant *S. epidermidis*
MSSA: methicillin-sensitive *S. aureus*
MSSE: methicillin-sensitive *S. epidermidis*
PDB: Protein Data Bank
RMSD: root mean square error
SAR: structure-activity relationship
SI: selectivity index
SI: selectivity index
Tc: Tanimoto coefficient
THF: tetrahydrofuran
TLC: thin-layer chromatography.

