## Supporting information for "Discovery of novel 1,4-dicarbonylthiosemicarbazides as DNA gyrase inhibitors for the treatment of MRSA infection"

Liang Peng

Department of Clinical Laboratory

The Fifth Affiliated Hospital of Guangzhou Medical University Guangzhou

Jie Xia & Song Wu

Institute of Materia Medica

Chinese Academy of Medical Sciences &

Peking Union Medical College

Beijing, China

### **Table S1**. Antibacterial activity of compound **1b** for a panel of gram-positive bacterial strains (clinical isolates).

| Strains^a^ | MIC(μg/ml) | |
| --- | --- | --- |
|  | **1b** | vancomycin |
| MRSA 15-1 | 4 | 2 |
| MRSA 15-2 | 4 | 2 |
| MRSA 15-3 | 4 | 1 |
| MRSA 17-1 | 2 | 0.5 |
| MRSA 17-2 | 4 | 0.5 |
| MRSA 17-3 | 2 | 1 |
| MRSA 17-4 | 2 | 1 |
| MRSA 17-5 | 2 | 0.5 |
| MRSA 17-6 | 2 | 0.5 |
| MRSA 17-7 | 2 | 0.25 |
| MRSA 17-8 | 2 | 0.5 |
| MRSA 17-9 | 2 | 0.5 |
| MRSA 17-10 | 2 | 1 |
| MRSA 16-1 | 2 | 1 |
| MRSA 16-2 | 2 | 1 |
| MSSA 16-1 | 1 | 0.5 |
| MSSA 16-2 | 1 | 0.5 |
| MSSA 15-1 | 1 | 1 |
| MSSA 15-2 | 2 | 1 |
| MSSA 15-5 | 2 | 0.5 |
| MSSA 15-6 | 2 | 0.5 |
| MSSA 15-7 | 2 | 1 |
| MSSA 15-8 | 2 | 0.5 |
| MRSE 17-1 | 0.5 | 1 |
| MRSE 17-3 | 1 | 2 |
| MRSE 15-1 | 4 | 2 |
| MRSE 15-2 | 4 | 2 |
| MRSE 15-3 | 2 | 2 |
| MRSE 15-76 | 2 | 2 |
| MRSE 15-78 | 2 | 1 |
| MRSE 15-80 | 1 | 2 |
| MRSE 15-81 | 0.5 | 2 |
| MRSE 15-82 | 1 | 1 |
| MRSE 15-83 | 1 | 1 |
| MSSE 15-1 | 1 | 1 |
| MSSE 15-2 | 1 | 2 |
| MSSE 15-3 | 1 | 2 |
| MSSE 15-4 | 1 | 2 |
| MSSE 15-58 | 1 | 2 |
| MSSE 15-59 | 1 | 1 |
| MSSE 15-61 | 1 | 1 |
| MSSE 15-64 | 8 | 1 |
| EFA 15-1 | 8 | 0.5 |
| EFA 15-2 | 8 | 0.5 |
| EFA 15-55 | 8 | 0.5 |
| EFA 15-57 | 4 | 2 |
| EFA 15-58 | 4 | 0.5 |
| EFA 15-60 | 8 | 8 |
| EFA 15-68 | 4 | 0.5 |
| EFA 15-66 | 4 | 2 |
| EFM 15-1 | 8 | 1 |
| EFM 15-2 | 4 | 2 |
| EFM 15-6 | 2 | 1 |
| EFM 15-7 | 2 | 0.5 |
| EFM 15-8 | 2 | 0.5 |
| EFM 15-9 | 4 | 0.5 |
| EFM 15-11 | 4 | 0.5 |
| EFM 15-52 | 8 | 2 |

^a^ MSSA, methicillin-sensitive S. aureus; MRSA, methicillin-resistant S. aureus; MSSE, methicillin-sensitive S. epidermidis; MRSE, methicillin-resistant S. epidermidis; EFA, E. faecalis; EFM, E. faecium.

### **Table S2**. MIC distribution for clinical isolates for **1b**.

| Strains^a^ | Compounds | MIC (µg/ml) distribution (Count of strains^c^) | | | | | |
| --- | --- | --- | --- | --- | --- | --- | --- |
|  |  | 8^b^ | 4 | 2 | 1 | 0.5 | 0.25 |
| MRSA  (15 strains) | **1b** |  | 4 | 11 |  |  |  |
|  | vancomycin |  |  | 2 | 6 | 6 | 1 |
| MSSA  (8 strains) | **1b** |  |  | 5 | 3 |  |  |
|  | vancomycin |  |  |  | 3 | 5 |  |
| MRSE  (11 strains) | **1b** |  | 2 | 3 | 4 | 2 |  |
|  | vancomycin |  |  | 7 | 4 |  |  |
| MSSE  (8 strains) | **1b** | 1 |  |  | 7 |  |  |
|  | vancomycin |  |  | 4 | 4 |  |  |
| EFA  (8 strains) | **1b** | 4 | 4 |  |  |  |  |
|  | vancomycin | 1 |  | 2 |  | 5 |  |
| EFM  (8 strains) | **1b** | 2 | 3 | 3 |  |  |  |
|  | vancomycin |  |  | 2 | 2 | 4 |  |

^a^ MSSA, methicillin-sensitive S. aureus; MRSA, methicillin-resistant S. aureus; MSSE, methicillin-sensitive S. epidermidis; MRSE, methicillin-resistant S. epidermidis; EFA, E. faecalis; EFM, E. faecium.

^b^ MIC values, μg/mL.

^c^ the number of bacterial strains that could be inhibited by a compound at a specific concentration.

### **Table S3**. LC-MS/MS method for *in vitro* metabolic stability of **1b**.

| LC-MS/MS analysis for Mouse liver microsomes stability | | | | |
| --- | --- | --- | --- | --- |
| LC methods | | | | |
| Mobile phase A | 0.1% Formic Acid in Water | | | |
| Mobile phase B | Acetonitrile | | | |
| Column | Zorbax Eclipse Plus C18, 50*2.1mm, 1.8μm | | | |
| Internal standard | Terfenadine | | | |
| LC system | Agilent 1290 | | | |
| Injection volume | 5 μL | | | |
| Gradient conditions | Time (min) | Flow (mL/min) | A (%) | B (%) |
|  | 0 | 0.3 | 50 | 50 |
|  | 2 | 0.3 | 5 | 95 |
|  | 2.5 | 0.3 | 5 | 95 |
|  | 2.51 | 0.3 | 50 | 50 |
|  | 4 | 0.3 | 50 | 50 |
| MS method | | | | |
| MS | Agilent 6495C | | | |
| Ionization model | AJS ESI | | | |
| Scan type | MRM | | | |
| Polarity | Positive | | | |
| Name | Ion pair (m/z) | Retention Time (min) | Frag (V) | CE (V) |
| **1b** | 324.4/191.0 | 1.7 | 166 | 10 |
| Terfenadine | 472.3/454.3 | 0.9 | 166 | 22 |
| LC-MS/MS analysis for Mouse plasma stability | | | | |
| LC methods | | | | |
| Mobile phase A | 0.1% Formic Acid in Water | | | |
| Mobile phase B | Acetonitrile | | | |
| Column | Synergi 4 µm Fusion-RP 80Å , 2 mm *50 mm | | | |
| Internal standard | Tolbutamide | | | |
| LC system | ExionLC | | | |
| Injection volume | 1 μL | | | |
| Gradient conditions | Time (min) | Flow (mL/min) | A (%) | B (%) |
|  | 0.00 | 0.8 | 75 | 25 |
|  | 0.90 | 0.8 | 5 | 95 |
|  | 1.20 | 0.8 | 5 | 95 |
|  | 1.21 | 0.8 | 75 | 25 |
|  | 1.50 | 0.8 | 75 | 25 |
| MS method | | | | |
| MS | Triple Quad 5500+ | | | |
| Ionization model | ESI | | | |
| Scan type | MRM | | | |
| Polarity | Negative | | | |
| Name | Ion pair (m/z) | Retention Time (min) | DP (eV) | CE (eV) |
| **1b** | 322.20/82.90 | 0.75 | -80 | -46 |
| Tolbutamide | 269.20/170.00 | 0.59 | -51 | -22 |

### **Table S4**. *In vitro* stability of **1b** in mice liver microsomes and mouse plasma.

| compounds | Mouse liver microsomes | | Mouse plasma |
| --- | --- | --- | --- |
|  | T_1/2_ (min) | *In vitro* CLint^a^ (µL/min/mg protein) | T_1/2_ (min) |
| Omeprazole | 9.7 | 143.4 | - |
| Testosterone | 1.3 | 1068.2 | - |
| Dextromethorphan | 7.7 | 179.0 | - |
| Diclofenac | 11.5 | 120.6 | - |
| Phenacetin | 2.6 | 531.2 | - |
| Propantheline bromide | - | - | 38.3 |
| 1b | 28.3 | 49.0 | >120 |
| ^a^ intrinsic clearance. | | | |

### **Table S5**. 12 proteins related to *S. aureus* and their corresponding inhibitors. For each protein, the inhibitor with the highest Tc value to **1b** and the co-crystal structure were shown.

| ID | Protein name | No. of ligands | Tc^a^ | Structure^b^ | PDB ID | Reso-lution  (Å) | Docking  Score ^c^ |
| --- | --- | --- | --- | --- | --- | --- | --- |
| 1 | Cell division protein ftsZ | 10 | 0.40 | 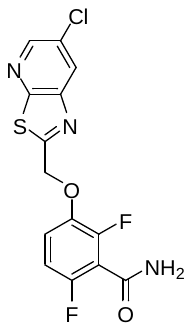 | 6KVP | 1.40 | -^d^ |
| 2 | Dehydrosqualene  synthase | 46 | **0.52** | 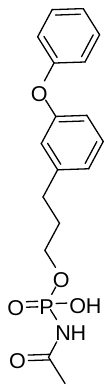 | 4F6V | 2.30 | **-7.28** |
| 3 | Dihydrofolate reductase | 149 | 0.45 | 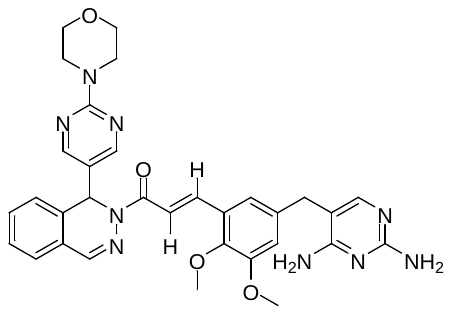 | 3FRD | 2.10 | - |
| 4 | Dihydroneopterin aldolase | 1 | 0.39 | 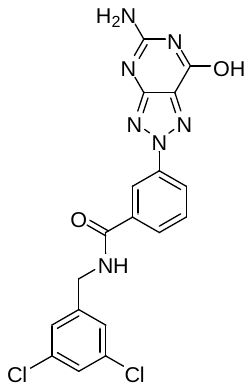 | 1RSD | 2.50 | - |
| 5 | DNA gyrase  subunit A | 47 | 0.44 | 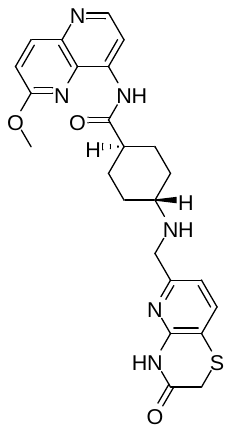 | 7MVS | 2.60 | - |
| 6 | DNA gyrase  subunit B | 253 | **0.56** | 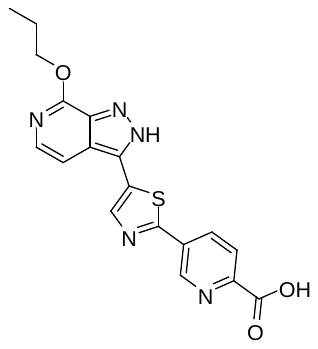 | 4URO | 2.59 | **-8.17** |
| 7 | DNA ligase | 47 | 0.44 | 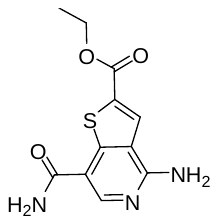 | 4CC6 | 2.01 | - |
| 8 | Enoyl-[acyl-carrier-protein] reductase (FABI) | 207 | 0.49 | 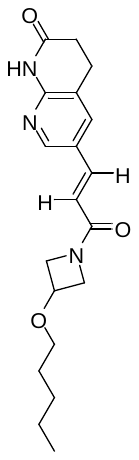 | 4FS3 | 1.80 | - |
| 9 | Methionine aminopeptidase | 2 | 0.29 | 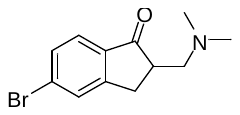 | 1QXY | 1.04 | - |
| 10 | Peptide  deformylase | 39 | **0.51** | 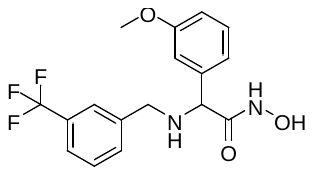 | 6JFG | 2.60 | **-6.88** |
| 11 | pyruvate kinase | 91 | 0.49 | 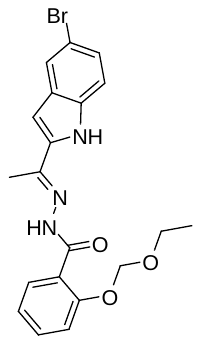 | 3T0T | 3.10 | - |
| 12 | UDP-N-acetylmuramate dehydrogenase | 6 | 0.44 | 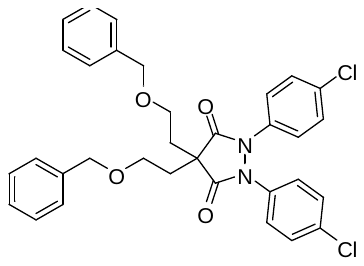 | 1HSK | 2.30 | - |
| ^a^: Tc: Tanimoto coefficient, the highest Tc value between **1b** and inhibitors of each protein.  ^b^: Structure with the highest Tc value is shown.  ^c^: FRED Chemgauss4 score  ^d^: Not determined | | | | | | | |

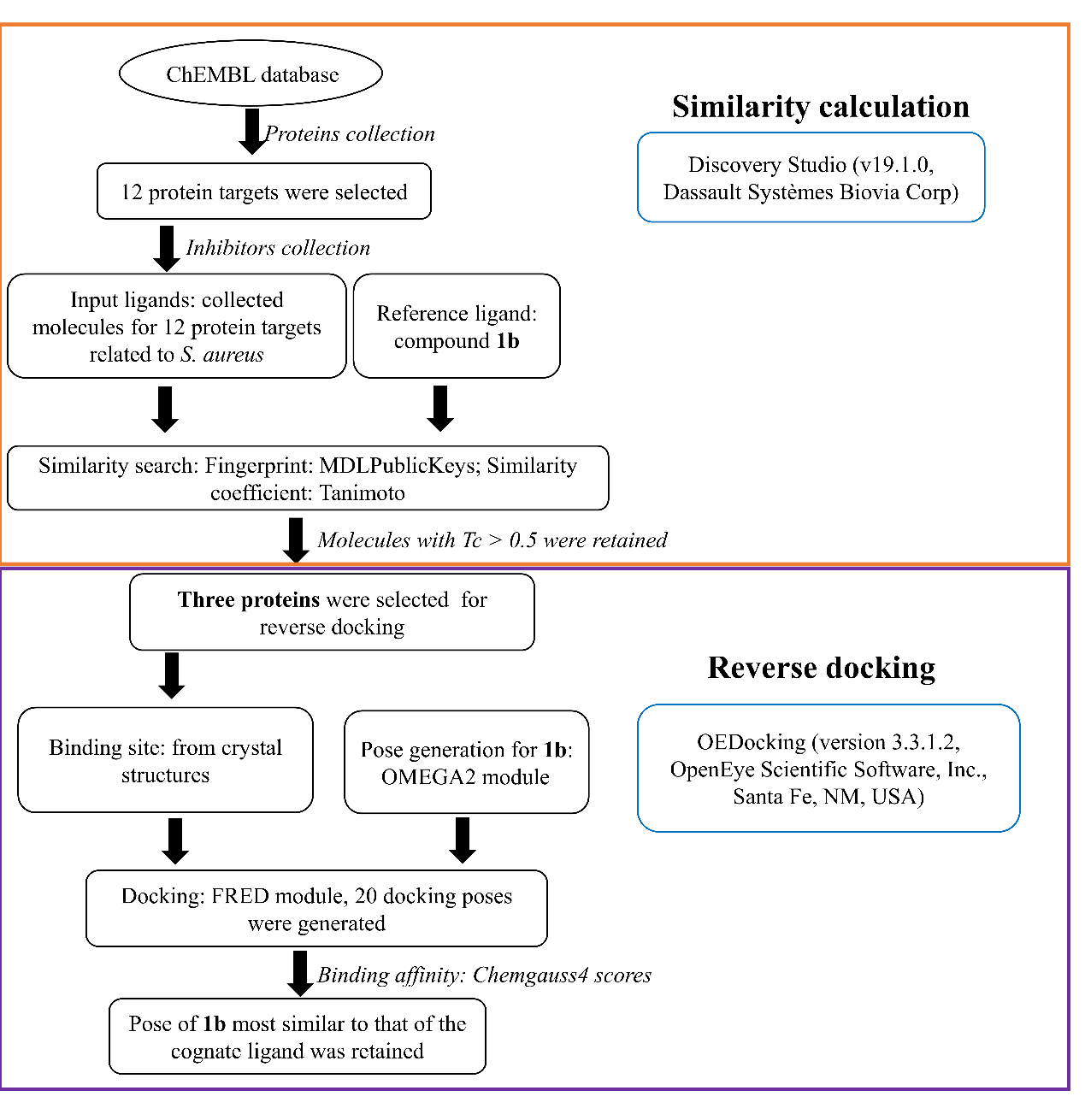

### **Fig. S1**. Our workflow for target prediction. (A) The protocol of similarity search. (B) The protocol of reverse docking.

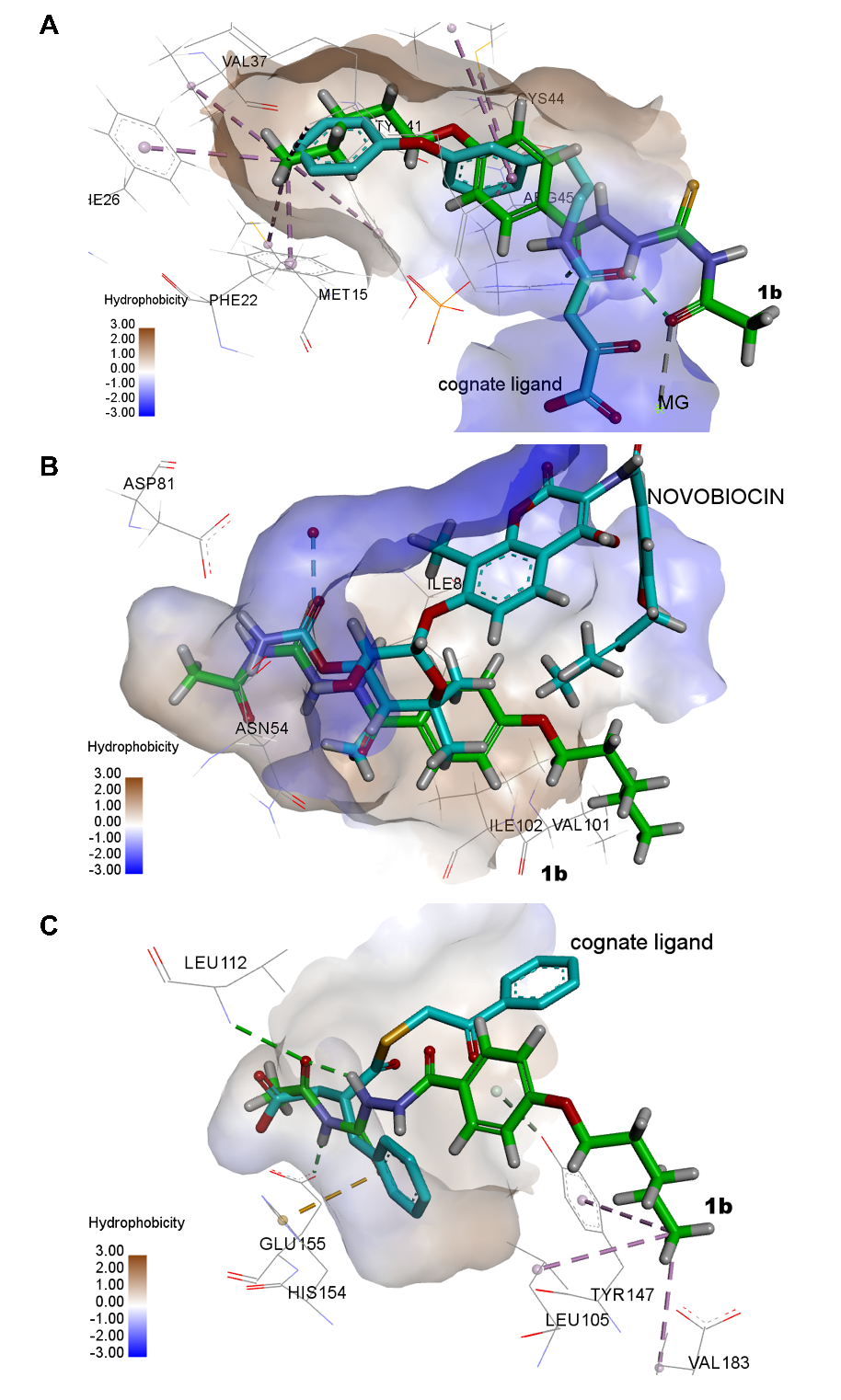

### **Fig. S2**. The binding modes of **1b** to three potential target proteins, respectively. For cognate ligands, carbon atoms were colored in blue. For compound **1b**, carbon atoms were colored in green. (A) The compound **1b** docked to 4F6V (dehydro-squalene synthase, FRED Chemgauss4 score: -7.28). (B) The compound **1b** docked to 4URO (DNA gyrase subunit B, FRED Chemgauss4 score: -8.17). (C) The compound **1b** docked to 6JFG (peptide deformylase, FRED Chemgauss4 score: -6.88).

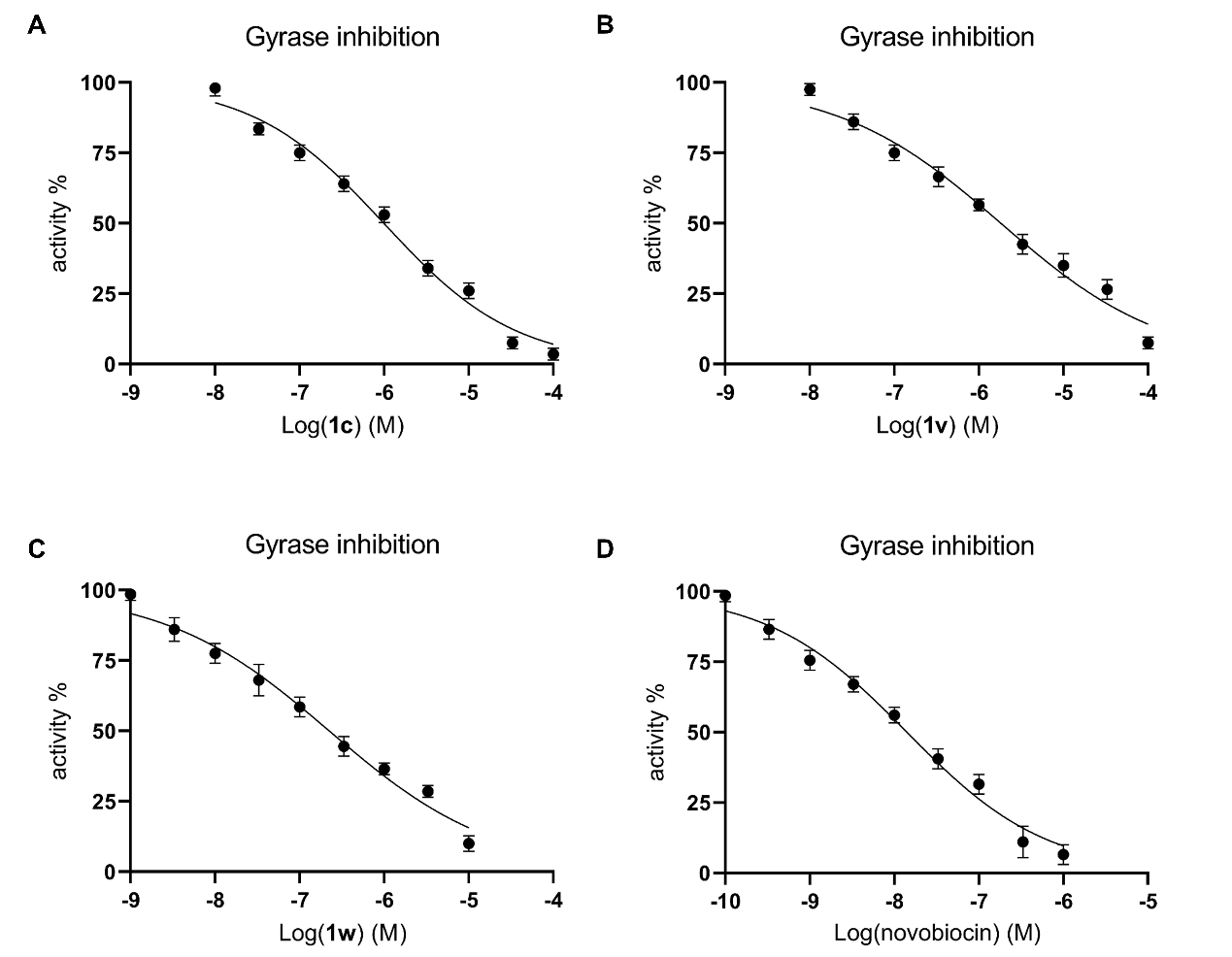

### **Fig. S3**. The activity of the tested derivatives against DNA gyrase. (A) IC_50_ value of **1c**: 1.01 μM. (B) IC_50_ value of **1v**: 1.89 μM. (C) IC_50_ value of **1w**: 0.23 μM. (D) IC_50_ value of novobiocin: 0.015 μM.

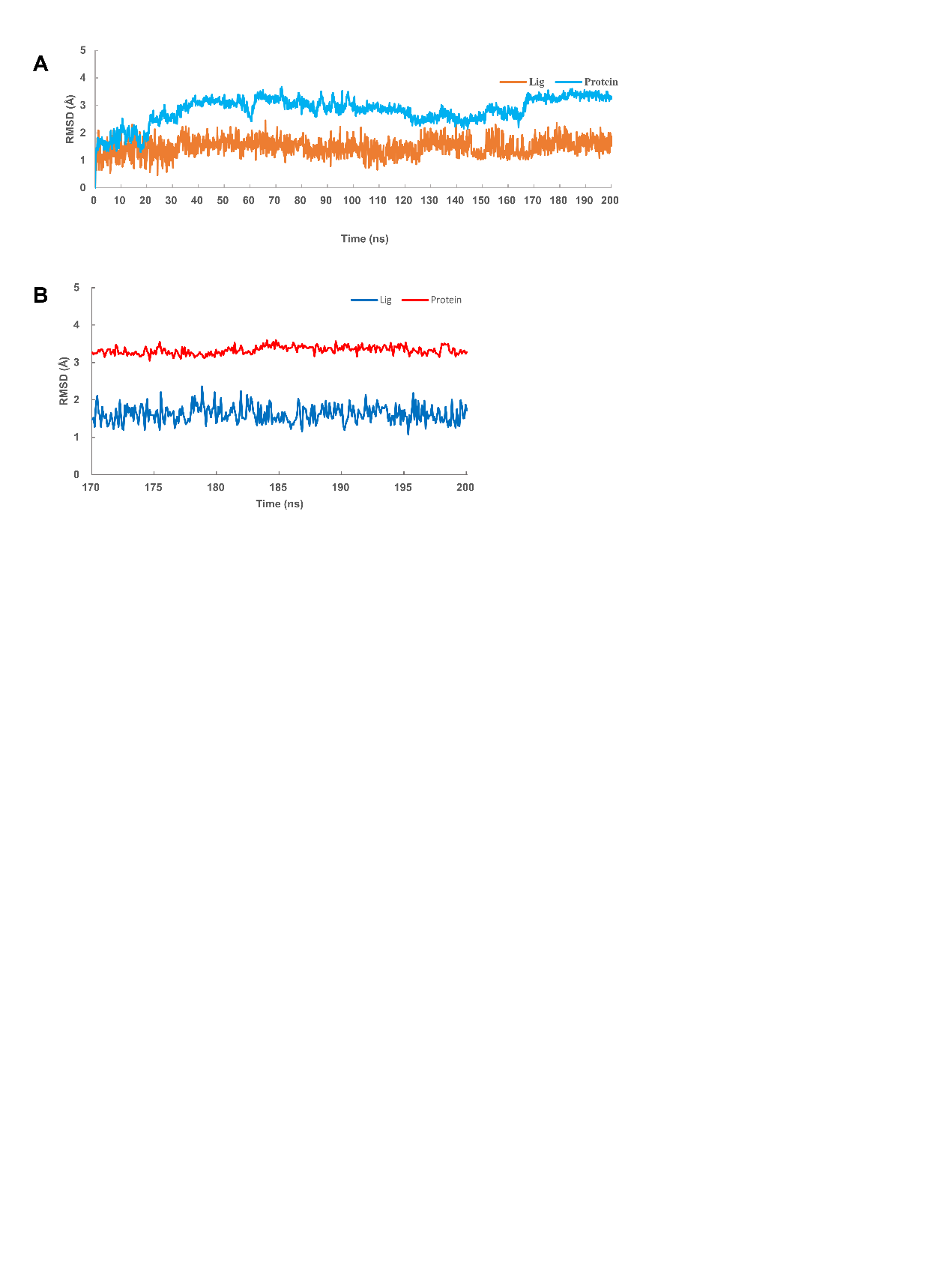

### **Fig. S4**. RMSD-Time plots for **1b** and gyrase ATPase domain in molecular dynamics simulation. (A) RMSD state from 0 ns to 200 ns. (B) Stable state from 170 ns to 200 ns.

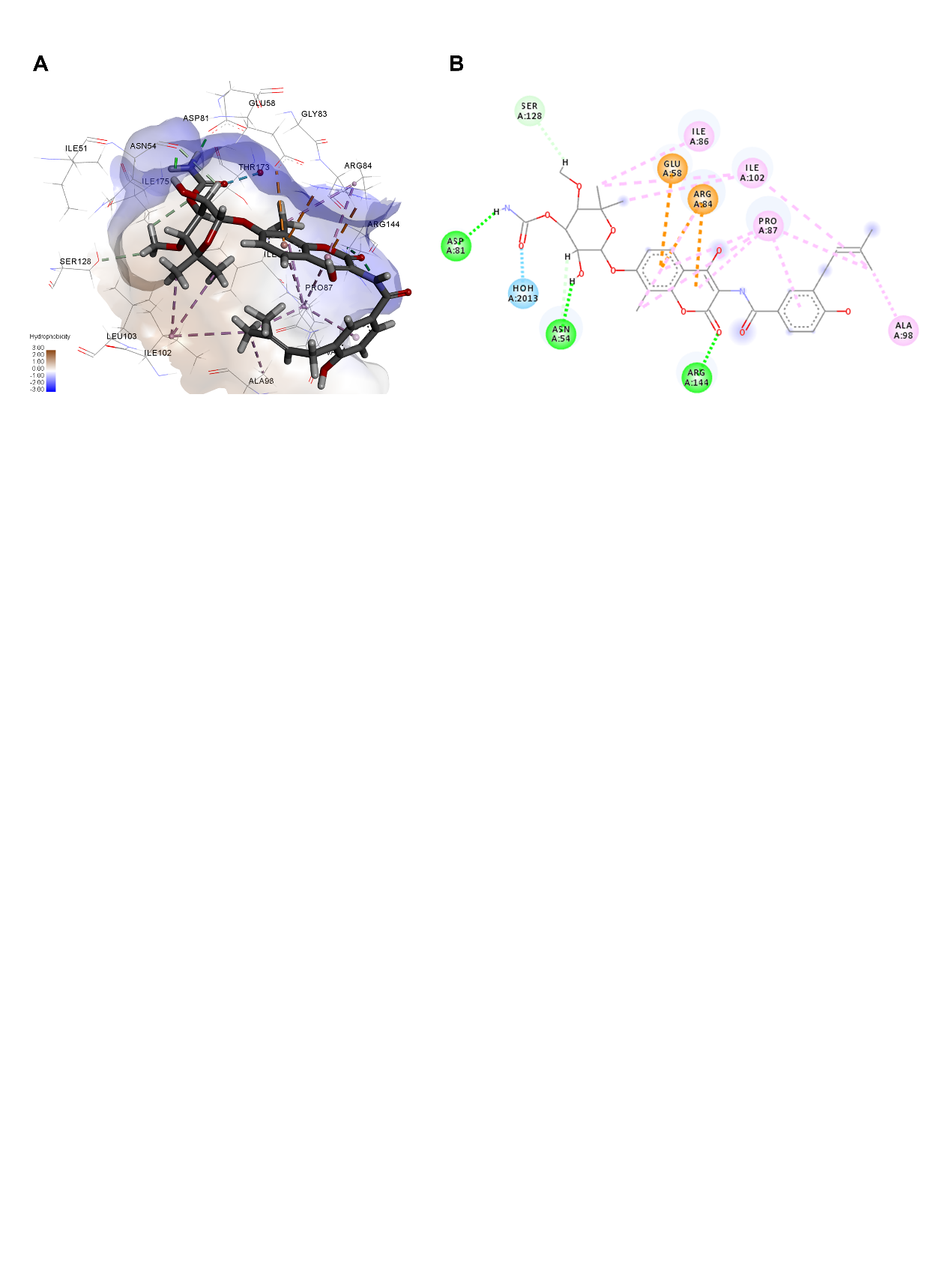

### **Fig. S5**. The co-crystal structure of novobiocin and gyrase ATPase domain (PDB ID: 4URO). (A) Novobiocin located in the ATP binding site of gyrase. The surface of the binding pocket was displayed according to pocket hydrophobicity. (B) 2D diagram of novobiocin bound to gyrase ATPase domain. Here, the interactions with ASP81, GLU58, ALA98 and PRO87 could also be observed in the binding of **1b** to gyrase ATPase domain.

### **Fig. S6**. ^1^H and ^13^C NMR spectra for all the synthesized 1,4-dicarbonylthiosemicarbazide derivatives.

^1^H-NMR spectrum of *N-(2-(4-n-butoxybenzoyl)hydrazine-1-carbonothioyl)acetamide (****1a****)*

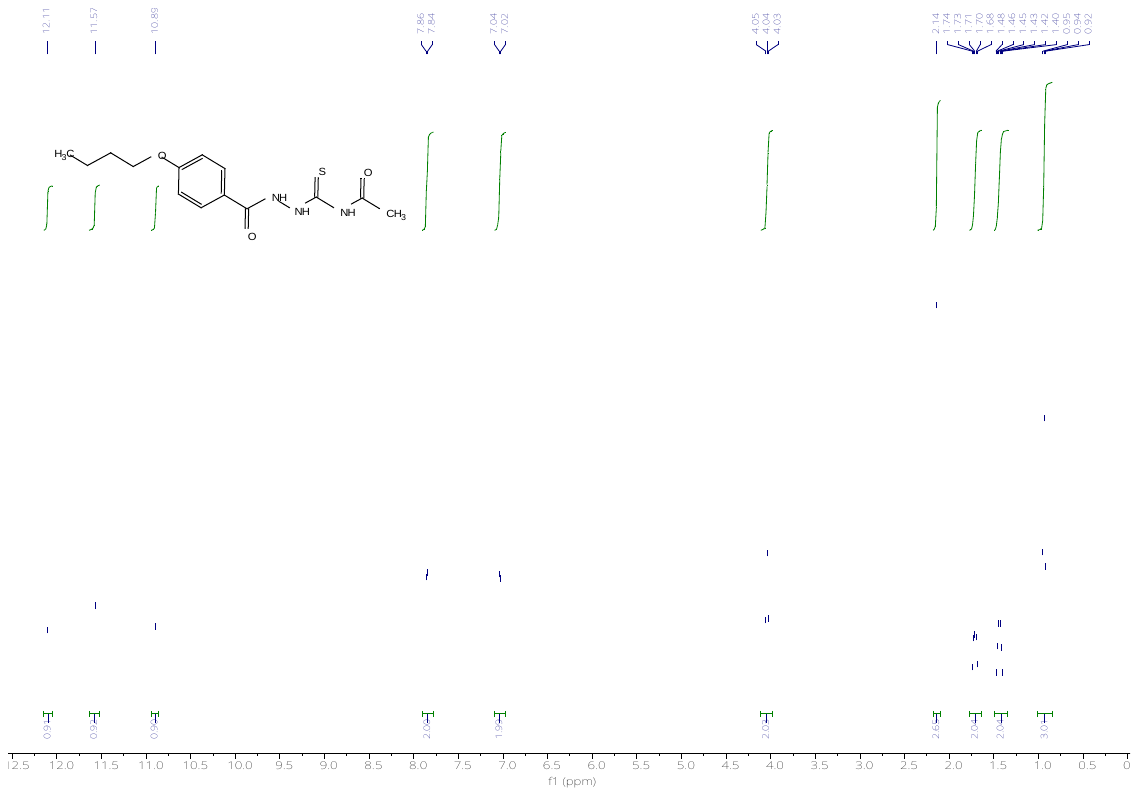

^13^C-NMR spectrum of *N-(2-(4-n-butoxybenzoyl)hydrazine-1-carbonothioyl)acetamide (****1a****)*

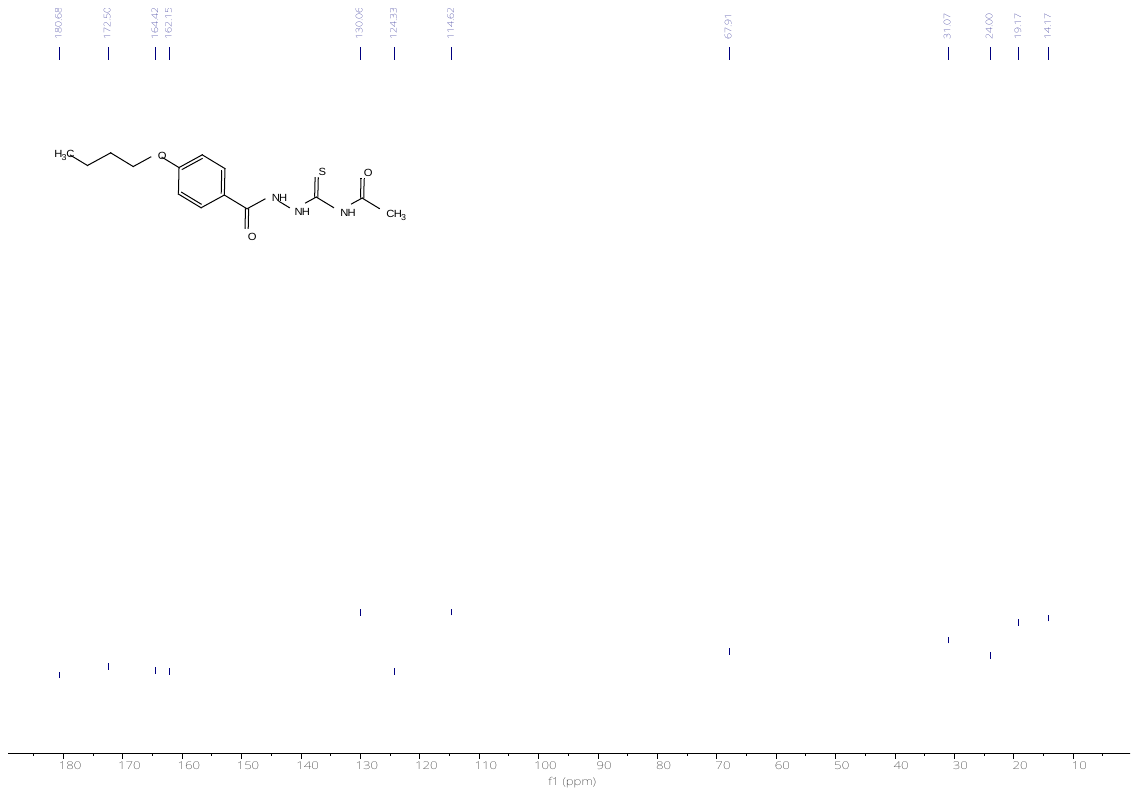

^1^H-NMR spectrum of *N-(2-(4-(n-pentoxy)benzoyl)hydrazine-1-carbonothioyl)acetamide (****1b****)*

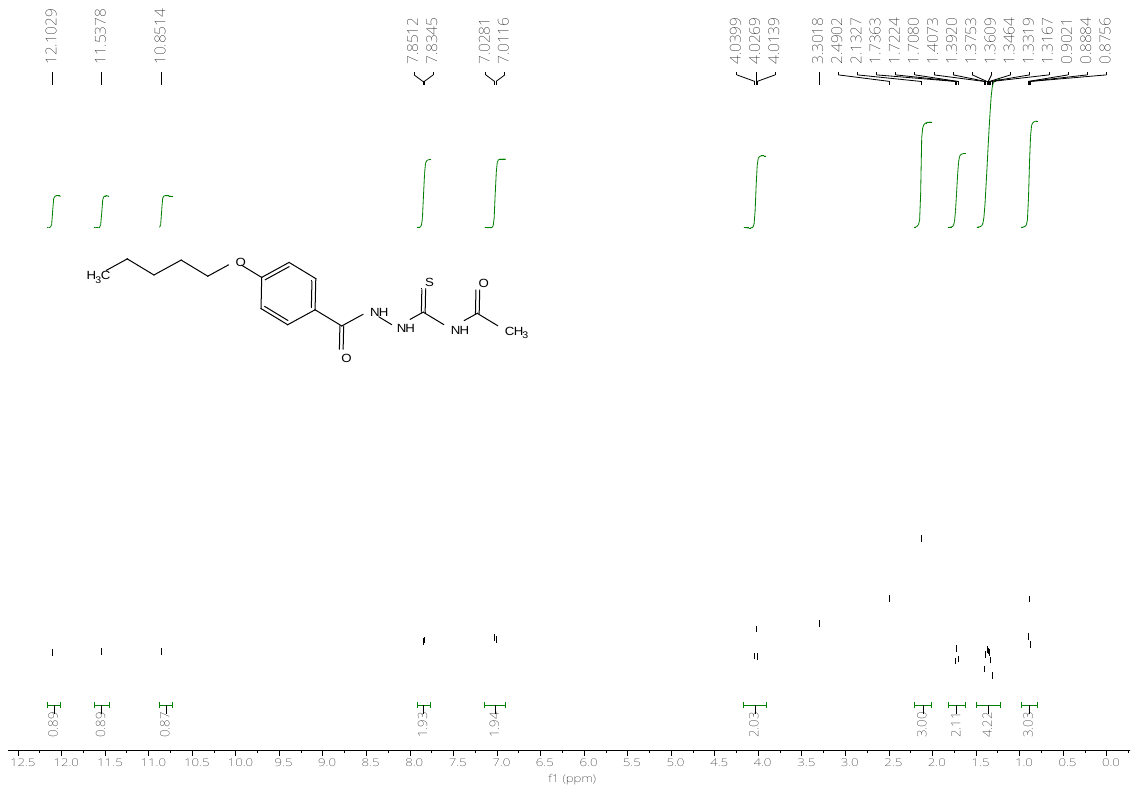

^13^C-NMR spectrum of *N-(2-(4-(n-pentoxy)benzoyl)hydrazine-1-carbonothioyl)acetamide (****1b****)*

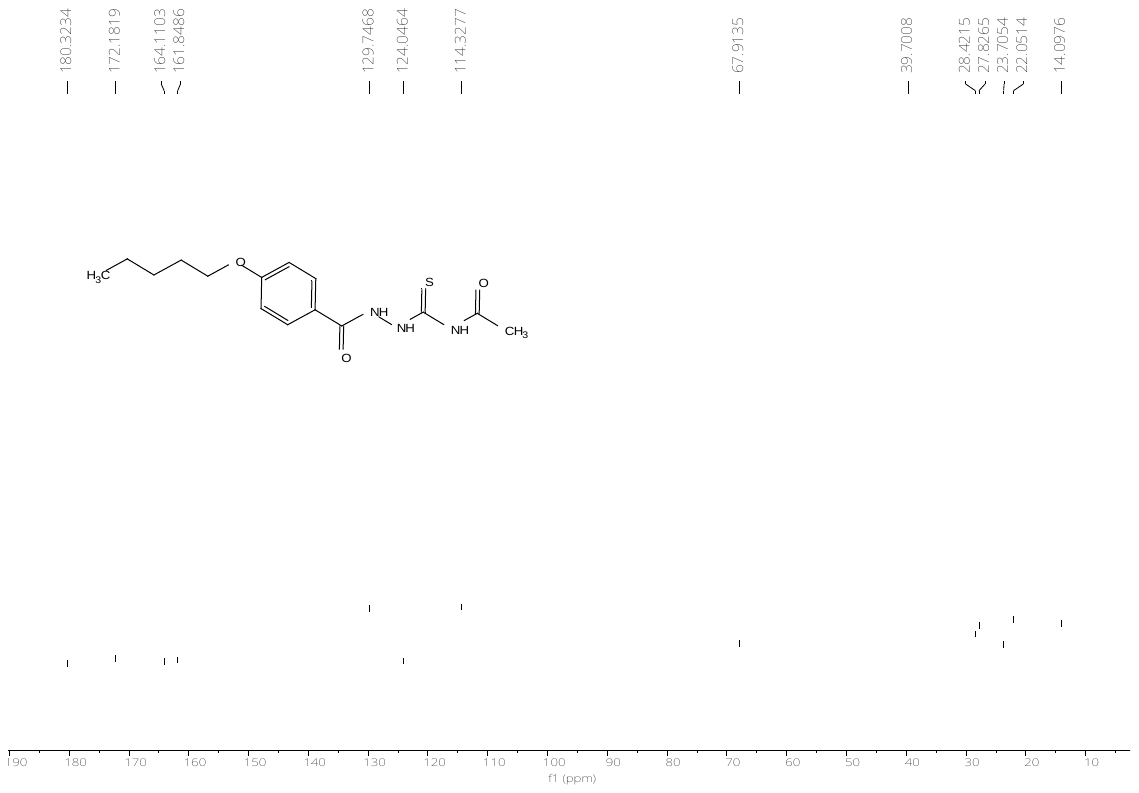

^1^H-NMR spectrum of *N-(2-(4-(n-hexoxy)benzoyl)hydrazine-1-carbonothioyl)acetamide (****1c****)*

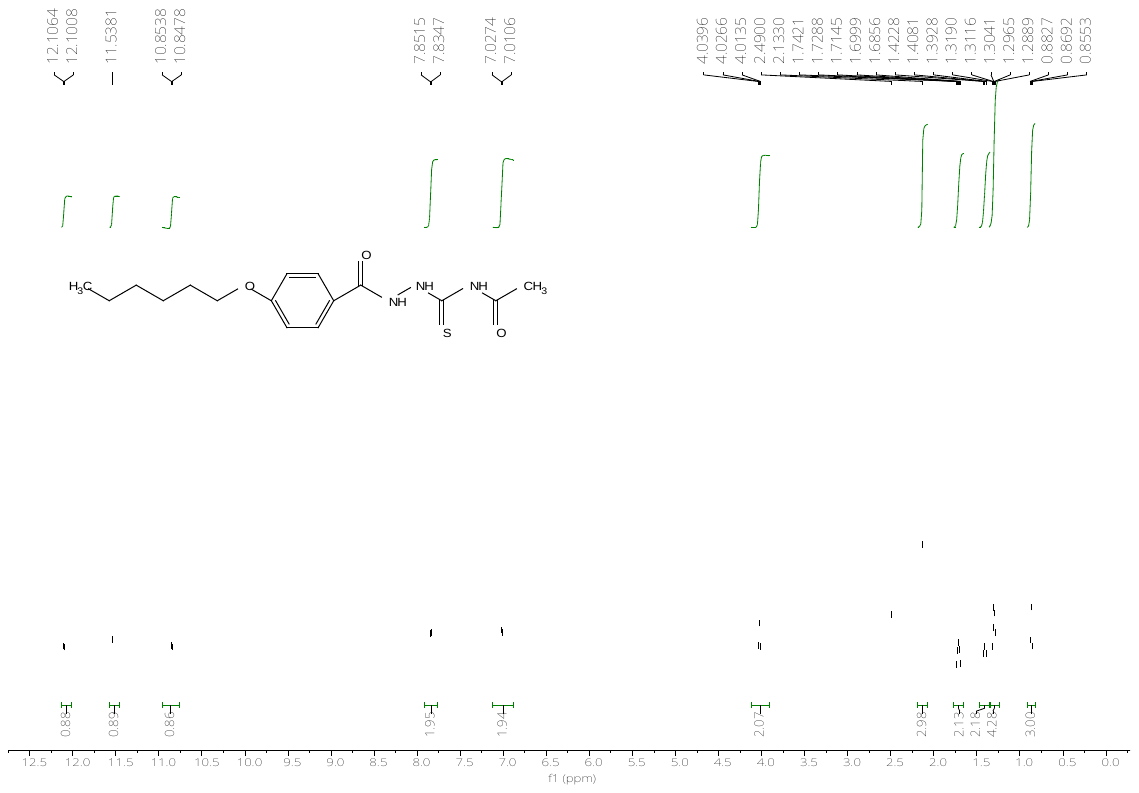

^13^C-NMR spectrum of *N-(2-(4-(n-hexoxy)benzoyl)hydrazine-1-carbonothioyl)acetamide (****1c****)*

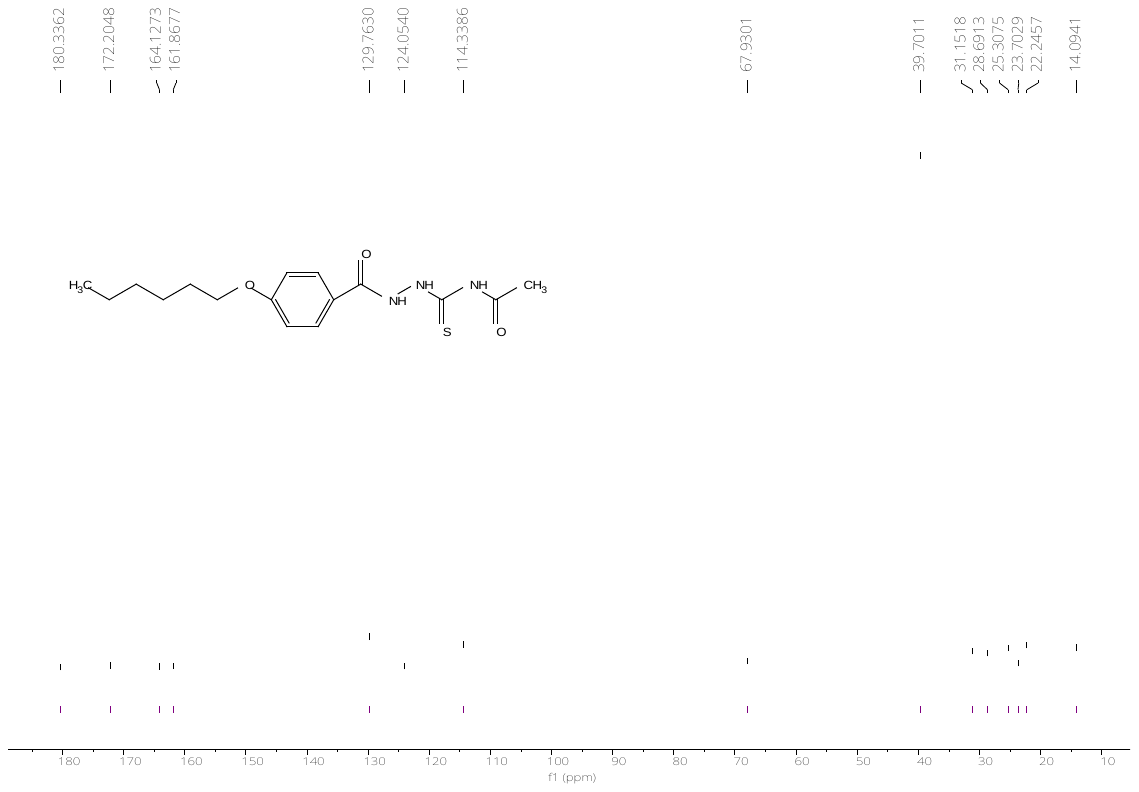

^1^H-NMR spectrum of *N-(2-(4-(n-undecoxy)benzoyl)hydrazine-1-carbonothioyl)acetamide (****1d****)*

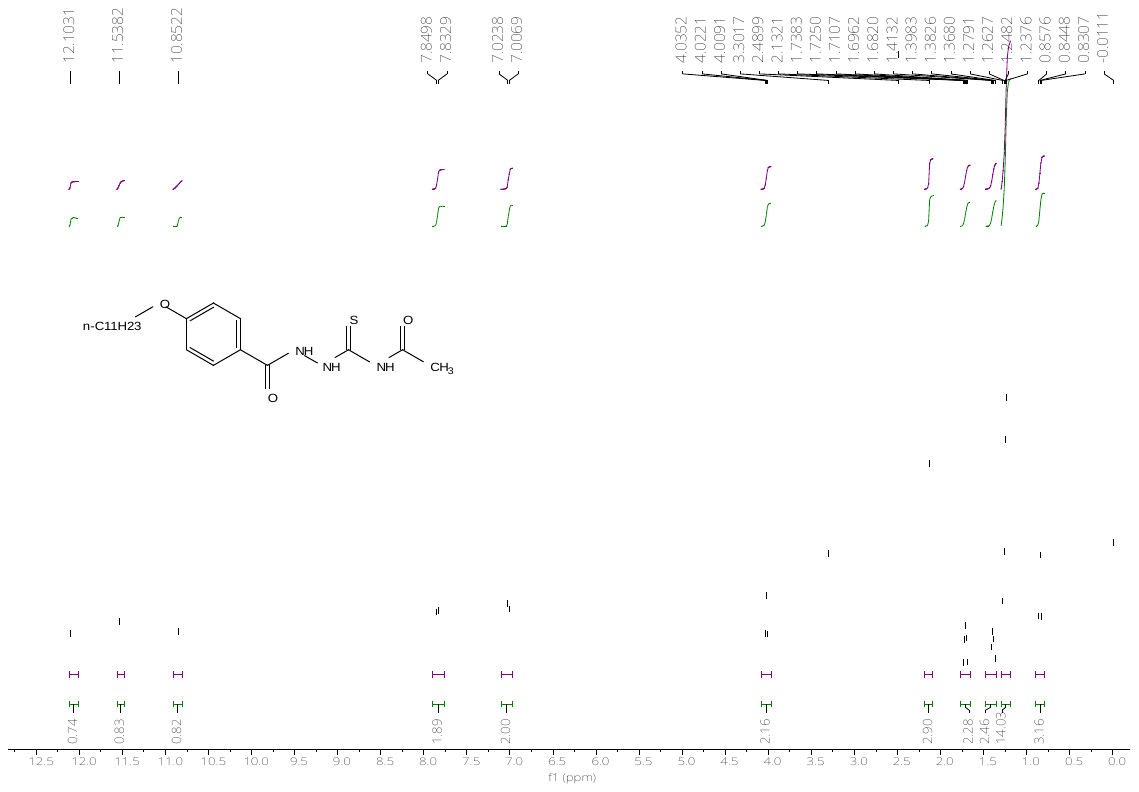

^13^C-NMR spectrum of *N-(2-(4-(n-undecoxy)benzoyl)hydrazine-1-carbonothioyl)acetamide (****1d****)*

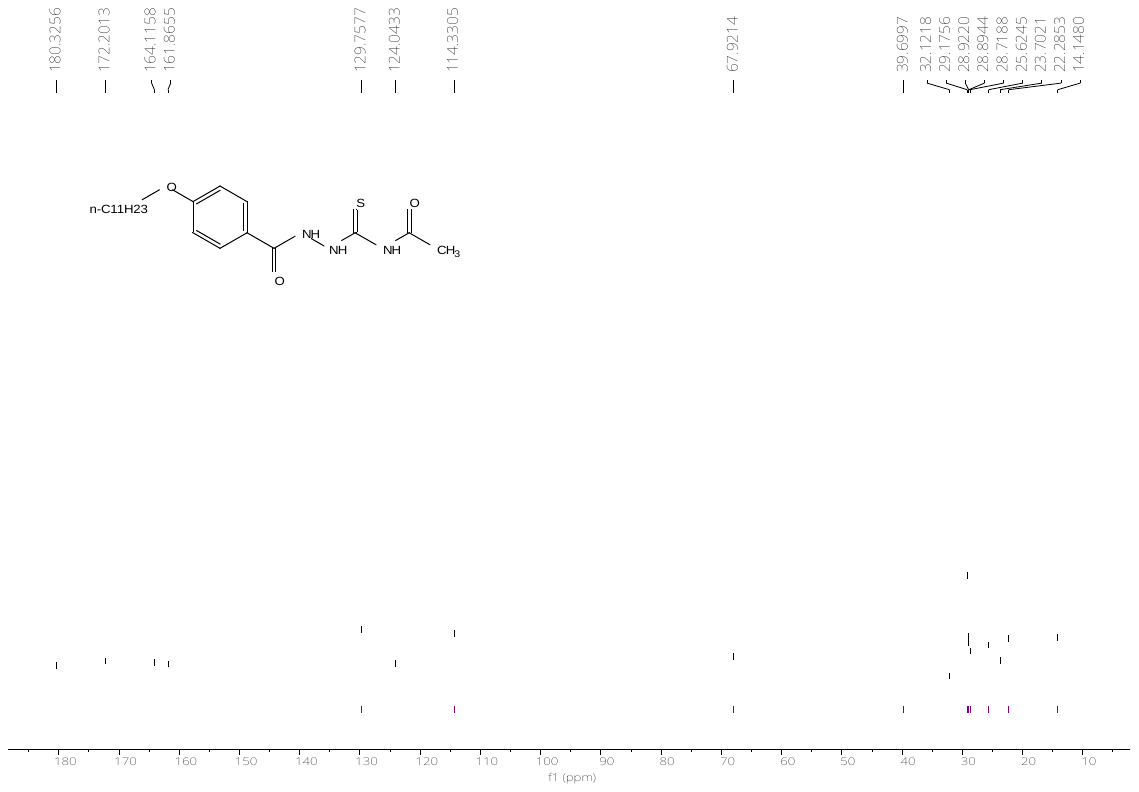

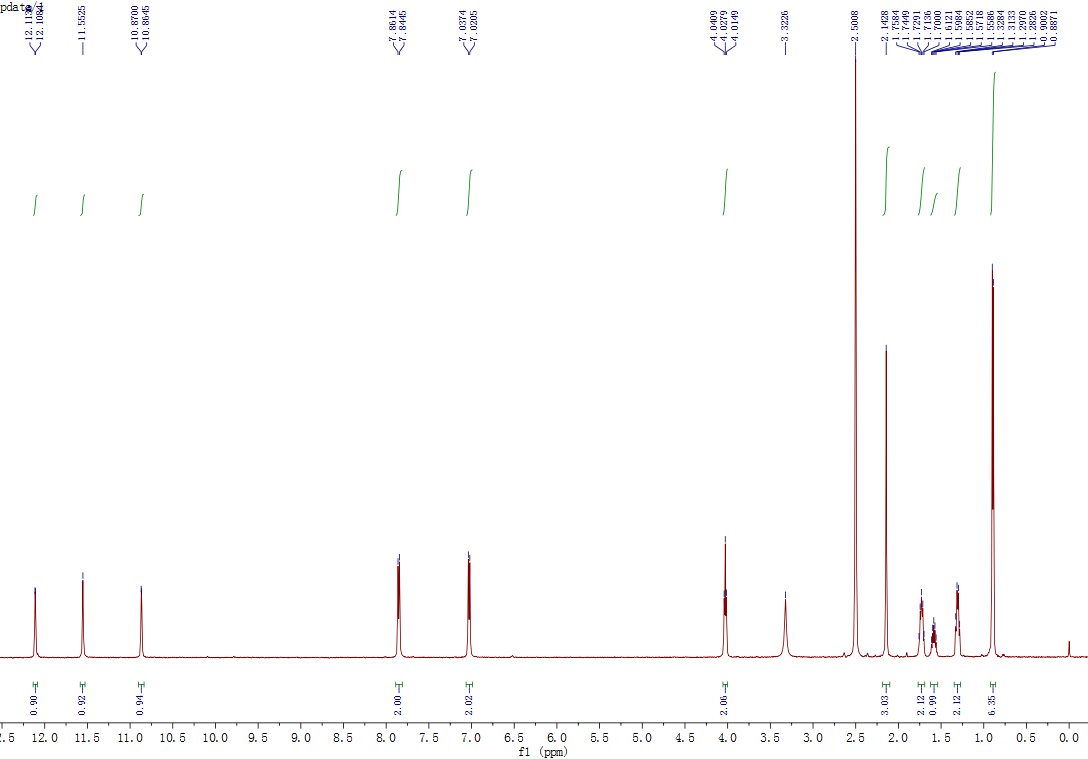
^1^H-NMR spectrum of
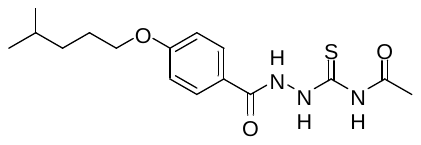
 *N-(2-(4-((4-methylpentyl)oxy)benzoyl)hydrazine-1-carbonothioyl)acetamide (****1e****)*

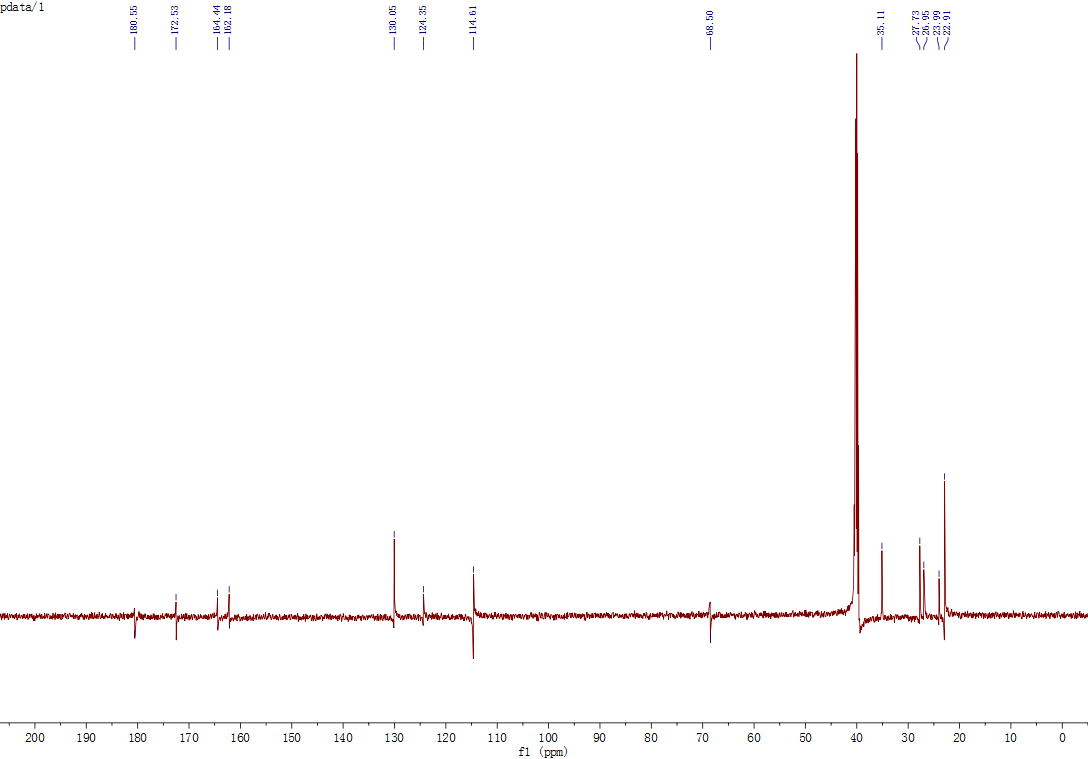

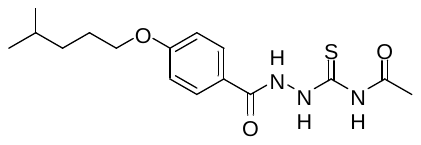
^13^C-NMR spectrum of *N-(2-(4-((4-methylpentyl)oxy)benzoyl)hydrazine-1-carbonothioyl)acetamide (****1e****)*

^1^H-NMR spectrum of *N-(2-(4-(3-methoxypropoxy)benzoyl)hydrazine-1-carbonothioyl)acetamide (****1f****)*

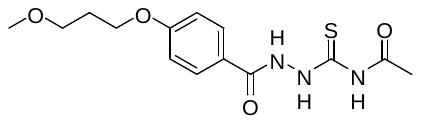

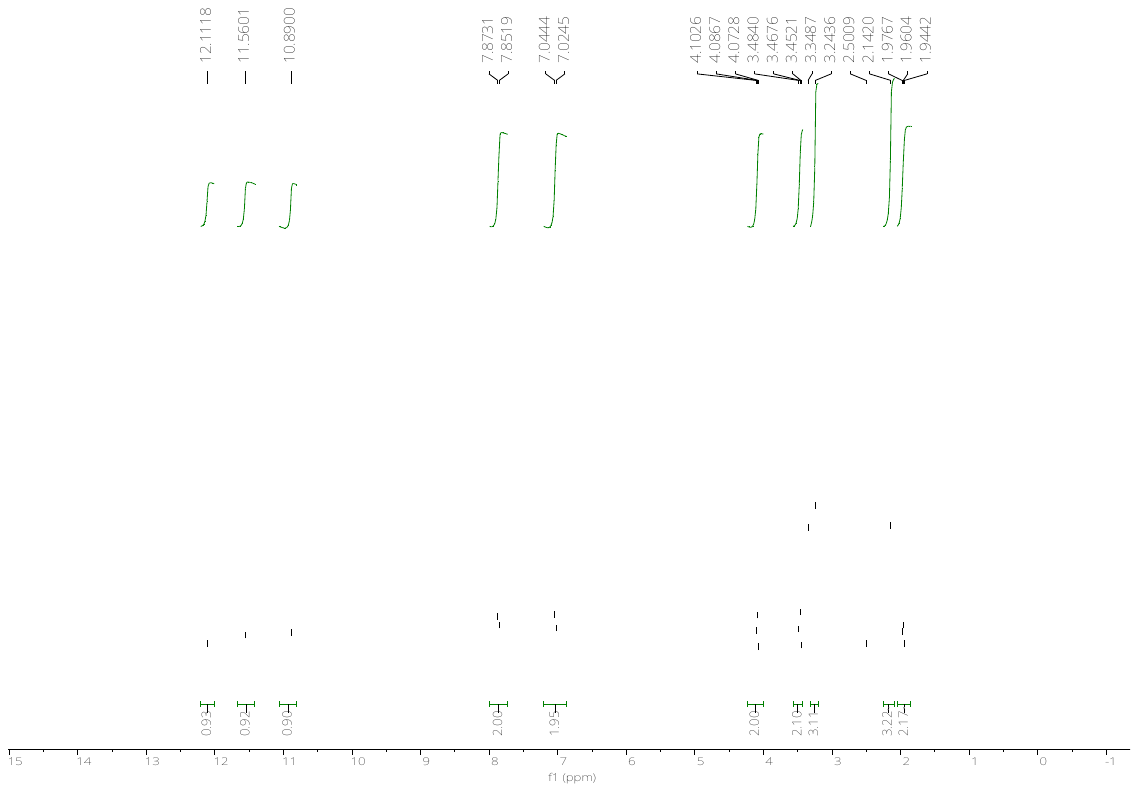

^13^C-NMR spectrum of *N-(2-(4-(3-methoxypropoxy)benzoyl)hydrazine-1-carbonothioyl)acetamide (****1f****)*

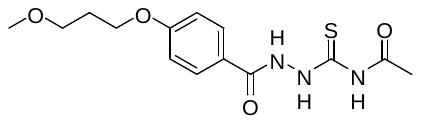

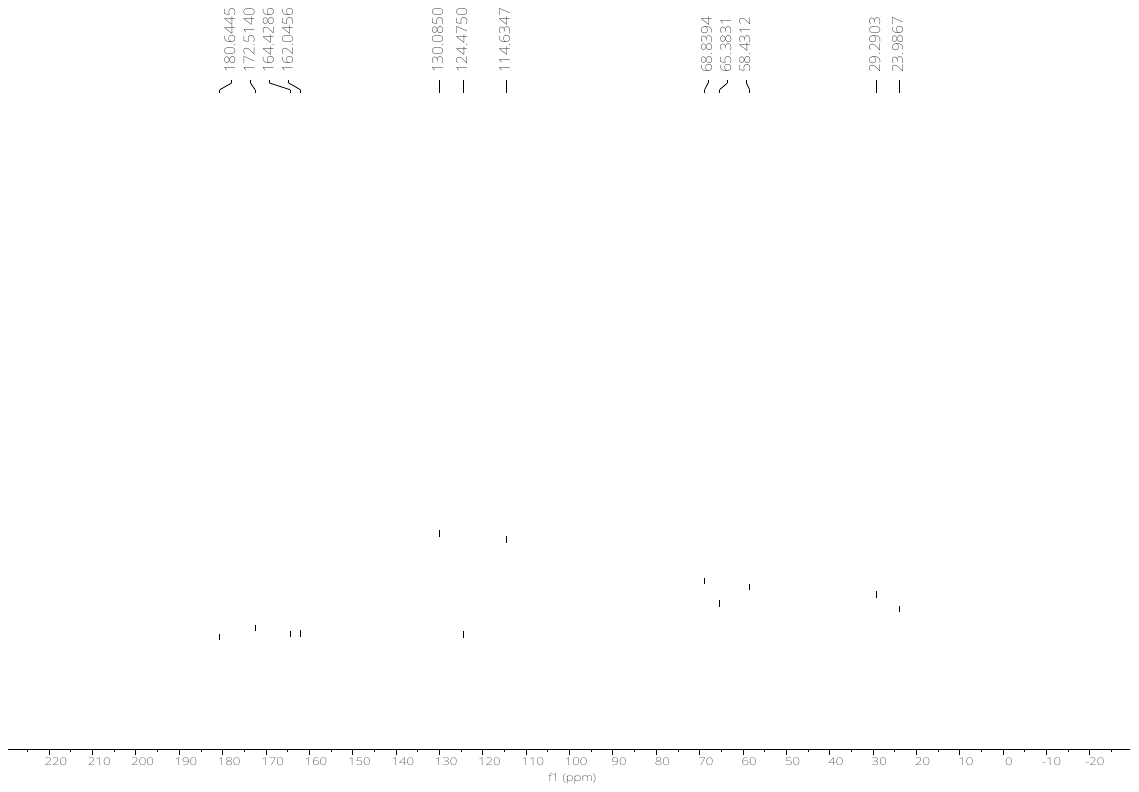

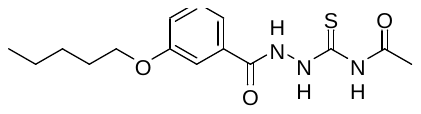
^1^H-NMR spectrum of *N-(2-(3-(n-pentoxy)benzoyl)hydrazine-1-carbonothioyl)acetamide (****1g****)*

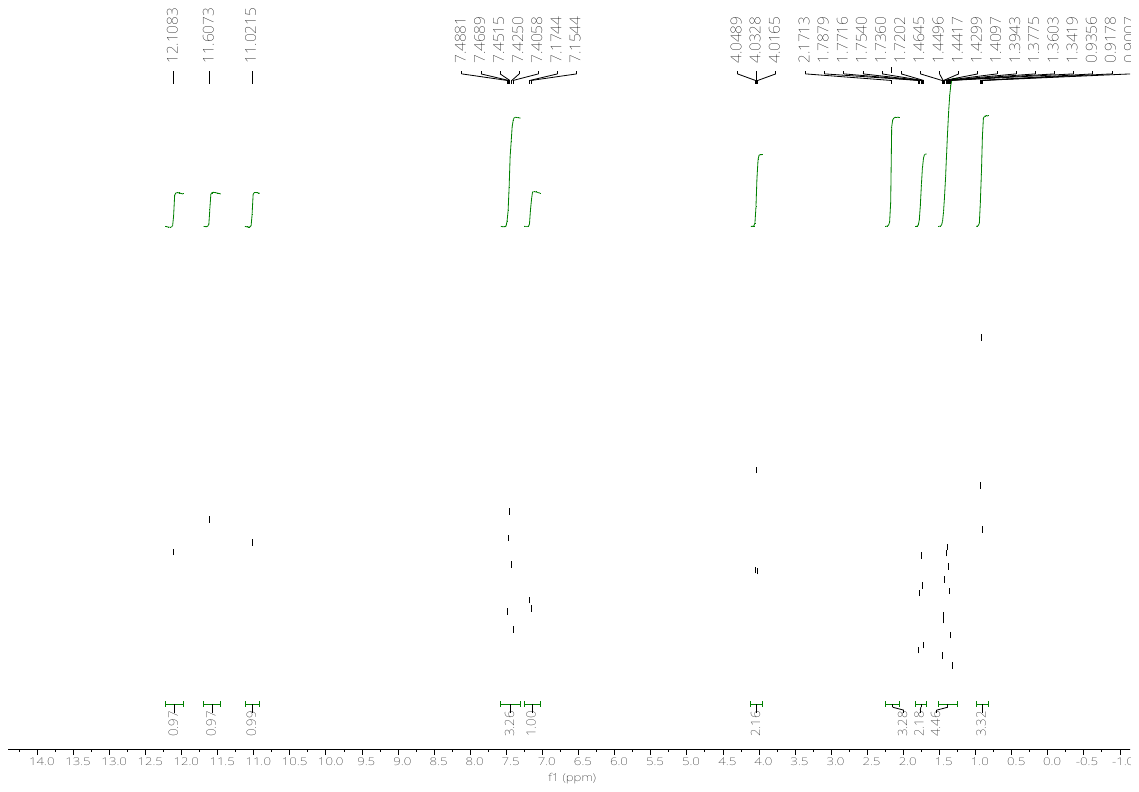

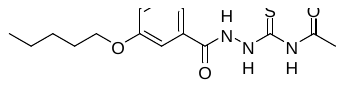
^13^C-NMR spectrum of *N-(2-(3-(n-pentoxy)benzoyl)hydrazine-1-carbonothioyl)acetamide (****1g****)*

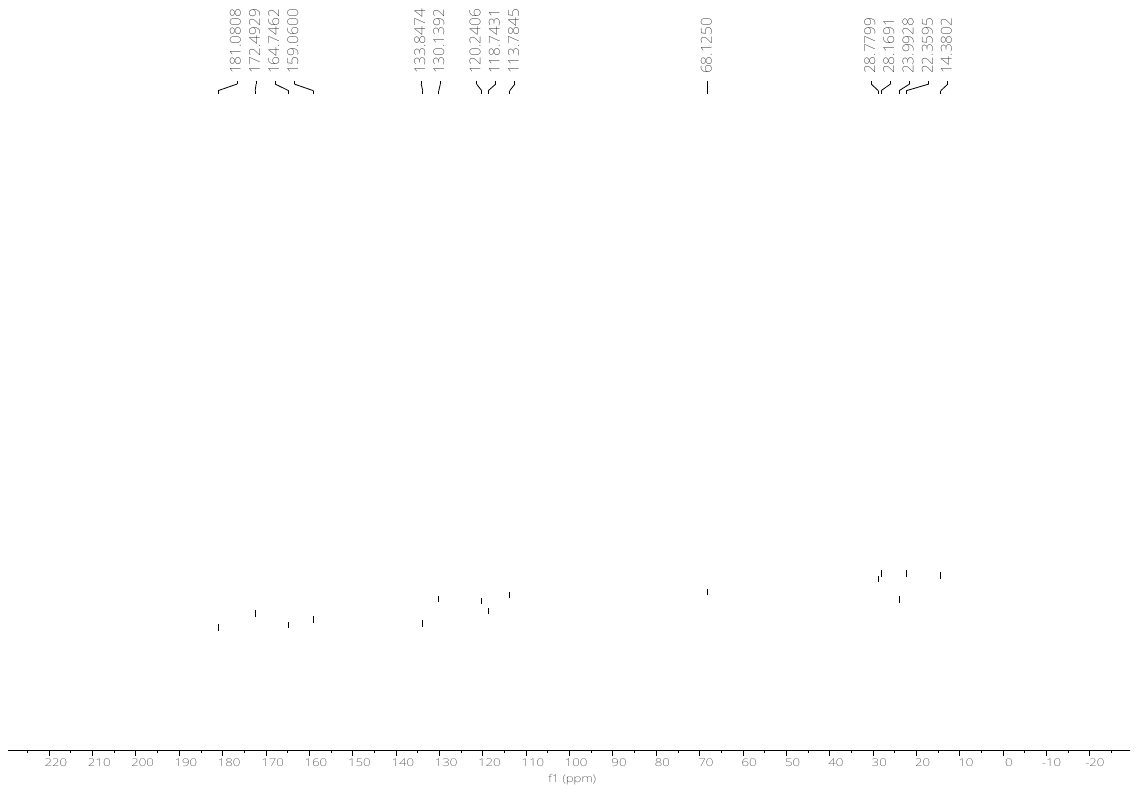

^1^H-NMR spectrum of *N-(2-(3,4-bis(n-pentoxy)benzoyl)hydrazine-1-carbonothioyl)acetamide (****1h****)*

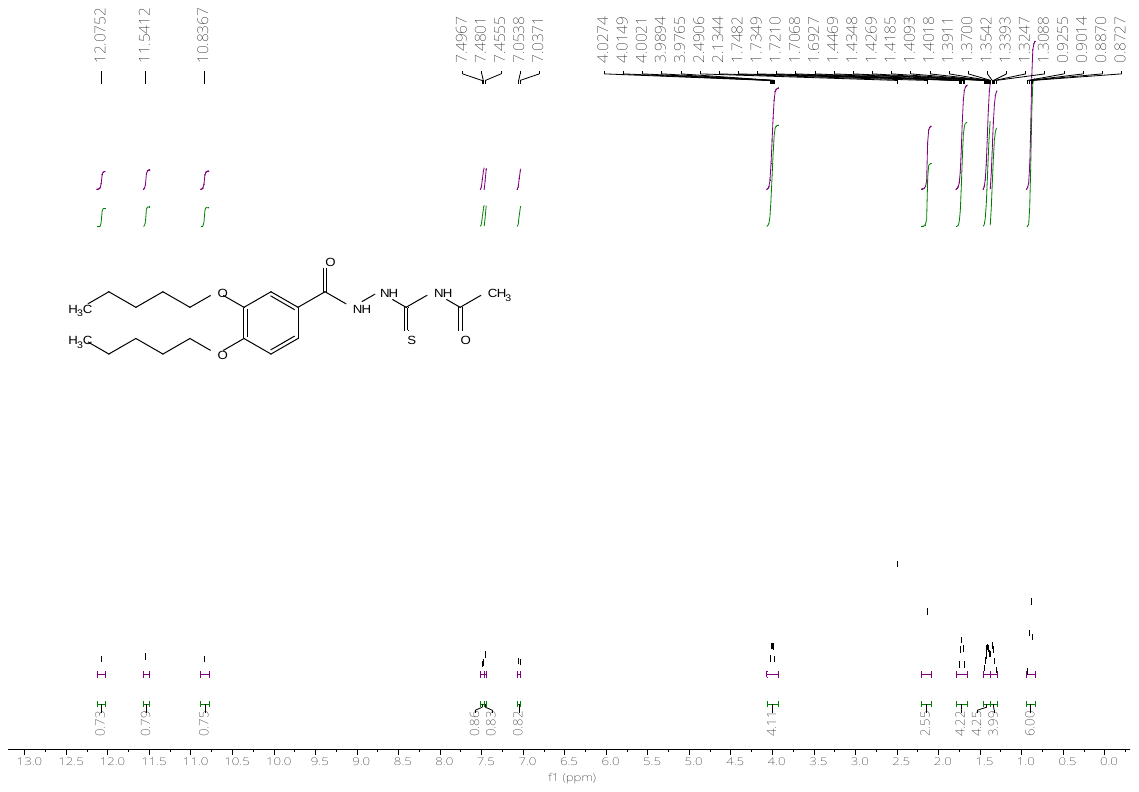

^13^C-NMR spectrum of *N-(2-(3,4-bis(n-pentoxy)benzoyl)hydrazine-1-carbonothioyl)acetamide (****1h****)*

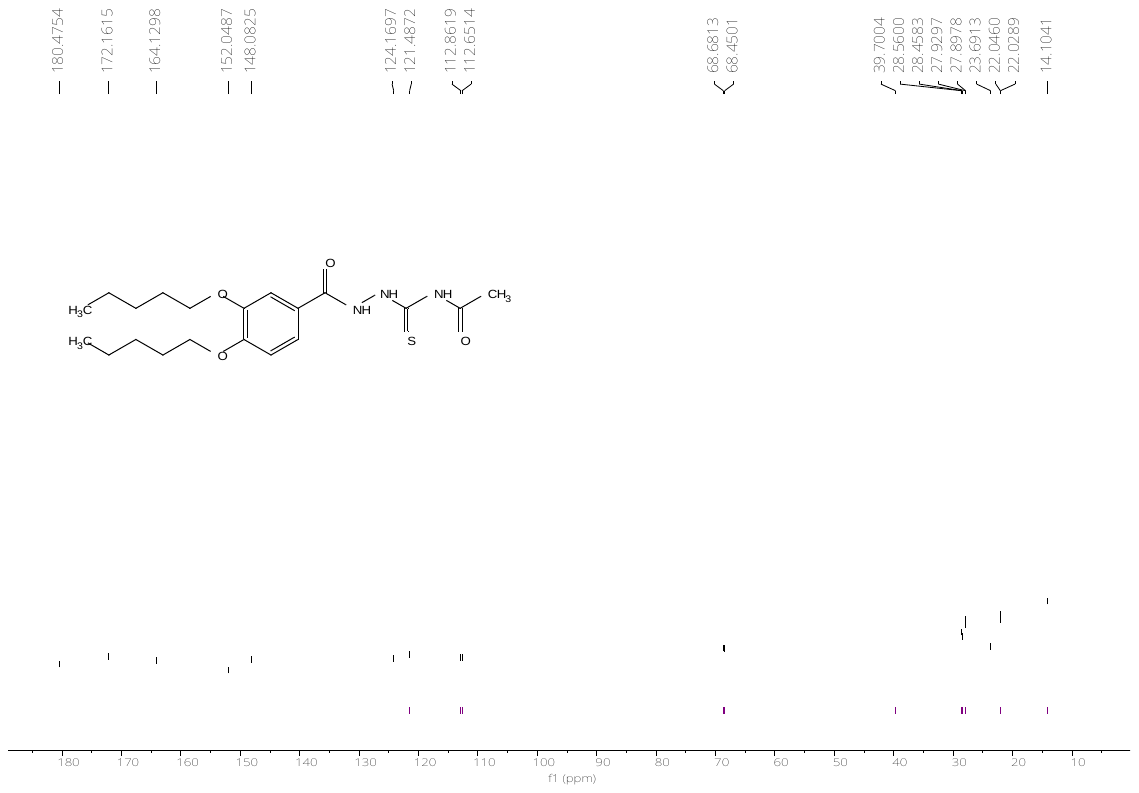

^1^H-NMR spectrum of *N-(2-(3-hydroxy-4-(n-pentoxy)benzoyl)hydrazine-1-carbonothioyl)acetamide (****1i****)*

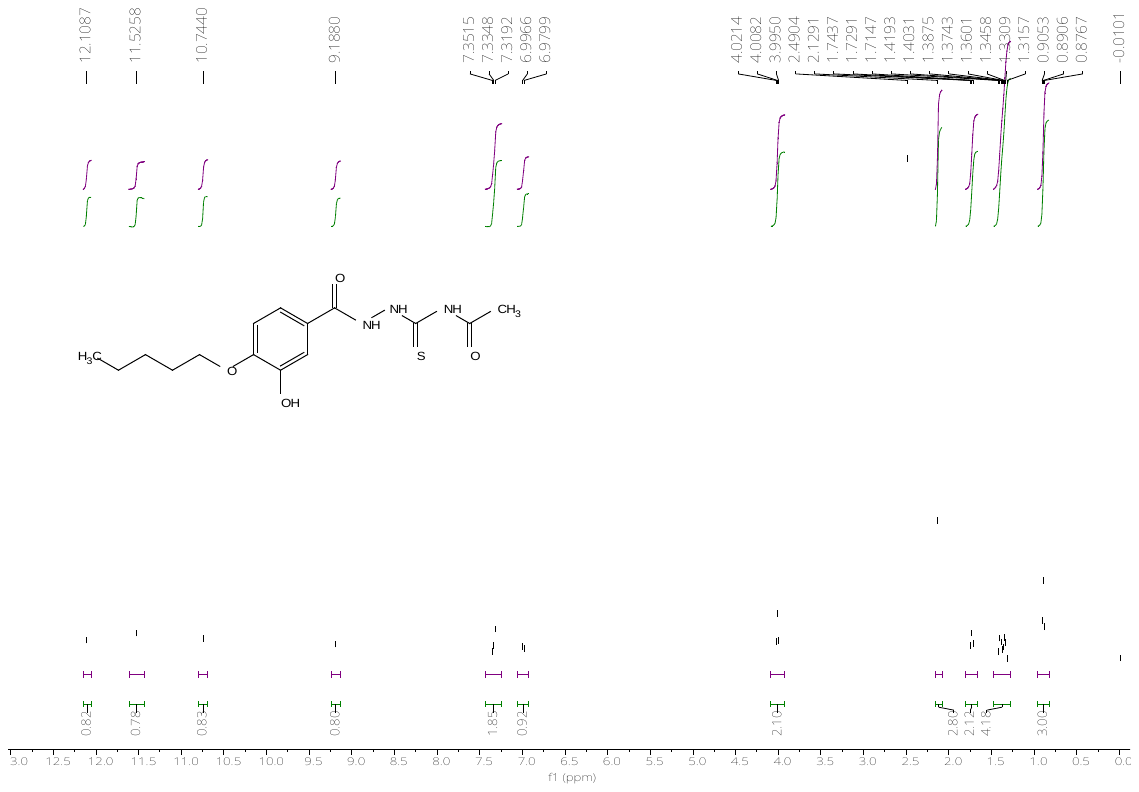

^13^C-NMR spectrum of *N-(2-(3-hydroxy-4-(n-pentoxy)benzoyl)hydrazine-1-carbonothioyl)acetamide (****1i****)*

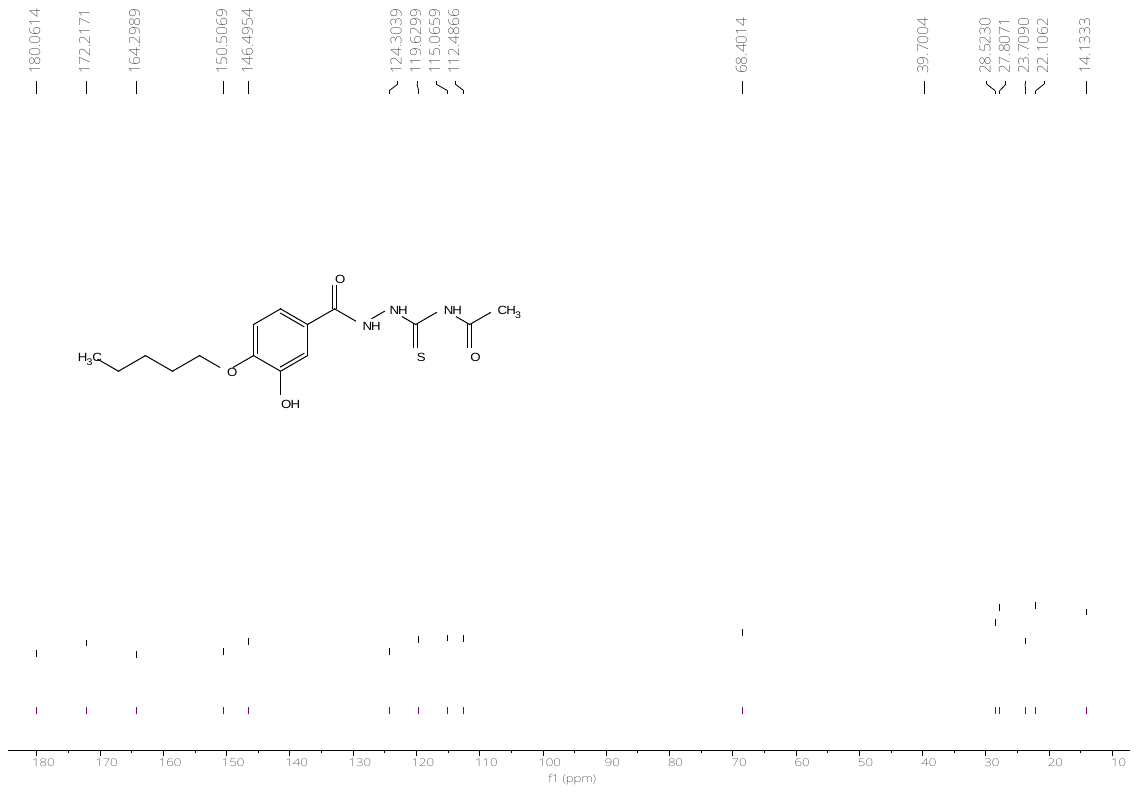

^1^H-NMR spectrum of *N-(2-(3,4-dihydroxybenzoyl)hydrazine-1-carbonothioyl)acetamide (****1j****)*

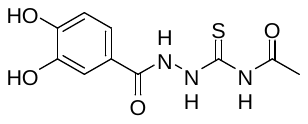

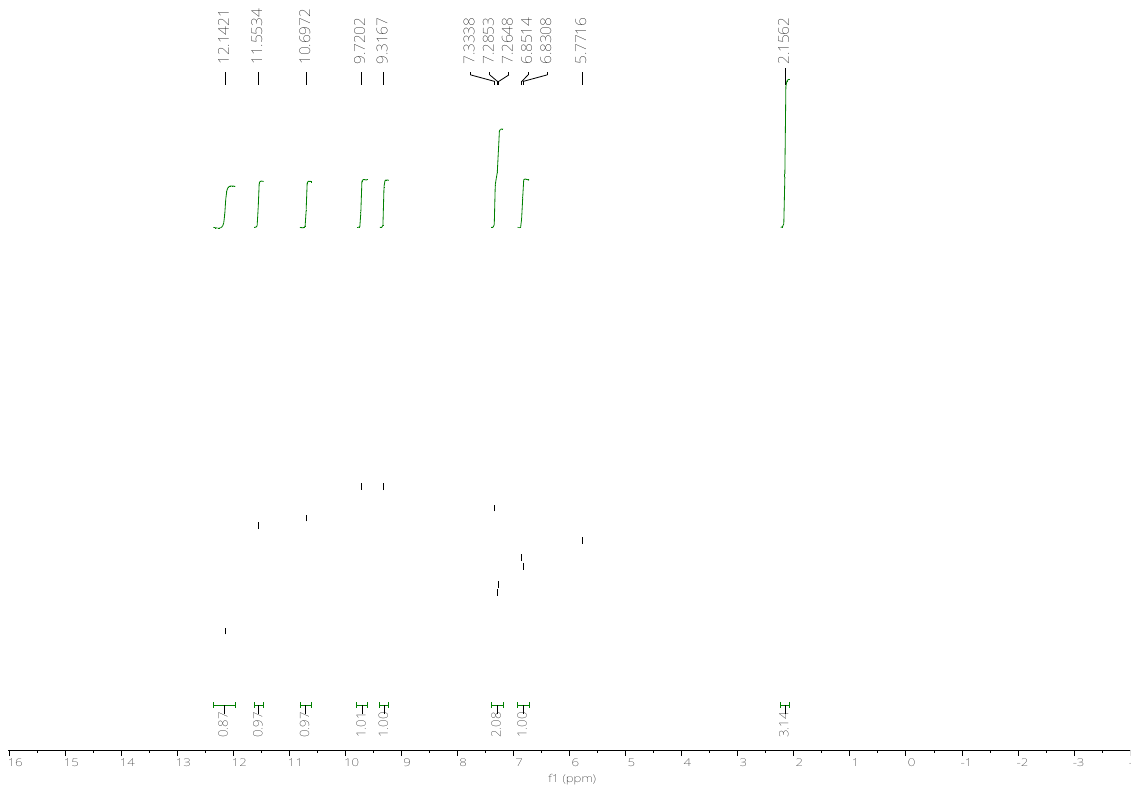

^13^C-NMR spectrum of *N-(2-(3,4-dihydroxybenzoyl)hydrazine-1-carbonothioyl)acetamide (****1j****)*

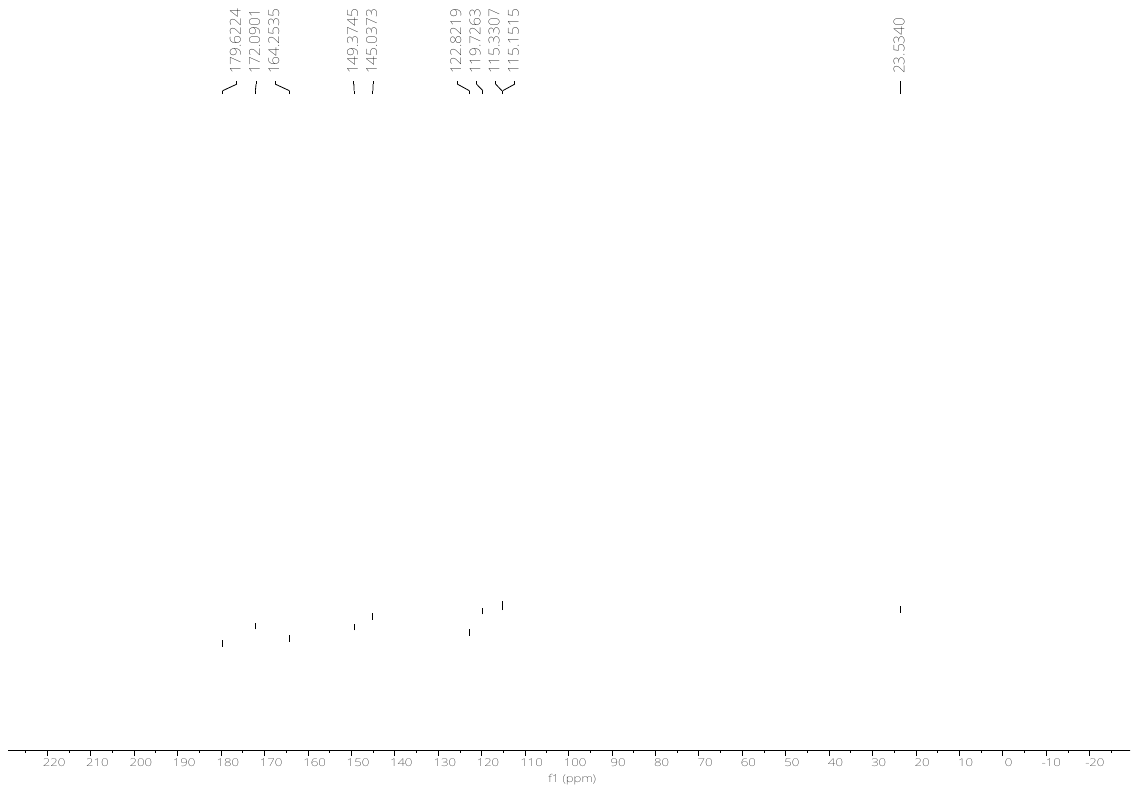

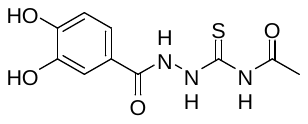

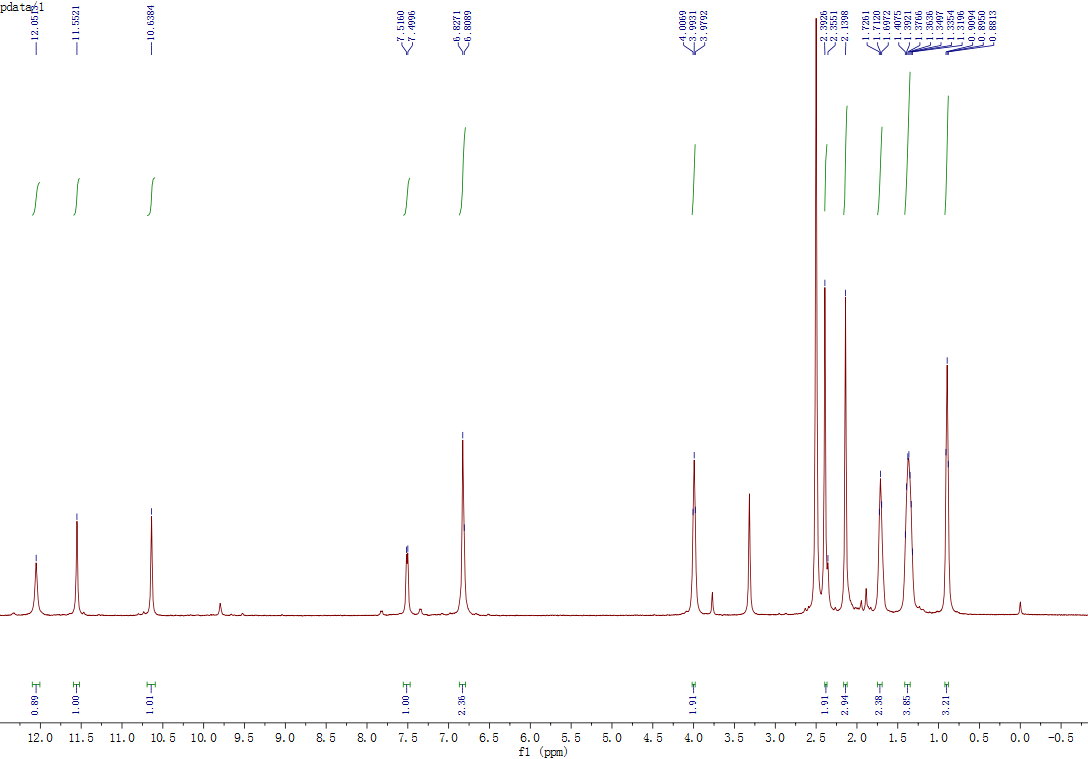

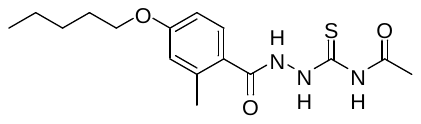
^1^H-NMR spectrum of *N-(2-(2-methyl-4-(n-pentoxy)benzoyl)hydrazine-1-carbonothioyl)acetamide (****1k****)*

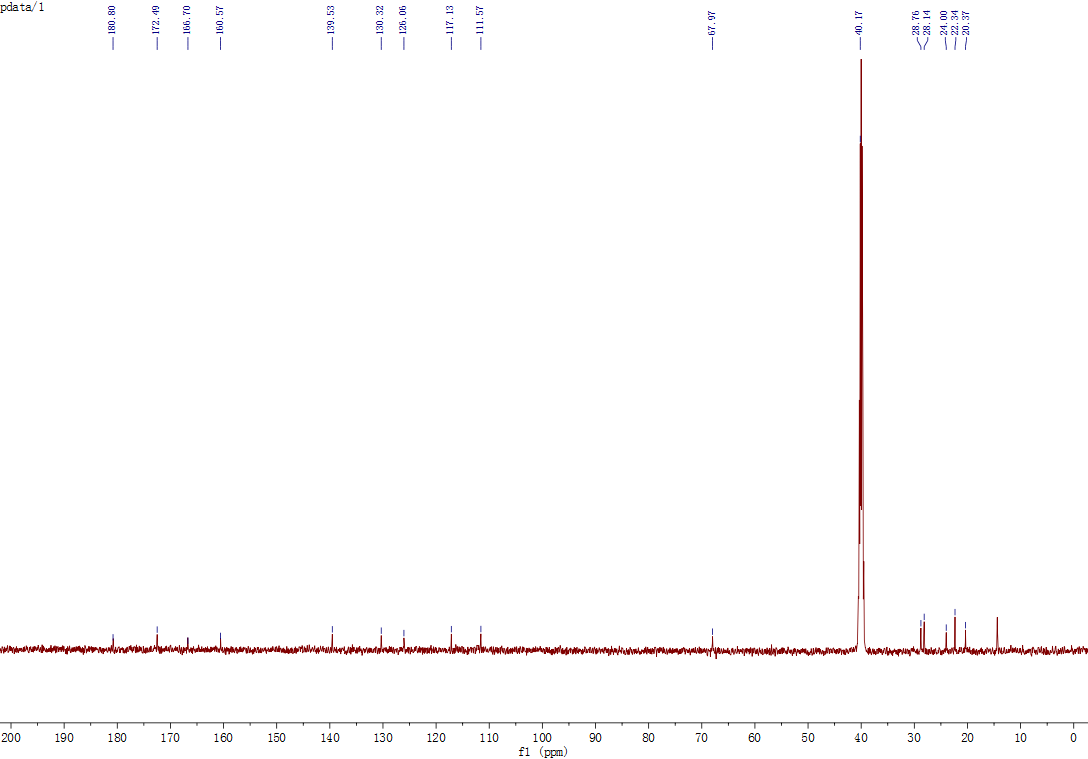

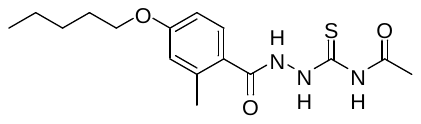
^13^C-NMR spectrum of *N-(2-(2-methyl-4-(n-pentoxy)benzoyl)hydrazine-1-carbonothioyl)acetamide (****1k****)*

^1^H-NMR spectrum of *N-(2-(2,4-bis(n-pentoxy)benzoyl)hydrazine-1-carbonothioyl)acetamide (****1l****)*

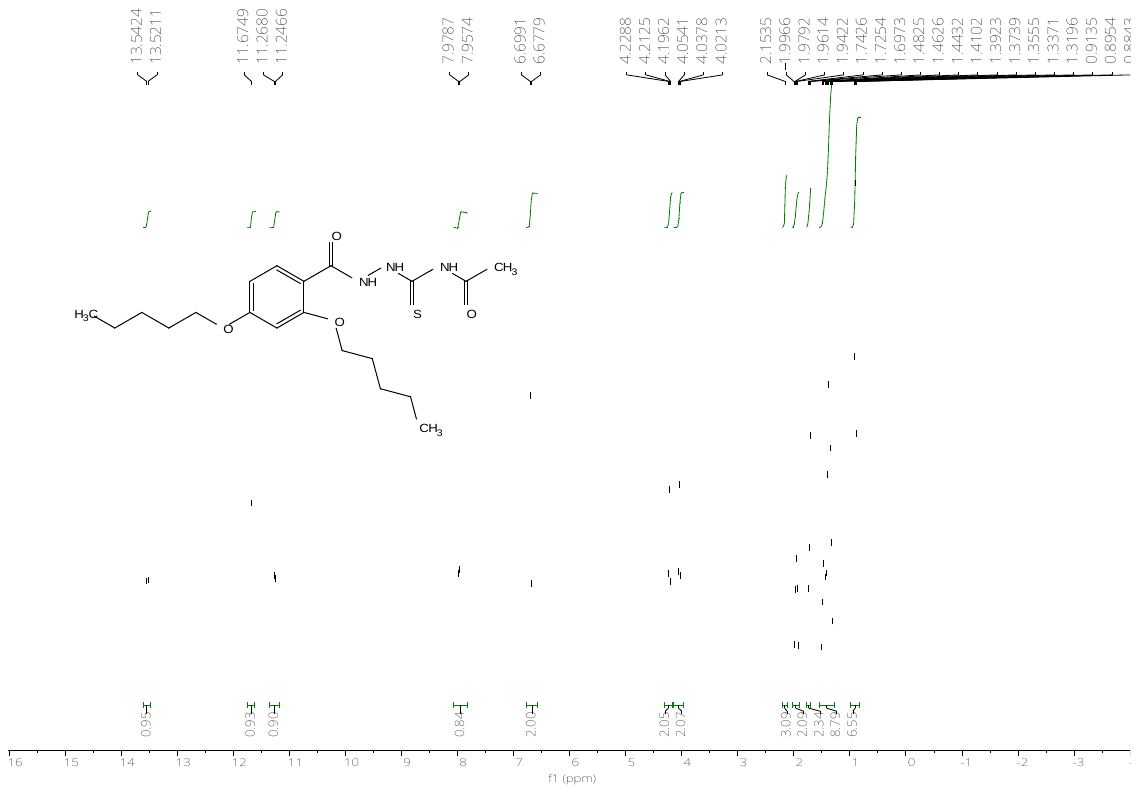

^13^C-NMR spectrum of *N-(2-(2,4-bis(n-pentoxy)benzoyl)hydrazine-1-carbonothioyl)acetamide (****1l****)*

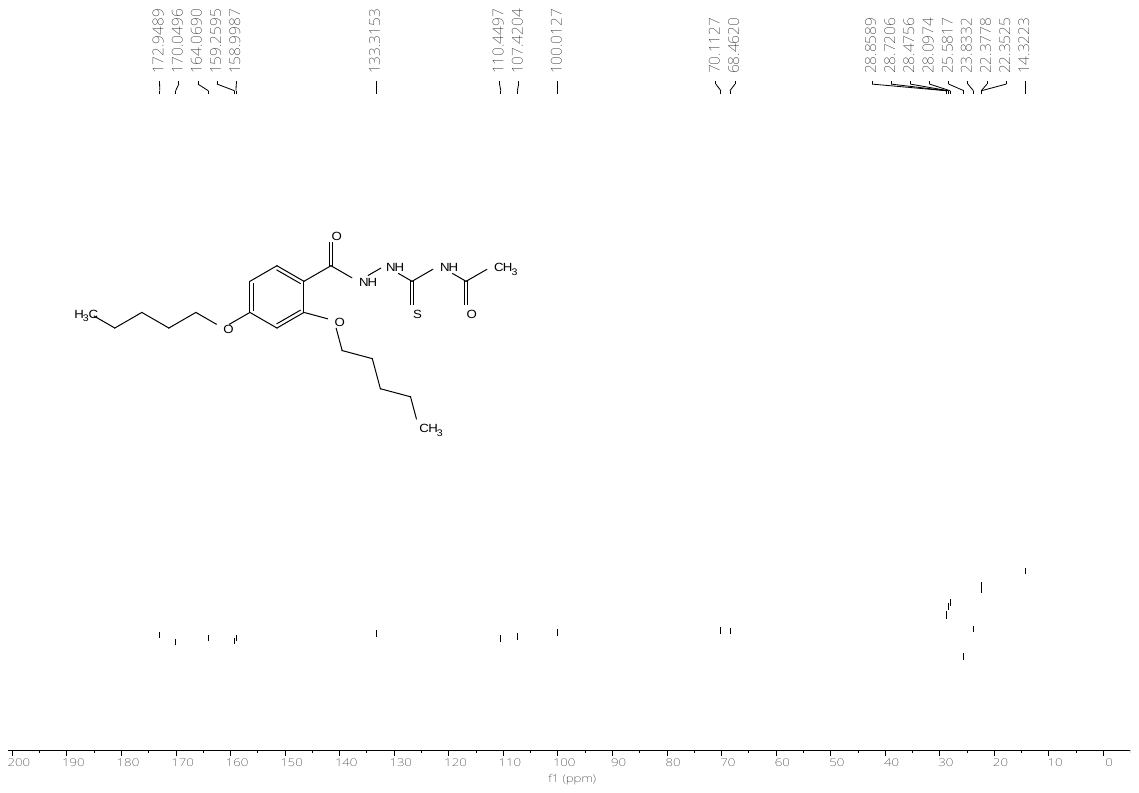

^1^H-NMR spectrum of *N-(2-(4-(cyclohexyloxy)benzoyl)hydrazine-1-carbonothioyl)acetamide (****1m****)*
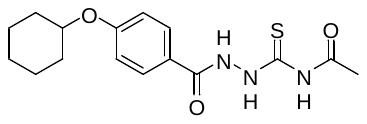

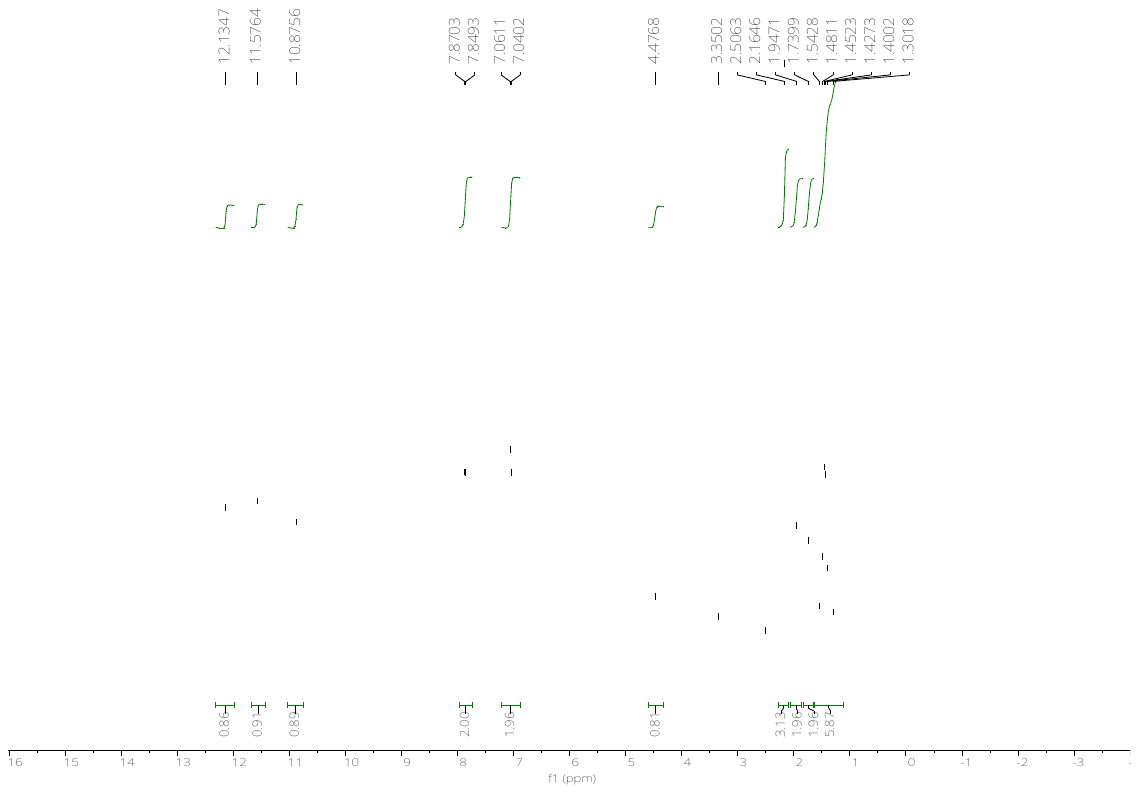

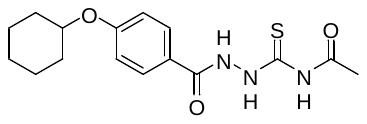
^13^C-NMR spectrum of *N-(2-(4-(cyclohexyloxy)benzoyl)hydrazine-1-carbonothioyl)acetamide (****1m****)*

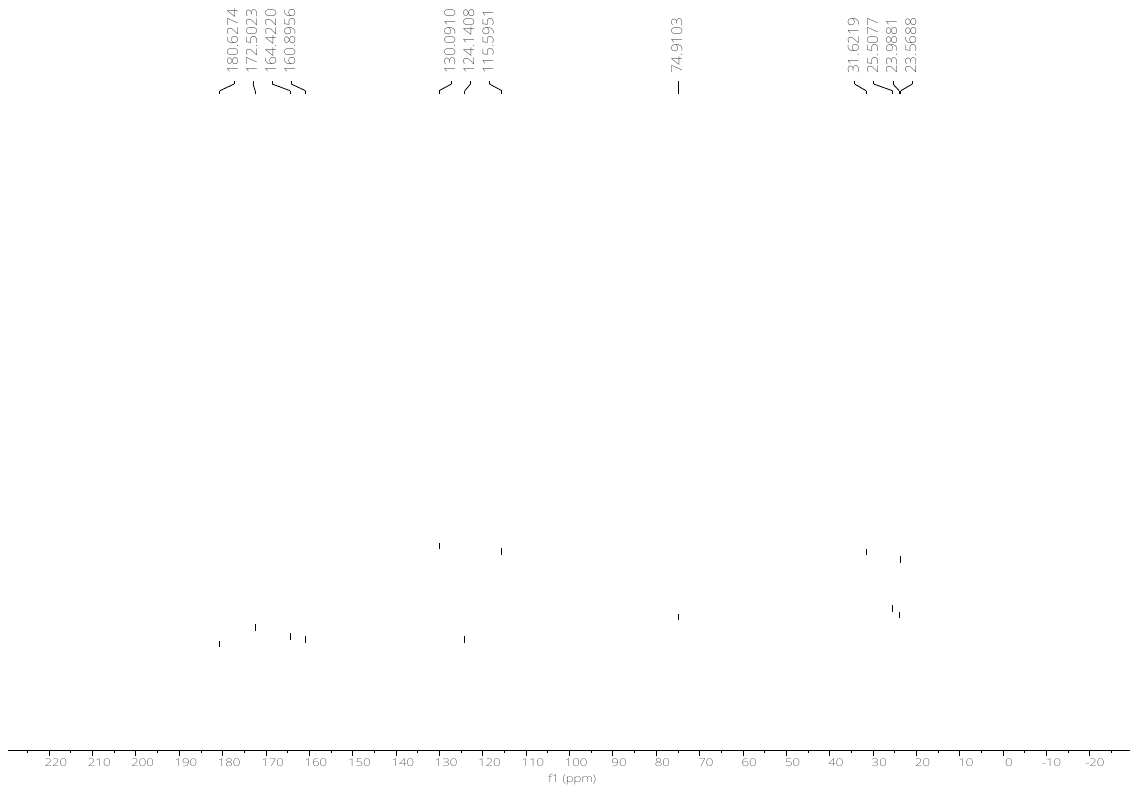

^1^H-NMR spectrum of *N-(2-(4-phenoxybenzoyl)hydrazine-1-carbonothioyl)acetamide (****1n****)*

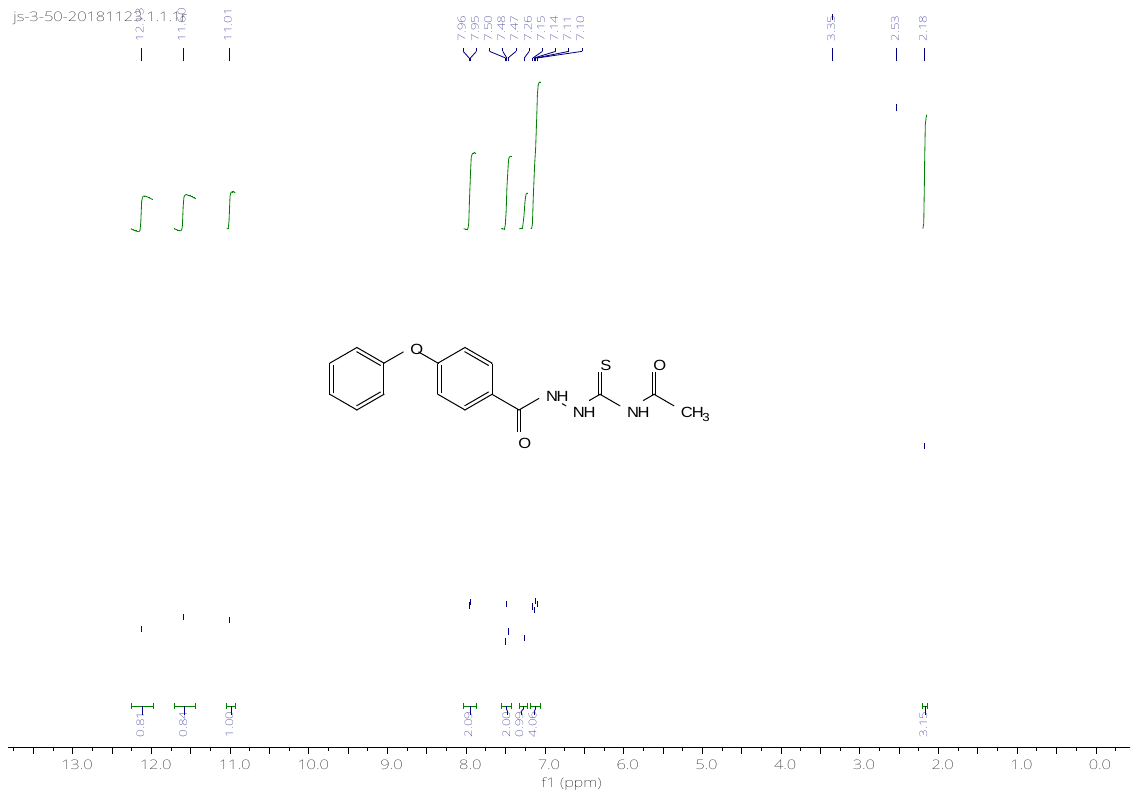

^13^C-NMR spectrum of *N-(2-(4-phenoxybenzoyl)hydrazine-1-carbonothioyl)acetamide (****1n****)*

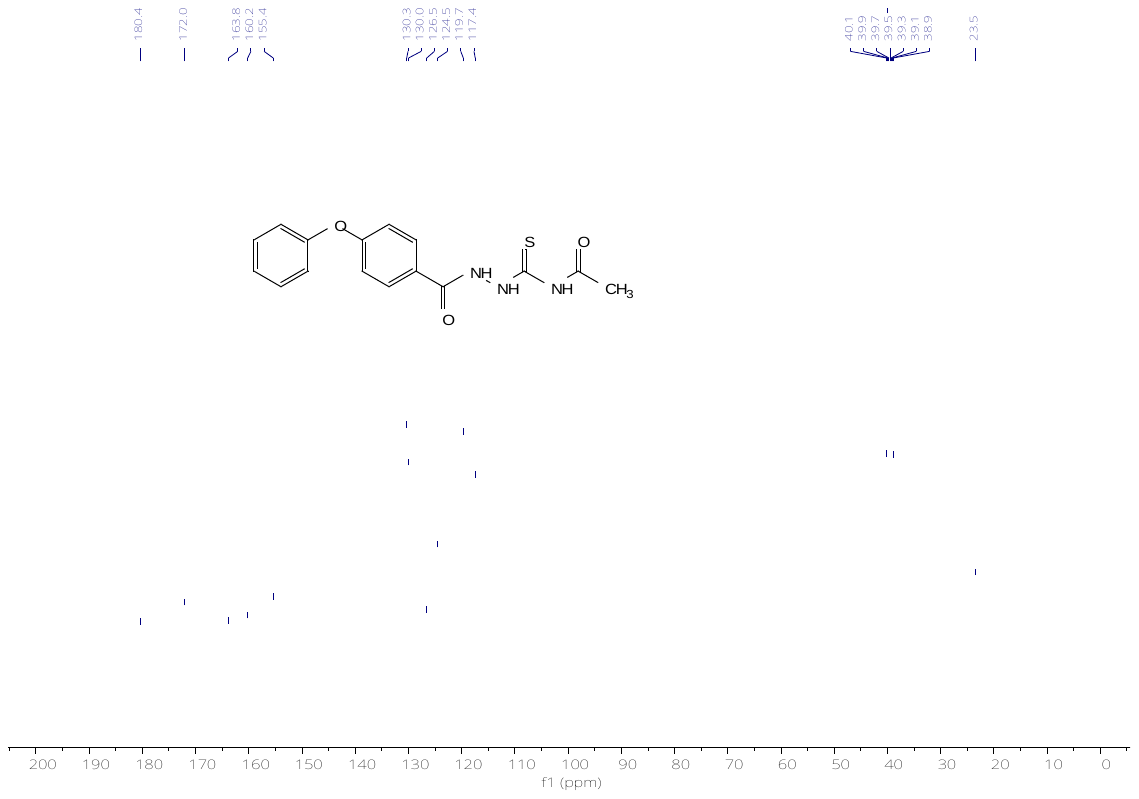

^1^H-NMR spectrum of *N-(2-(5-(n-pentoxy)picolinoyl)hydrazine-1-carbonothioyl)acetamide (****1o****)*

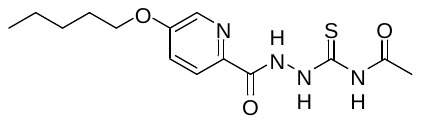

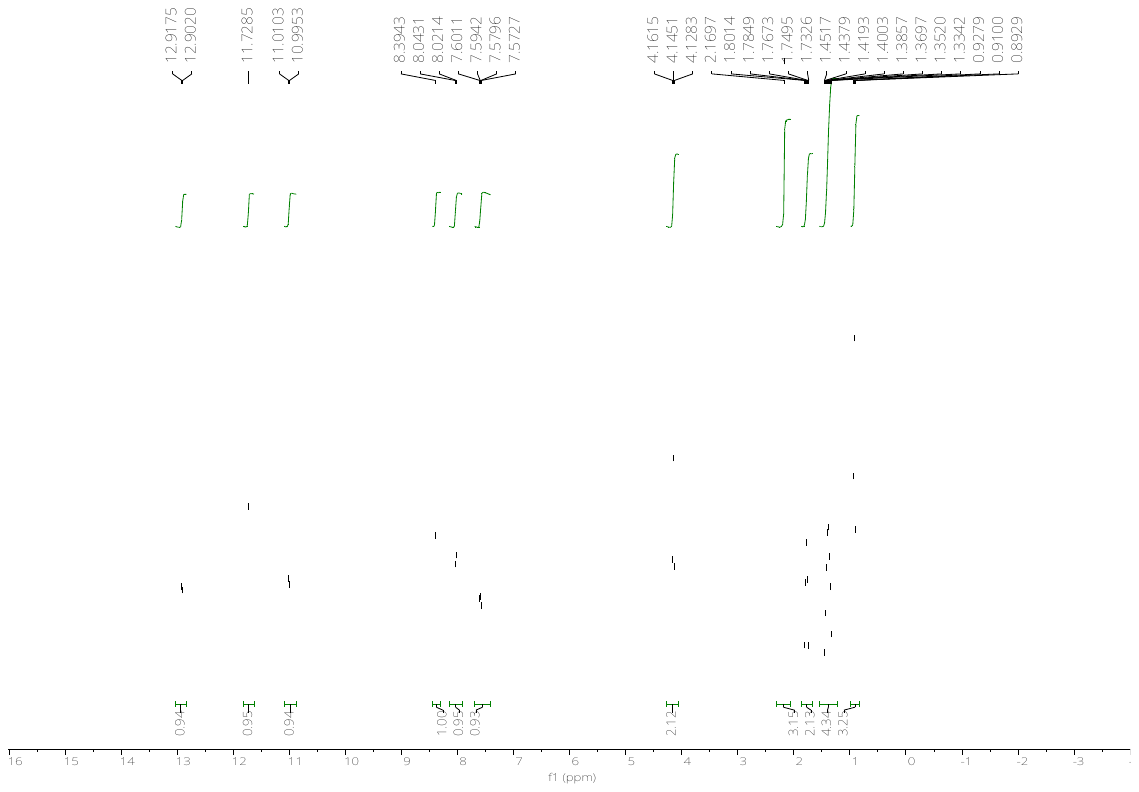

^13^C-NMR spectrum of *N-(2-(5-(n-pentoxy)picolinoyl)hydrazine-1-carbonothioyl)acetamide (****1o****)*

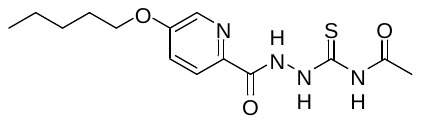

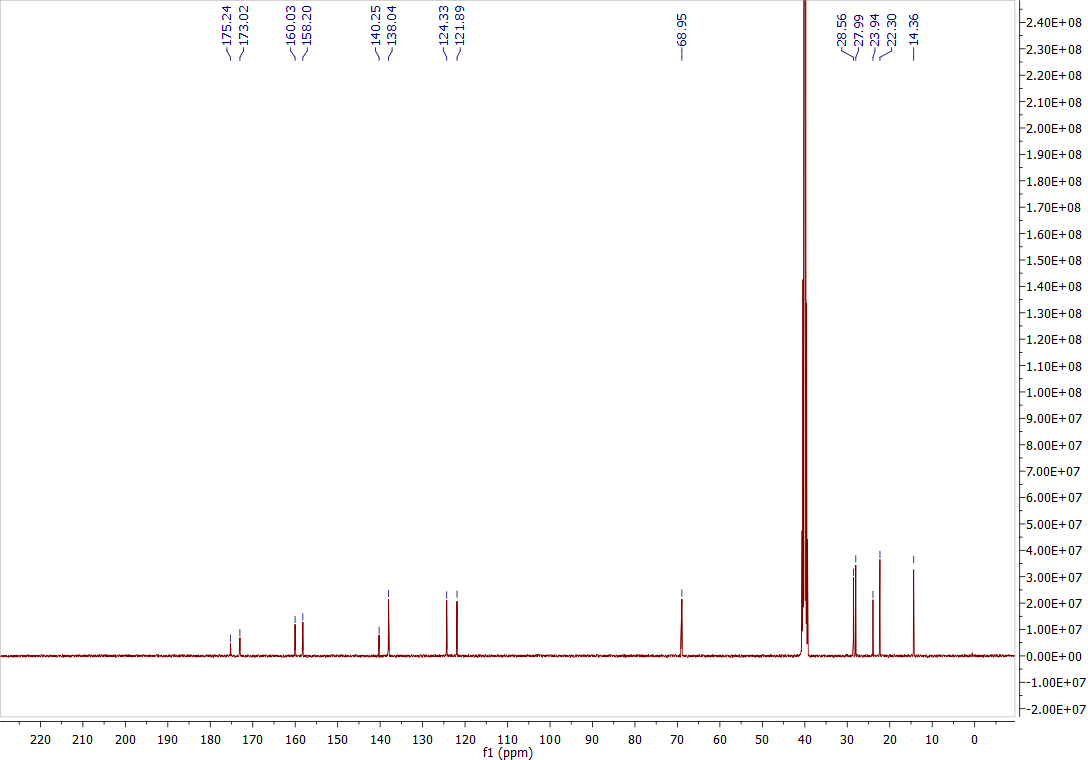

^1^H-NMR spectrum of *N-(2-(2-(4-(n-pentoxy)phenyl)acetyl)hydrazine-1-carbonothioyl)acetamide (****1p****)*

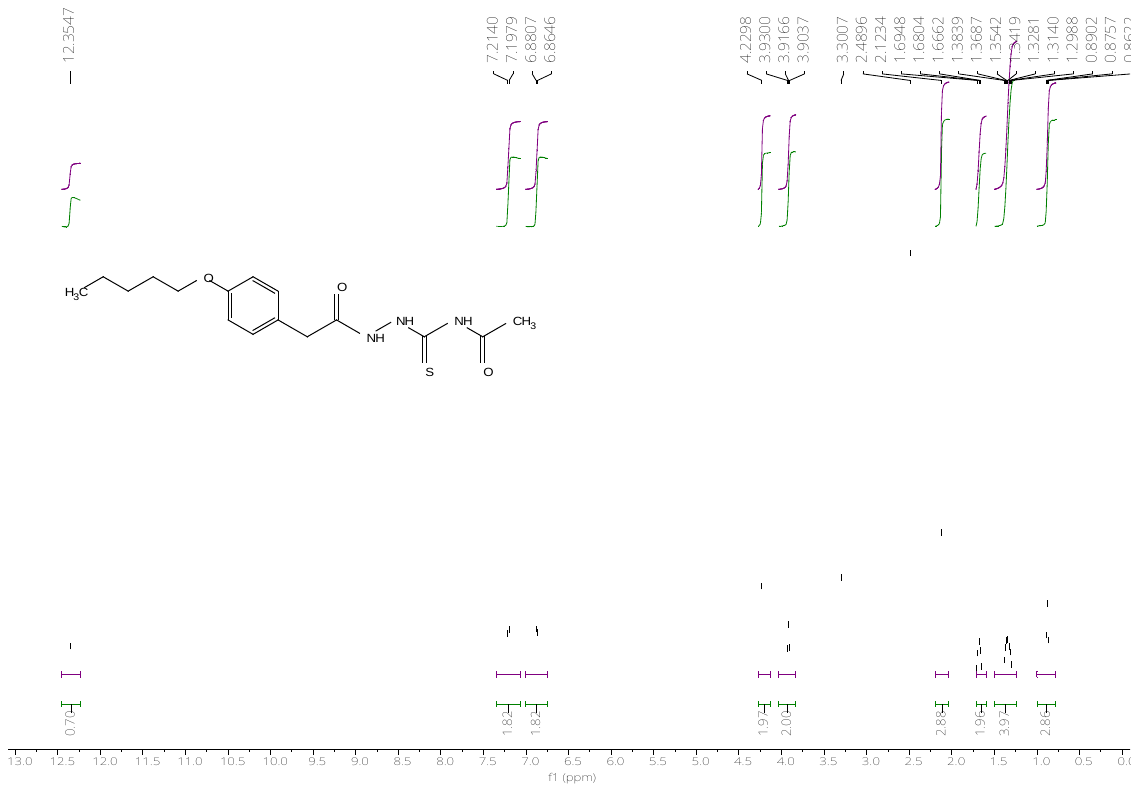

^13^C-NMR spectrum of *N-(2-(2-(4-(n-pentoxy)phenyl)acetyl)hydrazine-1-carbonothioyl)acetamide (****1p****)*

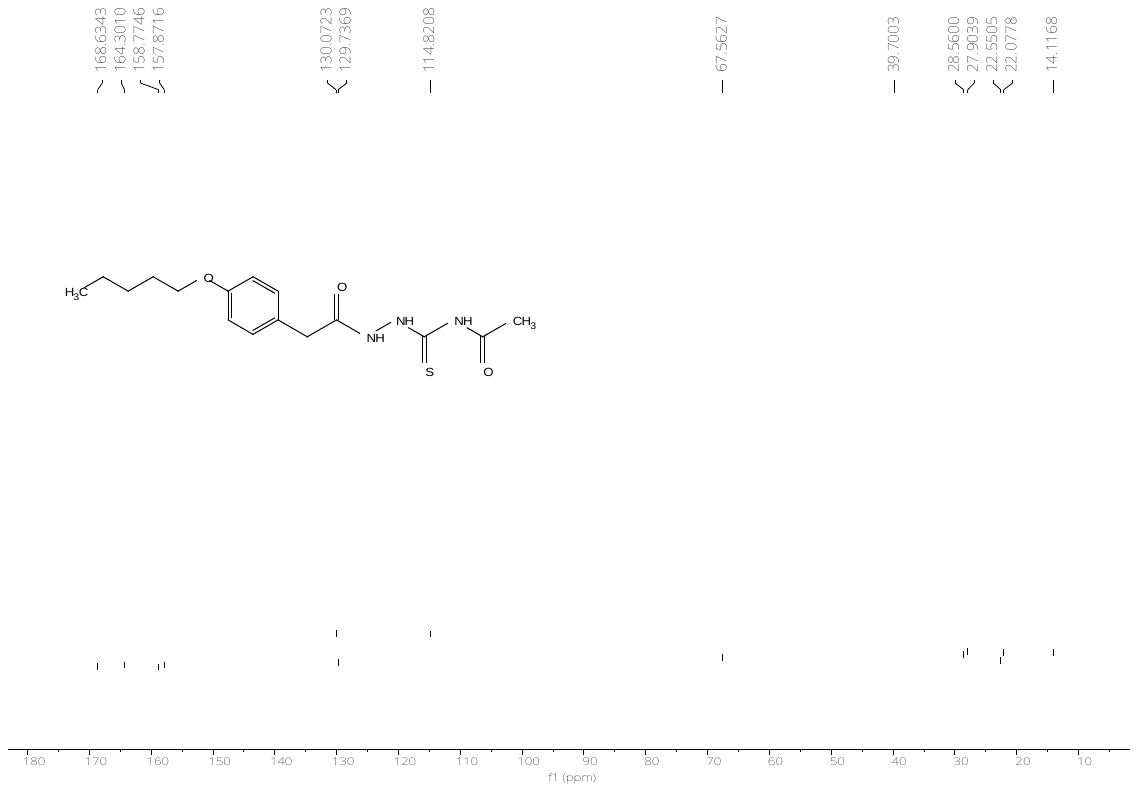

^1^H-NMR spectrum of *N-(2-(3-(4-(n-pentoxy)phenyl)propanoyl)hydrazine-1-carbonothioyl)acetamide (****1q****)*

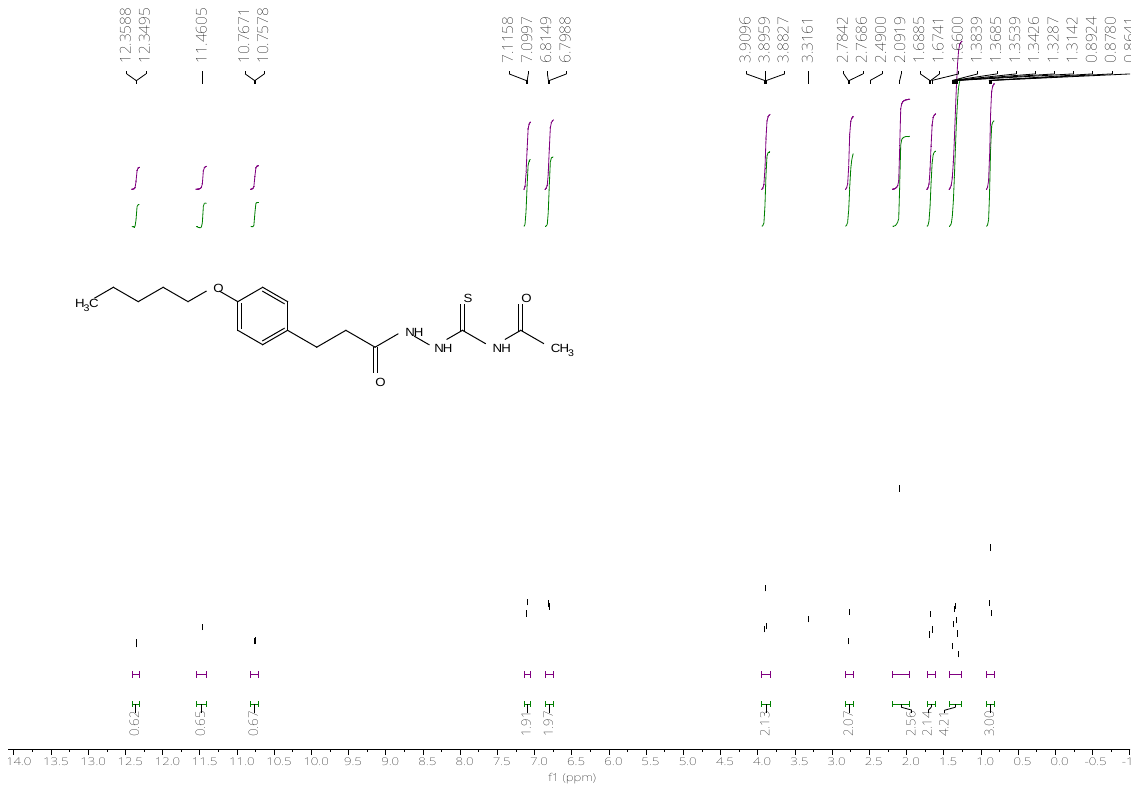

^13^C-NMR spectrum of *N-(2-(3-(4-(n-pentoxy)phenyl)propanoyl)hydrazine-1-carbonothioyl)acetamide (****1q****)*

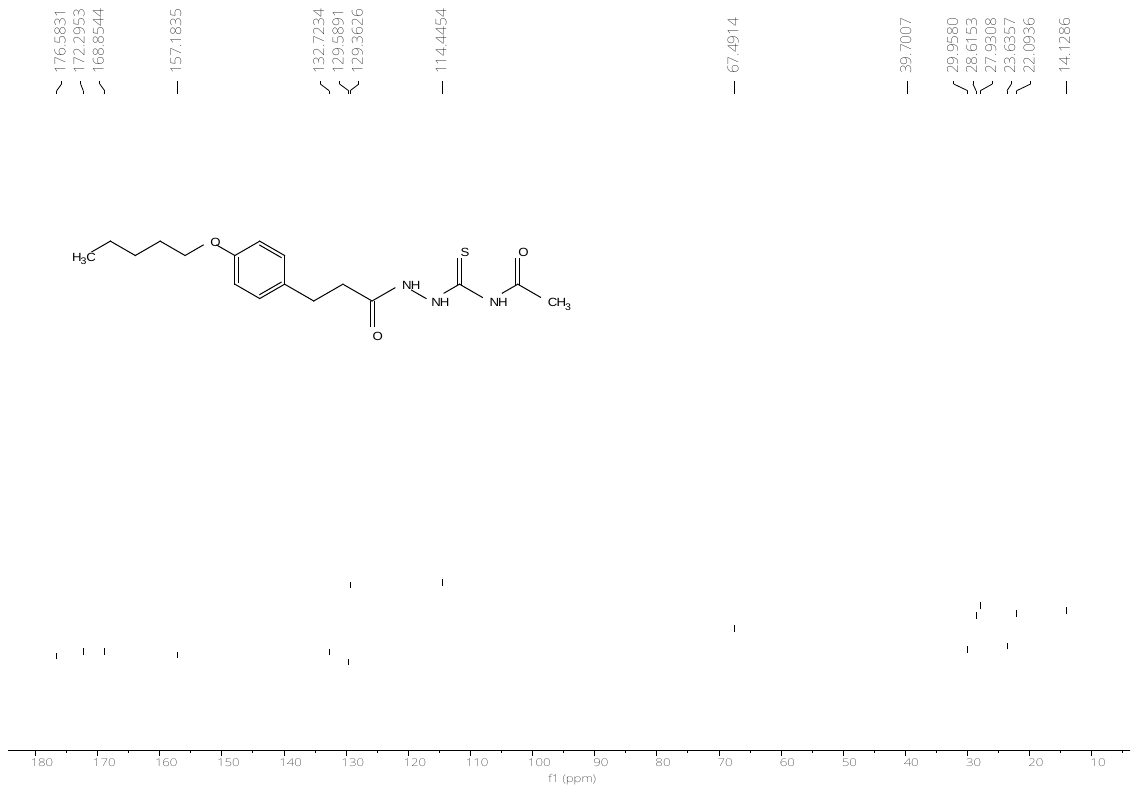

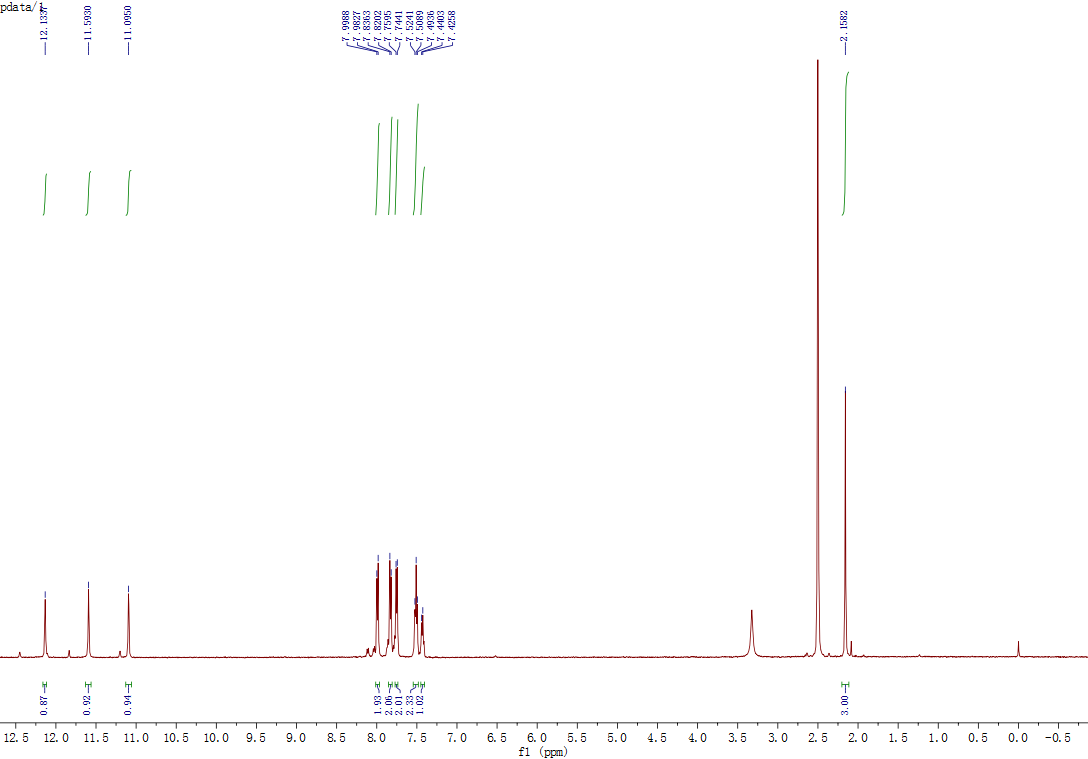

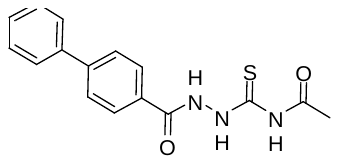
^1^H-NMR spectrum of *N-(2-([1,1'-biphenyl]-4-carbonyl)hydrazine-1-carbonothioyl)acetamide (****1r****)*

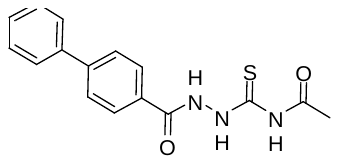

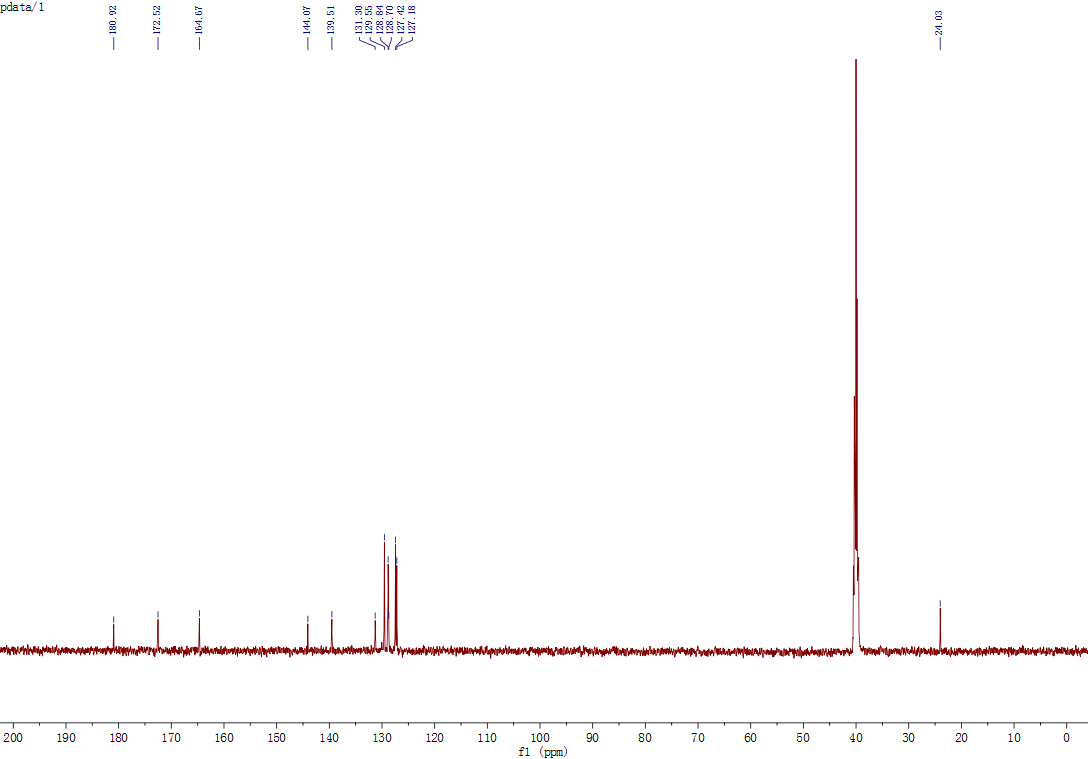
^13^C-NMR spectrum of *N-(2-([1,1'-biphenyl]-4-carbonyl)hydrazine-1-carbonothioyl)acetamide (****1r****)*

^1^H-NMR spectrum of *N-(2-([1,1'-biphenyl]-3-carbonyl)hydrazine-1-carbonothioyl)acetamide (****1s****)*
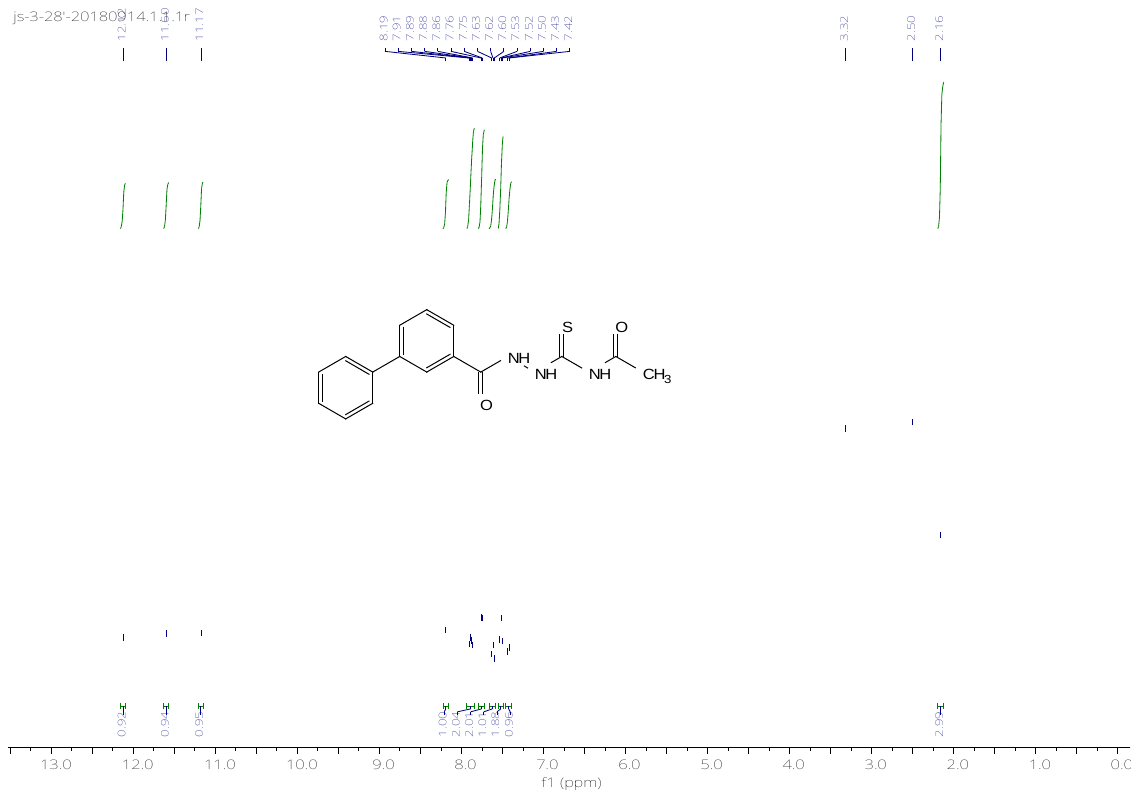

^13^C-NMR spectrum of *N-(2-([1,1'-biphenyl]-3-carbonyl)hydrazine-1-carbonothioyl)acetamide (****1s****)*

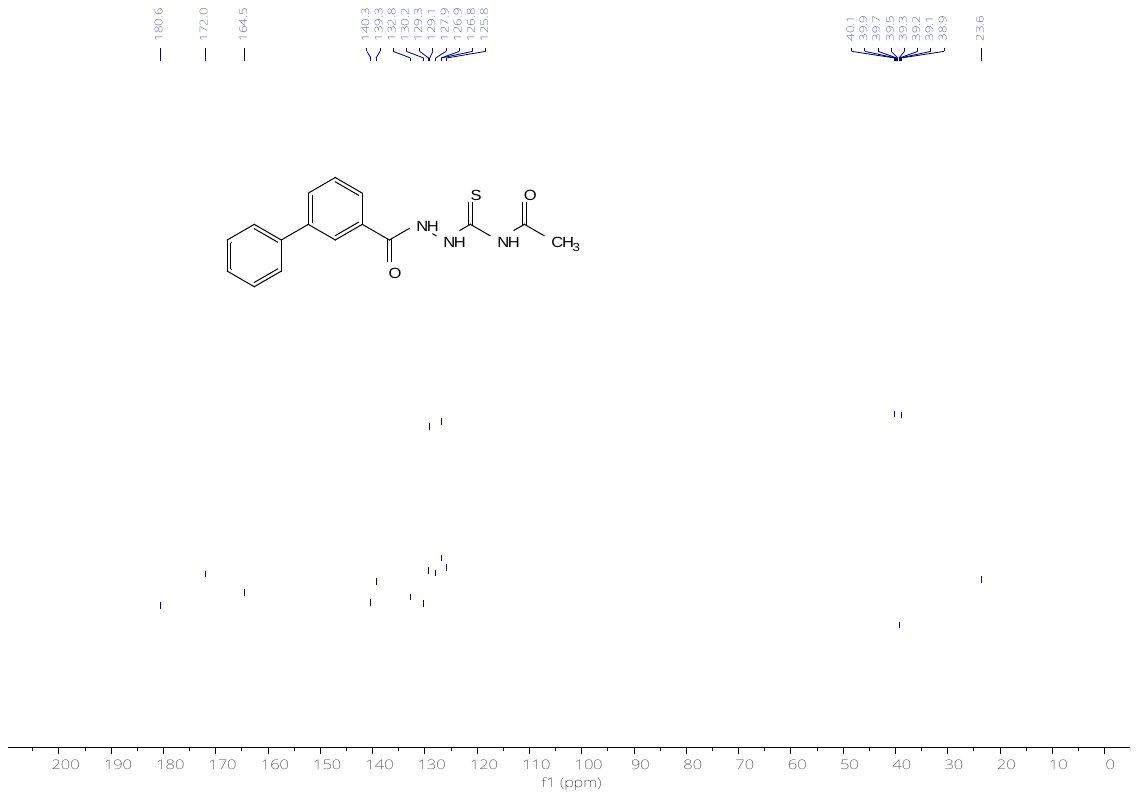

^1^H-NMR spectrum of *N-(2-(2-naphthoyl)hydrazine-1-carbonothioyl)acetamide (****1t****)*

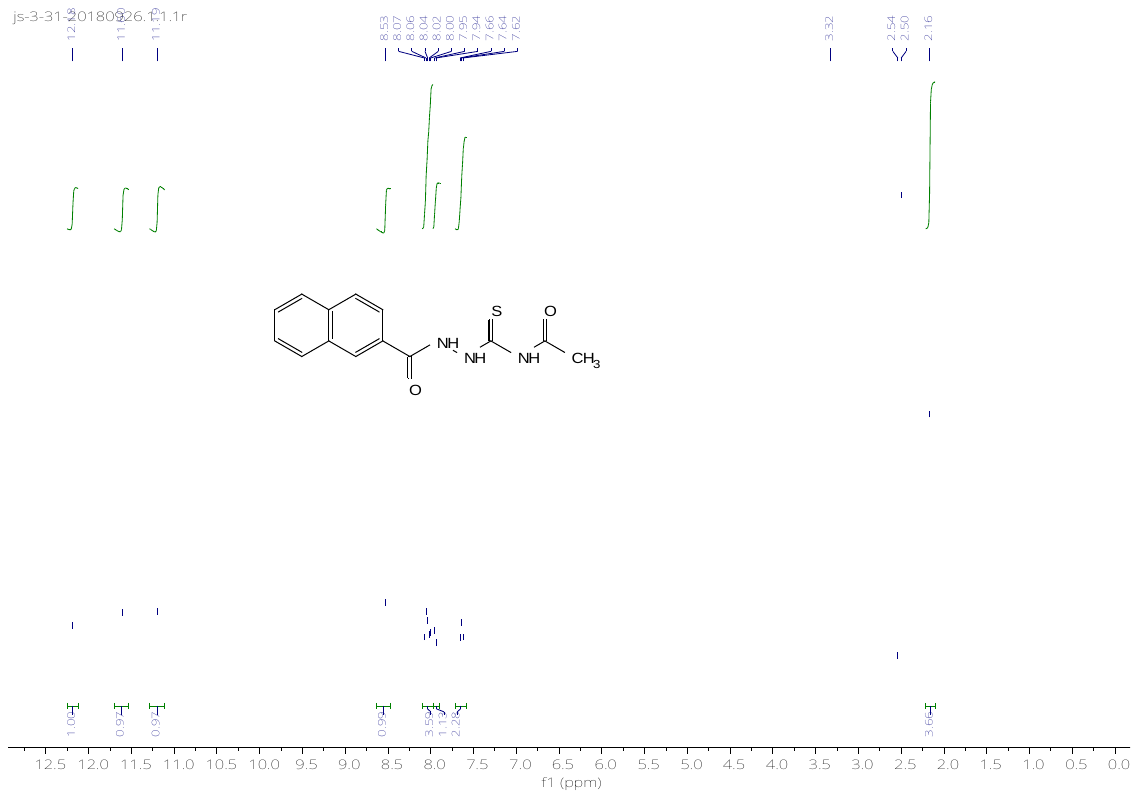

^13^C-NMR spectrum of *N-(2-(2-naphthoyl)hydrazine-1-carbonothioyl)acetamide (****1t****)*

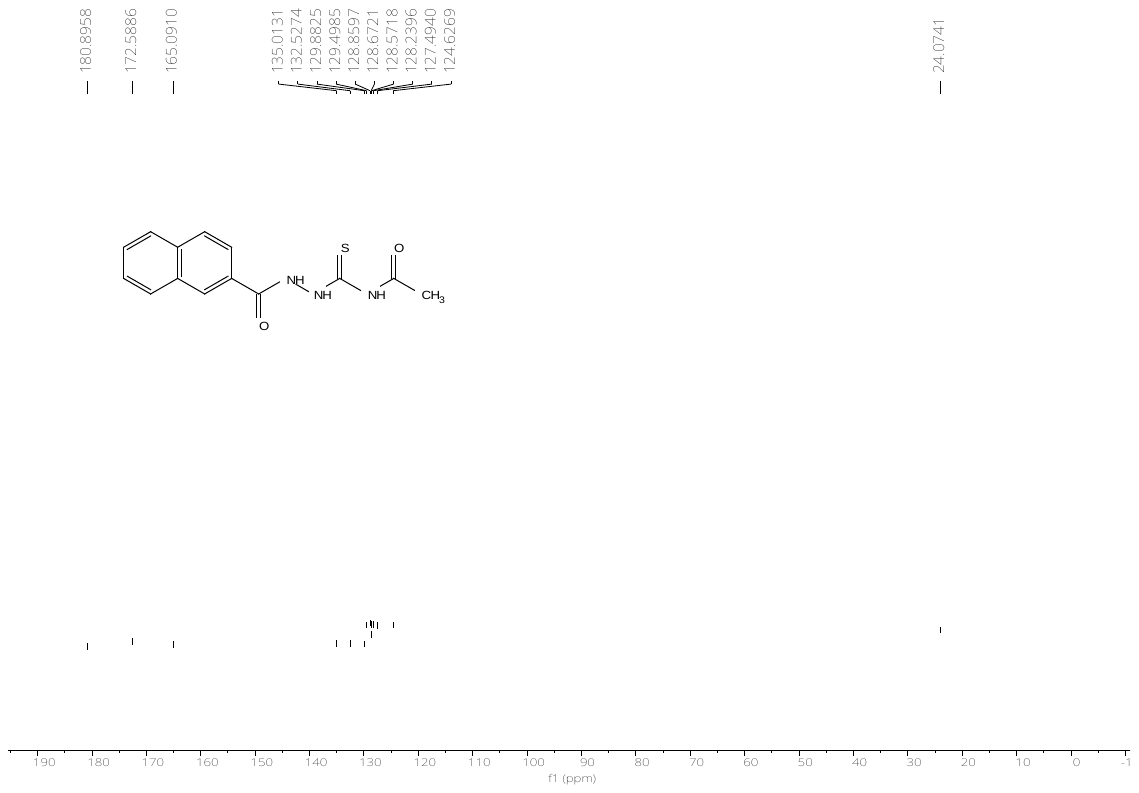

^1^H-NMR spectrum of *N-(2-(4-cyclopropylbenzoyl)hydrazine-1-carbonothioyl)acetamide (****1u****)*

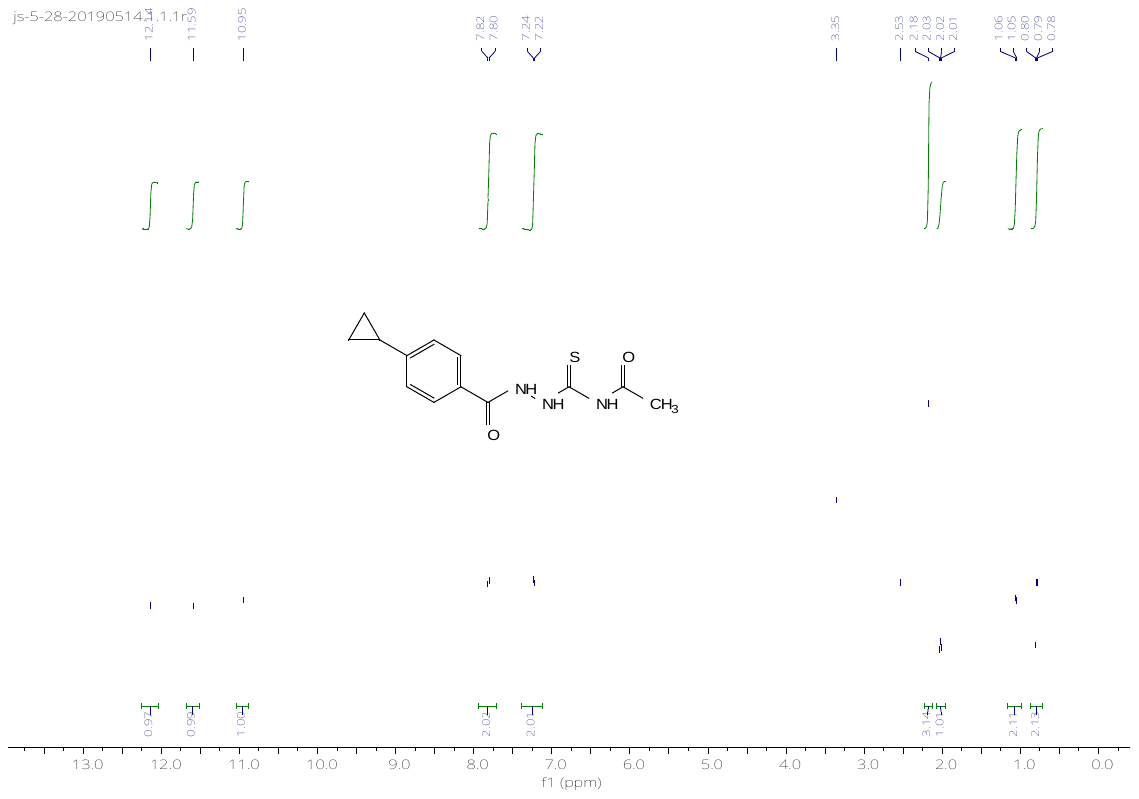

^13^C-NMR spectrum of *N-(2-(4-cyclopropylbenzoyl)hydrazine-1-carbonothioyl)acetamide (****1u****)*

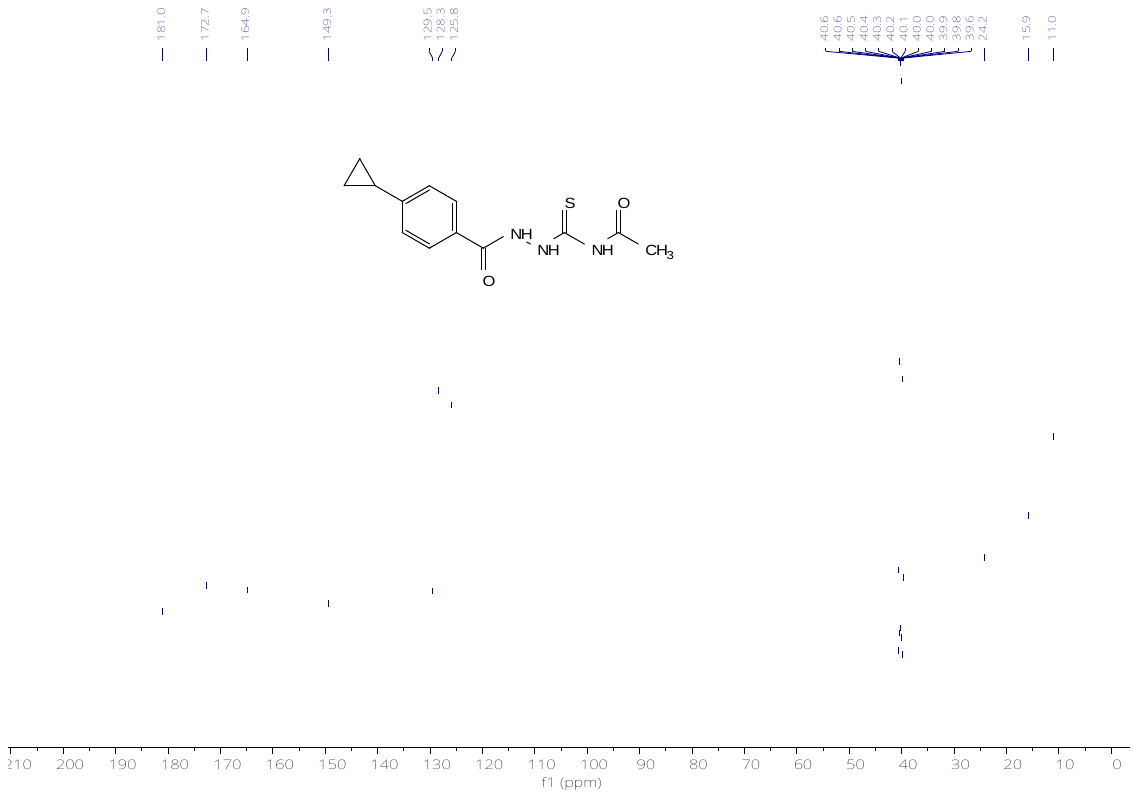

^1^H-NMR spectrum of *N-(2-(4-cyclohexylbenzoyl)hydrazine-1-carbonothioyl)acetamide (****1v****)*

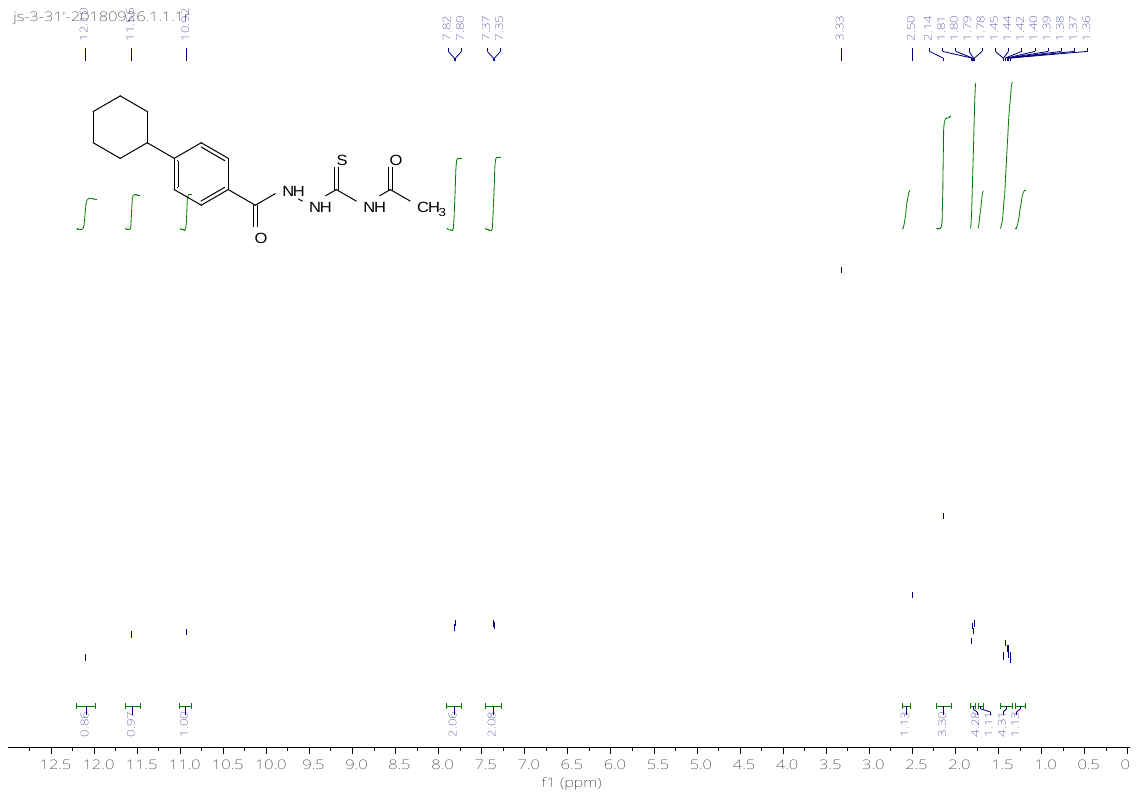

^13^C-NMR spectrum of *N-(2-(4-cyclohexylbenzoyl)hydrazine-1-carbonothioyl)acetamide (****1v****)*

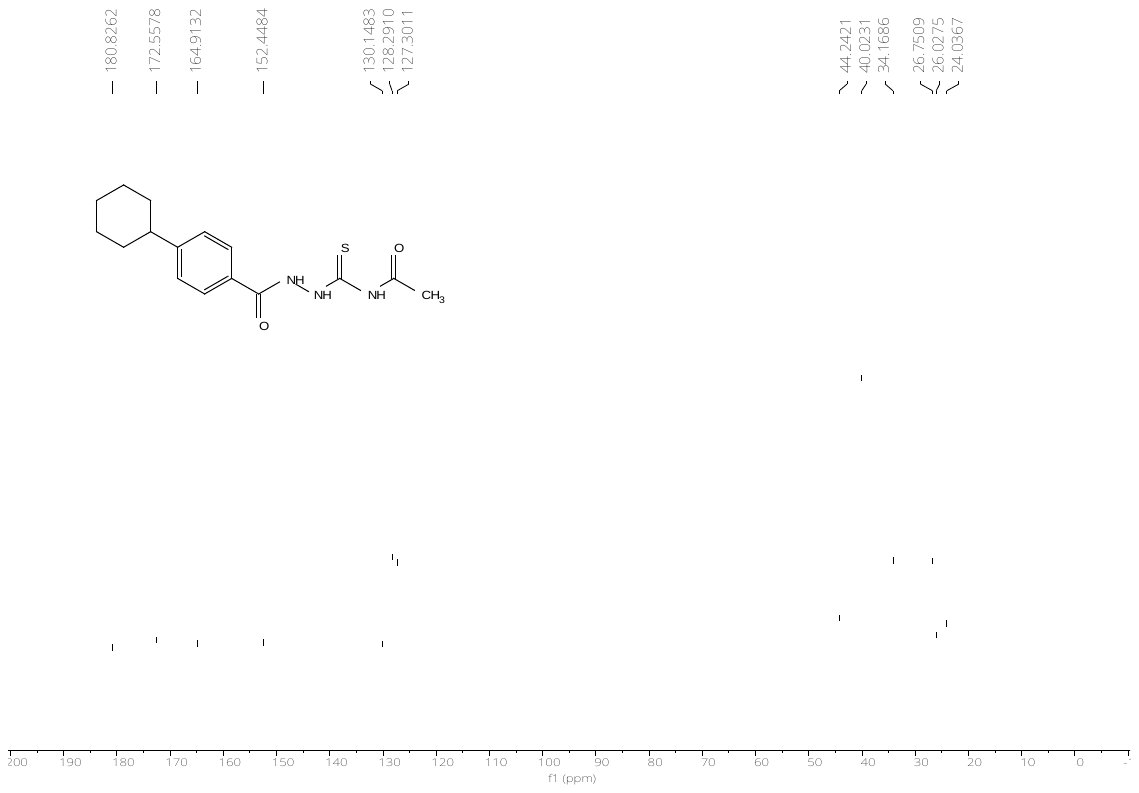

^1^H-NMR spectrum of *N-(2-(4-(4-ethylcyclohexyl)benzoyl)hydrazine-1-carbonothioyl)acetamide (****1w****)*

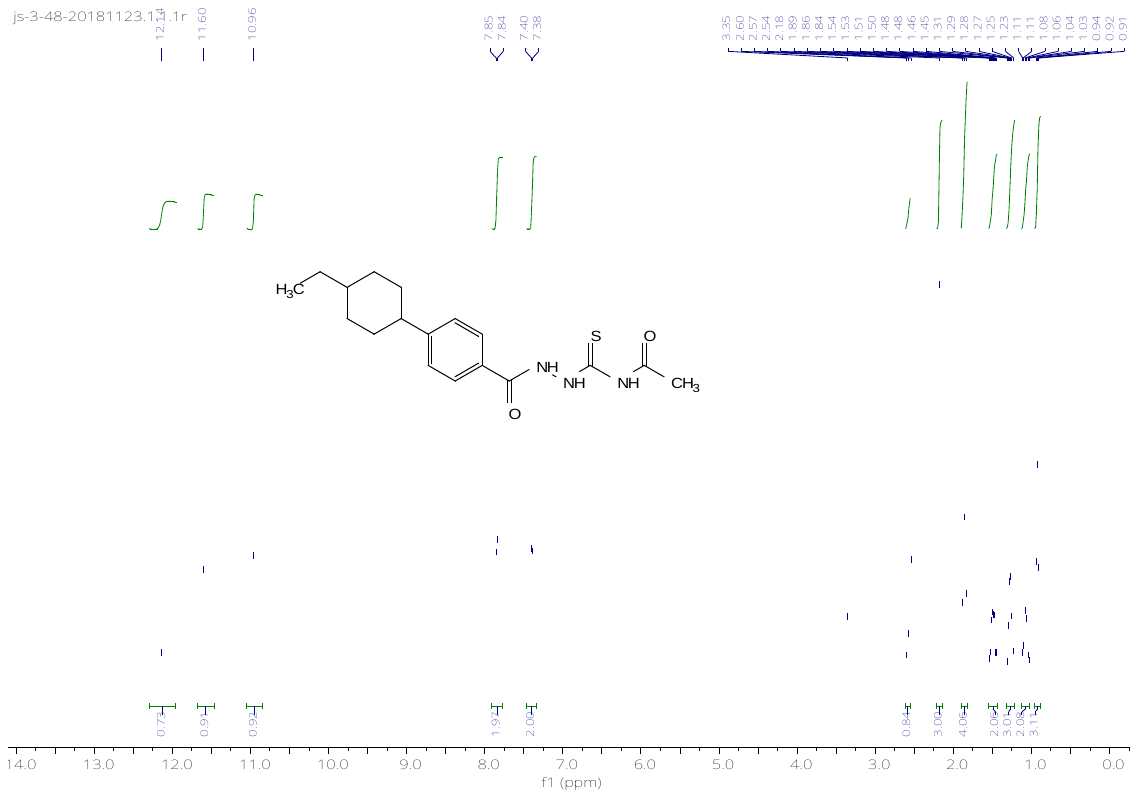

^13^C-NMR spectrum of *N-(2-(4-(4-ethylcyclohexyl)benzoyl)hydrazine-1-carbonothioyl)acetamide (****1w****)*

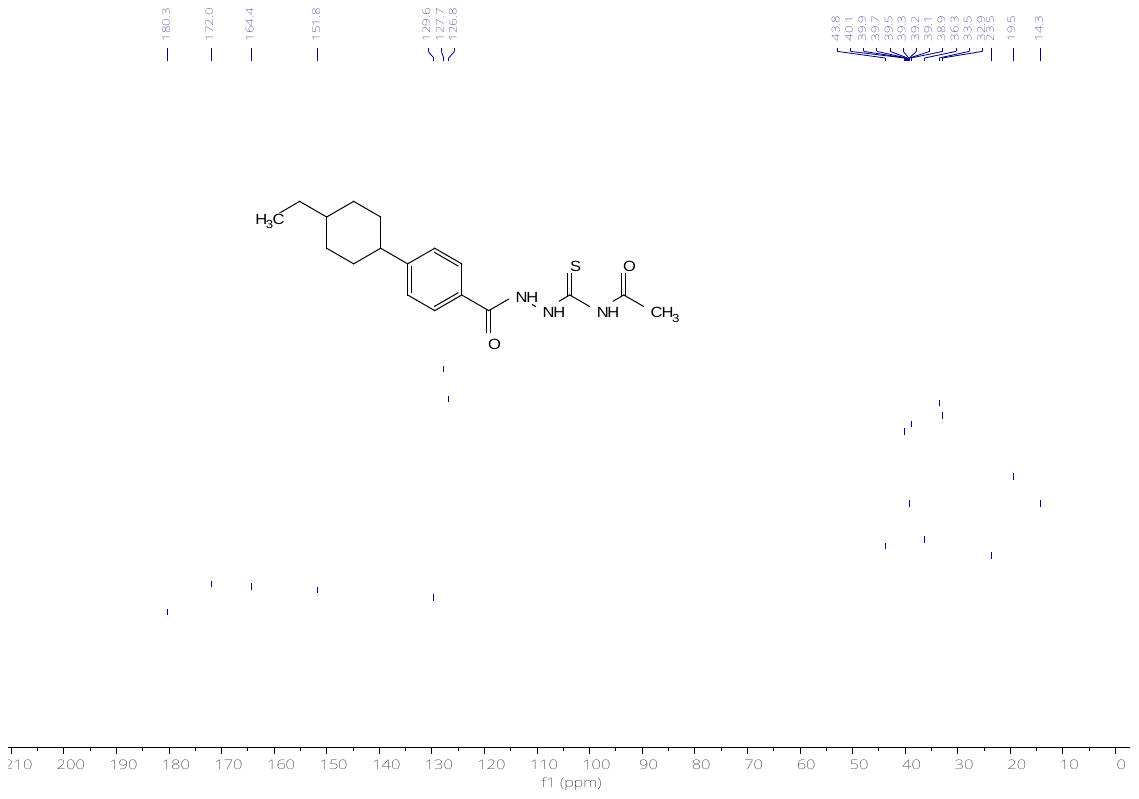

^1^H-NMR spectrum of *N-(2-(4-morpholinobenzoyl)hydrazine-1-carbonothioyl)acetamide (****1x****)*

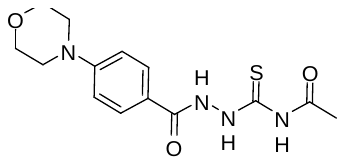

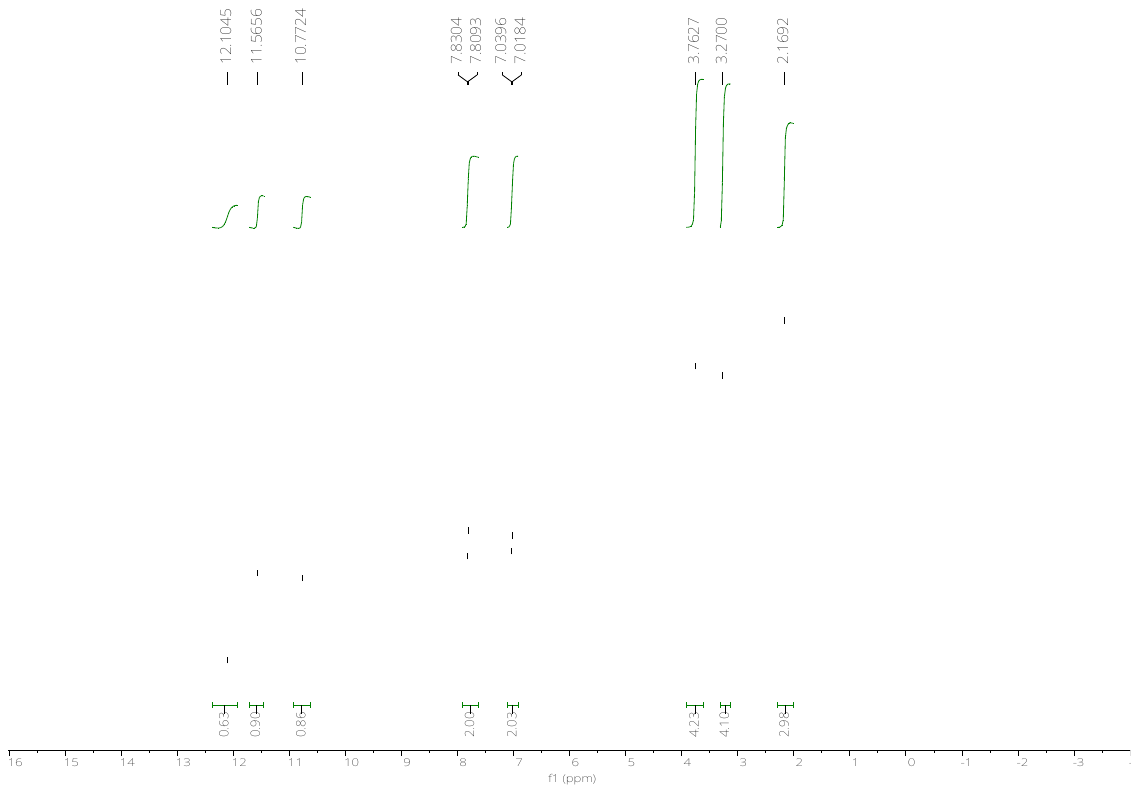

^13^C-NMR spectrum of *N-(2-(4-morpholinobenzoyl)hydrazine-1-carbonothioyl)acetamide (****1x****)*

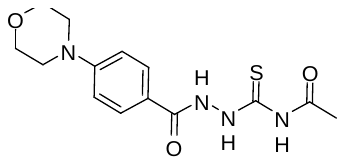

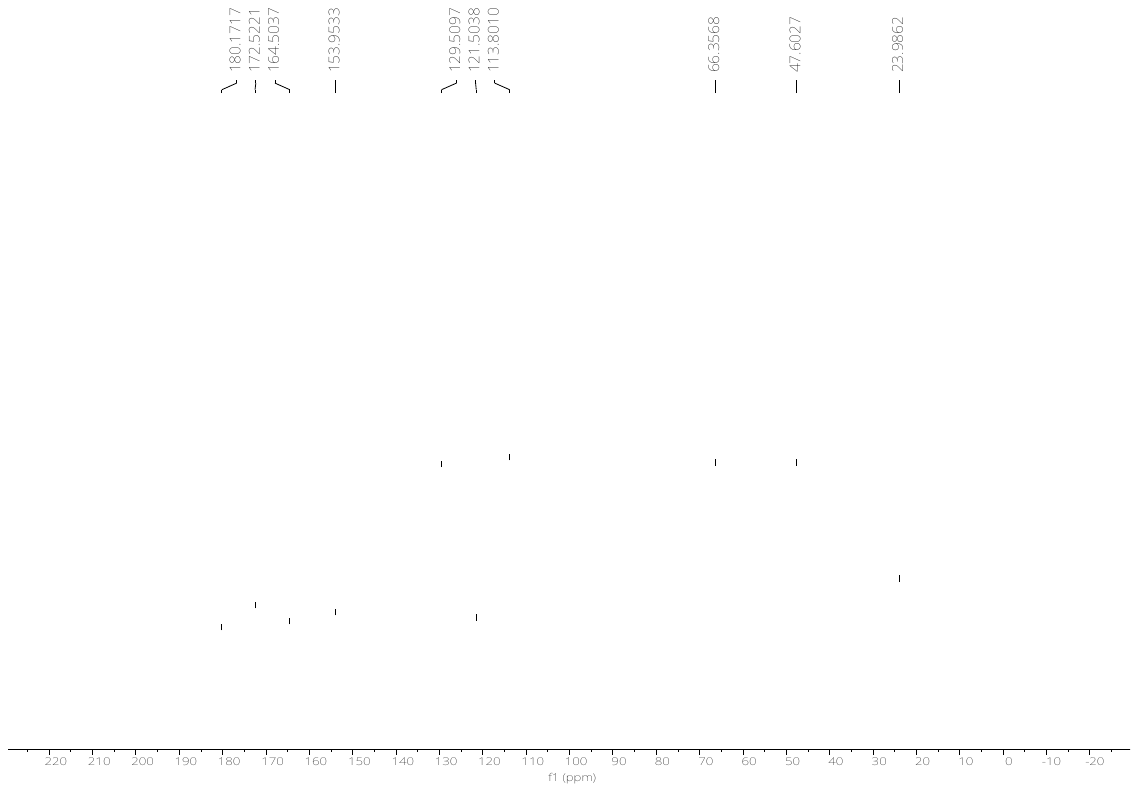

^1^H-NMR spectrum of *N-(2-(4-(n-pentoxy)benzoyl)hydrazine-1-carbonothioyl)propionamide (****2a****)*

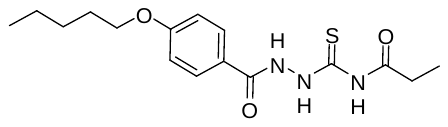

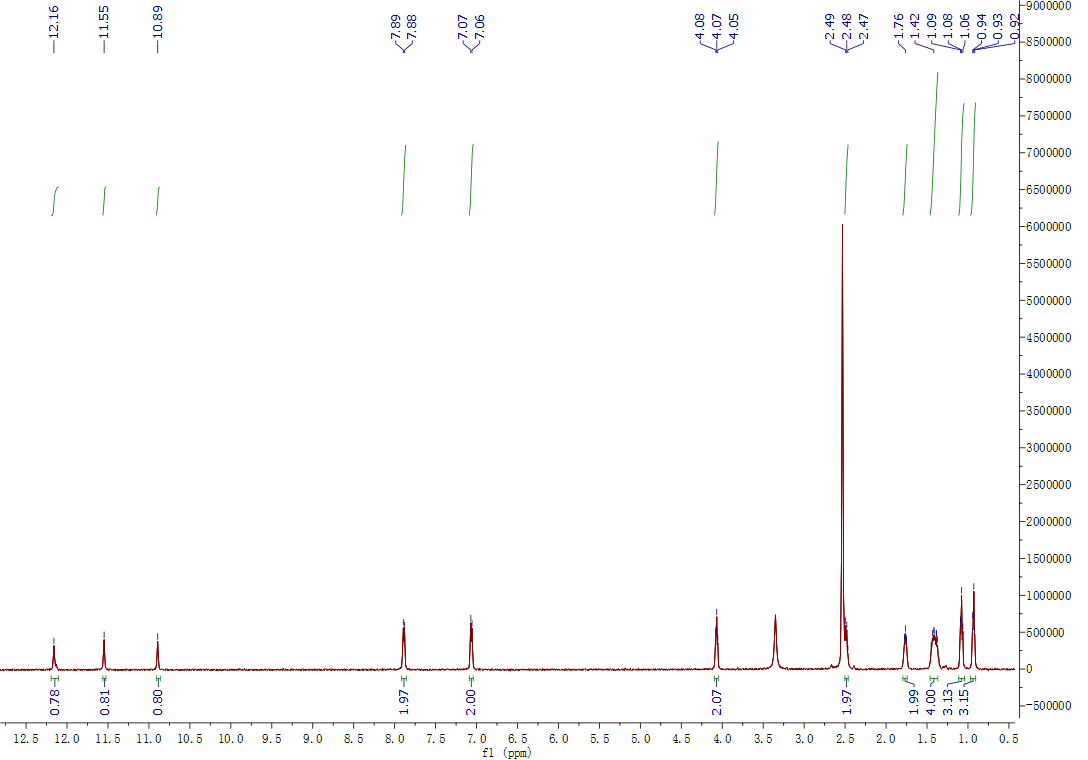

^13^C-NMR spectrum of *N-(2-(4-(n-pentoxy)benzoyl)hydrazine-1-carbonothioyl)propionamide (****2a****)*

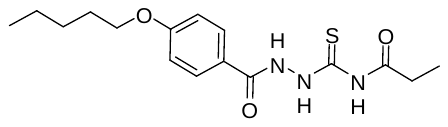

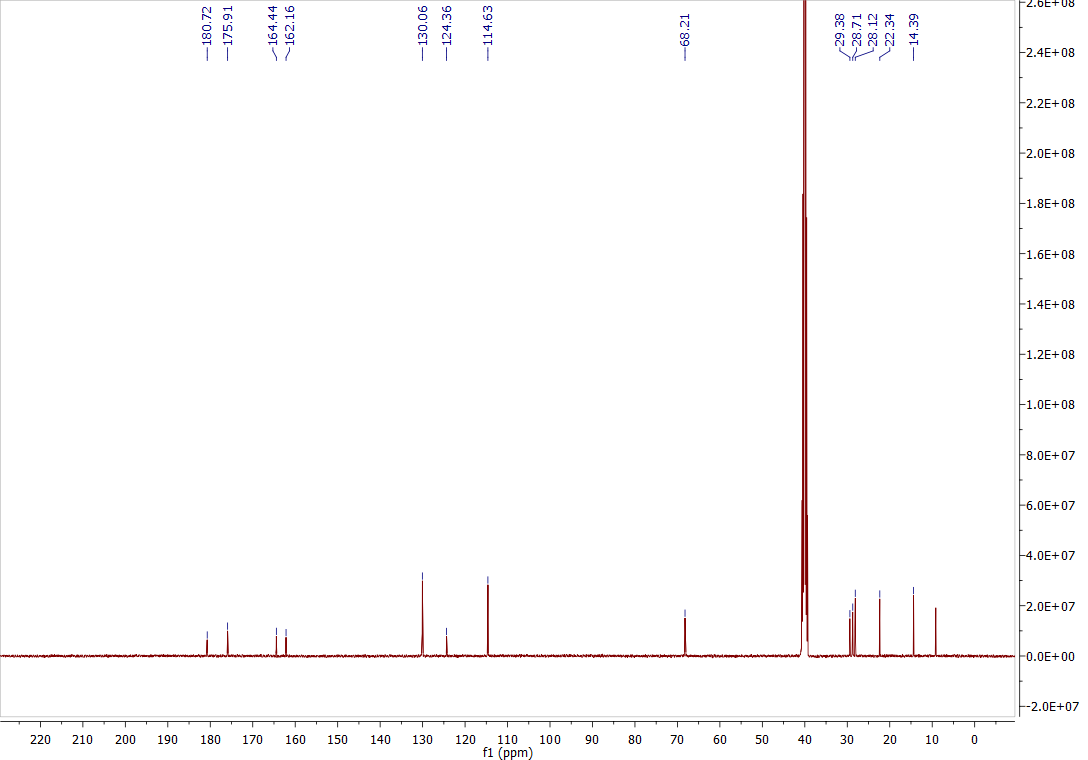

^1^H-NMR spectrum of *N-(2-(4-(n-pentoxy)benzoyl)hydrazine-1-carbonothioyl)butyramide (****2b****)*

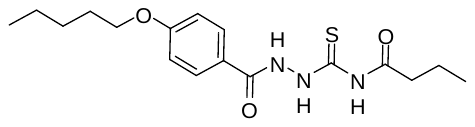

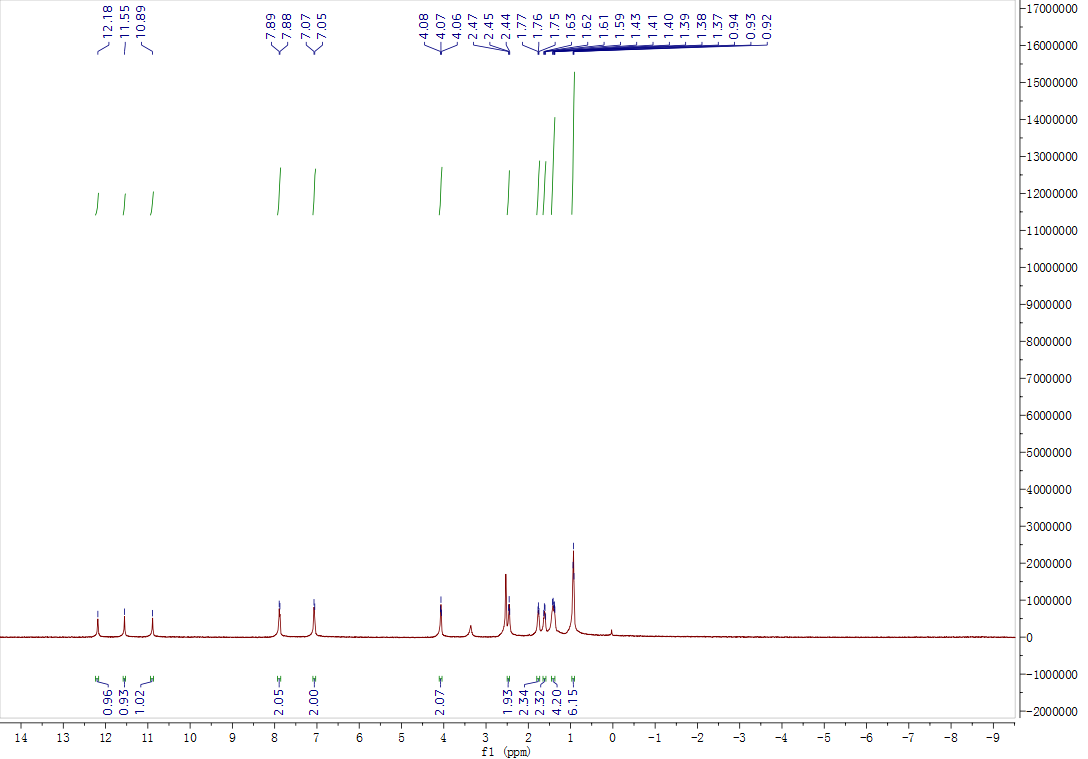

^13^C-NMR spectrum of *N-(2-(4-(n-pentoxy)benzoyl)hydrazine-1-carbonothioyl)butyramide (****2b****)*

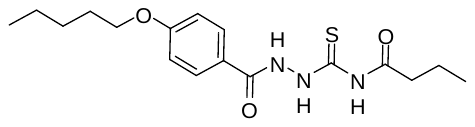

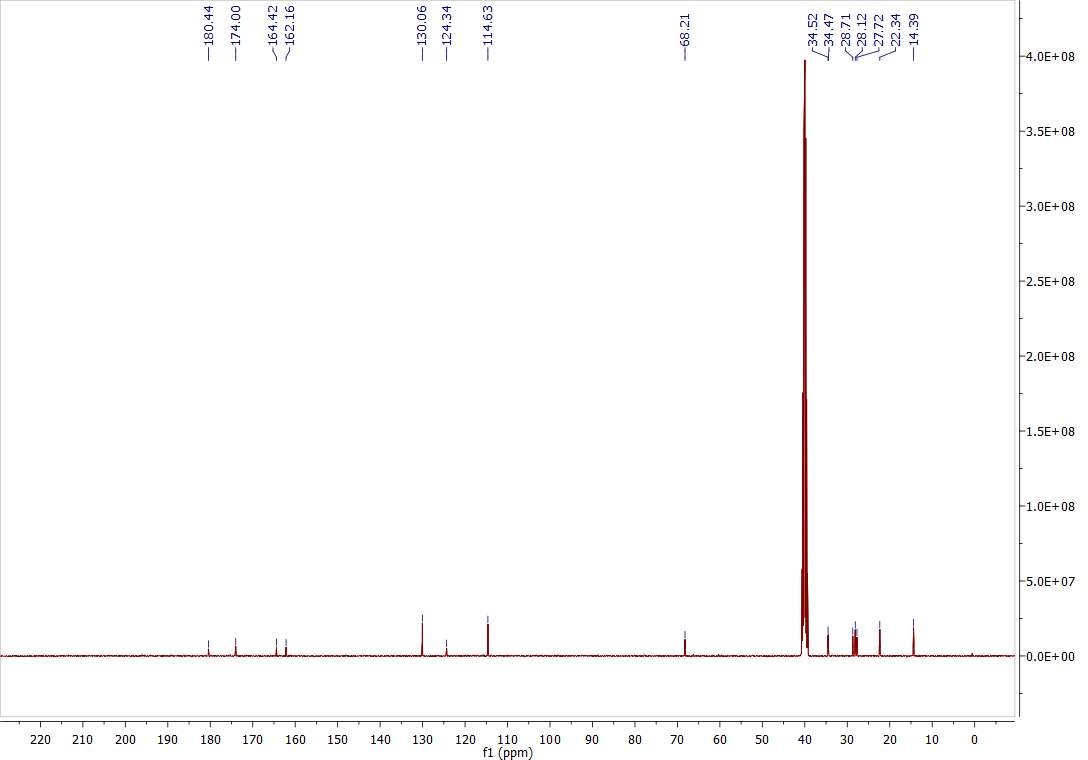

^1^H-NMR spectrum of *N-(2-(4-(n-pentoxy)benzoyl)hydrazine-1-carbonothioyl)isobutyramide (****2c****)*

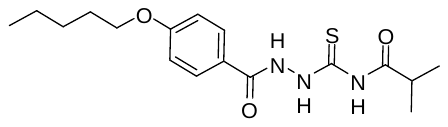

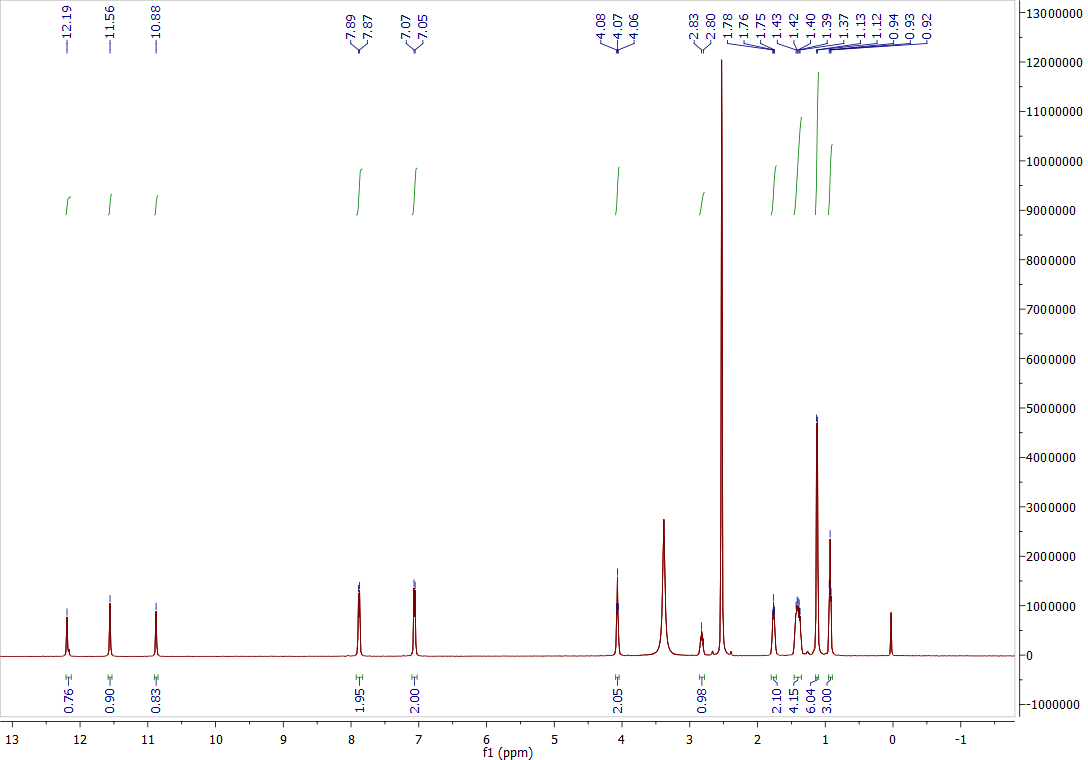

^13^C-NMR spectrum of *N-(2-(4-(n-pentoxy)benzoyl)hydrazine-1-carbonothioyl)isobutyramide (****2c****)*

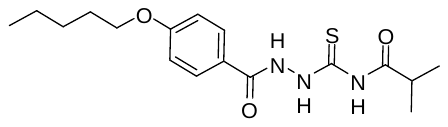

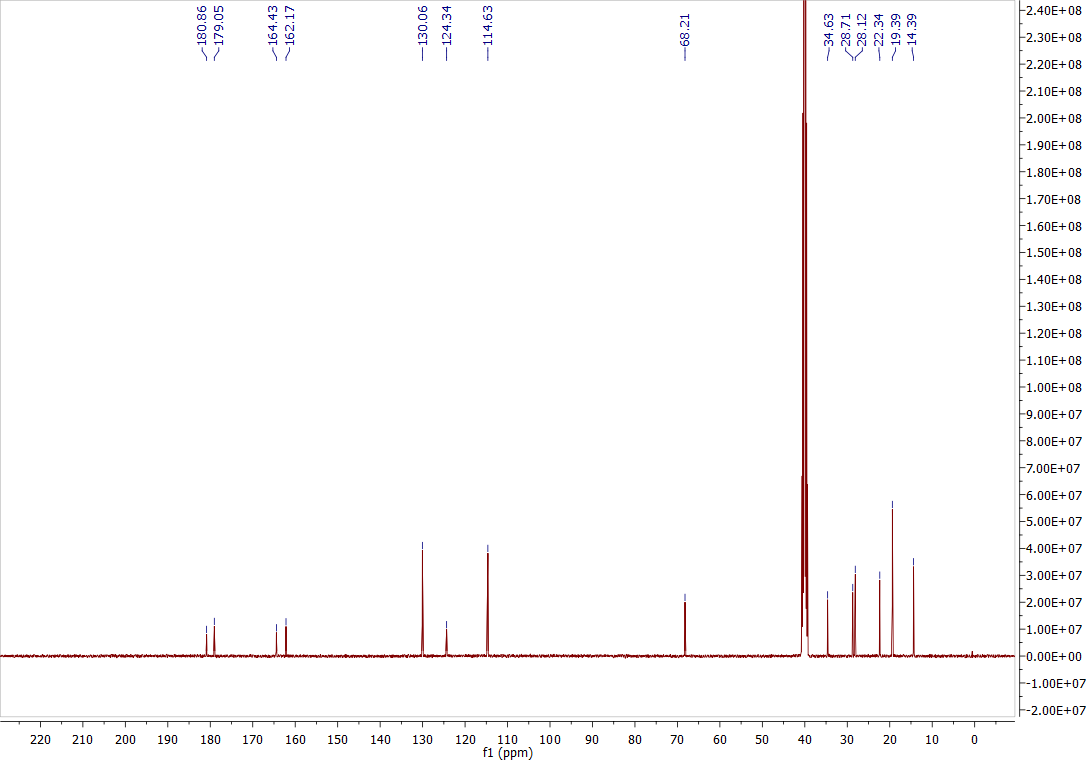

^1^H-NMR spectrum of *N-(2-(4-(n-pentoxy)benzoyl)hydrazine-1-carbonothioyl)cyclopropanecarboxamide (****2d****)*

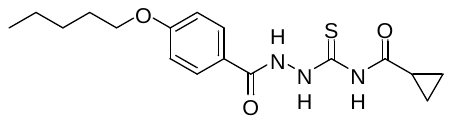

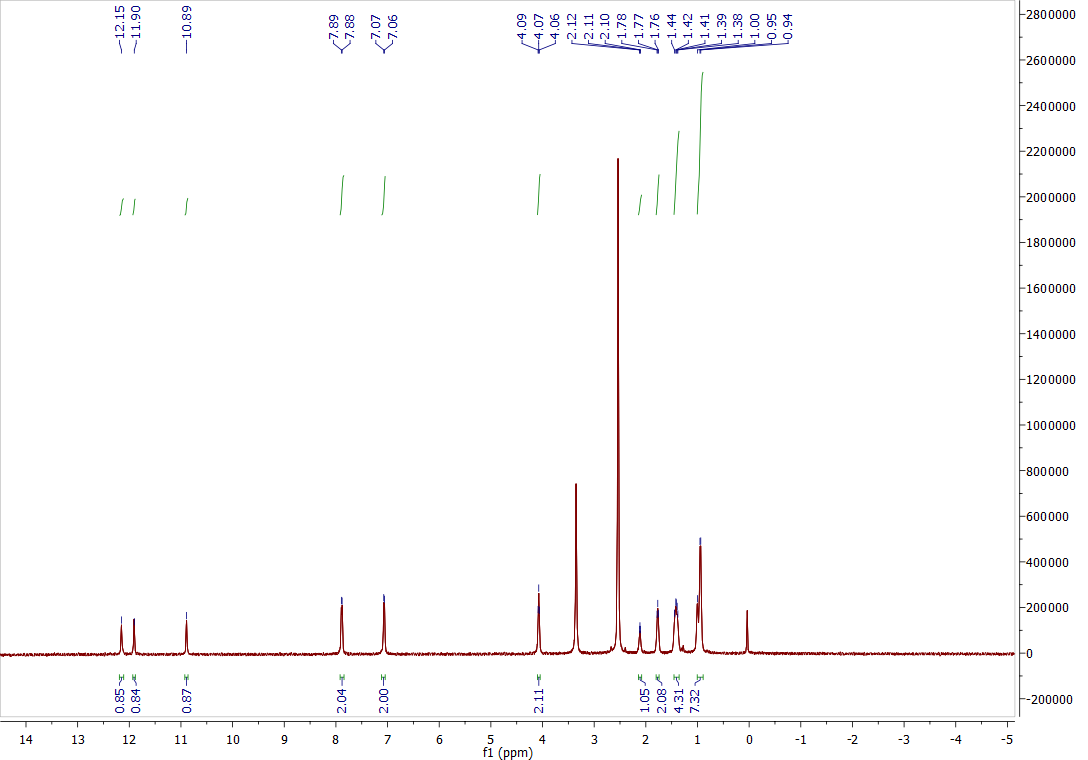

^13^C-NMR spectrum of *N-(2-(4-(n-pentoxy)benzoyl)hydrazine-1-carbonothioyl)cyclopropanecarboxamide (****2d****)*

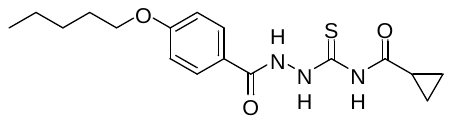

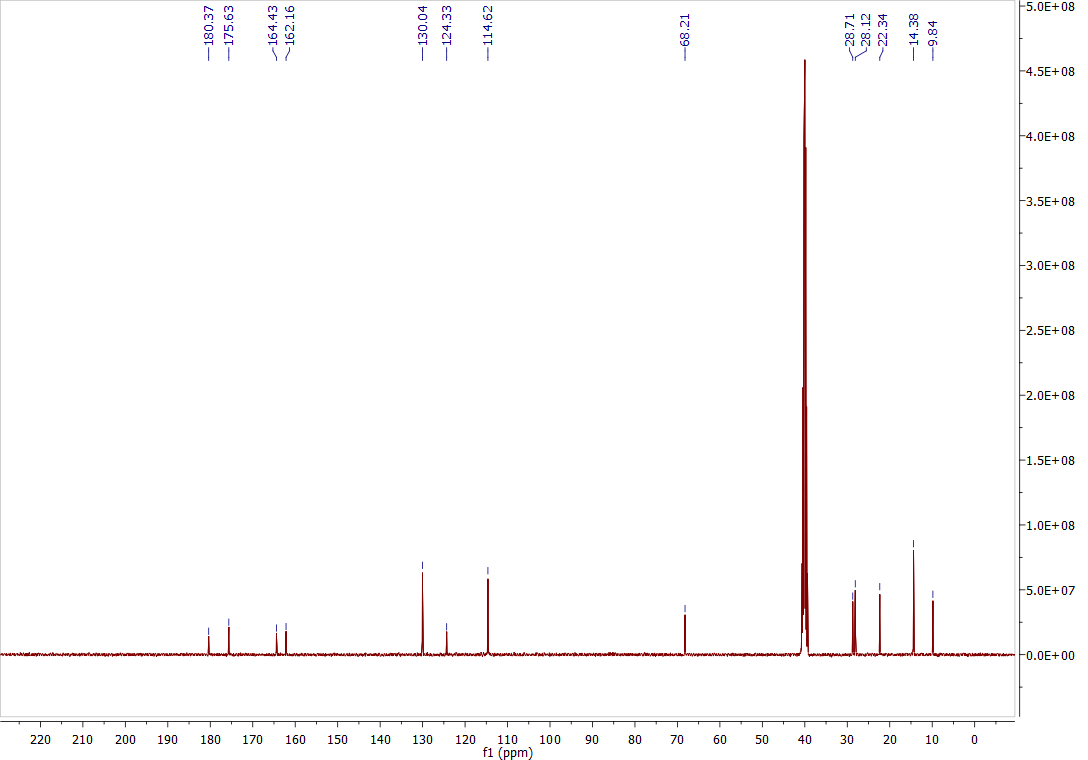

^1^H-NMR spectrum of *N-(2-(4-(n-pentoxy)benzoyl)hydrazine-1-carbonothioyl)cyclopentanecarboxamide (****2e****)*

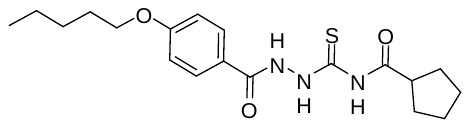

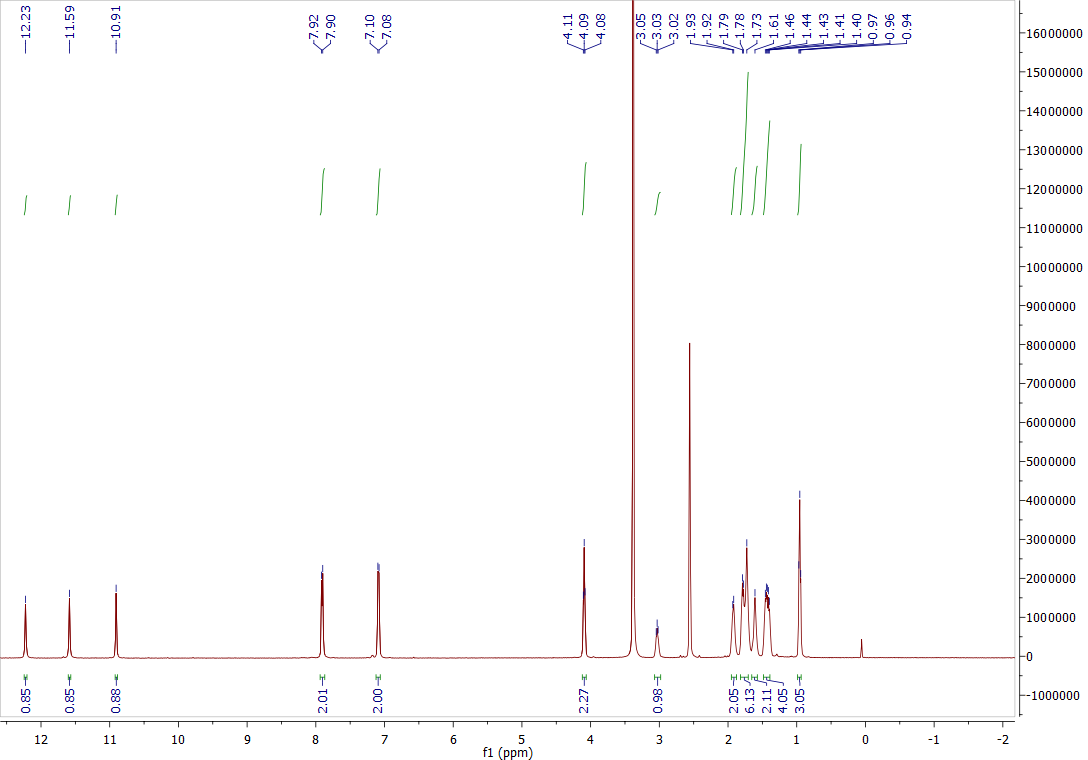

^13^C-NMR spectrum of *N-(2-(4-(n-pentoxy)benzoyl)hydrazine-1-carbonothioyl)cyclopentanecarboxamide (****2e****)*

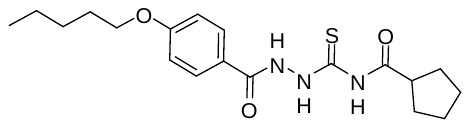

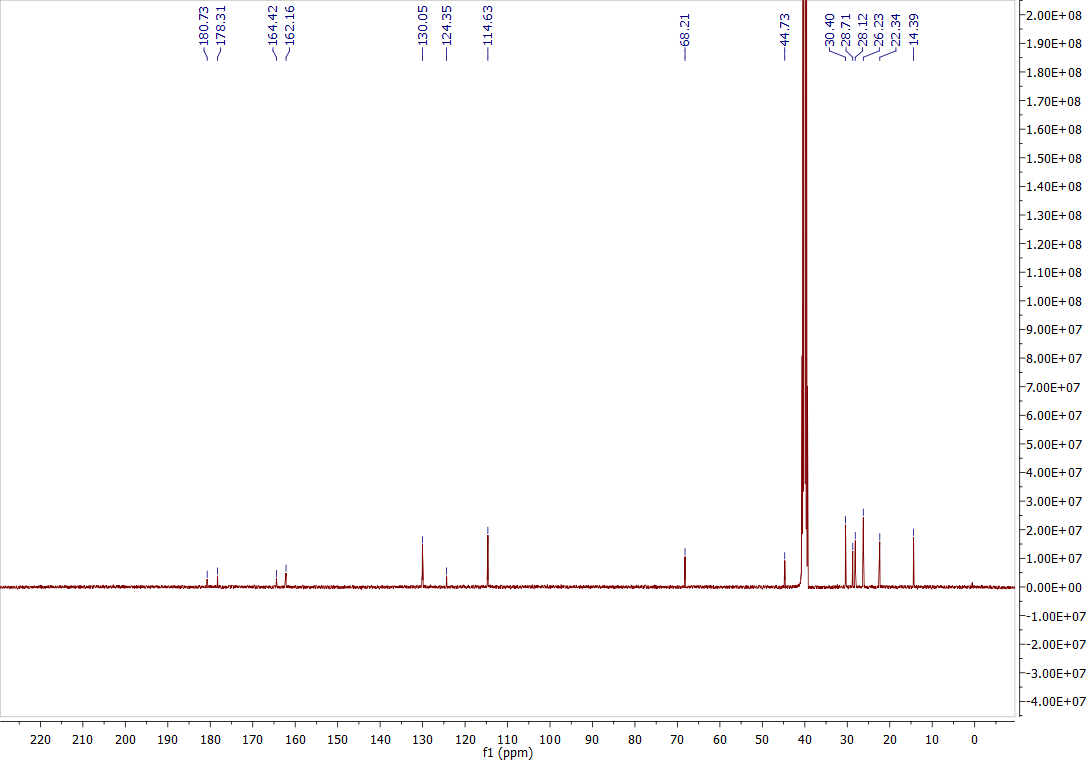

^1^H-NMR spectrum of *N-(2-(4-(n-pentoxy)benzoyl)hydrazine-1-carbonothioyl)-3-phenylpropanamide (****2f****)*

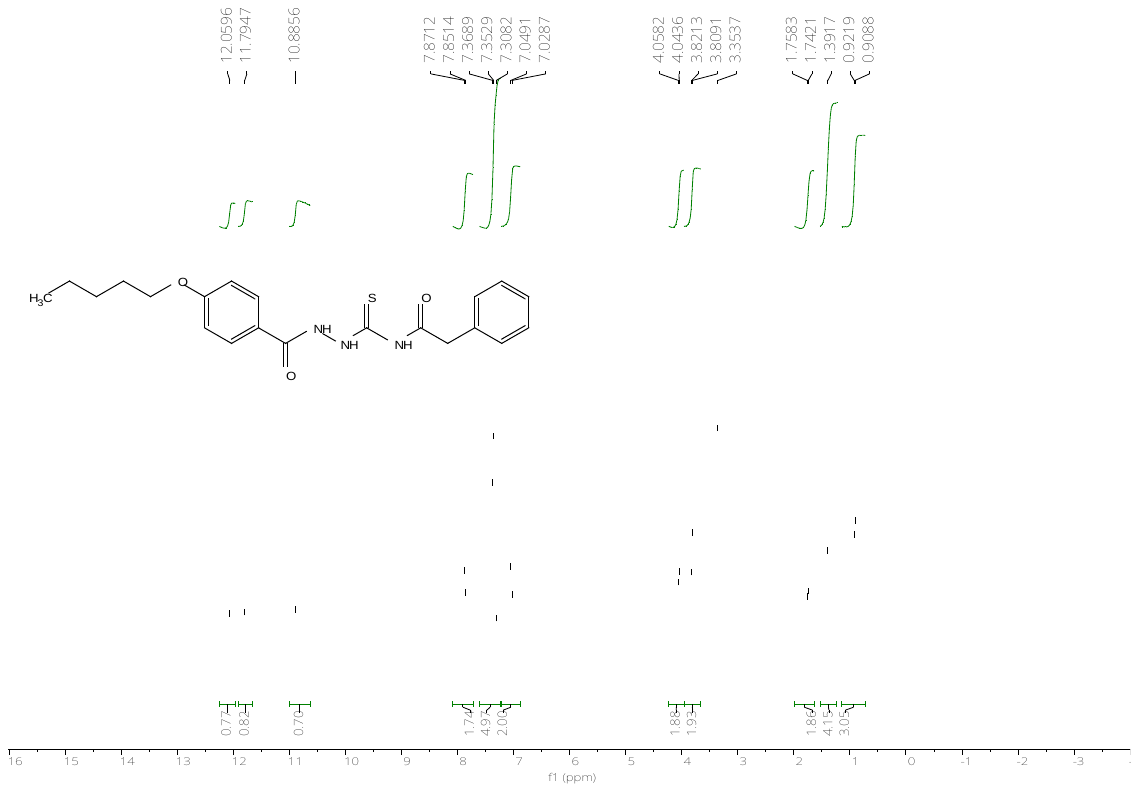

^13^C-NMR spectrum of *N-(2-(4-(n-pentoxy)benzoyl)hydrazine-1-carbonothioyl)-3-phenylpropanamide (****2f****)*

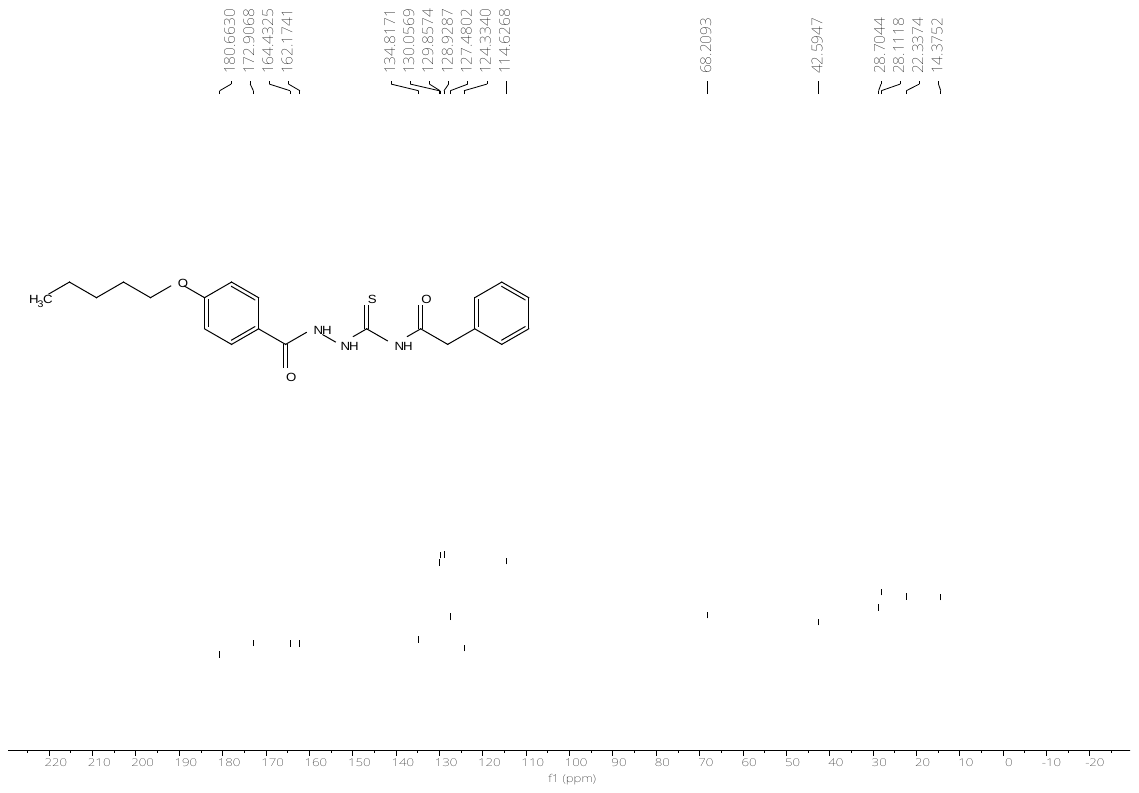

^1^H-NMR spectrum of *N-(2-(4-(n-pentoxy)benzoyl)hydrazine-1-carbonothioyl)thiophene-2-carboxamide (****2g****)*

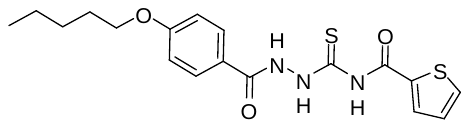

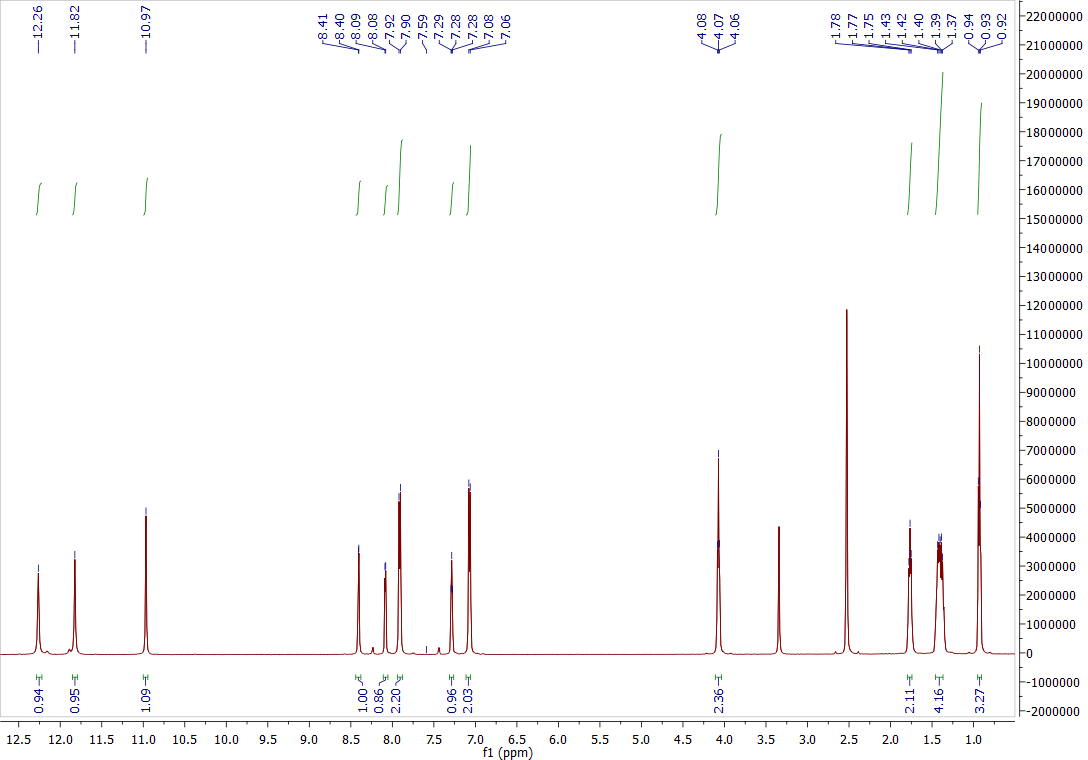

^13^C-NMR spectrum of *N-(2-(4-(n-pentoxy)benzoyl)hydrazine-1-carbonothioyl)thiophene-2-carboxamide (****2g****)*

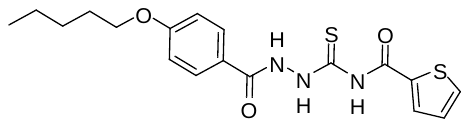

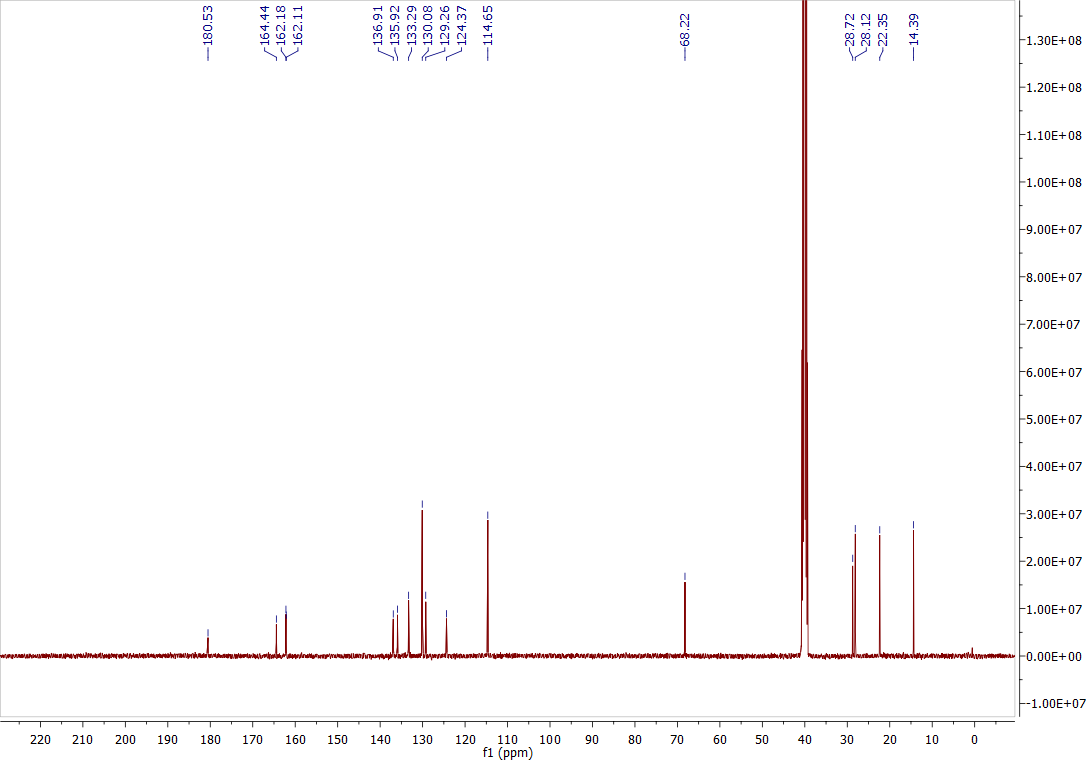

^1^H-NMR spectrum of *methyl 4-((2-(4-(n-pentoxy)benzoyl)hydrazine-1-carbonothioyl)carbamoyl)benzoate* *(****2h****)*

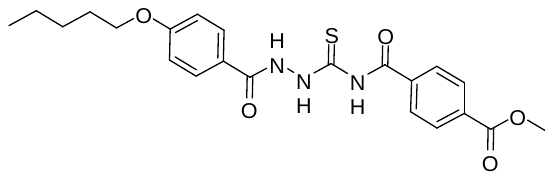

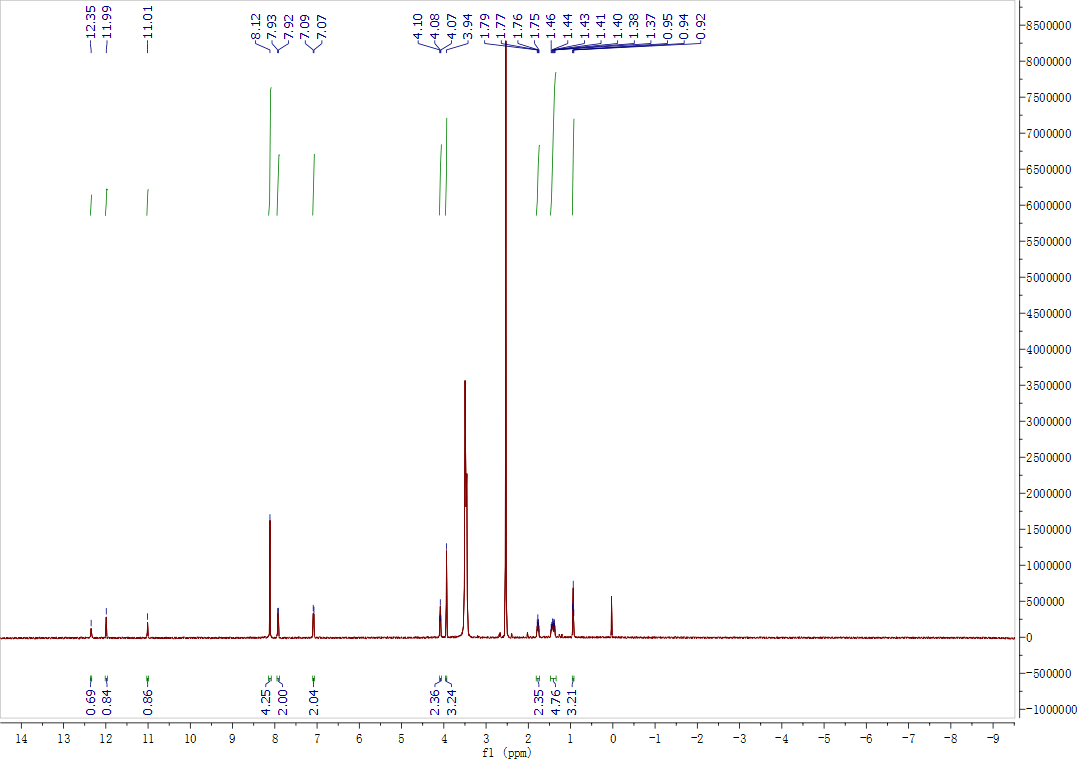

^13^C-NMR spectrum of *methyl 4-((2-(4-(n-pentoxy)benzoyl)hydrazine-1-carbonothioyl)carbamoyl)benzoate* *(****2h****)*

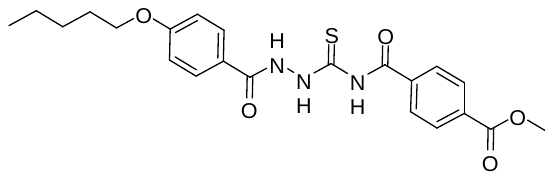

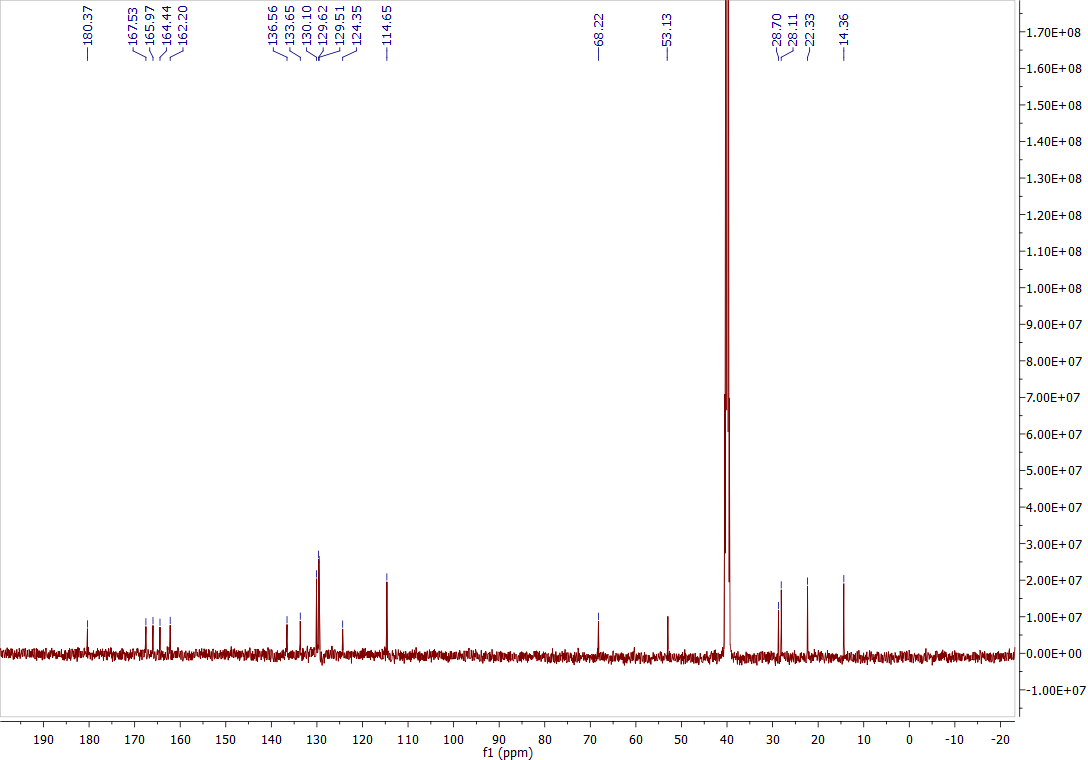

^1^H-NMR spectrum of *methyl 4-oxo-4-(2-(4-(n-pentoxy)benzoyl)hydrazine-1-carbothioamido)butanoate (****2i****)*

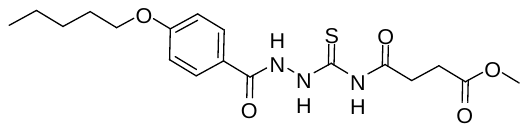

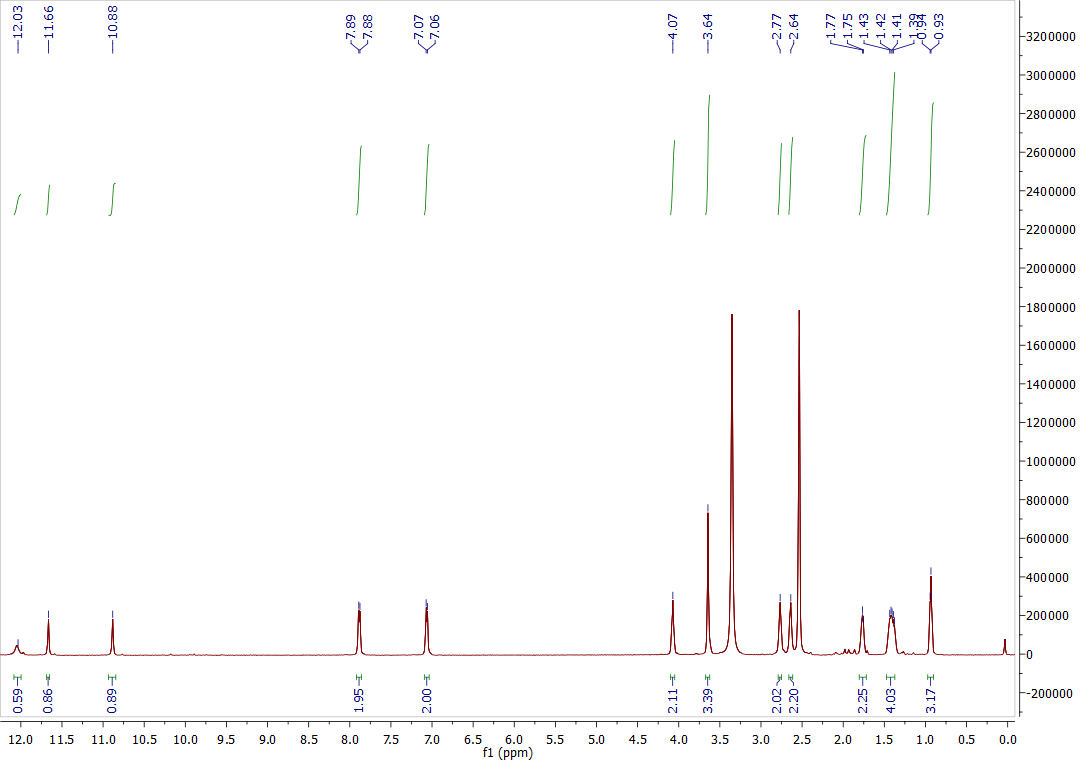

^13^C-NMR spectrum of *methyl 4-oxo-4-(2-(4-(n-pentoxy)benzoyl)hydrazine-1-carbothioamido)butanoate (****2i****)*

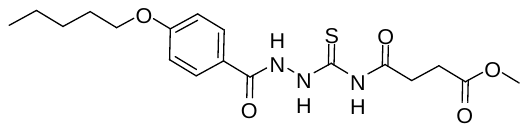

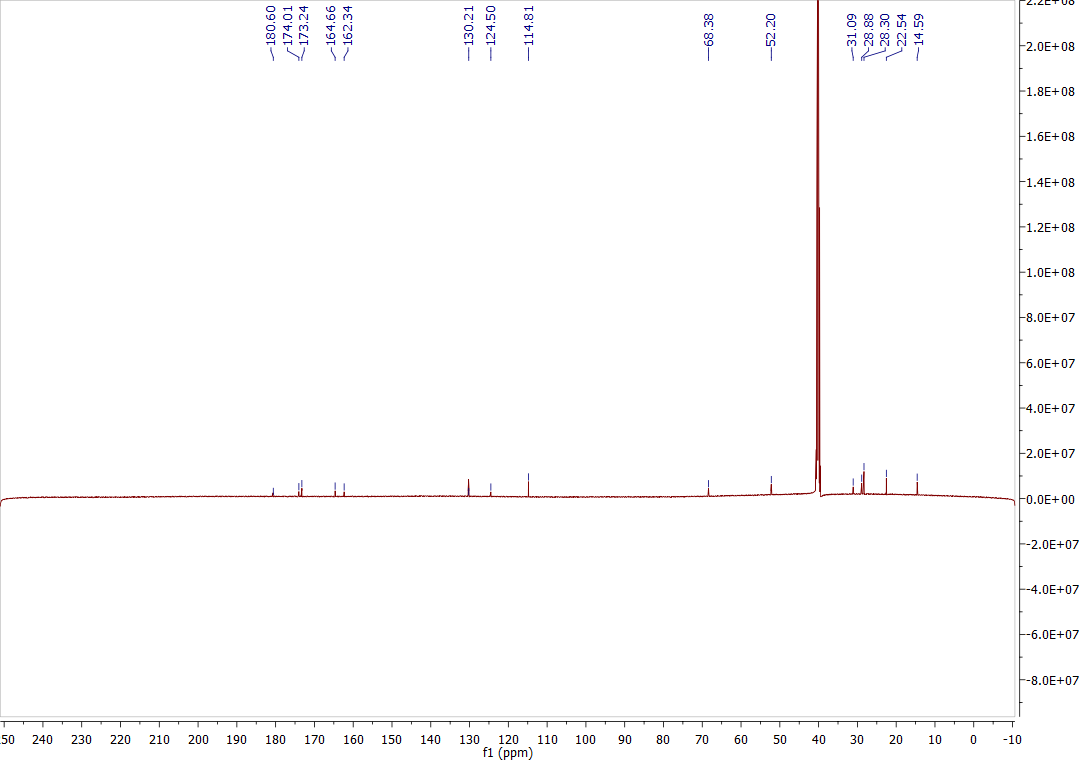

^1^H-NMR spectrum of *4-((2-(4-(n-pentoxy)benzoyl)hydrazine-1-carbonothioyl)carbamoyl)benzoic acid (****2j****)*

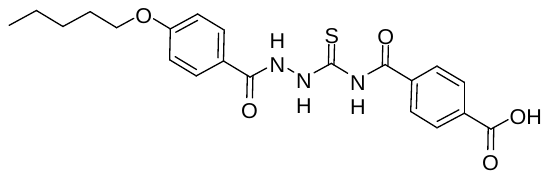

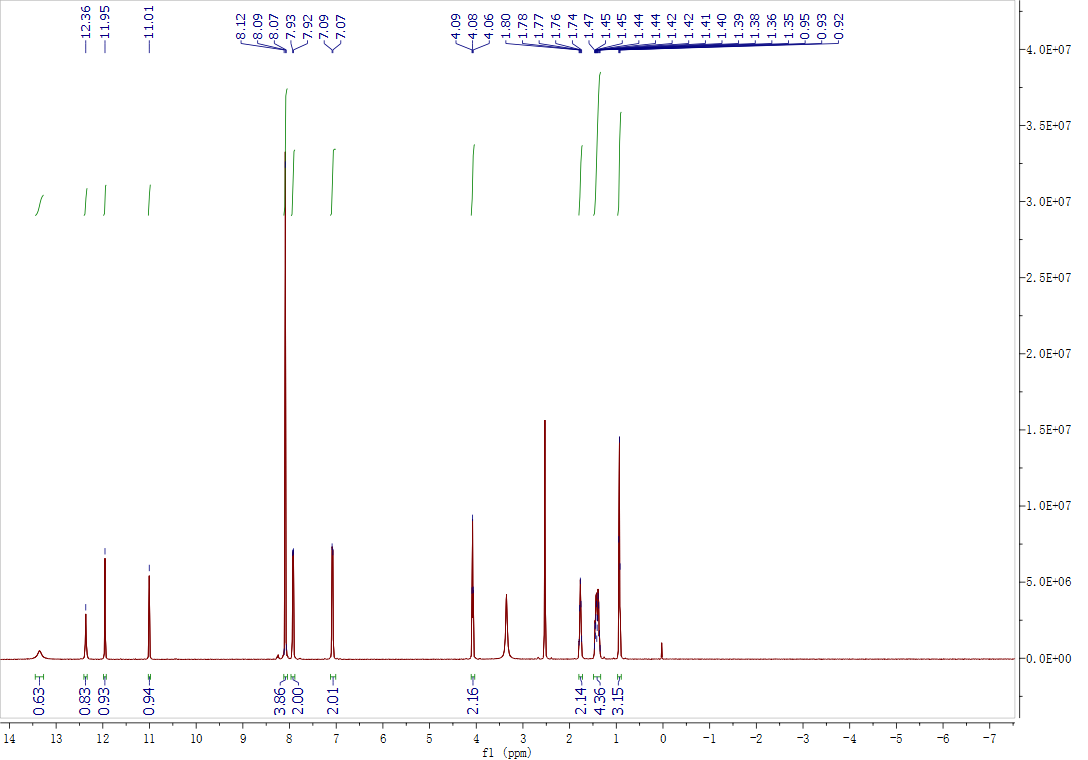

^13^C-NMR spectrum of *4-((2-(4-(n-pentoxy)benzoyl)hydrazine-1-carbonothioyl)carbamoyl)benzoic acid (****2j****)*

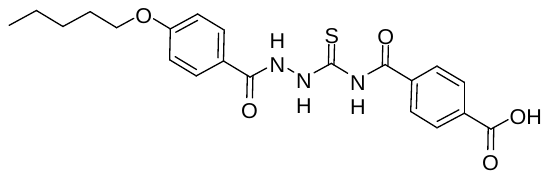

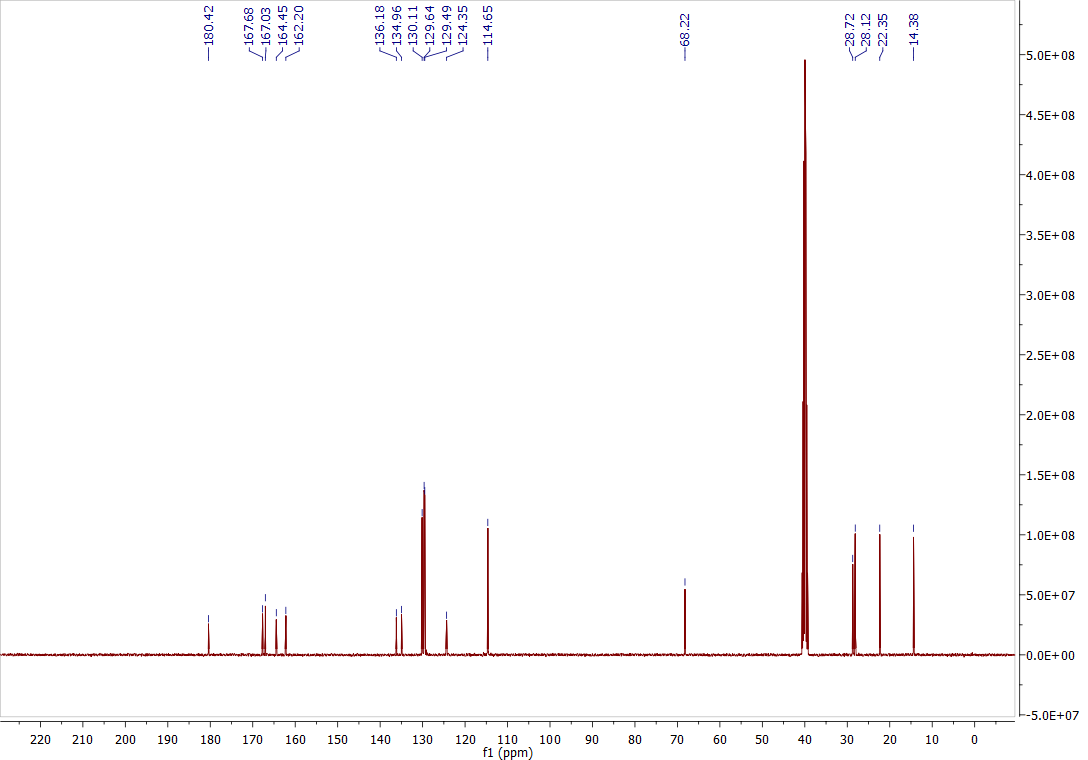

^1^H-NMR spectrum of *4-oxo-4-(2-(4-(n-pentoxy)benzoyl)hydrazine-1-carbothioamido)butanoic acid (****2k****)*

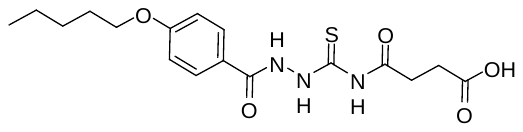

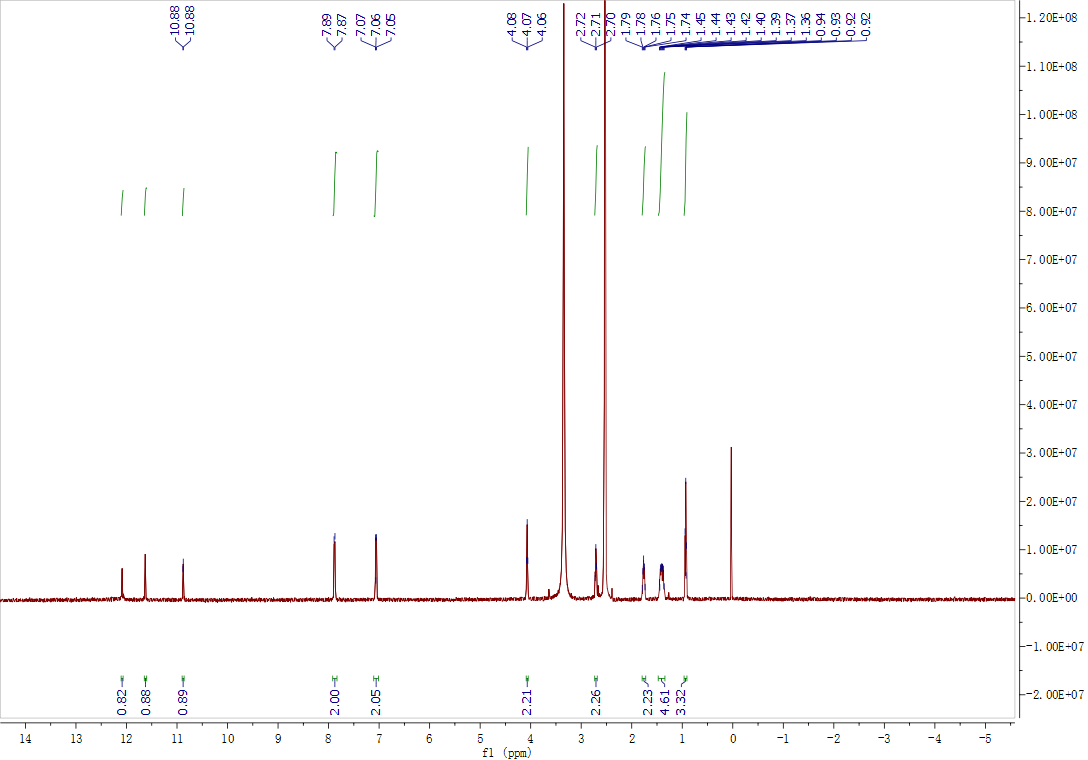

^13^C-NMR spectrum of *4-oxo-4-(2-(4-(n-pentoxy)benzoyl)hydrazine-1-carbothioamido)butanoic acid (****2k****)*

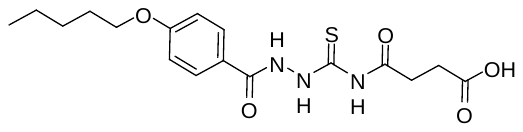

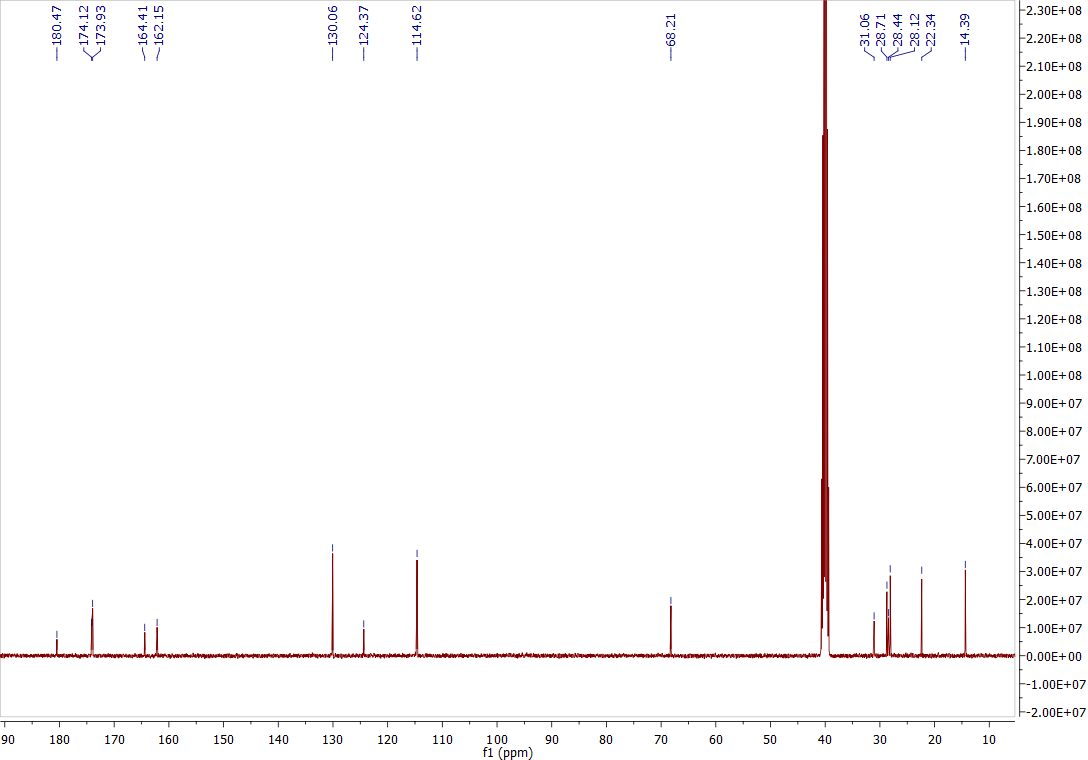

^1^H-NMR spectrum of *N-(5-(4-(n-hexoxy)phenyl)-1,3,4-oxadiazol-2-yl)acetamide (****3a****)*

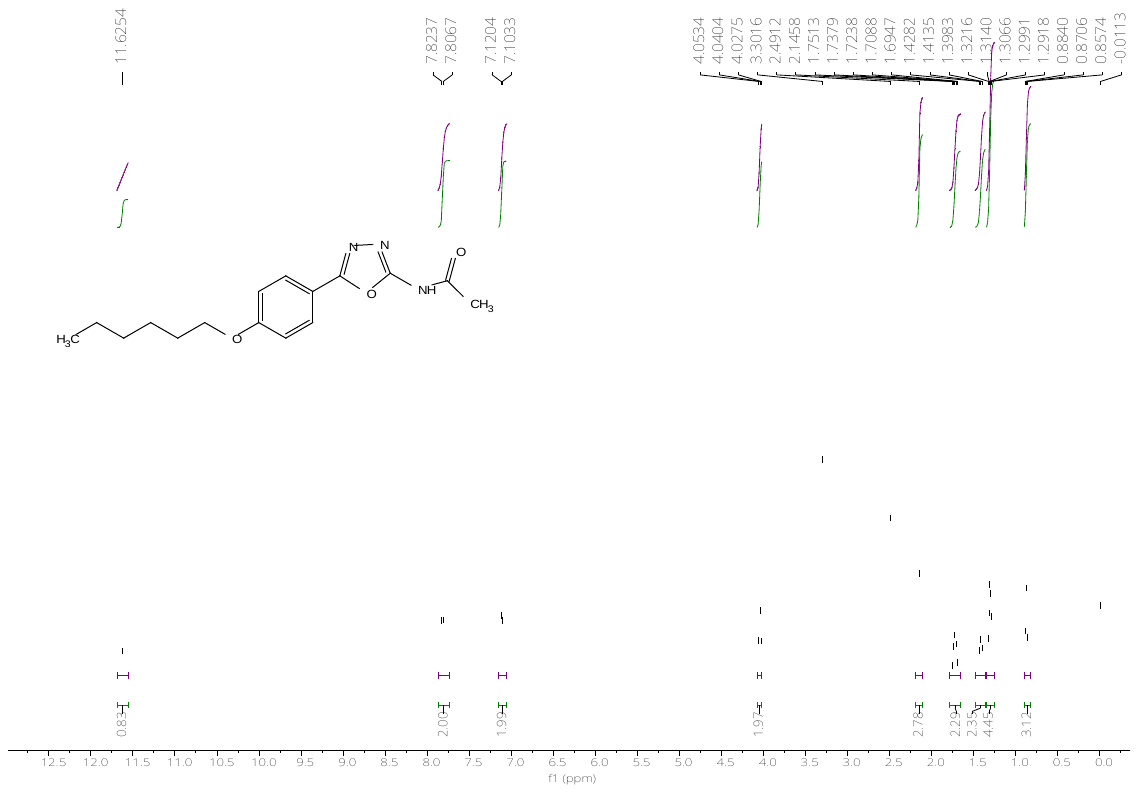

^13^C-NMR spectrum of *N-(5-(4-(n-hexoxy)phenyl)-1,3,4-oxadiazol-2-yl)acetamide* *(****3a****)*

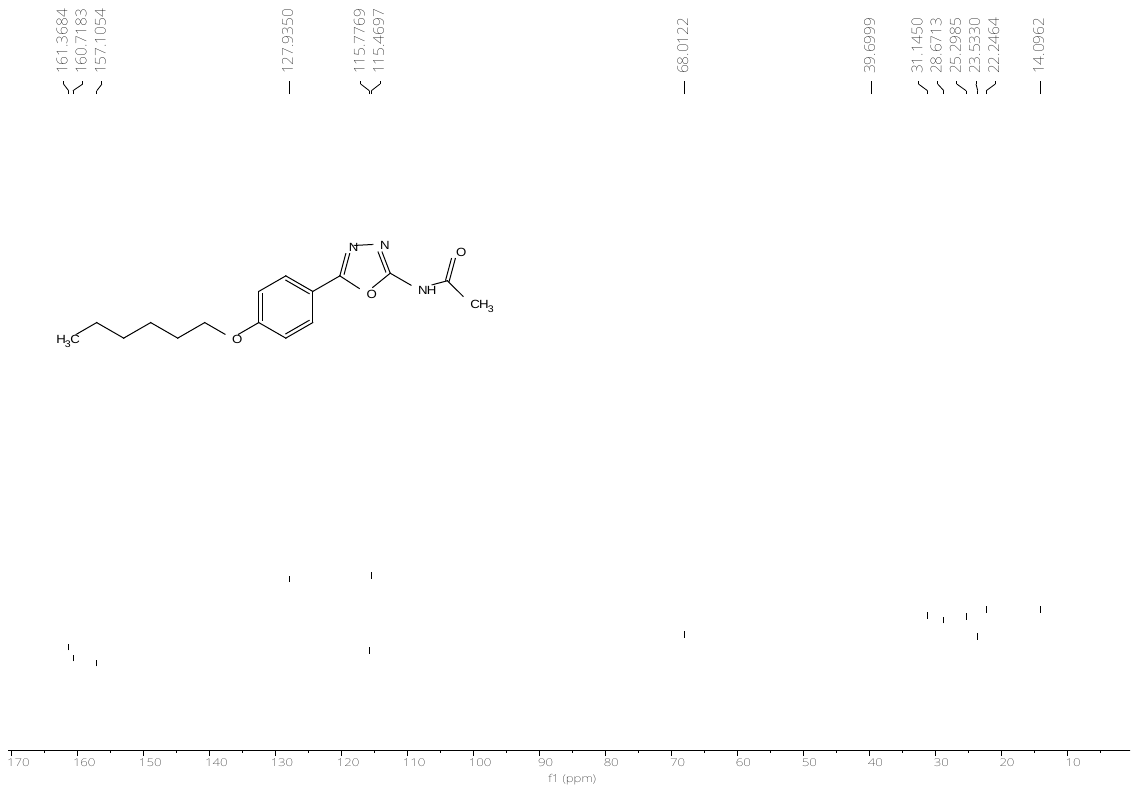

^1^H-NMR spectrum of *N-(5-(4-(n-hexoxy)phenyl)-1,3,4-thiadiazol-2-yl)acetamide (****3b****)*

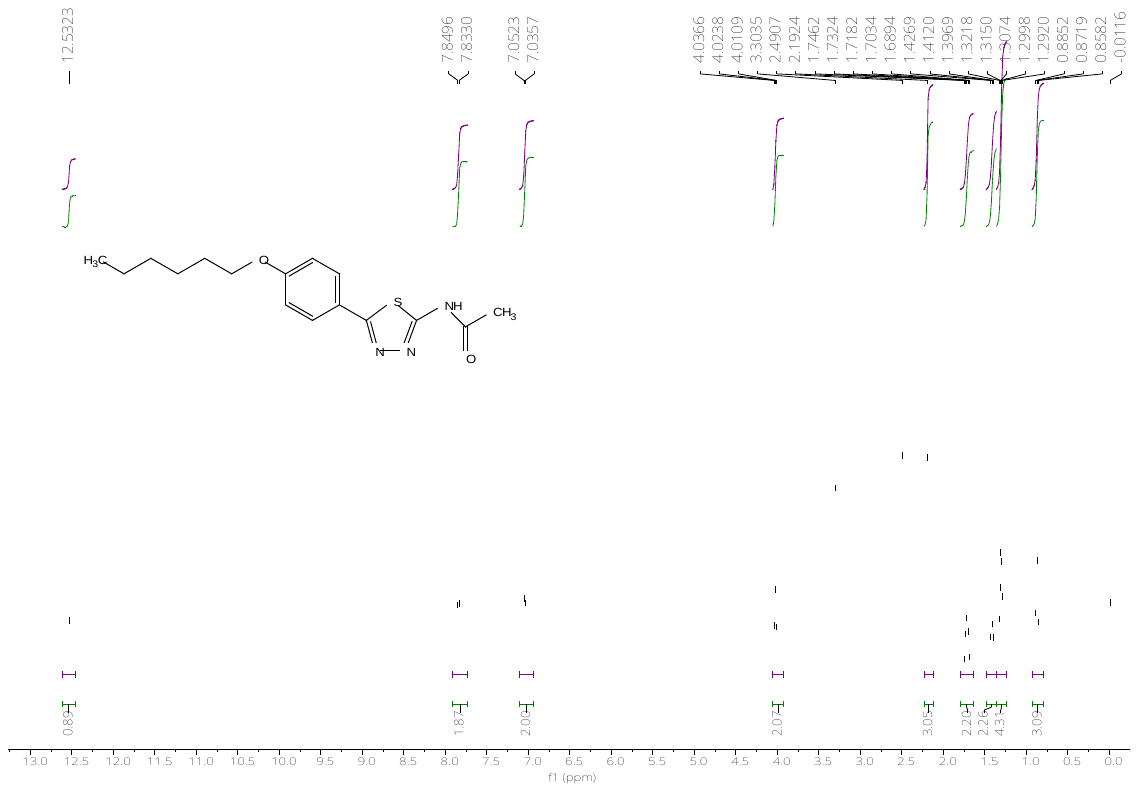

^13^C-NMR spectrum of *N-(5-(4-(n-hexoxy)phenyl)-1,3,4-thiadiazol-2-yl)acetamide (****3b****)*

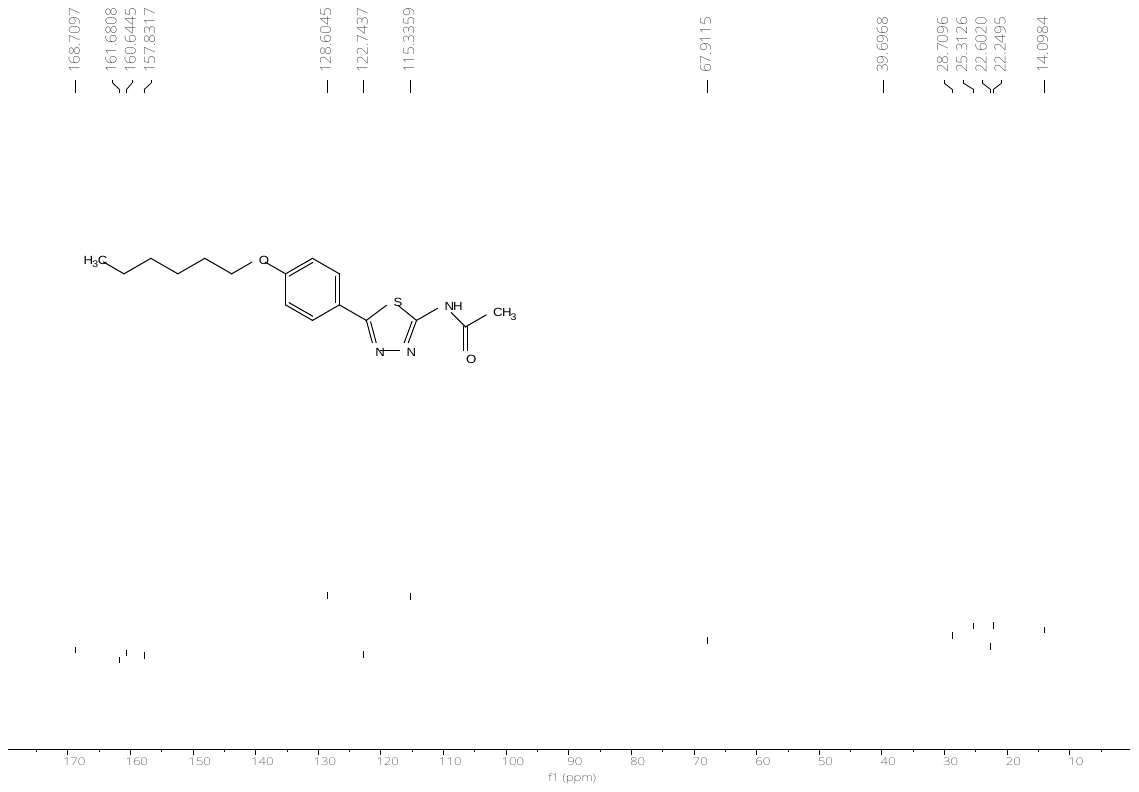

^1^H-NMR spectrum of *N-(2-(4-(n-hexoxy)benzoyl)-1-methylhydrazine-1-carbonothioyl)acetamide (****3c****)*

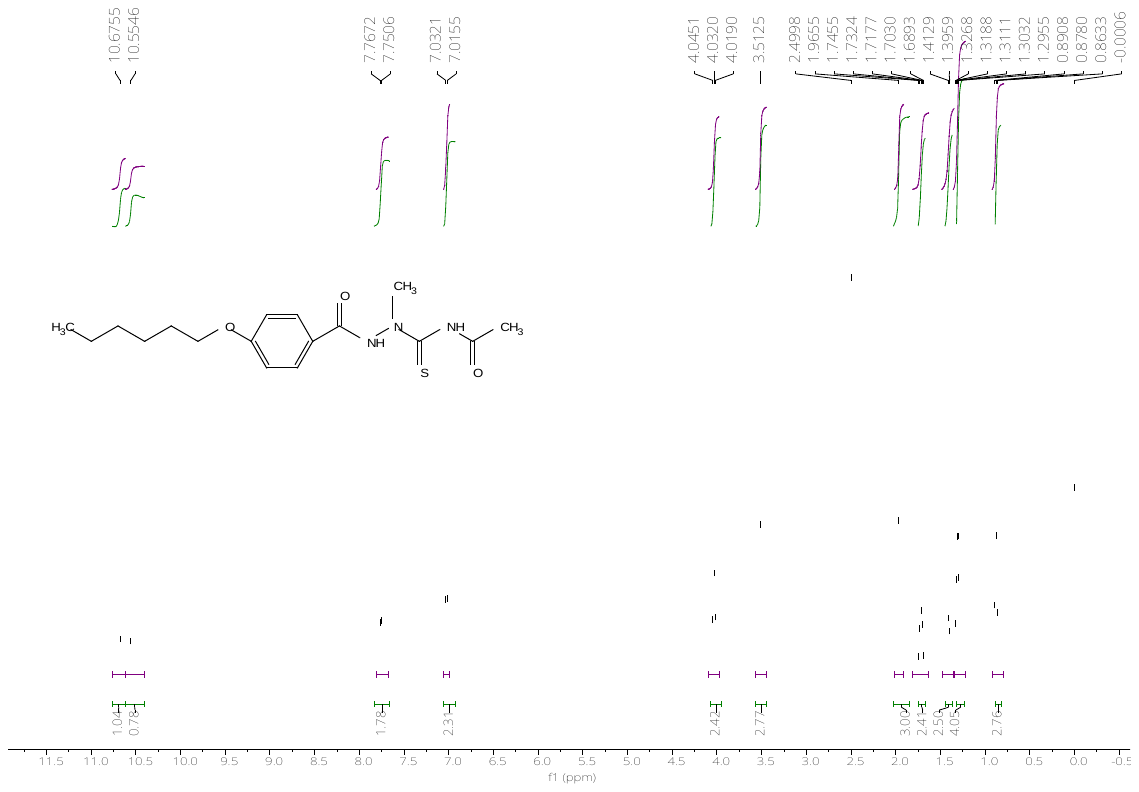

^13^C-NMR spectrum of *N-(2-(4-(n-hexoxy)benzoyl)-1-methylhydrazine-1-carbonothioyl)acetamide (****3c****)*

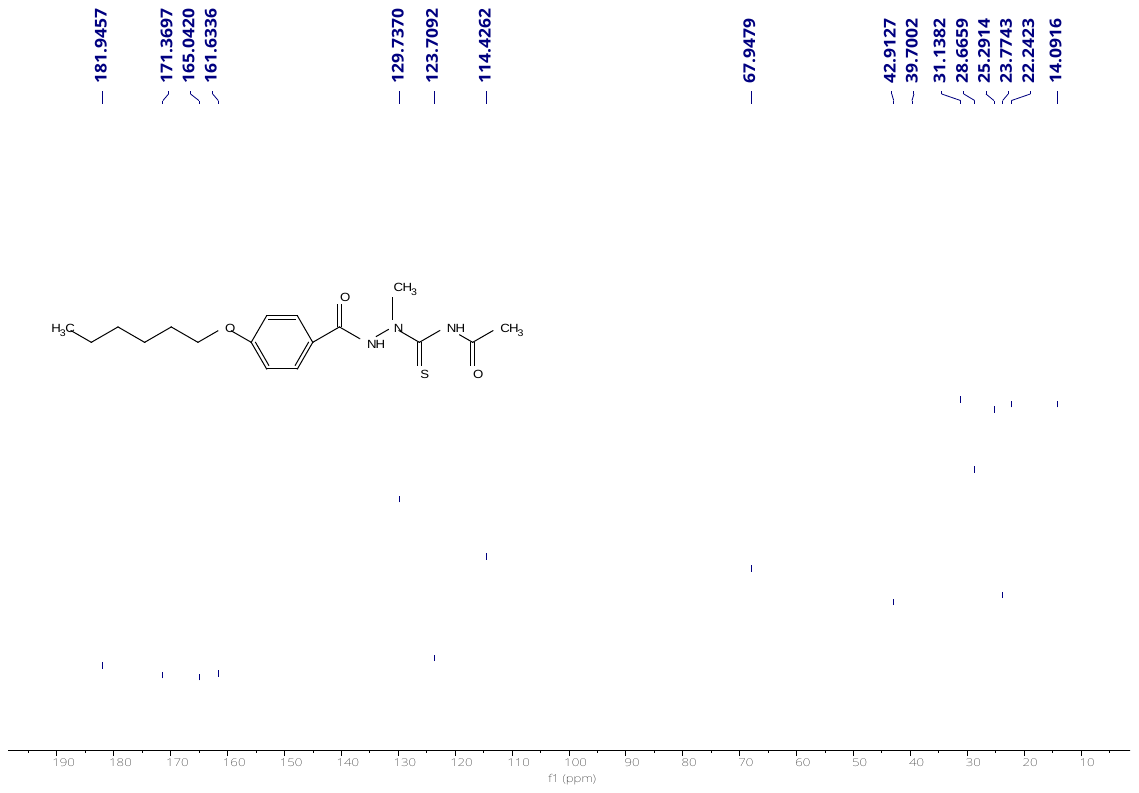

^1^H-NMR spectrum of *N-(2-(4-(n-hexoxy)benzoyl)-2-methylhydrazine-1-carbonothioyl)acetamide (****3d****)*

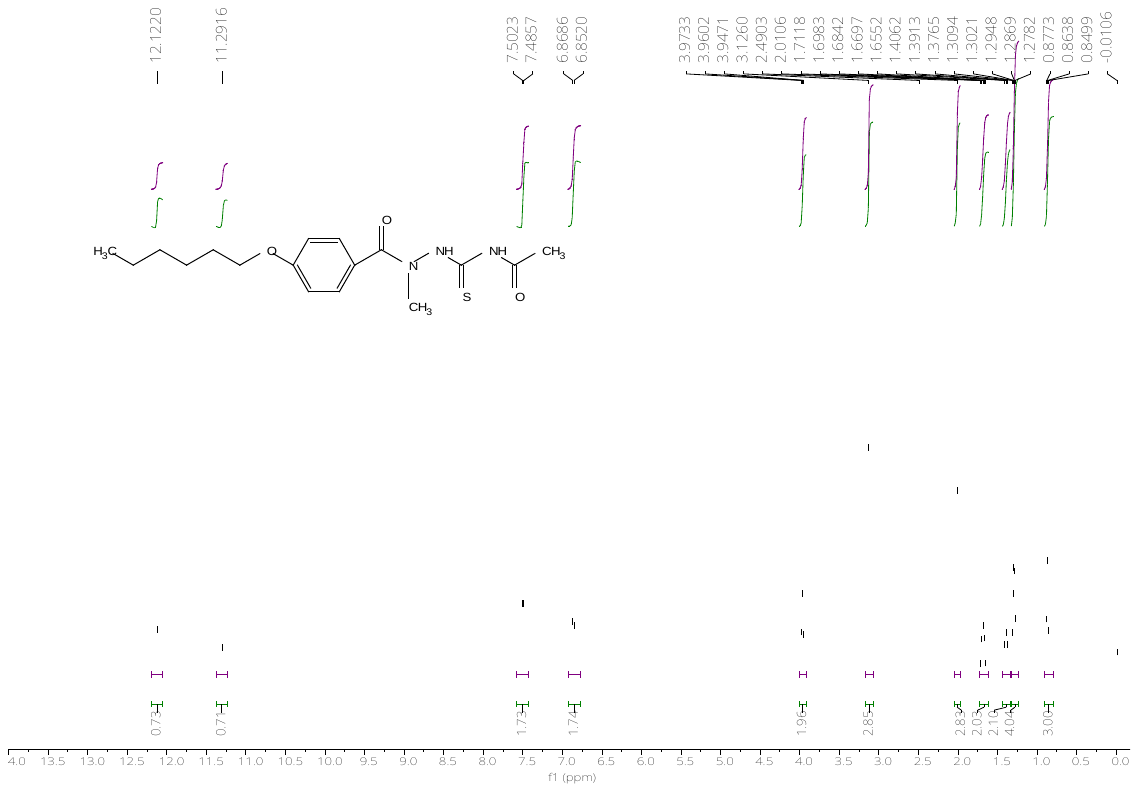

^13^C-NMR spectrum of *N-(2-(4-(n-hexoxy)benzoyl)-2-methylhydrazine-1-carbonothioyl)acetamide (****3d****)*

^1^H-NMR spectrum of *N-ethyl-2-(4-(n-hexoxy)benzyl)hydrazine-1-carbothioamide (****3e****)*

^13^C-NMR spectrum of *N-ethyl-2-(4-(n-hexoxy)benzyl)hydrazine-1-carbothioamide (****3e****)*

^1^H-NMR spectrum of *N-ethyl-2-(4-(n-hexoxy)benzoyl)hydrazine-1-carbothioamide (****3f****)*

^13^C-NMR spectrum of *N-ethyl-2-(4-(n-hexoxy)benzoyl)hydrazine-1-carbothioamide (****3f****)*

^1^H-NMR spectrum of *N-acetyl-2-(4-(n-hexoxy)benzoyl)hydrazine-1-carboxamide (****3g****)*

^13^C-NMR spectrum of *N-acetyl-2-(4-(n-hexoxy)benzoyl)hydrazine-1-carboxamide (****3g****)*

^1^H-NMR spectrum of *N-(methylsulfonyl)-2-(4-(n-pentoxy)benzoyl)hydrazine-1-carbothioamide (****3h****)*

^13^C-NMR spectrum of *N-(methylsulfonyl)-2-(4-(n-pentoxy)benzoyl)hydrazine-1-carbothioamide (****3h****)*

^1^H-NMR spectrum of *N-(2-acetylhydrazine-1-carbonothioyl)-4-(n-pentoxy)benzamide (****3i****)*

^13^C-NMR spectrum of *N-(2-acetylhydrazine-1-carbonothioyl)-4-(n-pentoxy)benzamide (****3i****)*

^1^H-NMR spectrum of *2-(4-(n-hexoxy)benzoyl)hydrazine-1-carbothioamide (****3j****)*

^13^C-NMR spectrum of *2-(4-(n-hexoxy)benzoyl)hydrazine-1-carbothioamide (****3j****)*

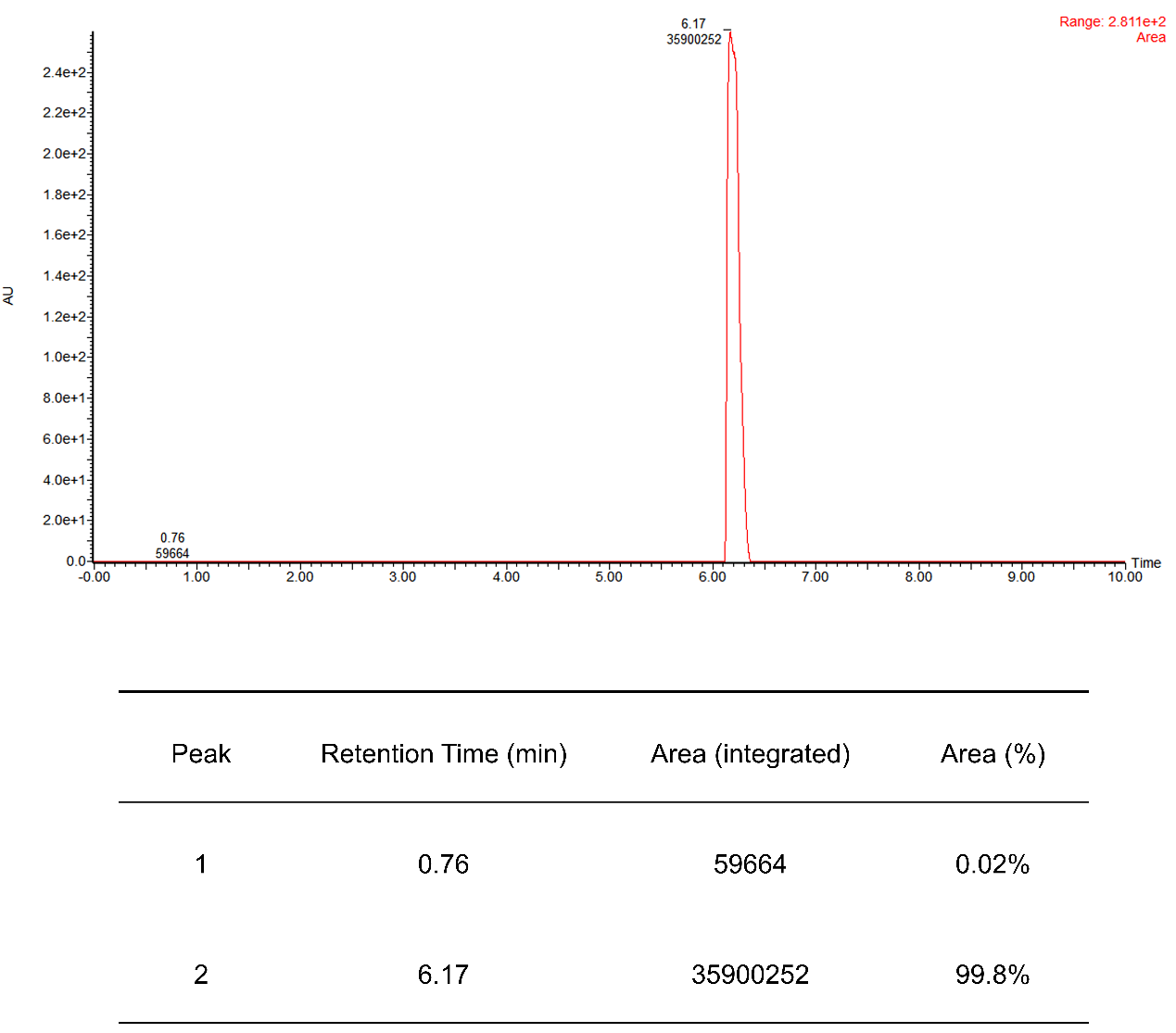

### **Fig. S7**. HPLC purity data of compound **1b**.
